# Genetic architecture and gene mapping of cyanide in cassava (*Manihot esculenta Crantz*.)

**DOI:** 10.1101/2020.06.19.159160

**Authors:** Alex C Ogbonna, Luciano Rogerio Braatz de Andrade, Ismail Y. Rabbi, Lukas A. Mueller, Eder Jorge de Oliveira, Guillaume J. Bauchet

## Abstract

Cassava is a root crop originating from South America and a major staple crop in the Tropics, including marginal environments. In this study, we focused on South American and African cassava germplasm and investigated the genetic architecture of Hydrogen Cyanide (HCN), a major component of tuber quality. HCN is a plant defense component against herbivory but also toxic for human consumption. We genotyped 3,354 landraces and modern breeding lines originating from 26 Brazilian states and 1,389 individuals were phenotypically characterized across multi-year trials for HCN. All plant material was subjected to high density genotyping using Genotyping-by-sequencing (GBS). We performed genome wide association mapping (GWAS) to characterize the genetic architecture and gene mapping of HCN. Field experiment revealed strong broad and narrow-sense trait heritability (0.82 and 0.41 respectively). Two major loci were identified, encoding for an ATPase and a MATE protein and contributing up to 7% and 30% of the cyanide concentration in roots, respectively. We developed diagnostic markers for breeding applications, validated trait architecture consistency in African germplasm and investigated further evidence for domestication of sweet and bitter cassava. Fine genomic loci characterization indicate; (i) a major role played by vacuolar transporter in regulating HCN content, (ii) co-domestication of sweet and bitter cassava major alleles to be geographical zone dependant, and (ii) major loci allele for high cyanide cassava in *Manihot esculenta Crantz* seems to originate from its ancestor, *M. esculenta* ssp. *flabellifolia.* Taken together these findings expand insights on cyanide in cassava and its glycosylated derivatives in plants.

**One-sentence summary:** Identification of an intracellular transporter gene and its allelic variation allow to point out cultivars with up to 30 percent decrease in cassava root cyanide content, toxic for human consumption.

## Introduction

Cassava (*Manihot esculenta Crantz.*) is a starchy root crop widely grown throughout the tropics (Southeast Asia, Latin America, the Caribbean and sub-Saharan Africa) for human and livestock consumption, and as feedstock for biofuels and other bio-based materials (Howeler, Lutaladio, and Thomas 2013). Mostly cultivated by low-income, smallholder farmers, cassava is a staple food crop for over 800 million people worldwide. Cassava is an efficient crop in marginal areas where poor soils and unpredictable rainfall dominates (FAO 2018). Cassava has developed defense mechanisms against herbivores and pathogens, including the biosynthesis of cyanogenic glucosides (CG) (Takos et al. 2011). However, some of the major challenges in cassava includes low tuber protein and carotenoid content as well as high content of linamarin and lotaustralin CG (K. Jørgensen et al. 2005; Blomstedt et al. 2012; Gleadow and Møller 2014). CG, characterized as α-hydroxynitriles, are secondary metabolites of plant products derived from amino acids (Gleadow and Møller 2014). Cyanogenesis is the release of toxic hydrogen cyanide in cassava upon tissue disruption. Its concentration is usually higher in young plants, when nitrogen is in ready supply, or when growth is constrained by nonoptimal growth conditions (Gleadow and Møller 2014).

Cultivars with cyanide content of < 100 mg kg^−1^ fresh weight (FW) are called ‘sweet cassava’ while cultivars with 100-500 mg kg^-1^ are ‘bitter cassava’ (Wheatley et al., 1993). In Brazil, cassava’s center of diversity, the difference in preference of bitter and sweet cassava appears to be in its role in subsistence in regions they dominate. Areas where sweet cassava dominates, it is a component of a diet; whereas areas where bitter cassava dominates, it is the main carbohydrate source, generally complemented by a protein, such as a fish (Mühlen et al. 2019).

Cyanide in cassava is synthesized in the leaf and transported to the roots via the phloem (Jørgensen, Nour-Eldin, and Halkier 2015). The most abundant CG is linamarin (>85%), and total CG concentration varies according to the cultivar, environmental conditions, cultural practices and plant age (McMahon, White, and Sayre 1995). Degradation of linamarin is catalyzed by the enzyme linamarase, which is found in cassava tissues, including intact roots. The compartmentalization of linamarase in cell walls and linamarin in cell vacuoles prevents the accidental formation of free cyanide. Disruption of tissues ensures that the enzyme comes into contact with its substrate, resulting in rapid production of free cyanide via an unstable cyanohydrin intermediary (Wheatley, Chuzel, and Zakhia 2003). Therefore, careful processing is required to remove Hydrogen Cyanide (HCN), especially in communities with poor nutritional status (Jørgensen et al. 2005; Blomstedt et al. 2012; Gleadow and Møller 2014). Incomplete processing could result in acute to chronic exposure to HCN (Leavesley et al. 2008). High dietary cyanogen consumption from insufficiently processed roots of bitter cassava combined with a protein-deficient diet leads to a neglected disease known as konzo (Kashala-Abotnes et al. 2019). Konzo is a distinct neurological disease characterized by an abrupt onset of an irreversible, non-progressive limbs paralysis (Tshala-Katumbay et al. 2001; Nzwalo and Cliff 2011; Kashala-Abotnes et al. 2019). Juice extraction, heating, fermentation, drying or a combination of these processing treatments aid in reducing the HCN concentration to safe levels (Wheatley, Chuzel, and Zakhia 2003). Gleadow and Moller (2014), reported efforts in cassava breeding programmes to actively select for varieties with lower levels of HCN. However some farmers favour cassava varieties with higher cyanide content as a source of resistance against herbivores and theft by humans (Lebot 2009). Modern breeding has not yet succeeded in developing cassava cultivars totally free of CG (Nweke, Lynam, and Spencer 2002; K. Jørgensen et al. 2005). Previous studies (Kizito et al. 2007; Whankaew et al. 2011) on cyanide using Quantitative Trait Locus (QTL) approach, could not provide a conclusive information on the genetic basis for cyanide variation in cassava, owing to the available genomic resources and narrow dataset background used so far.

In this study, we seek to (1) comprehensively understand the genetic architecture of HCN in cassava, (2) map gene(s) associated to CG variation, (3) develop a fast, cost effective molecular diagnostic toolkit for breeding purposes to increase selection efficiency, and (4) investigate evidences of HCN domestication.

## Results

### Large scaled analysis of Brazilian population for HCN content

Phenotypic distribution and variation for cyanide content was measured in a Brazilian population of 1,246 individuals using picrate titration method, on which a scale of 1 to 9 indicates the concentration of HCN content (1 and 9 representing extremes of low and high HCN, respectively) (M. G. Bradbury, Egan, and Howard Bradbury 1999). Based on a determined scale, the cyanide concentration varies from 2 to 9 with an average of 5.6 with individuals coming from across Brazilian states **(Figure 1A-B)**. About two-thirds of the total plots of 28,203 had missing values, with 9,139 plots having HCN observations (**Supplementary Table 1; Supplementary Table 2**). Broad-sense heritability (*H*^2^) was calculated to 0.82 for cyanide content, similar to previous observations reported on several species (Barnett and Caviness 1968; Goodger, Ades, and Woodrow 2004; Gleadow and Møller 2014). Using genotyping data previously recorded for this population (Ogbonna et al., in press), we observed a genotype variance (V_G_) higher than genotype-by-year variance (V_GxY_), with their ratio (V_GxY_/V_G_) showing an interaction of 0.29. HCN deregressed best linear unbiased prediction (BLUP) shows a very high correlation with non-deregressed BLUP with Pearson’s correlation coefficient of 0.99, indicating a balanced replication of individuals among the studied population. See deregressed BLUPs (**Supplementary Table 3)**.

**Figure. 1.**
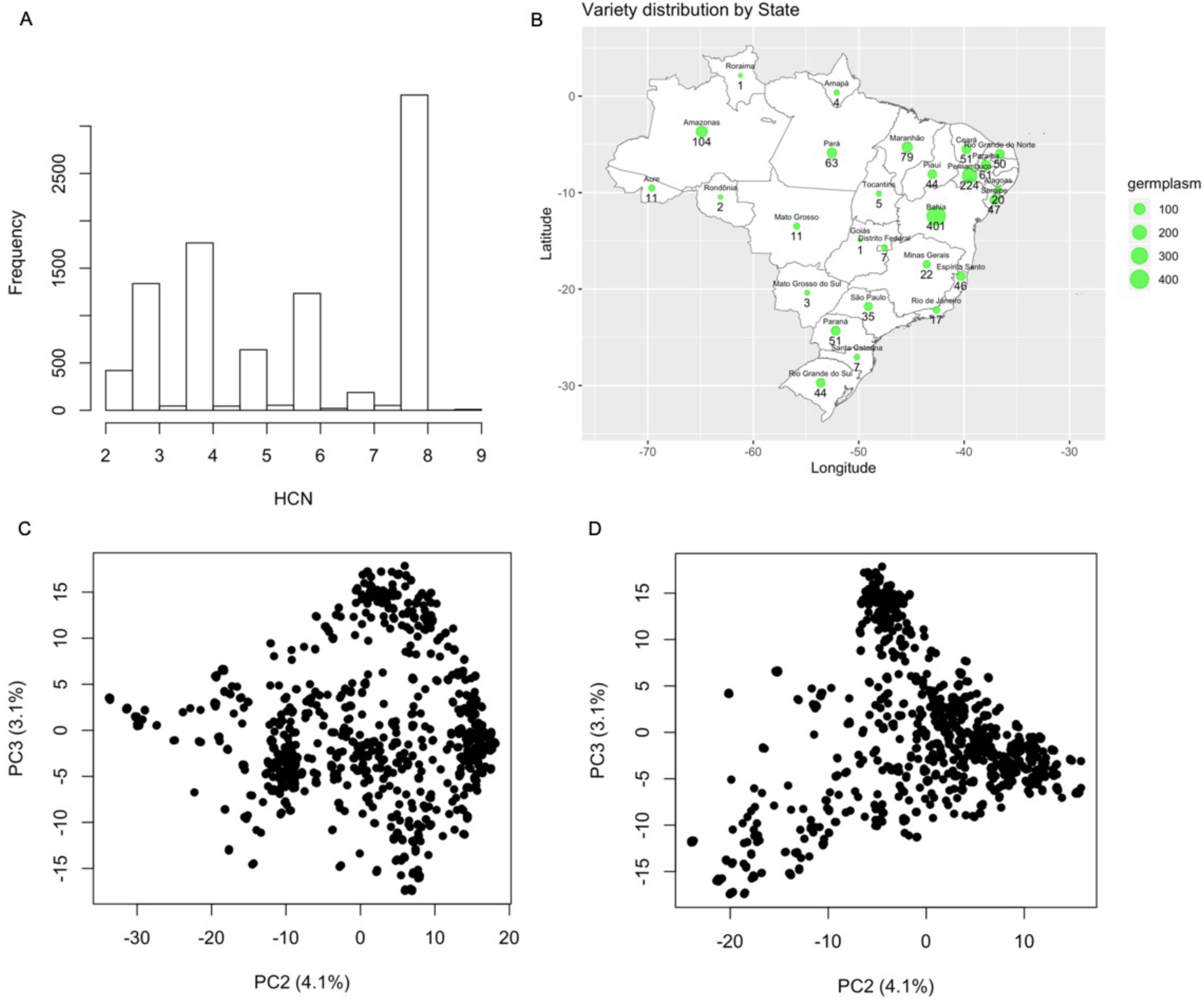
Geographic distribution and population structure of Latin American (Brazilian) germplasm. **(A)** Distribution of assayed HCN. HCN assayed phenotype score ranges from 2 to 9. **(B)** Distribution of germplasm based Brazilian states. A total of 1,821 cassava accessions have valid geographic information. The green dots show where the accessions come from while the size of the dots represent the number of clones from that state. The black numbers show how many accessions were sampled from each location. Population structure reveals the first three axes of the principal component analysis (PCA) explains about 15.3% of the variations in the population of 1246 individuals, 9,686 SNPs after filtering for Hardy–Weinberg equilibrium and LD of 0.01 and 0.2 respectively. **(C)** shows the first and second axis, while **(D)** shows the first and the third axis.

### GWAS analysis revealed two SNPs associated with HCN accumulation

Single Nucleotide Polymorphisms (SNPs) calling using Tassel version 5, identified a total of 343,707 variants of which 30,279 were selected for phasing and imputation. After imputation, a total of 27,045 bi-allelic SNPs with an allelic correlation of 0.8 or above were kept for downstream analysis. The first three Principal Components (PCs) accounted for over 15.3% genetic variation (**Figure 1C-D; Supplementary Note1**).

To identify genetic correlation between HCN content and genotypic variation, mixed model Genome Wide Association Mapping (GWAS) was performed using GCTA software (Yang et al. 2011) with bonferroni correction as a test of significant SNPs. After the Bonferroni correction (-log10(0.05/27045) threshold, 5.733117), two significant peaks were identified on chromosomes 14 and 16 with 45 and 12 significant associated markers, respectively **(Figure 2A, Supplementary Table 4)**. Subsequent regional linkage disequilibrium (LD) analysis on chromosome 16 gives a 3.6 Mb interval and local LD analysis gives a 248 Kb interval (with *r*^2^ threshold of > 0.8) on which 6 genes are annotated **(Table 1, Supplementary Figure 1A)**. The optimal strongest *p-value* indicates the SNP S16_773999 (p-value: 7.53E-22) located within the Manes.16G007900 gene. Manes.16G007900 is annotated as a Multidrug and Toxic Compound Extrusion or Multi-Antimicrobial Extrusion protein (MATE). MATE transporters are a universal gene family of membrane effluxers present in all life kingdoms. MATE transporters have been implicated directly or indirectly in mechanisms of detoxification of noxious compounds and able to transport CG (Darbani et al. 2016). Interestingly, S16_773999 SNP is predicted to induce a missense variant (A>G) in exon 4 **(Figure 2B – red star-gene model)**. This mutation causes an amino acid change from Threonine to Alanine, predicted as deleterious. A second MATE gene (Manes.16G008000) located at 22Kb from the candidate MATE gene (**Figure 2B – annotation panel**) also shows a high LD (pairwise correlation of 0.96; **Supplementary Figure 2A)**. The second MATE gene could be suggested as a paralog of our candidate gene following reported cassava genome duplication (Bredeson et al. 2016).

**Table 1.**
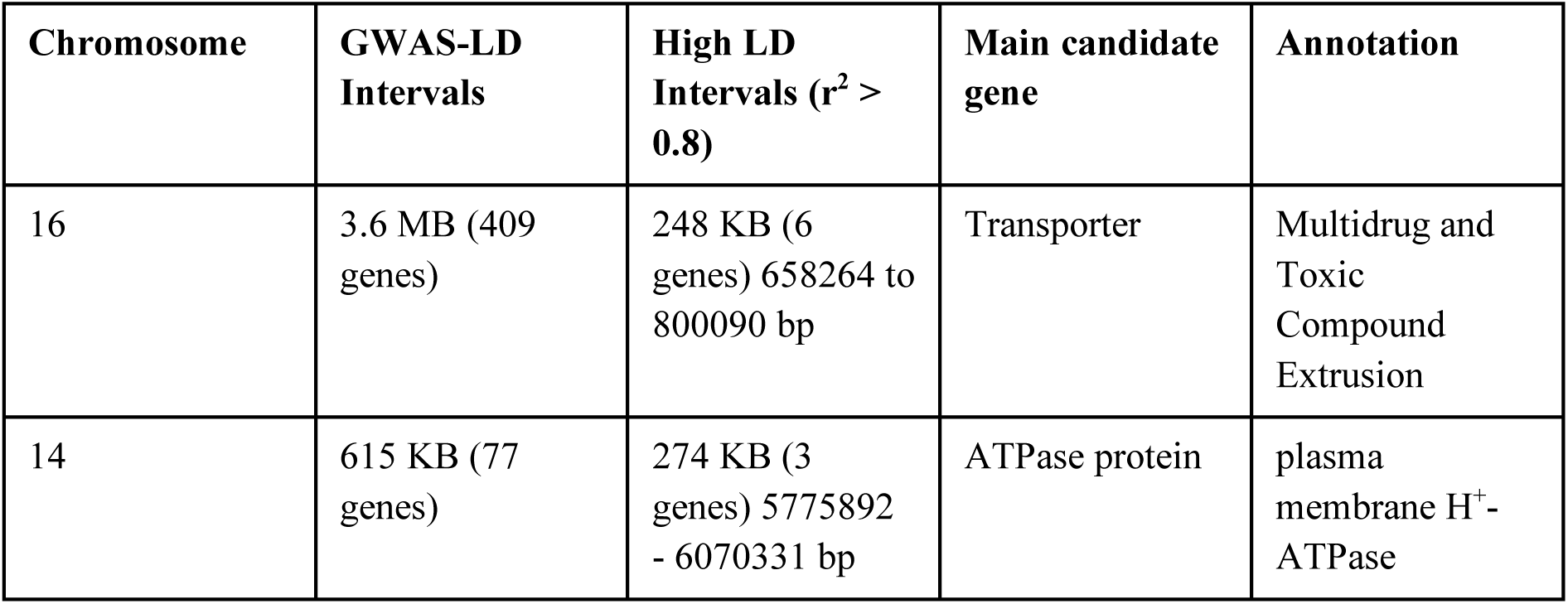
Summary of linkage disequilibrium analysis within the regions (chromosomes 14 and 16) associated with HCN variation in cassava for Brazilian germplasm.

| Chromosome | GWAS-LD Intervals | High LD Intervals ($r^2 > 0.8$) | Main candidate gene | Annotation |
| --- | --- | --- | --- | --- |
| 16 | 3.6 MB (409 genes) | 248 KB (6 genes) 658264 to 800090 bp | Transporter | Multidrug and Toxic Compound Extrusion |
| 14 | 615 KB (77 genes) | 274 KB (3 genes) 5775892 - 6070331 bp | ATPase protein | plasma membrane H<sup>+</sup>-ATPase |

**Figure 2.**
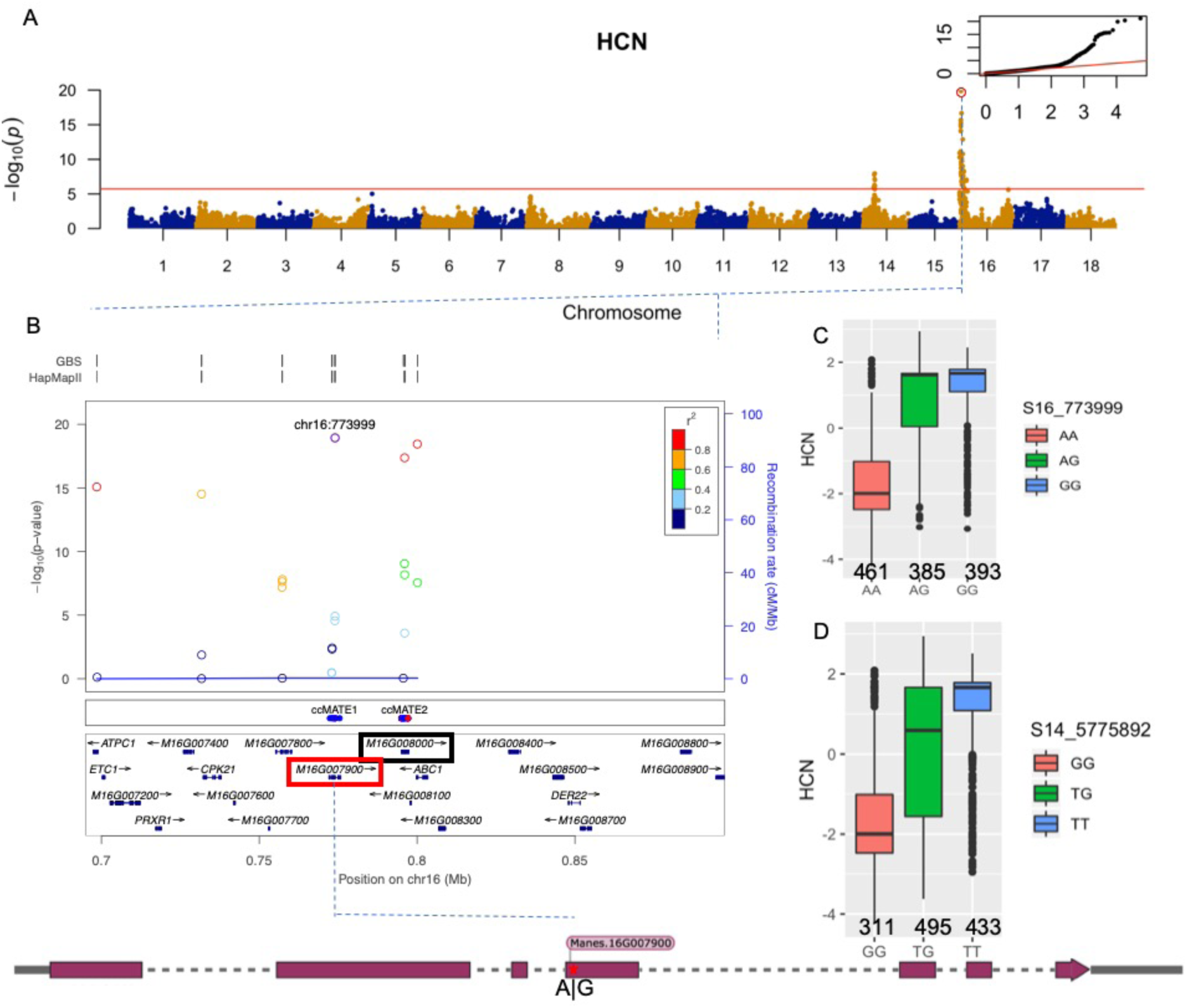
GWAS of HCN for LA germplasm. **(A)** Manhattan plot from a mixed linear model (MLM-LOCO) with the chromosome on which the candidate SNP is located, excluded from calculating the genetic relationship matrix (GRM). Bonferroni significance threshold is shown in red. A quantile-quantile plot is inserted to demonstrate the observed and expected −log10 of P-value for HCN. The red circle indicates the candidate SNP. **(B)** LocusZoomplot shows the HCN chromosome 16 associated region (-log_10_ *P* value) around the candidate gene. Top two lines show genomic coverage at the locus, with each vertical tick representing directly genotyped from GBS and HapMap SNPs. Each circle represents a SNP, with the color of the circle indicating the correlation between that SNP and the candidate SNP at the locus (purple). Light blue lines indicate estimated recombination rate (hot spots) in GBS. The middle panel shows 36 single point mutations (red are deleterious) between the region spanning ccMATE1 and ccMATE2. Bottom panel shows genes at each locus as annotated in the cassava genome version 6.1. The red and black rectangle indicate Manes.16G007900 and Manes.16G008000 gene respectively, with Pearson correlation coefficient of 0.96 (r^2^) between both genes. The lower figure is the gene model with the position indicated of the associated SNP within the 4th exon. (**C** and **D**) Boxplot showing candidate SNP effect for HCN between each genotype class at the top markers, S14_6050078 and S16_773999, respectively.

The second peak in chromosome 14 shows an association with log *p-values* of 1.08e-08, gives an interval of 615 Kb and the local LD analysis reduced it to 274 Kb on which 3 genes are located **(Table 1, Supplementary Figure 1B)**. The first candidate SNP indicates that S14_6050078 (*p-value* 1.08e-08) is located in Manes.14G074300, a gene coding for an integral membrane HPP family protein involved in nitrite transport activity (Maeda et al. 2014). In a recent study, Obata and colleagues (2020), highlighted that linamarin, an abundant variant of CG in cassava, contains nitrogen and serves as a nitrogen storage compound (Obata et al. 2020) as previously hypothesized (Siritunga and Sayre 2004). This confirms previous observations that application of nitrate fertilizer to cassava plants increases cyanide accumulation in the shoot apex (K. Jørgensen et al. 2005). The second candidate SNP indicates that S14_6021712 (*p-value* 7.32E-08) is located in Manes.14G073900.1, coding for a plasma membrane H(+)-ATPase. H+-ATPase mediated H+influx associated with Al-induced citrate efflux coupled with a MATE co-transport system (Zhang et al. 2017). Wu et al. (2014) found that transgenic *Arabidopsis* lines containing *Brassica oleracea* MATE gene had stronger citrate exudation coupled with a higher H+efflux activity than wild-type plants (Wu et al. 2014).

As a validation step, we used a subset of 523 unique individuals (from the core Panel of 1,536 unique individuals [Ogbonna et al., in press]), with phenotypic and genotypic information to perform GWAS. Results **(Figure 4-LA Unique; Supplementary Table 5)** revealed the same loci (as was observed in the larger dataset of 1,246 individuals) associated with HCN variation in our initial GWAS dataset, indicating that the core unique panel represents the overall genetic variation for HCN in the Brazilian germplasm collection. However, it detected less significant loci (only 46%) than those detected using a dataset of 1,246 individuals. This indicates that additional small effects QTL were captured with the larger dataset conferring increased statistical power.

**Figure 3.**
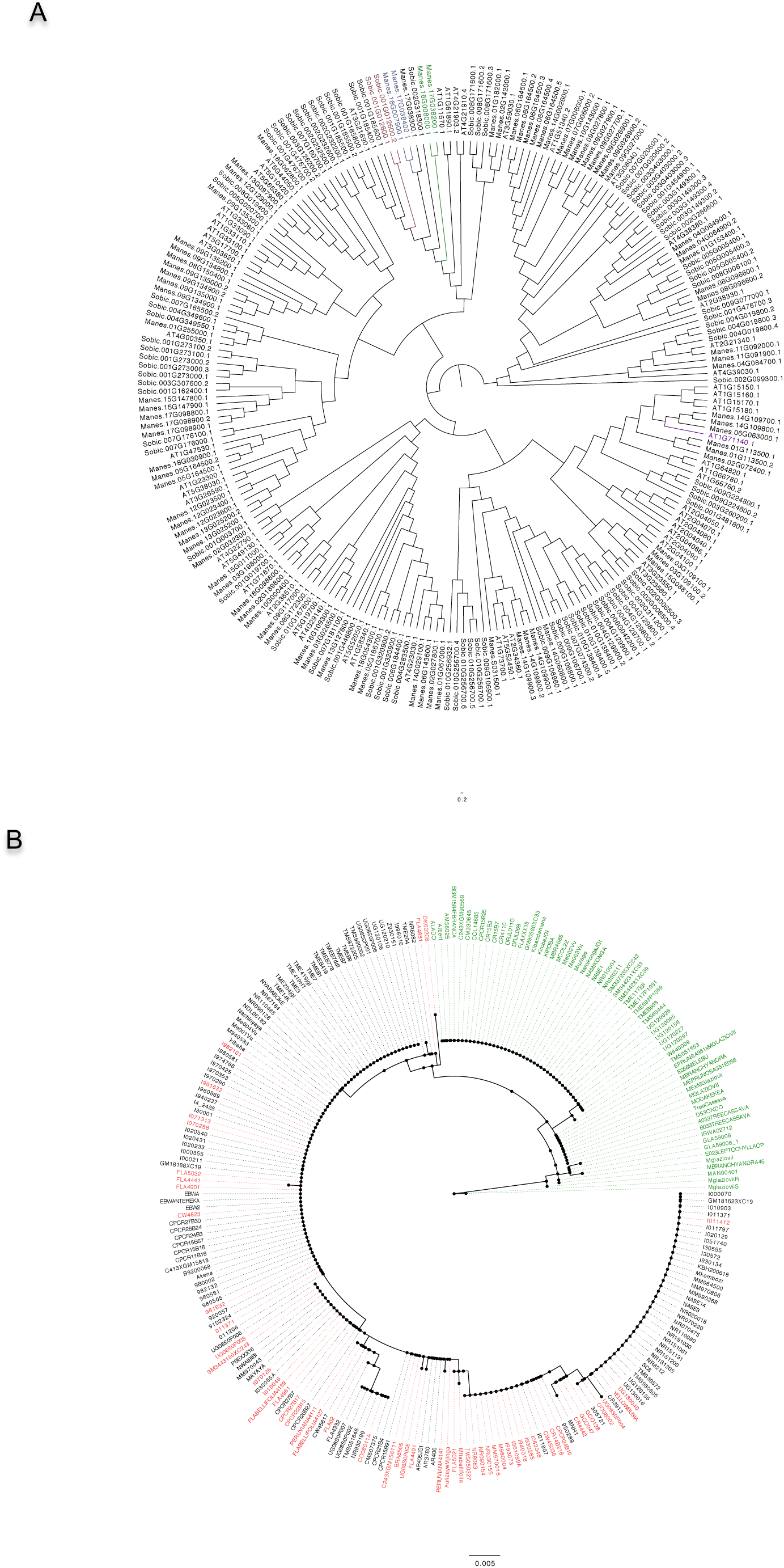
Phylogeny analysis. **(A)** Protein sequences alignment of MATE genes in cassava, sorghum and Arabidopsis. Protein Alignment and comparative phylogeny show a close sequence homology between the GWAS candidate gene and SbMATE2 (Sobic.001G012600), a characterized vacuolar membrane MATE transporter in Sorghum, functions in the accumulation of plant specialized metabolites such as flavonoids and alkaloids. **(B)** Genomic sequence of Manes.16G007900 for the 242 HapMap accessions. Accessions highlighted in red are homozygous GG for SNP16_773999, identified as having high HCN content. Accession highlighted in green are homozygous AA for SNP16_773999, identified as low HCN content. Accessions in black are heterozygotes AG or GA for SNP16_773999.

**Figure 4.**
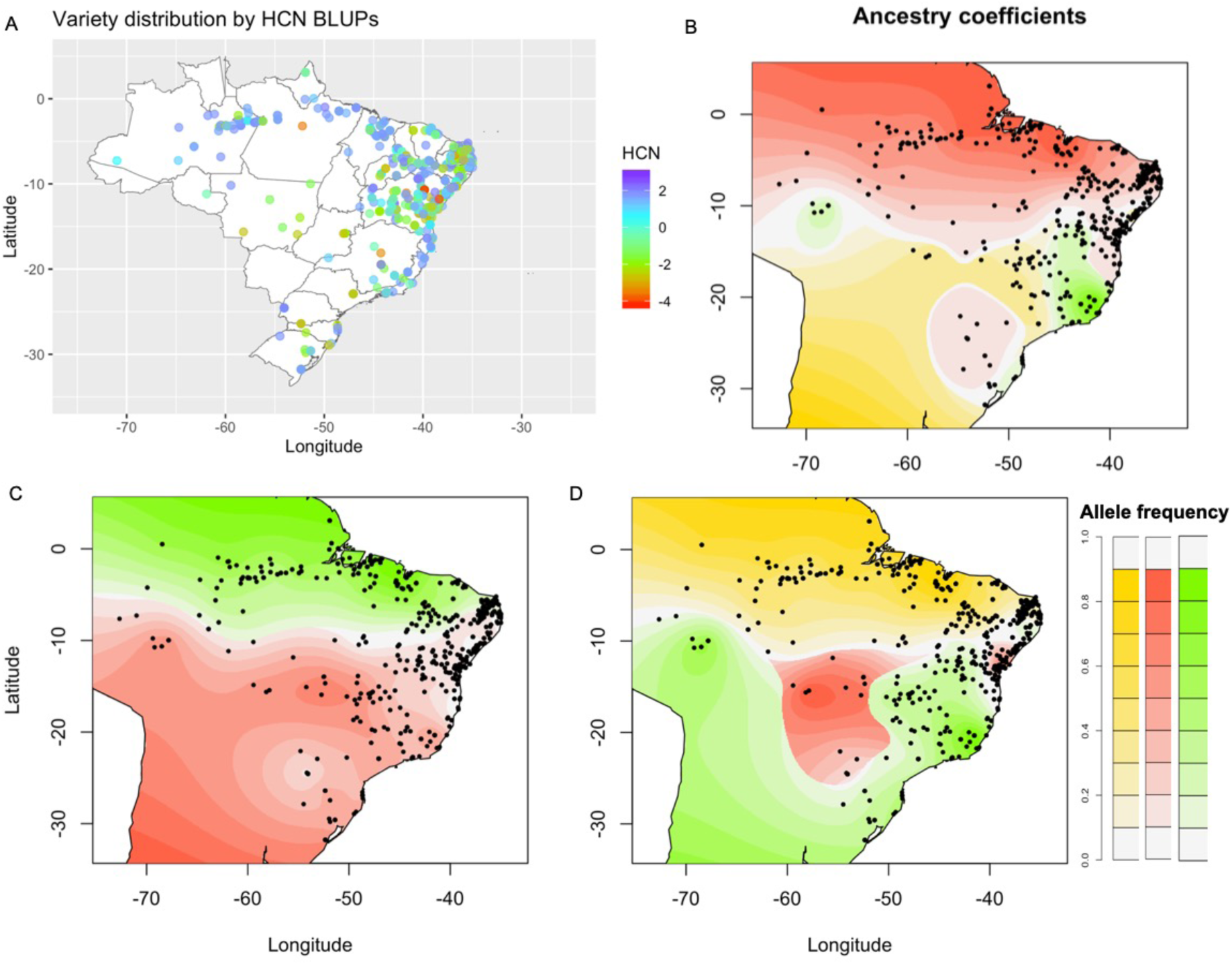
Spatial distribution of ancestral coefficients for HCN candidate SNPs using 1,657 accessions. **(A)** Distribution of germplasm based on best linear unbiased prediction (BLUP) of HCN. Accessions with high cyanide contributing alleles are grouped around the Amazonas and low cyanide contributing alleles are grouped in other areas of Brazil. **(B)** Spatial distribution of allele frequency for HCN candidate loci in chromosome 14. **(C)** Spatial distribution of allele frequency for HCN candidate loci in chromosome 16. **(D)** Interactions of HCN candidate loci in chromosome 14 and 16.

The alleles driving high cyanide at S16_773999 and S14_6050078 loci show dominance and additive patterns, respectively (**Figure 2C-D**); homozygotes with alternate alleles for both loci show higher cyanide content than heterozygotes, while homozygotes with reference alleles show lower cyanide. This indicates that cyanogenic cassava can be alternate allele homozygous or heterozygous at these loci, while acyanogenic cassava plants are more likely reference allele homozygous at these loci. Joint allelic substitution effects at the associated loci for cyanide did not show any interaction between the two loci as shown in **Supplementary Figure 1C**.

### Variance explained and evidence for Domestication in HCN reveals chromosome 16 as a good candidate for KASP marker development

To calculate narrow-sense heritability, the proportion of variance explained was calculated using parametric mixed model multiple kernel approach (Akdemir and Jannink 2015). Single kernel mixed model explained 0.41 of the marker based (narrow-sense heritability, *h*^2^) proportion of the variance for HCN across the genome. A multi kernel mixed model with the top significant SNPs in chromosome 16 and 14 (S16_773999 and S14_5775892) as the first and second kernel with the rest of the genome as the third kernel, explained 30%, 7% and 63% of the marker based variance respectively. A three kernel mixed model to determine the variance explained by chromosome 14, 16 and the rest of the genome, showed that the proportion of variance explained by the three kernels are 16%, 50% and 34% respectively. Chromosome 14 and 16 tag SNPs for the candidate SNPs explains 8% and 36% proportions of variance, respectively; while the rest of the genome explains 56%. We found evidence for local interactions within chromosome 16 which is most likely as a result of high LD around the region (**Supplementary Method1 for M&M**).

To validate the local interaction found in chromosome 16, we performed intra-chromosomal epistasis interaction using FaST-LMM (Lippert et al. 2011, 2013). Chromosome 16 revealed 242 significant interactions above the bonferroni corrected threshold (-log10(0.05/1131*(1131-1)/2); 1.6024), with three interactions clearly separated by 1 Mb between each pair of SNPs (**Supplementary Figure 1D; Table 2, Supplementary Table 9**). A biosynthetic gene cluster in cassava (Genome draft version Cassava4.1) was earlier identified by Andersen et al. (2000), which we identified to be on chromosome 12 in genome version 6.1 as shown in **supplementary figure 3A-B**. Inter-chromosomal epistasis interactions analysis involving about 400 million tests, did not reveal any significant interactions, neither for bonferroni or FDR threshold. Over 27 million tests had *p-values* less than 0.05 significant level (**Supplementary Method2 for M&M**).

**Table 2.** Chromosome 16 interactions SNP Pairs for Separated (1 Mb apart) Epistasis Interactions in Chromosome 16. SNP 1 and SNP 2 Showed Strong Significant Interactions. The table also contains Single-Locus P-value for the interaction SNPs (SNP).

| SNP 1 |  | SNP 2 |  | Interactions |
| --- | --- | --- | --- | --- |
| SNP | Single-Locus p Value | SNP | Single-Locus p Value | Interaction p Value |
| S16_12540 | 1.00E-05 | S16_1063230 | 4.31E-15 | 5.207858e-10 |
| S16_1298874 | 2.01E-07 | S16_12540 | 1.00E-05 | 2.969058e-09 |
| S16_1298876 | 7.72E-08 | S16_12540 | 1.00E-05 | 8.371752e-10 |

**Table 3.**
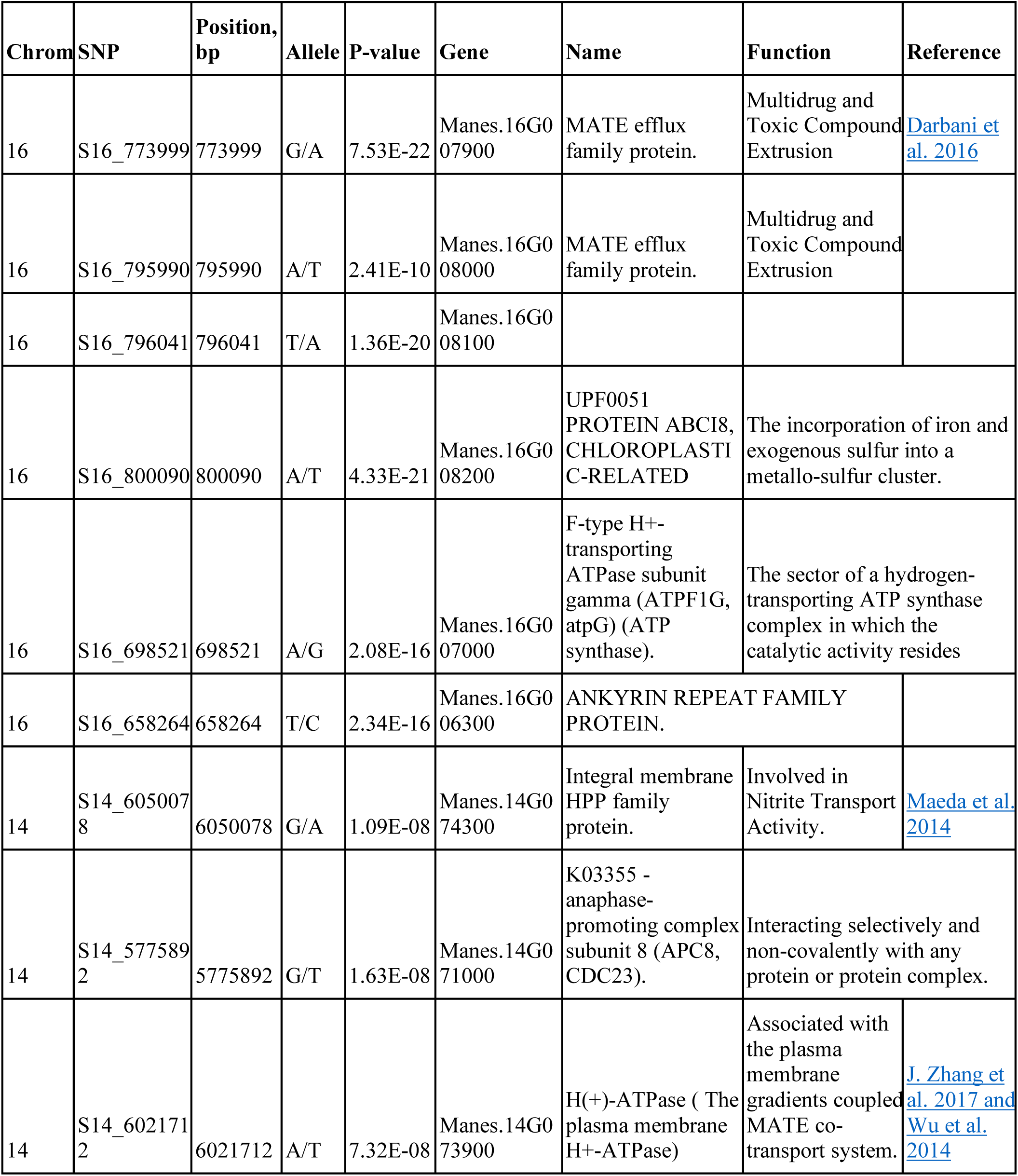
Summary of genes within the regions associated with HCN variation in Brazilian cassava.

Investigating evidence for domestication in HCN, we carried out differentiating loci analysis using cassava HapMap reference lines (Ramu et al. 2017) for cultivated *M. esculenta* and wild *M. esculenta* ssp. *flabellifolia* (**Supplementary Table 6**). We identified 294 biallelic ancestry-informative single-nucleotide markers that represent fixed or nearly fixed differences between cultivated and wild accessions (**Supplementary Figure 4)**. Interestingly, we observed high fixed loci (89) differentiating between the two groups in chromosome 16, of which over 54 of them are approximately 0.37 Mb away from the candidate MATE gene for HCN regulation (**Supplementary Figure 4).** Together these results indicate that (i) epistasis is observed within chromosome 16 around the main GWAS peak (**Supplementary Figure 1D**) and (ii) the identified epistatic region colocalizes with differentiating loci between *M. esculenta* and wild *M. esculenta* ssp. *flabellifolia* (**Supplementary Table 6**; **Supplementary Method3 for M&M**).

The Kompetitive Allele-Specific PCR assay (KASP) is a robust, high throughput and cost efficiency PCR based marker technology (He, Holme, and Anthony 2014; Neelam, Brown-Guedira, and Huang 2013). We used KASP to develop and validate diagnostic markers for HCN content, based on association peaks, local linkage disequilibrium and allelic effect. Candidate SNPs from the GWAS were subjected to KASP marker design (**Supplementary Table 7**) and assayed on Embrapa Breeding populations for a total of 576 individuals. The average percentage genotype score or call rate was 96.59% with a maximum of 97.92% and a minimum of 92.71% validated across population and validate allelic segregation for HCN content (**Supplementary Table 8, Supplementary Method4 for M&M**).

### Phylogenetics and mutation predictions reveal altered function of MATE transporter

To identify homologues of the MATE transporter Manes.16G007900, a protein alignment and comparative phylogeny analysis were performed for genome-wide MATEs in cassava, sorghum and *Arabidopsis* using CLUSTAL OMEGA (Sievers et al. 2011). Results showed a close sequence homology between three additional MATE transporters in the cassava genome: Manes.16G007900, Manes.17G038400, Manes.17G038300 and Manes.16G00800 with percentage identity of 91.09%, 78.05%, 68.59%, respectively. Highest interspecific homology analysis found SbMATE2 from sorghum (Sobic.001G012600; percentage identity of 67.84% [first isoform] and 71.00% [second isoform]) (Darbani et al. 2016) and AtMATE from *Arabidopsis* (AT3G21690; percentage identity of 72.80%) (Liu et al. 2009), characterized as vacuolar membrane transporters **(Figure 3A, Supplementary Note2 on Phylogeny Tree)**. Manes.16G007900 and Manes.16G00800 predicted topology of 12 transmembrane helices supporting their annotation (**Supplementary Figure 5A-B**), as previously reported for *Arabidopsis (Li et al. 2002), sorghum (Darbani et al. 2016)* and blueberry (Chen et al. 2015) (**Supplementary Method5 for M&M**). Maximum likelihood tree using protein sequences from 241 HapMap individuals displayed distinct clade distribution of 64 homozygote individuals for SNP S16_773999 G:G allele (high Cyanide) in color red and 114 homozygotes for SNP S16_773999 A:A allele (low Cyanide) in green color. *M. esculenta* ssp. *flabellifolia* individuals (homozygote G:G) and other wild accessions *M.glaziovii and M.pruinosa* (homozygote A:A) clustered in distinct clades (**Figure 3B**).

The stability of a protein to denaturation is calculated by measuring changes in free energy, and the higher and more positive the change in the free energy is, the more stable the protein is against denaturation (Quan, Lv, and Zhang 2016). We mined 36 single point mutation predictions in GBS and whole genome resequencing data (Ramu et al. 2017) for Manes.16G007900 and Manes.16G008000 proteins. In the observed 36 single point mutations across the two proteins, this value ranges from 0.26 to −4.00 with an average of −1.57 (**Supplementary Figure 5C(1-4), Method6, Table 10**). The deleterious point mutations showed higher negative values in their structural change prediction. Mutations with sensitive stability changes can affect the motion and fluctuation of the target residues. All 36 point mutations except one (**Figure 2B, middle panel**), had a negative change in free energy, indicating loss of stability conferring fluctuations in the protein function (**Supplementary Method6 for M&M**).

### Sweet and Bitter cassava Geographical Distribution

We represented the geographical distribution and HCN content of Brazilian germplasm, recently characterized (Ogbonna et al, in press) and presented a contrasted distribution (Figure 4A). Accessions with high cyanide contributing alleles are grouped mostly around the Amazonas and low cyanide contributing alleles are grouped in other areas of Brazil. Specifically, individuals with high cyanide are mostly found around the Amazonian rivers and the coastal areas while more variation in HCN content was observed in other regions of Brazil. The ancestry coefficient distribution for S16_773999, S14_5775892 and joint haplotypes (S16_773999 and S14_5775892) revealed 3 different ancestry coefficients for the candidate SNP S14_5775892 (Figure 4B) following an additive response (Figure 2C). Two different ancestry coefficients were observed for the candidate SNP S16_773999 (Figure 4C) following the complete dominant response observed (Figure 2D). Pseudohaplotype of candidate SNPs in chromosome 14 and 16 shows the distribution of 3 ancestry coefficients (Figure 4D), indicating low, intermediate and high cyanide ancestry coefficients (**Supplementary Method7 for M&M and Note3 for Discussion**).

Leveraging from open source data (available at cassavabase.org, see **Supplementary Method8 for M&M**), we explored the distribution of cyanide across sub-Saharan Africa datasets, including assayed individuals originating from 26 countries (**Supplementary Table 11, Supplementary Figure 6A**) and field trials carried out in different locations across Nigeria. This analysis indicated that Central and Southern Africa showed on average higher cyanide varieties compared to West Africa (**Supplementary Figure 6B**), while a trend of lowering cyanide was detected on landraces compared to improved varieties **(Supplementary Figure 6C)**.

### Validating GWAS results in African and Joint Africa, Latin America population

Phenotypic distribution and variation for cyanide content was measured in African population of 636 individuals using the picrate titration method. Cyanide concentration varies from 1 to 9 with an average of 5.1 in the African population (**Supplementary Table 12**). *H*^2^ and *h*^2^ for cyanide content were 0.27 and 0.26 respectively, less than observed in Brazilian germplasm (**Supplementary Table 2**). The genotype variance (*V*_*g*_) was higher than genotype-by-environment variance (*G*_*gxe*_) with their ratio (*V*_*gxe*_ / *V*_*g*_) showing a high interaction of 0.86. The estimated deregressed BLUPs ranged from 0.0009 to 2.5638 with an average of 0.5242 (**Supplementary Table 13**). After the Bonferroni correction (-log10(0.05/53547) threshold, 6.029765), two significant peaks were identified on chromosomes 14 and 16 respectively **(Figure 5 AF-panel; Supplementary Table 14)**. A third peak was observed in chromosome 11 but was not up to the significant threshold. GWAS dataset for HCN in African accessions showed peaks on chromosome 14 and 16 with SNP S14_6612442 and SNP S16_1298874 showing the highest *p*-values, congruent with the Brazilian GWAS dataset.

**Figure 5.**
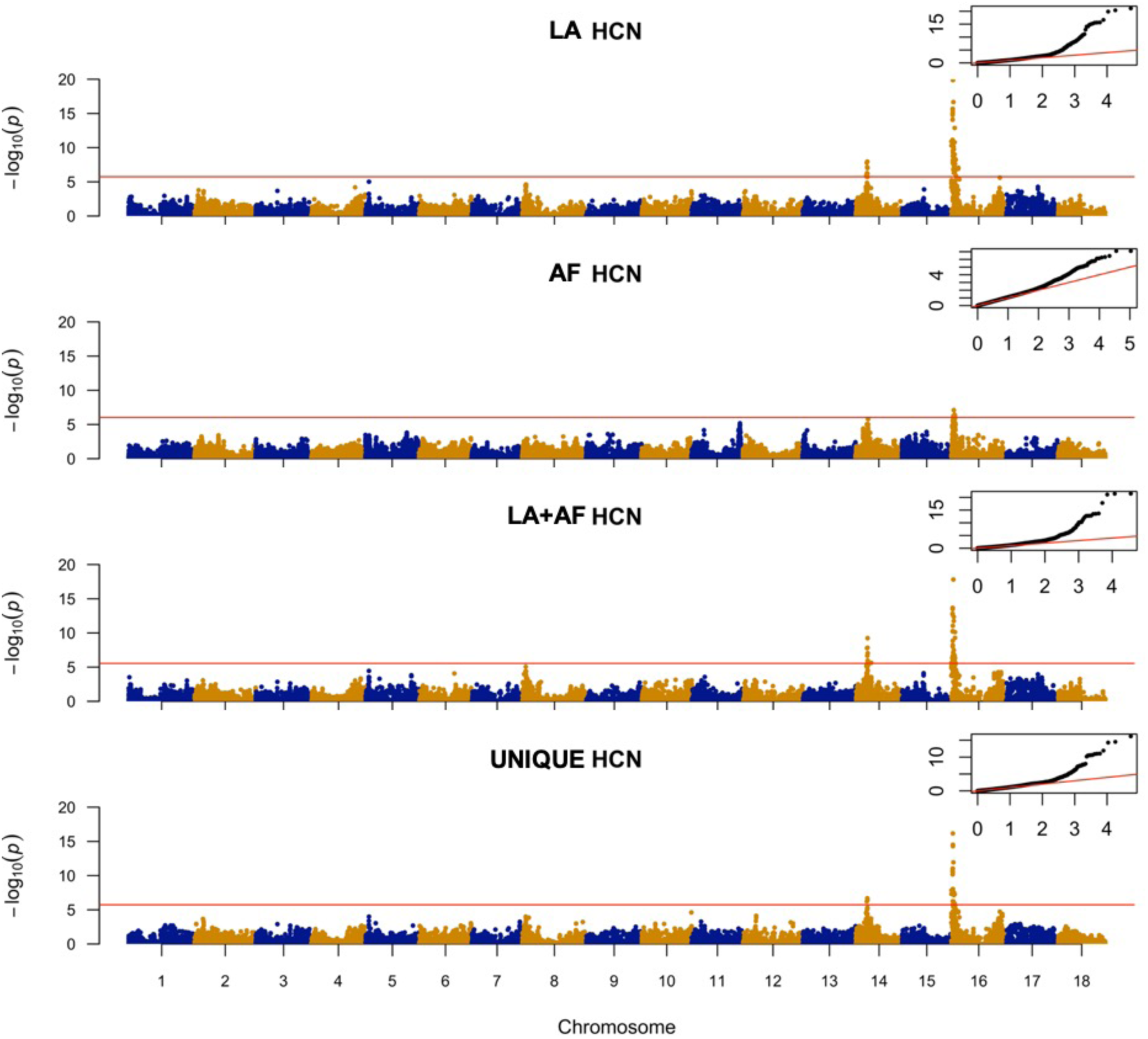
Manhattan plot from a mixed linear model (MLM-LOCO) with the chromosome on which the candidate SNP is located, excluded from calculating the genetic relationship matrix (GRM). The MLM-LOCO summarizes the genome-wide association results for HCN in **Latin American (LA, Brazilian), African (AF), joint Latin American + African (LA+AF)** and **Latin American unique (LA UNIQUE)** germplasms. Bonferroni significance threshold is shown in red. A quantile-quantile plot is inserted to demonstrate the observed and expected −log10 of P-value for HCN.

For Africa and Latin America combined analysis, phenotypic variation ranged between 1 to 9 with an average of 5.2 (**Supplementary Table 15**). *H*^2^ and *h*^2^ heritability for cyanide content in Africa and Brazilian combined analysis was 0.50 and 0.38 respectively. The genotype variance (*V*_*g*_) was higher than genotype-by-environment variance (*G*_*gxe*_) with a ratio (*V*_*gxe*_ / *V*_*g*_) showing a lower interaction of 0.42 compared to that of African population alone (**Supplementary Table 2**). The estimated deregressed BLUPs (for the 1,875 individuals used in GWAS) ranged from 0.0027 to 4.2266 with an average of 1.2545 (**Supplementary Table 16**). After the Bonferroni correction, two significant peaks were identified on chromosomes 14 and 16 respectively, corresponding to the earlier reported candidate SNPs **(Figure 5 LA+AF-panel; Supplementary Table 17)**. A whole genome imputation of the African-Brazilian dataset using the hapmap as a reference panel for chromosome 16 **(Supplementary Figure 7A)**, further validate Manes.16G007900 and the associated SNP S16_773999, based on optimal *p*-value (4.74E-22) (**Supplementary Table 18**; **Supplementary Method8 for M&M)**. See distributions of phenotypes and deregressed BLUPs (**Supplementary Figure 8).**

We requested available open source RNA-sequencing dataset on the molecular identities for 11 cassava tissue/organ types using the TMEB204 (TME204) cassava variety to evaluate gene expression (Wilson et al. 2017). Both Manes.16G007900 and Manes.16G008000 showed differential expression between storage and fibrous root with p-values of 5.00E-05 and 0.00065, respectively (**Figure 6A-B**). Manes.16G007900 is differentially expressed between fibrous root and leaf with FPKM values of 13.9219 and 89.5362 respectively, whereas Manes.16G008000 is not and shows low expression level (**Figure 6A-B**). Selective sweeps detection using HapMap WGS between cassava progenitors, Latin American and African accessions do not show sweeps overlapping with candidate and biosynthetic regions (**Supplementary Figure 9&10)**.

**Figure 6.**
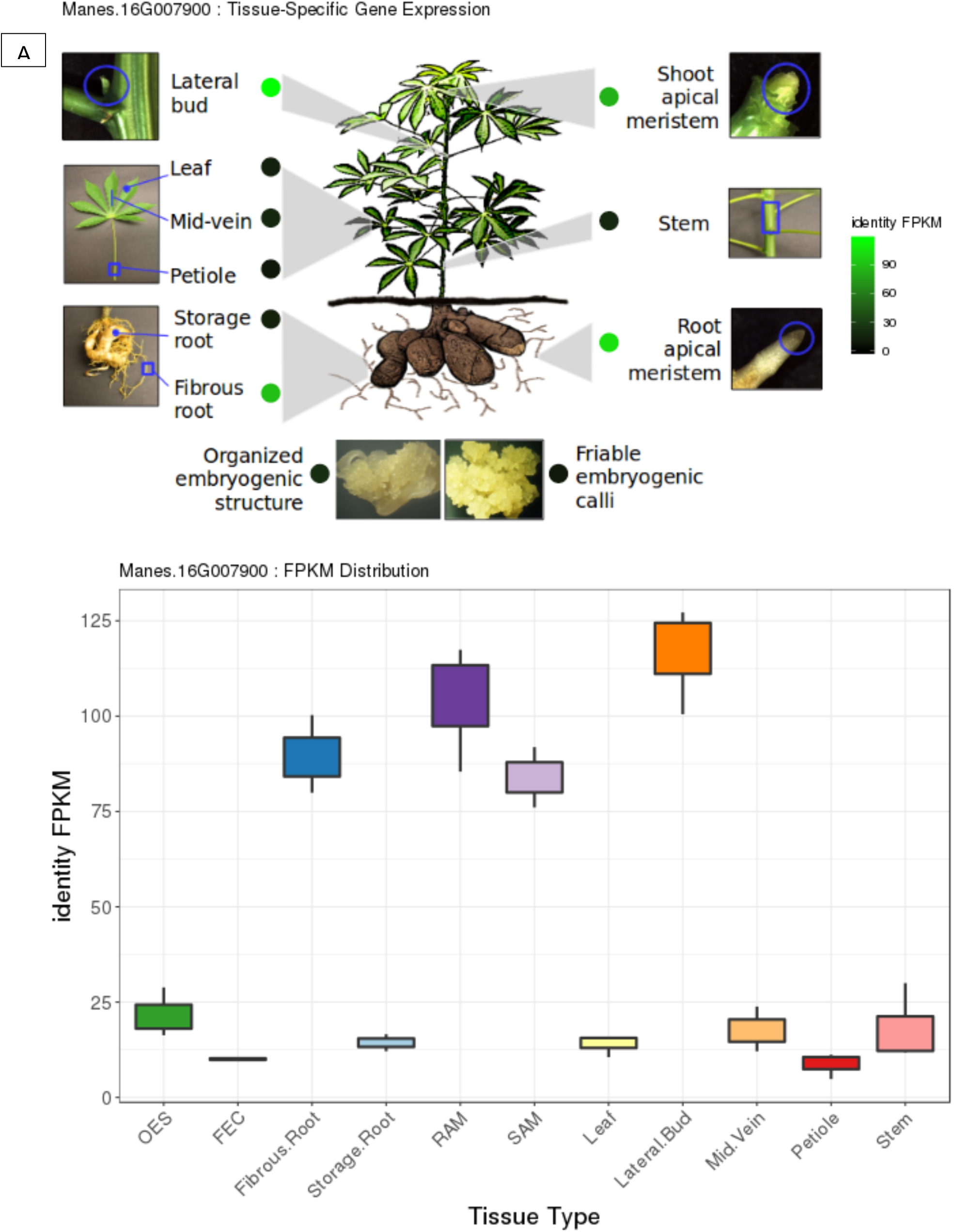
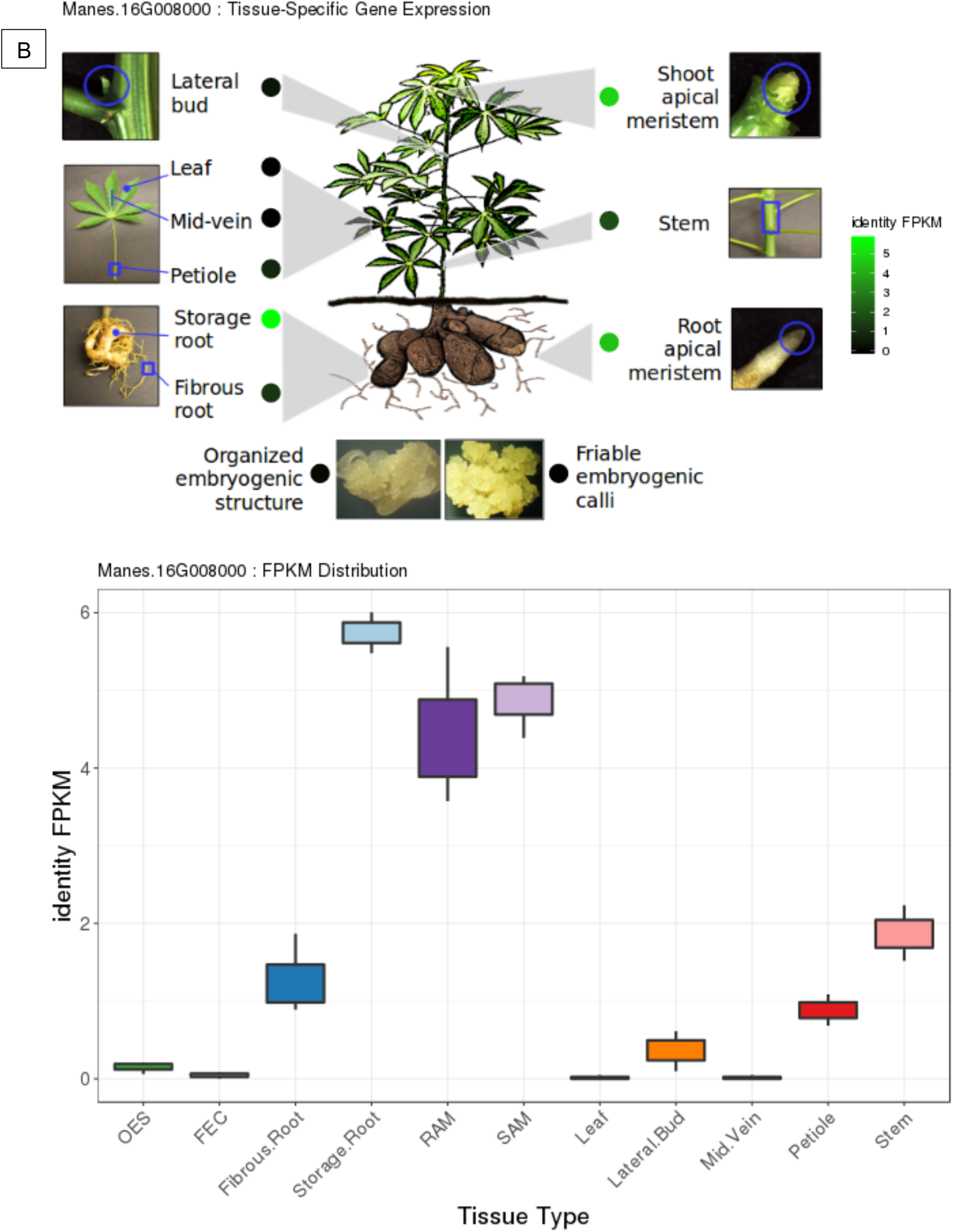
**(A)** Manes.16G007900 and (B) Manes.16G008000 issues/organs expression profiles, three month after planting (FPKM) of African cassava accession TMEB204 (*Manihot esculenta*) sampled for gene expression (Wilson et al. 2017). TMEB204, an African variety, was assayed for HCN in a 1997 field experiment carried out at IITA Mokwa location (Nigeria) and forms part of the individuals in our African dataset with average cyanide content of 5.67 (min=5, max=7). TMEB204 allelic profile for candidate SNP S16_773999 on chromosome 16 is heterozygous, indicating dominance of Manes.16G007900 high cyanide alleles.

## Discussion

The potential of CG content in cassava varieties varies, even among the roots of the same plant (Gleadow and Møller 2014). These variations are partly due to genetics, environmental conditions and soil type (Bokanga et al. 1994; K. Jørgensen et al. 2005; Nzwalo and Cliff 2011). While germplasm from Latin America shows higher genetic variance and heritability (Brazil, *V*_*g*_=2.59, *H*^2^=0.82, *h*^2^=0.41; Colombia, *V*_*g*_=1.58, *H*^2^=0.69), their African counterparts showed much less genetic variance and heritability (*V*_*g*_=0.21, *H*^2^=0.27, *h*^2^=0.26). Latin America/Brazil is the primary center of domestication from where cassava was solely introduced in the XVI century into Africa, which could explain a probable genetic bottleneck and the observed difference between the two populations (Bredeson et al. 2016). In addition, sweet and bitter cassava landraces are differentiated in Latin America but not in Africa. This is attributed to post-introduction hybridization between sweet and bitter cassava and the inconsistent transfer of ethnobotanical knowledge of use-category management to Africa (Bradbury et al. 2013). Mislabeling of germplasm in Africa (Yabe, Iwata, and Jannink 2018) may also have contributed to the observed difference. These differences were also observed with the distribution plots for the individuals assayed in our analysis for HCN in Latin American (bimodal distribution) and African (almost normally distributed) populations. The observed differences between broad and narrow-sense heritability estimates, attributed to missing heritability, could be explained by the local epistasis interactions involving few major genes explaining the variance for HCN in chromosome 16 (Akdemir and Jannink 2015), the large numbers of rare variants omitted through imputation (Yang et al. 2015) and the use of only bi-allelic subsets of filtered SNPs, leaving behind multi-allelic loci that may have explained additional variance.

Previous studies on the genetic architecture of HCN, found two QTL linked to loci SSRY105 and SSRY242 explaining 7 and 20% of the genetic variation in an S_1_ population (Kizito et al. 2007). Blasting the sequences of the loci, revealed SSRY105 location on chromosome 14 (57,582,253 bp) of cassava genome version 6.1 (http://phytozome.jgi.doe.gov) and was congruent with the region found on chromosome 14 associated with HCN variation in our current datasets. Whankaew et al. (2011), found five QTL (CN09R1, CN09R2, CN09L1, CN09L2, CN08R1) across two environments and years, but without any consistent QTL. Their corresponding locations on cassava genome version 6.1 were chromosomes 12 (CN09R1), 9 (CN09L1), 3 (CN09L2), and 4 (CN09R2). The sequence of SSRY242 and CN08R1 QTL could not be found to specify their locations in the genome. These studies could not provide a comprehensive information on the genetic basis for root cyanide variation in cassava given that; (1) cyanide content is affected by environment, (2) the use of population with distinct genetic background, (3) the stage of the field trials at which cyanide was assayed, and (4) the use of low marker density which limited the resolution and QTL detection power.

To provide comprehensive genetic architecture of HCN in cassava, we performed a GWAS using multi-year trials conducted in Brazil in 2016 through 2019 on individuals assayed for root cyanide using picrate titration method. Two major regions associated with HCN variation were identified in our dataset, the stronger one in chromosome 16 (within MATE efflux transporter coding region) and another in chromosome 14 (within an integral membrane HPP family protein and H+ ATPase coding regions). The validation of the genetic architecture of HCN in an African population, joint GWAS analysis between Africa and Latin America (Brazil) and whole genome imputation of the African-Brazilian dataset using HapMap as a reference for chromosome16 confirms results and shows that the genetic architecture of cyanide is conserved (based on our datasets). Homozygous reference alleles at identified loci showing lower cyanide content is in agreement with the finding that acyanogenic plants are homozygous recessive at one of the loci (Gleadow and Møller 2014). However, such homozygous cassava variety was yet to be identified given that they are recessive and difficult to discover because of cassava polyploid make-up (Jennings and Iglesias, 2002). Cyanide is maintained in cultivated cassava populations from Africa and Latin America via the selection of high and low cyanide phenotypes under different environmental and herbivore pressures, leading to a balanced selection. This phenomenon has been previously reported for cyanide in white clover (Corkhill L. 1942), *Sorghum bicolor* (Hansen et al. 2003) and *trifolium* (Kakes 1997) for cyanide. More recently, selective sweeps results between cultivated and cassava progenitors suggested that selection during domestication decreased CG content (Ramu et al. 2017).

Genome wide phylogenetic analysis of MATE genes in cassava, sorghum and arabidopsis suggested homology between our candidate gene and SbMATE2, a characterized vacuolar membrane transporter in sorghum for cyanogenic glucoside dhurrin (**Figure 3A)**. SbMATE2 functions in the accumulation of plant specialised metabolites such as flavonoids and alkaloids, and exports dhurrin and other hydroxynitrile glucosides, protecting against the self-toxic biochemical nature of chemical defence compounds. The transport of the pH-dependent unstable cyanogenic glucoside from its cytoplasmic site of production to the acidic vacuole likely contributes to reducing self-toxicity (Darbani et al. 2016). Mechanistic studies on MATE transporters, such as sorghum SbMATE gene, strongly suggest that its transport cycle could be driven by proton and/or cation (H^+^ or Na^+^) gradients (Doshi et al. 2017). SbMATE shows high affinity for Na^+^ & H^+^, and H^+^ constitute the main electrochemical driving force in plants, hence, it is likely that H^+^ constitutes the main coupling ion for SbMATE. Darbani et al. (2016), reported that the biosynthetic gene cluster for dhurrin additionally includes a gene encoding a MATE transporter and glutathione S-transferase gene for dhurrin uptake in sorghum bicolor.

Our study identified a MATE transporter on chromosomes 16 and the Na^+^ (from integral membrane HPP family protein) and a plasma membrane H^+^-ATPase-coupled on chromosome 14, as involved in cyanide content regulation. In the cassava genome version 6.1, the cyanide biosynthesis gene cluster is located on chromosome 12 within 75 kb interval, including a couple of changes in orientation and gene arrangement (**Supplementary Figure 3B).** Interestingly, genome-wide epistasis study did not reveal interactions with other parts of the genome, including the biosynthesis gene cluster region on chromosome 12. This finding contrasts with sorghum, where cyanide biosynthesis and transport have been characterized within the same gene cluster (Darbani et al. 2016). This suggests a distinct evolutionary path for cyanide regulation in cassava than in sorghum. In view of this observation, we speculate that perhaps, cassava domestication targeted upstream or downstream genetic regulation steps of cyanide bio-synthesis. In cassava, CGs are synthesized in the shoot apex (Andersen et al. 2000) and then transported to the fibrous roots (Nartey 1968; Koch et al. 1992; K. Jørgensen et al. 2005). Jorgensen et al (2005), reported a reduction of cyanogenic content in leaves of RNAi transgenic cassava plants, but not in the roots, indicating a tissue-specific regulation of cyanide accumulation in roots. Candidate Manes.16G007900 (chromosome16) showed local epistasis interaction with a 1.36Mb region located at 772055 – 775833 bp downstream. Epistatic effects that arise from alleles in gametic disequilibrium, between closely located loci can contribute to long-term response since recombination disrupts allelic combinations that have specific epistatic effects and the detection of epistasis is a key factor for explaining the missing heritability (Akdemir, Jannink, and Isidro-Sánchez 2017; Santantonio, Jannink, and Sorrells 2019). This region spans over 54 biallelic ancestry-informative single-nucleotide markers fixed or nearly fixed between *M. esculenta* and *M. flabellifolia* (Ogbonna et al, in press), suggesting that domestication can impact metabolic content targeting transport regulation (Wang et al. 2019), as earlier reported in maize and rice (Sosso et al. 2015). In view of the above findings, we speculate that cassava domestication may have specifically targeted downstream genetic regulation steps of cyanide biosynthesis. This is supported by the fact that root size (starch storage) and cyanide content are the major traits of cassava domestication (Ramu et al. 2017). HCN is regulated in an oligogenic manner with two major loci explaining the variation across our datasets. To facilitate their use in breeding pipelines, SNPs tagging the major QTL loci were converted to robust, high-throughput and easy to use competitive allele-specific PCR (KASP) assays. The diagnostic markers for cyanide are available (**Supplementary Table 7**) to the global cassava improvement community through a commercial genotyping service provider under the High Throughput Genotyping Project (https://excellenceinbreeding.org/htpg) via Intertek (https://www.intertek.com). We also observed that the closest homology observed for MATEs in cassava is in line with the results of the MATE protein alignment which displays the highest homology between MATE gene on chromosome 16 and chromosome 17 (Figure 3A). This is congruent with previously identified paleo tetraploidy in the cassava genome, where chromosomes 14 and 16 present partial conserved synteny with chromosome 6 and 17, respectively (Bredeson et al. 2016). We found the candidate gene to be paralog (68.59%) with Manes.16G008000 and homeolog (91.09%) with Manes.17G038400, indicating that our candidate had undergone double duplication events. This finding would need further investigation to clarify the potential fate of the observed tandem duplication (ie: subfunctionalization, neofunctionalization). MATE candidate gene topology prediction suggests that our candidate MATE protein shares a similar topology in the membrane as those observed in the MATE protein family and functions as an efflux carrier that mediates the extrusion of toxic substances (Brown, Paulsen, and Skurray 1999; Morita et al. 2000; Li et al. 2002). Further functional characterization of the putative cyanide transporters in cassava need to be performed.

Allele mining and mutation prediction (**Figure 2B)** on the HapMap dataset ensures that the current study captures the diversity of the HapMap panel. Moreover, DNA sequence analysis of Manes.16G007900 across HapMap individuals shows that *M. esculenta* ssp. *flabellifolia* individuals are preferentially homozygous G:G (high cyanide allele) for candidate SNP S16_773999, which is in line with its phenotypic characterization for cyanide content by Perrut-Lima and colleagues (Perrut-Lima, Mühlen, and Carvalho 2014). Interestingly, for the same candidate SNP, *M. glaziovii* and *M. pruinosa* individuals gene sequences are all homozygous A:A (low cyanide alleles) and cluster separately from *M. esculenta subsp. flabellifolia* (**FIgure 3B**). However, sweeps on HapMap data groups (Latin American, African and progenitors) did not reveal selective sweeps associated with GWAS loci and biosynthesis clusters. Phenotypic spatial distribution analysis results for sweet and bitter cassava in Brazil, suggested clinal variation occurred along subregions gradient separating ancestral coefficients across ecoregions and agrees with the candidate marker response in the region regulating cyanide variation in cassava. This reflects the role environmental conditions and herbivore pressure had played on cyanide regulation and its synergy in maintaining balanced selection of cyanide traits in cassava **(see further discussion: Supplementary Note3)**.

## Conclusion

In this study, we deciphered the genetic architecture of cyanide in cassava and mapped the genetic region in chromosome 16 and 14. The GWAS peak in chromosome 16 is strongly associated with the coding region of a MATE efflux protein, a transporter able to transport cyanogenic glucosides. In addition, the peaks on chromosome 14 is associated with the coding region of an integral membrane HPP family protein involved in Nitrite Transport Activity and a plasma membrane H^+^-ATPase mediated H^+^ influx which potentially associated with MATE to participate in a cyanide glucosides cotransport system.

Haplotype defined from the region in chromosome 16 and 14 explained 36 and 8% of the total variance explained by the markers, while loci associated with the optimal p-values explained 30 and 7% variance respectively. Selected individuals carrying the alleles for high and low cyanide in chromosomes 16 and 14 were further validated by designing KASP markers for breeding applications. This approach also found the same regions explaining the variance in an African dataset for cyanide, a joint dataset for African and Latin American germplasm and a whole genome imputation of the African-Brazilian dataset for chromosome 16, validating the candidate SNP. Sweet and bitter cassava distribution have maintained pre-conquest distribution in Brazil, with breeding activities around Northern and Central regions creating a more balanced population with low, intermediate and high cyanide clones.

The broader impact of this study was to understand the genetic mechanism of cyanide regulation in cassava root and the identification of closely linked SNP markers to enhance efficiency and cost effectiveness through marker assisted selection. Further steps can include (1) deployment of diagnostic markers for breeding applications; (2) develop co-expression studies to further assess the source/sink relationship of cyanide metabolism in multi-environmental conditions on impact of low cyanide on pest and disease control in cassava. (3) breeding and introduction of low cyanide cassava varieties that are high yielding and disease resistant to regions often affected by agricultural and health related crisis such as konzo, especially in sub-Saharan Africa. Altogether the present study consolidates our understanding of the genetic control of HCN variation in cassava and provides new insights using genomics of diverse genetic background populations.

## Material and Method

### Plant material

A first dataset including a total of 1,389 accessions from the Cassava Germplasm Banks (CGB) of Brazilian Agricultural Research Corporation (Embrapa), located in Cruz das Almas, Bahia, Brazil were used for this study (**Figure 1B**). The region is tropical with an average annual temperature of 24.5°C, relative humidity of 80%, and annual precipitation of 1,250 mm. The germplasm were collected from different cassava growing regions and ecosystems of Brazil, and consisted of land races and modern breeding lines (de Oliveira et al. 2014; Albuquerque et al. 2018).

A second data including 1,363 African accessions was obtained from the open source cassava breeding database, cassavabase.org. This dataset comprises plant material from the International Institute of Tropical Agriculture (IITA).

### DNA extraction

DNA extraction was performed following protocol described in Albuquerque et al (Albuquerque et al. 2018) and Ogbonna et al (in press) on the Embrapa CGB collection. Briefly, from young leaves according to the CTAB protocol (cetyltrimethylammonium bromide) as described by Doyle and Doyle (1987). The DNA was diluted in TE buffer (10 mM Tris-HCl and 1 mM EDTA) to a final concentration of 60 ng/µL, and the quality was checked by digestion of 250 ng of genomic DNA from 10 random samples with the restriction enzyme *Eco*RI (New England Biolabs, Boston, MA).

### Genotyping

Genotyping, imputation, filtering methods and parameters were performed as described in Ogbonna et al (in press). Briefly, Genotyping-By-Sequencing (Elshire et al. 2011) was conducted using the *Ape*KI restriction enzyme (Rabbi et al. 2014) and Illumina sequencing read lengths of 150 bp. Marker genotypes were called with the TASSEL GBS pipeline V5 (Glaubitz et al. 2014) using cassava reference genome version 6.1 available on Phytozome (http://phytozome.jgi.doe.gov). After filtering (mean depth values >5, missing data < 0.2 and minor allele frequency <0.01 per loci) and Imputation (AR^2^>0.8) (Browning and Browning 2009), remaining markers were retained for downstream analysis.

### Phenotyping

#### Brazilian dataset

Phenotypic data were collected on 1,389 accessions over 4 trials in a single location with 3 replications each in 2016, 2017, 2018 and 2019. A total of 1,246 accessions had both phenotypic and genotypic information and were retained for further analysis. Cyanide content was measured using picrate titration methods (M. G. Bradbury, Egan, and Howard Bradbury 1999) as earlier described by Fukuda et al. (2010). Briefly, it involves a qualitative determination of HCN potential in cassava root, and given that HCN potential varies considerably in plants, we assayed 5 to 6 plants in a plot and 3 roots per plant. Cross sectional 1 cm^3^ cut is made at mid-root position foreach root, between the peel and the center of the parenchyma. The cut root-cube and 5 drops of toluene (methylbenzene or phenyl methane) are added to a glass test tube respectively and tightly sealed with a stopper. To determine the qualitative score of HCN potential on a color scale of 1 to 9, a strip of whatman filter paper is dipped into a freshly prepared alkaline picrate mixture until saturation. The saturated filter paper is then placed above the cut root cube in the glass tube and tightly sealed for 10 to 12 hours before recording the color intensity (Maziya-Dixon, Dixon, and Adebowale 2007). See **Supplemental Table 1** for a HCN assay for Brazilian germplasm across four years.

### African and Colombian datasets

African phenotypic data was collected from a breeding database cassavabase (https://cassavabase.org) and include 18 locations, 23 years and 393 trials for a total of 8,244 accessions for a total of 33,523 observations from the Institute of Tropical Agriculture (IITA) (**Supplementary Figure 8C**). Colombian phenotypic data includes 41 locations, 11 years and 155 trials for a total of 13,111 observations from the Centro Internacional de Agricultura Tropical (CIAT). The Phenotyping protocol was performed using the same protocol as for the Brazilian dataset. A total of 636 unique accessions with phenotypic and genotypic information from 228 trials were retained for further analysis for the African dataset.

### Statistical Analyses

Trials across years were combined and BLUPs were estimated for each clone from 9,138 observations on 1,389 genotypes for HCN. We used the lme4 (Bates et al. 2015) package in R (R. Core Team 2015) version 3.4.2 (2017-09-28) to fit a mixed linear model (MLM) following the method described in (Wolfe et al. 2016): *Y*_*ijkl*_ = *μ* + *c*_*i*_ + *β*_*j*_ + *r*_*k*_ + *d*_*l*_ + *ϵ* _*ijkl*_, where 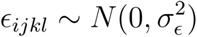 is the residual variance and assumed to be randomly distributed, *Y*_*ijkl*_ represents the phenotypic observations, *μ* is the grand mean, *c*_*i*_ is the random effects for clone with 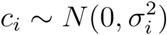, *β*_*j*_ is a random effect for clone nested in combination of year, *r*_*k*_ is a random effect for combination of year and rep and assumed to be normally distribution with 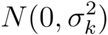 and *d*_*l*_ is a fixed effect for years. Variance components from our mixed model were used to compute the broad-sense heritability according to the method described in (Holland, Nyquist, and Cervantes-Martínez 2010) Briefly, 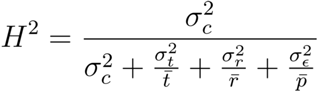, where 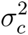 is the clone variance, 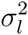 is variance due to clone by year, 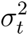 is the variance due to years by replications and 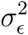 is the variance due to error. While 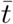, 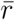 and 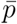 are the harmonic mean number of years, replications and plots in which the clone was observed, respectively. Given that the number of observations per clones varies across the four years dataset (replication varies from 1 to 9 with an average of 6), bias induced by precorrection and induced-heterogeneous residual variance (de Los Campos et al. 2013), estimated BLUPs (differentially shrunken to the mean) were deregressed using:

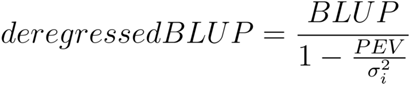

Where *PEV* is the prediction error variance for each clone and 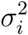 is the variance for the clonal component. **Supplementary Figure 8** shows the deregressed BLUPs distribution that was further used in the GWAS.

### GWAS analysis

We carried out a mixed model Genome Wide Association Mapping using GCTA software (Yang et al. 2011). Specifically, we used the mixed linear model based association analysis with the chromosome, on which the candidate SNP is located, excluded from calculating the genetic relationship matrix (GRM). The model is *y* = *a* + *bx* + *g*^−^ + *e* where *y* is the deregressed-BLUP estimate, *a* is the mean term, *b* is the additive effect (fixed effect) of the candidate SNP to be tested for association, *x* is the SNP genotype indicator variable, *g*^−^ is the accumulated effect of all SNPs except those on the chromosome where the candidate SNP is located, and *e* is the residual. We plotted Manhattan with the Bonferroni threshold used as association tests of significant SNPs and compared the observed −log10(p-value) against the expected using the quantile-quantile plot. Local LD analysis will be performed on GWAS significant region based on *r*^2^ threshold of > 0.8 to identify candidate genes. GWAS was also performed on a unique set of 1,536 individuals (GU panel) from Ogbonna et al., (in press). This unique set was selected based on duplicate (identity-by-state) analysis on the total population of 3,354 individuals to ensure efficient germplasm and resource management at the Brazilian cassava program and to balance individual genetic contribution to population structure definition (Ogbonna et al., In press).

### Candidate Gene Analysis

To investigate further GWAS candidate regions, we used the genomic resource from the cassava HapMap data (Ramu et al. 2017) to perform allele mining and predict genome-wide allelic mutation effect using SNPeff (Cingolani et al. 2012) and SIFT (Kumar, Henikoff, and Ng 2009).

### Phylogenetic analysis of candidate gene sequence

We obtained MATE whole genome protein sequences from *Arabidopsis thaliana* (v.10), *Sorghum bicolor* (v3.11) and *Manihot esculenta* (v.6.1) genomes from phytozome (https://phytozome.jgi.doe.gov/pz/portal.html). The sequences were submitted to the transporter TP prediction tool (http://bioinfo3.noble.org/transporter/) for membrane domain identification and gene curation according to the TCDB guidelines (Saier et al. 2016). Sequences were aligned with CLUSTAL OMEGA (Sievers et al. 2011) and a phylogenetic analysis was performed using a Neighbour-joining tree without distance corrections (**Supplementary Sequence Data, Supplementary Multiple Alignment**). In addition, we generatedMATE candidate, Manes.16G007900, protein sequences from the cassava hapmap (Ramu et al. 2017). Briefly Manes16.G007900 annotated variants from HapMap II (ftp://ftp.cassavabase.org/HapMapII/) were used to generate coding sequences (CDS) and translated protein sequence for each 241 accessions in a fasta format. Subsequent alignment and maximum-likelihood phylogenetic trees were generated using MAFFT (Katoh and Standley 2013) and PhyML (Guindon et al. 2010) through the NGphylogeny portal (Lemoine et al. 2019).

## Supporting information

Supplementary Multiple Sequence Alignment

Supplementary Sequence Dataset

## Supplemental Data

The following supplemental materials are available.

**Supplementary Figure 1.** Manhattan plot and LD plots for chromosomes 16 and 14.

**Supplementary Figure 2.** Pearson correlation of top 5 significant SNPs.

**Supplementary Figure 3.** Schematic representation of the clustering of cyanogenic glucoside biosynthetic genes.

**Supplementary Figure 4.** Differentiating loci between cultivated and cassava progenitor.

**Supplementary Figure 5.** TMHMM posterior probability for transmembrane Protein and mutation prediction.

**Supplementary Figure 6.** Distribution of sweet and bitter cassava in Sub-Saharan Africa.

**Supplementary Figure 7.** Manhattan plot for whole-genome imputed chromosomes 16.

**Supplementary Figure 8.** Distribution of HCN assayed for Latin American and African.

**Supplementary Figure 9.** Selective sweeps between cassava Progenitors and Latin American.

**Supplementary Figure 10.** Selective sweeps between Latin American and African cassava.

**Supplementary Figure 11.** Genetic (cM) vs. Physical (bp) positions.

**Supplementary Table 1.** Raw HCN dataset from Latin America (Embrapa-Brazil).

**Supplementary Table 2.** Summary statistics, variance components and broad-sense heritability for HCN.

**Supplementary Table 3.** 1,389 BLUPs for Latin American (Embrapa-Brazil) dataset and the list of 1,246 BLUPs with genotype information used for GWAS.

**Supplementary Table 4.** Significant SNPs from Latin American dataset (Embrapa-Brazil).

**Supplementary Table 5.** Significant SNPs from GWAS on 523 Unique individuals.

**Supplementary Table 6.** Cultivated and Cassava Progenitor Differenting loci comparison; *M. esculenta, M. esculenta* ssp. *flabellifolia*.

**Supplementary Table 7.** Designed KASP Marker Sequences.

**Supplementary Table 8.** HCN kaspar ségrégation results.

**Supplementary Table 9.** 242 Significant epistasis interactions pair of SNPs higher than bonferroni correction threshold (2 way test result).

**Supplementary Table 10.** Single point mutation prediction for Manes.16G007900 and Manes.16G008000

**Supplementary Table 11.** List of Countries and regions in Sub-Saharan Africa with their average BLUP values.

**Supplementary Table 12.** Raw African dataset phenotypes.

**Supplementary Table 13.** African BLUPs used for GWAS analysis.

**Supplementary Table 14.** Significant SNPs from African Germplasm GWAS analysis.

**Supplementary Table 15.** Raw African (IITA) and Latin American (Embrapa) phenotypes.

**Supplementary Table 16.** 1882 Combined BLUPs for Africa (IITA) and Latin America (Embrapa) GWAS.

**Supplementary Table 17.** Significant SNPs from African and Brazil Germplasm.

**Supplementary Table 18.** Significant SNPs from Whole Genome Imputation of chromosome 16 GWAS using HapMapII and raw GBS dataset. 5000 SNP window was used.

**Supplementary Sequence Dataset.** Whole-genome sequence dataset for all MATE genes in Cassava, Arabidopsis and Sorghum.

**Supplementary Multiple Sequence Alignment.** Multiple sequence alignment for all MATE genes in Cassava, Arabidopsis and Sorghum.

Supplementary Method 1 Proportion of Variance Explained by Markers

Supplementary Method 2 Genome-wide Epistasis Interactions

Supplementary Method 3 Cultivated and Cassava Progenitor Differentiating Loci Analysis

Supplementary Method 4 KasparMarker Design and Assessment

Supplementary Method 5 Candidate gene protein Topology and Structure Prediction.

Supplementary Method 6 Single Point Mutation Prediction

Supplementary Method 7 Geographical Distribution of HCN

Supplementary Method 8 GWAS in African Population and Joint Africa, Latin America Analysis

Supplementary Note 1 Population Structure Analysis

Supplementary Note 2 Phylogenetic tree

Supplementary Note 3 Sweet and Bitter cassava Geographical Distribution

## Availability of data and material

Genotyping (SNP) data used in this study were deposited on cassavabase.org hosted at “ftp://ftp.cassavabase.org/manuscripts/Ogbonna_et_al_2020/gwas_manuscript“.

## Acknowledgements

The authors appreciate Jean-Luc Jannink and Deniz Akdemir both from Cornell University, Ithaca, NY for their invaluable advice. The authors thank Kelly Robbins and Victoria Gomez for reviewing this manuscript. We are grateful to Embrapa Cruz das Almas breeding team for field experiment management. We thank the IITA cassava team and particularly Alfred Dixon, Peter Kulakow, Prasad Peteti, cassava breeders and data manager for making the historical African dataset publicly available. We thank the CIAT Breeding team and breeder Hernan Ceballos for making the historical Colombian dataset publicly available. This work was supported through Boyce Thompson Institute for plant science in collaboration with Embrapa Mandioca e Fruticultura of Brazil and partially supported by the NEXTGEN Cassava project, through a grant to Cornell University by the Bill & Melinda Gates Foundation and the UK Department for International Development.

## Authors Contributions

Designed experiment: AO, EJO, GB; Performed experiment: AO, GB, LBRA. Project supervision: LM, GB, EJO; First draft of the manuscript: AO; IYR provided technical assistance. GB and EO agree to serve as authors responsible for contact and ensure communication. All authors reviewed and approved the manuscript.

