## Supplementary Multiple Sequence Alignment for "Genetic architecture and gene mapping of cyanide in cassava (*Manihot esculenta Crantz*.)"

CLUSTAL O(1.2.4) multiple sequence alignment for whole-genome MATE genes in Arabidopsis, Sorghum and Cassava

```
Sobic.002G099300.1      ------------------------------------------------------------ 0
AT4G39030.1             ---MLIKSQRLT--------LFSPLLSKTRRIPVNSHQTLVAES---------------- 33
Manes.04G084700.1       -----MQIKSMA--------LSS------PTTPL--QNPFLKKKFI-S---SFIFKNPPY 35
Manes.11G091900.1       --------------------MNIQSLLHSSHTPFRTHEPIFLSQ---S---LILSKKSPL 34
Sobic.004G019800.2      ------------------------------------------------------------ 0
Sobic.004G019800.3      ------------------------------------------------------------ 0
Sobic.004G019800.4      ------------------------------------------------------------ 0
AT2G21340.1             ---MQIQCKTLT--------FTVSSIPCNPKLPFPS------------------------ 25
Manes.11G092000.1       ----MMQIK---------------TLAHRPPLISPQK-PNFKH----------YYSQSPL 30
Sobic.001G476700.3      ------------------------------------------------------------ 0
Sobic.009G077000.1      ------------------------------------------------------------ 0
Sobic.001G454900.1      -----MED----------------------------------NGAA-AA-----NAAPAT 15
Sobic.003G403000.1      -----MEE----------------------------------HRSP-AHAKP-EAEQPPQ 19
Sobic.003G403000.2      -----MEE----------------------------------HRSP-AHAKP-EAEQPPQ 19
Sobic.003G403000.3      -----MEE----------------------------------HRSP-AHAKP-EAEQPPQ 19
Sobic.007G020600.1      ------------------------------------------------------------ 0
Sobic.007G020600.2      ------------------------------------------------------------ 0
AT3G08040.1             ------------------------------------------------------------ 0
Manes.09G027700.1       ------------------------------------------------------------ 0
Manes.09G027800.1       ------------------------------------------------------------ 0
Manes.09G027900.1       ------------------------------------------------------------ 0
Manes.09G026900.1       ------------------------------------------------------------ 0
Manes.09G026900.2       ------------------------------------------------------------ 0
Manes.09G027000.1       ------------------------------------------------------------ 0
Manes.07G006000.1       ------------------------------------------------------------ 0
Manes.07G006000.2       ------------------------------------------------------------ 0
Manes.10G143000.1       ------------------------------------------------------------ 0
AT1G51340.2             ------------------------------------------------------------ 0
Manes.06G164500.1       ------------------------------------------------------------ 0
Manes.06G164500.2       ------------------------------------------------------------ 0
Manes.06G164500.3       ------------------------------------------------------------ 0
Manes.06G164500.4       ------------------------------------------------------------ 0
Manes.06G164500.5       ------------------------------------------------------------ 0
Manes.14G002600.1       ------------------------------------------------------------ 0
Sobic.008G006100.1      -----MAA------TSPTP-TRPAAAAHAR-TLSA------------------GP--SRV 27
Sobic.005G005400.1      ------MV------TSPLVTVRAAAFALTL-TPSS------------------RL--SRA 27
Sobic.005G005400.3      ------MV------TSPLVTVRAAAFALTL-TPSS------------------RL--SRA 27
Sobic.005G005400.2      ------MV------TSPLVTVRAAAFALTL-TPSS------------------RL--SRA 27
AT2G38330.1             -----MAA------V-------ATSFCFSP-HRS-----------------PSRFGNPNS 24
Manes.08G096600.1       -----MAT------VQSLKLLSTHSFHTNP-FPKS----------------QSFTANPNF 32
Manes.08G096600.2       -----MAT------VQSLKLLSTHSFHTNP-FPKS----------------QSFTANPNF 32
AT4G38380.1             ---------------------------------MESSRVVVGGGLPLANRRNSSFAKPKI 27
Sobic.002G286800.1      --------MEFAAGAGVAPRLRLRSLPAGP-VPGRSGRVMVGGGGAIGIGWARRATAPGL 51
Sobic.003G149300.2      ---------------------MLHLHHHSP-LVSRHGAVCPPTRLACS--CICNKANSRI 36
Sobic.003G149300.1      ---------------------MLHLHHHSP-LVSRHGAVCPPTRLACS--CICNKANSRI 36
Sobic.003G149300.3      ---------------------MLHLHHHSP-LVSRHGAVCPPTRLACS--CICNKANSRI 36
Sobic.003G149300.4      ------------------------------------------------------------ 0
Manes.01G153400.1       ------------------------------------------------------------ 0
Manes.04G064900.1       -----MTTRQFS-GSNLSRGLARKSSVHNE-AA--------KGTRRFSLGCPSEVVK--- 42
Manes.04G064900.2       -----MTTRQFS-GSNLSRGLARKSSVHNE-AA--------KGTRRFSLGCPSEVVK--- 42
Sobic.001G003700.1      ------------------------------------------------------------ 0
AT4G22790.1             --------------------------MS-------------------------------- 2
Manes.02G032300.1       --------------------------MS-------------------------------- 2
Manes.10G000400.1       ------------------------------------------------------------ 0
AT2G38510.1             ------------------------------------------------------------ 0
Manes.08G172300.1       ------------------------------------------------------------ 0
Manes.09G117000.1       ------------------------------------------------------------ 0
Sobic.010G167800.1      --------------------------MCTTPVPSPP----A------V-A-LV-GVKGGH 21
AT5G19700.1             ------------------------------------------------------------ 0
AT4G29140.1             --------------------------MCNPSTTTTT----T------G---SE-NQESRT 20
Manes.03G026500.1       --------------------------MCNPDTTAAL----L------T-EKPA-GQKPQT 22
Manes.16G109300.1       --------------------------MCNSDITTTL----L------I-EKPP-QK---P 19
Sobic.007G181100.1      --------------------------MCHCHCSEP--------------------TQCHH 14
Manes.13G127800.1       --------------------------MTREVSS--------------------------- 7
Sobic.001G446800.1      --------------------------MCNAAAGTD--SS-----------LPA-PAAPFG 20
AT5G52050.1             -------------------------------------------------------MSQSN 5
AT1G58340.1             --------------------------MCNSKPSSA--SSSL------L-SCKD-KTHISK 24
```

```
Manes.05G186700.1       --------------------------MCNPKPSSQ--TSFL---------CPN-KT----  18
Manes.18G054300.1       --------------------------MCNPKPSSP--SSFL---------CPK-TT----  18
Sobic.001G320900.1      --------------------------MCEGLVG-----KLL------P-PCLC-NAKDGG  21
Sobic.001G320900.2      --------------------------MCEGLVG-----KLL------P-PCLC-NAKDGG  21
Sobic.004G283500.1      -----MSSC-----------AGATV-ALRDAAEEA------------L-NASL-LSK--A  27
Sobic.006G184400.1      -----MTSC-----------AGDATVACRDDARPHGDAACG------V-SCPL-LAKPAG  36
AT4G23030.1             ------------------------------------------------------------  0
Manes.01G067000.1       --------------------------MCQLASPRD--------------SCKS-NLDQST  19
Manes.02G027800.1       --------------------------MCQLSSPPH--------------SCKT-NLDPST  19
Manes.06G143600.1       --------------------------MCQ---LNS---SLS------S-KCPC-LLSIQD  20
Manes.14G029100.1       --------------------------MCHRQPNSS---ALS------S-RCPC-LLSIKD  23
AT5G49130.1             ------------------------------------------------------------  0
Manes.03G198000.1       ------------------------------------------------------------  0
Manes.15G011000.1       ------------------------------------------------------------  0
Sobic.001G019700.1      ---------------------------------------MA------I-PLPG-KALPRH  13
AT1G71870.1             ------------------------------------------------------------  0
Manes.02G189800.1       ------------------------------------------------------------  0
Manes.18G098800.1       ------------------------------------------------------------  0
Sobic.010G256932.1      ------------------------------------------------------------  0
Sobic.010G256700.1      ------------------------------------------------------------  0
Sobic.010G256700.5      ------------------------------------------------------------  0
Sobic.010G256700.6      ------------------------------------------------------------  0
Sobic.010G256700.4      ------------------------------------------------------------  0
Sobic.009G106900.1      ------------------------------------------------------------  0
Manes.S031500.1         ------------------------------------------------------------  0
Manes.14G109900.1       ------------------------------------------------------------  0
Manes.14G109900.2       ------------------------------------------------------------  0
Manes.14G109900.3       ------------------------------------------------------------  0
AT1G73700.1             ------------------------------------------------------------  0
AT2G34360.1             ------------------------------------------------------------  0
AT5G52450.1             ------------------------------------------------------------  0
Sobic.009G106960.1      ------------------------------------------------------------  0
Manes.14G060800.1       ------------------------------------------------------------  0
Sobic.009G106800.1      ------------------------------------------------------------  0
Sobic.007G074300.2      ------------------------------------------------------------  0
Sobic.009G106700.1      ------------------------------------------------------------  0
Sobic.010G138400.4      ------------------------------------------------------------  0
Sobic.010G138400.1      ------------------------------------------------------------  0
Sobic.010G138400.5      ------------------------------------------------------------  0
Sobic.004G129900.1      ------------------------------------------------------------  0
Sobic.004G129900.2      ------------------------------------------------------------  0
Sobic.006G042200.1      ------------------------------------------------------------  0
Sobic.004G129700.1      ------------------------------------------------------------  0
Sobic.004G129800.1      ------------------------------------------------------------  0
Sobic.004G129800.2      ------------------------------------------------------------  0
AT3G23550.1             ------------------------------------------------------------  0
AT3G23560.1             ------------------------------------------------------------  0
Manes.15G088100.1       ------------------------------------------------------------  0
Manes.03G109100.1       ------------------------------------------------------------  0
Manes.03G109100.2       ------------------------------------------------------------  0
Sobic.002G311200.1      -----MRQC-----------RAAIFIFFPIHSRLHA--MLR------AESHLH-AIIPRG  35
Sobic.002G006500.3      ------------------------------------------------------------  0
Sobic.002G006500.4      ------------------------------------------------------------  0
AT2G04090.1             ------------------------------------------------------------  0
AT2G04100.1             ------------------------------------------------------------  0
AT2G04066.1             ------------------------------------------------------------  0
AT2G04040.1             ------------------------------------------------------------  0
AT2G04080.1             ------------------------------------------------------------  0
AT2G04050.1             ------------------------------------------------------------  0
AT2G04070.1             ------------------------------------------------------------  0
Sobic.001G481800.1      ------------------------------------------------------------  0
Sobic.003G260200.1      ------------------------------------------------------------  0
Sobic.009G224800.1      ------------------------------------------------------------  0
Sobic.009G224800.2      ------------------------------------------------------------  0
AT1G66760.2             ------------------------------------------------------------  0
AT1G64820.1             ------------------------------------------------------------  0
AT1G66780.1             ------------------------------------------------------------  0
Manes.02G072400.1       ------------------------------------------------------------  0
Manes.01G113500.1       ------------------------------------------------------------  0
Manes.01G113500.2       ------------------------------------------------------------  0
AT1G15150.1             ------------------------------------------------------------  0
```

```
AT1G15160.1             ------------------------------------------------------------ 0
AT1G15170.1             ------------------------------------------------------------ 0
AT1G15180.1             ------------------------------------------------------------ 0
AT1G71140.1             ------------------------------------------------------------ 0
Manes.06G063000.1       ------------------------------------------------------------ 0
Manes.14G109700.1       ------------------------------------------------------------ 0
Manes.14G109800.1       ------------------------------------------------------------ 0
Sobic.008G171600.1      ------------------------------------------------------------ 0
Sobic.008G171600.2      ----------------------------------------------------MLDQKKFR 8
Sobic.008G171600.3      ------------------------------------------------------------ 0
AT3G59030.1             ------------------------------------------------------------ 0
Manes.01G182000.1       ------------------------------------------------------------ 0
Manes.02G142000.1       ------------------------------------------------------------ 0
AT4G21903.2             ------------------------------------------------------------ 0
AT4G21910.4             ------------------------------------------------------------ 0
AT1G11670.1             ------------------------------------------------------------ 0
AT1G61890.1             ------------------------------------------------------------ 0
Manes.16G008000.1       ------------------------------------------------------------ 0
Manes.17G038200.1       ------------------------------------------------------------ 0
Sobic.002G318300.1      ------------------------------------------------------------ 0
Manes.17G038300.1       ------------------------------------------------------------ 0
Manes.16G007900.1       ------------------------------------------------------------ 0
Manes.17G038400.1       ------------------------------------------------------------ 0
Sobic.001G012600.1      ------------------------------------------------------------ 0
Sobic.001G012600.2      ------------------------------------------------------------ 0
Sobic.001G185600.1      ------------------------------------------------------------ 0
Sobic.001G185400.1      ------------------------------------------------------------ 0
AT3G21690.1             ------------------------------------------------------------ 0
Sobic.001G185800.1      ------------------------------------------------------------ 0
Sobic.001G185500.1      ------------------------------------------------------------ 0
Sobic.001G185500.2      ------------------------------------------------------------ 0
Sobic.007G165500.2      ------------------------------------------------------------ 0
AT4G00350.1             ------------------------------------------------------------ 0
Manes.01G255000.1       MAILISRSTNTA----ASRHMHALLIVTNKKEIISNKKKIS------K-SWKEKKEKKER 49
Sobic.004G349550.1      ------------------------------------------------------------ 0
Sobic.004G349600.1      ------------------------------------------------------------ 0
Sobic.007G176000.1      ------------------------------------------------------------ 0
Sobic.007G176100.1      ------------------------------------------------------------ 0
AT1G47530.1             ------------------------------------------------------------ 0
Manes.05G164500.1       ------------------------------------------------------------ 0
Manes.05G164500.2       ------------------------------------------------------------ 0
Manes.18G030900.1       ------------------------------------------------------------ 0
AT1G23300.1             ------------------------------------------------------------ 0
AT3G26590.1             ------------------------------------------------------------ 0
AT5G38030.1             ------------------------------------------------------------ 0
Manes.12G023400.1       ------------------------------------------------------------ 0
Manes.12G023500.1       ------------------------------------------------------------ 0
Manes.12G023600.1       ------------------------------------------------------------ 0
Manes.13G025200.1       ------------------------------------------------------------ 0
Manes.13G025200.2       ------------------------------------------------------------ 0
Sobic.001G273100.1      ------------------------------------------------------------ 0
Sobic.001G273100.2      ------------------------------------------------------------ 0
Sobic.001G273000.2      ------------------------------------------------------------ 0
Sobic.001G273000.1      ------------------------------------------------------------ 0
Sobic.001G273000.3      ------------------------------------------------------------ 0
Manes.15G147800.1       ------------------------------------------------------------ 0
Manes.15G147900.1       ------------------------------------------------------------ 0
Manes.17G098800.1       ------------------------------------------------------------ 0
Manes.17G098900.1       ------------------------------------------------------------ 0
Manes.17G098900.2       ------------------------------------------------------------ 0
Sobic.001G162400.1      ------------------------------------------------------------ 0
Sobic.003G307600.2      ------------------------------------------------------------ 0
Sobic.002G232200.1      ------------------------------------------------------------ 0
Sobic.002G232500.1      ------------------------------------------------------------ 0
Sobic.002G232600.1      ------------------------------------------------------------ 0
Sobic.007G160700.1      ------------------------------------------------------------ 0
Sobic.003G126200.2      ------------------------------------------------------------ 0
Sobic.001G476700.1      ------------------------------------------------------------ 0
Sobic.001G476700.2      ------------------------------------------------------------ 0
Manes.18G062800.1       ------------------------------------------------------------ 0
AT5G44050.1             ------------------------------------------------------------ 0
```

```
AT5G10420.1          ------------------------------------------------------------  0
AT5G65380.1          ------------------------------------------------------------  0
Manes.12G129000.1    ------------------------------------------------------------  0
Manes.13G097900.1    ------------------------------------------------------------  0
Sobic.005G020700.1   ------------------------------------------------------------  0
Sobic.008G019400.1   ------------------------------------------------------------  0
Manes.09G135300.1    ------------------------------------------------------------  0
AT1G33080.1          ------------------------------------------------------------  0
AT1G33090.1          ------------------------------------------------------------  0
AT1G33100.1          ------------------------------------------------------------  0
AT1G33110.1          ------------------------------------------------------------  0
AT3G03620.1          ------------------------------------------------------------  0
AT5G17700.1          ------------------------------------------------------------  0
Manes.09G135200.1    ------------------------------------------------------------  0
Manes.09G134800.1    ------------------------------------------------------------  0
Manes.08G150400.1    ------------------------------------------------------------  0
Manes.09G134900.1    ------------------------------------------------------------  0
Manes.09G135000.1    ------------------------------------------------------------  0
Manes.09G134900.2    ------------------------------------------------------------  0
Manes.09G135000.2    ------------------------------------------------------------  0


Sobic.002G099300.1   ------------------------------------------------------------  0
AT4G39030.1          -VI--T-RRTL--GAITATPSFHKN------PV-----------VIRRRI---KLE----  63
Manes.04G084700.1    F-------------------SLPLPLSPPSLRI-----------SYRTSQ---HFS----  58
Manes.11G091900.1    SLA--S-PK--------SFPS---SLLPPS---------------LRTIK---SRS----  58
Sobic.004G019800.2   ------------------------------------------------------------  0
Sobic.004G019800.3   ------------------------------------------------------------  0
Sobic.004G019800.4   ------------------------------------------------------------  0
AT2G21340.1          -SL--T-LRSW--N--PSFPSFRSSAVSGP----------------KSSL---KLN----  54
Manes.11G092000.1    HLL--T-LRTH--S--SIFPS----SVHLS----------------APKY---RRN----  56
Sobic.001G476700.3   ------------------------------------------------------------  0
Sobic.009G077000.1   ------------------------------------------------------------  0
Sobic.001G454900.1   S----------------MLPAAE------------------------------KHV----  25
Sobic.003G403000.1   Q----------------QVPAAMA--------VAVAVD------VAAPAAL-QNST----  44
Sobic.003G403000.2   Q----------------QVPAAMA--------VAVAVD------VAAPAAL-QNST----  44
Sobic.003G403000.3   Q----------------QVPAAMA--------VAVAVD------VAAPAAL-QNST----  44
Sobic.007G020600.1   ------------------------------------------------------------  0
Sobic.007G020600.2   ------------------------------------------------------------  0
AT3G08040.1          ------------------------------------------------------------  0
Manes.09G027700.1    ------------------------------------------------------------  0
Manes.09G027800.1    ------------------------------------------------------------  0
Manes.09G027900.1    ------------------------------------------------------------  0
Manes.09G026900.1    ------------------------------------------------------------  0
Manes.09G026900.2    ------------------------------------------------------------  0
Manes.09G027000.1    ------------------------------------------------------------  0
Manes.07G006000.1    ------------------------------------------------------------  0
Manes.07G006000.2    ------------------------------------------------------------  0
Manes.10G143000.1    ------------------------------------------------------------  0
AT1G51340.2          ------------------------------------------------------------  0
Manes.06G164500.1    ------------------------------------------------------------  0
Manes.06G164500.2    ------------------------------------------------------------  0
Manes.06G164500.3    ------------------------------------------------------------  0
Manes.06G164500.4    ------------------------------------------------------------  0
Manes.06G164500.5    ------------------------------------------------------------  0
Manes.14G002600.1    ------------------------------------------------------------  0
Sobic.008G006100.1   S-----------------------------LTSCRCHC---------RRSTPPRWS----  45
Sobic.005G005400.1   S-----------------------------FRACLAQR---------RR---RRW-----  41
Sobic.005G005400.3   S-----------------------------FRACLAQR---------RR---RRW-----  41
Sobic.005G005400.2   S-----------------------------FRACLAQR---------RR---RRW-----  41
AT2G38330.1          S-----------------------------IR-RTIVC---------K-SSPRDES----  40
Manes.08G096600.1    S-----------------------------VR-SLAAA---------PKSSPQK-S----  48
Manes.08G096600.2    S-----------------------------VR-SLAAA---------PKSSPQK-S----  48
AT4G38380.1          QQG--T-----------FLPLSRINNVSAPQKCSLHTNPNPMF----PFVTRRKS-----  65
Sobic.002G286800.1   SLA----------------PAV--------ARRAVSAAGGGH--LLHGRV----V-----  76
Sobic.003G149300.2   T-------------------------------AAQSTNNNANLLLLRGWSSTRRR-----  60
Sobic.003G149300.1   T-------------------------------AAQSTNNNANLLLLRGWSSTRRR-----  60
Sobic.003G149300.3   T-------------------------------AAQSTNNNANLLLLRGWSSTRRR-----  60
Sobic.003G149300.4   ------------------------------------------------------------  0
Manes.01G153400.1    ------------------------------------------------------------  0
Manes.04G064900.1    ----------------------GLGRRDVASNCCLSADYR---KVLSPSVTRRKK-----  72
```

```
Manes.04G064900.2      ----------------------GLGRRDVASNCCLSADYR---KVLSPSVTRRKK-----  72
Sobic.001G003700.1     ------------------------------------------------------------  0
AT4G22790.1            --------------------------------E-TS------------------------  5
Manes.02G032300.1      --------------------------------T-AT------------------------  5
Manes.10G000400.1      ------------------------------------------------------------  0
AT2G38510.1            ------------------------------------------------------------  0
Manes.08G172300.1      ------MR---------------------ARGE-TQ------------------------  8
Manes.09G117000.1      ------------------------------------------------------------  0
Sobic.010G167800.1     HVY----------VTVPQVPEGD----D--------------------------------  35
AT5G19700.1            ----------------METPNII----S--------------------------------  8
AT4G29140.1            GLF----------LDLFSINSFE----P--------------------------------  34
Manes.03G026500.1      QLY----------MHLLSLPSTI----K--------------------------------  36
Manes.16G109300.1      HRY----------LDRLSLPTTV----K--------------------------------  33
Sobic.007G181100.1     Q--------------VLLPP----------------------------------------  20
Manes.13G127800.1      ----------------ASVPLLG----SYSS----Q------------------------  19
Sobic.001G446800.1     --------------------------------S---------------------------  21
AT5G52050.1            -------RVR---DEV--------------------------------------------  11
AT1G58340.1            ------------------------------------------------------------  24
Manes.05G186700.1      ------------------------------------------------------------  18
Manes.18G054300.1      ------------------------------------------------------------  18
Sobic.001G320900.1     GVGDDGHRLR---AVVVSPDVAV----VVVLNE-ST------------------------  49
Sobic.001G320900.2     GVGDDGHRLR---AVVVSPDVAV----VVVLNE-ST------------------------  49
Sobic.004G283500.1     DVRLPV---------EDVPPVLT----SKPPGR-FA------------------------  49
Sobic.006G184400.1     EVHIPVAG-----EDAPAVPVLT----TKSPGR-LA------------------------  62
AT4G23030.1            --------------------MAA----PLLM-----------------------------  7
Manes.01G067000.1      -------R-------LHKSDMLI----PLVS-----------------------------  32
Manes.02G027800.1      -------K-----DEVQKPDTLT----PLVL-----------------------------  34
Manes.06G143600.1      -------P-----EANMHTPFLI----PKSPTS-KQ------------------------  39
Manes.14G029100.1      -------P-----AESDIQTFLI----PKSPAP-KQ------------------------  42
AT5G49130.1            ------------------------------------------------------------  0
Manes.03G198000.1      ------------------------------------------------------------  0
Manes.15G011000.1      ------------------------------------------------------------  0
Sobic.001G019700.1     DVS---------------------------------------------------------  16
AT1G71870.1            ------------------------------------------------------------  0
Manes.02G189800.1      ------------------------------------------------------------  0
Manes.18G098800.1      ------------------------------------------------------------  0
Sobic.010G256932.1     ------------------------------------------------------------  0
Sobic.010G256700.1     ------------------------------------------------------------  0
Sobic.010G256700.5     ------------------------------------------------------------  0
Sobic.010G256700.6     ------------------------------------------------------------  0
Sobic.010G256700.4     ------------------------------------------------------------  0
Sobic.009G106900.1     ------------------------------------------------------------  0
Manes.S031500.1        ------------------------------------------------------------  0
Manes.14G109900.1      ------------------------------------------------------------  0
Manes.14G109900.2      ------------------------------------------------------------  0
Manes.14G109900.3      ------------------------------------------------------------  0
AT1G73700.1            ------------------------------------------------------------  0
AT2G34360.1            ------------------------------------------------------------  0
AT5G52450.1            ------------------------------------------------------------  0
Sobic.009G106960.1     ------------------------------------------------------------  0
Manes.14G060800.1      -----------------------------------------------------------M  1
Sobic.009G106800.1     ------------------------------------------------------------  0
Sobic.007G074300.2     ------------------------------------------------------------  0
Sobic.009G106700.1     ------------------------------------------------------------  0
Sobic.010G138400.4     ------------------------------------------------------------  0
Sobic.010G138400.1     ------------------------------------------------------------  0
Sobic.010G138400.5     ------------------------------------------------------------  0
Sobic.004G129900.1     ------------------------------------------------------------  0
Sobic.004G129900.2     ------------------------------------------------------------  0
Sobic.006G042200.1     ------------------------------------------------------------  0
Sobic.004G129700.1     ------------------------------------------------------------  0
Sobic.004G129800.1     ------------------------------------------------------------  0
Sobic.004G129800.2     ------------------------------------------------------------  0
AT3G23550.1            --------------------------------------------MA--------------  2
AT3G23560.1            --------------------------------------------MA--------------  2
Manes.15G088100.1      ------------------------------------------MIMY--------DKKHQQ  10
Manes.03G109100.1      -----------------------------------------------------------M  1
Manes.03G109100.2      ------------------------------------------------------------  0
Sobic.002G311200.1     RLRQRGRGRAHDISTAAAAPRFL----VGSRRS-AAAAGSRRLLTGGHRRRPF-AASSGG  89
Sobic.002G006500.3     ------MG-----EEEAAAPLLV----P---------------------------GSNGD  18
Sobic.002G006500.4     ------MG-----EEEAAAPLLV----P---------------------------GSNGD  18
```

```
AT2G04090.1          ------------------------------------------------------------  0
AT2G04100.1          ------------------------------------------------------------  0
AT2G04066.1          ------------------------------------------------------------  0
AT2G04040.1          ------------------------------------------------------------  0
AT2G04080.1          ------------------------------------------------------------  0
AT2G04050.1          ------------------------------------------------------------  0
AT2G04070.1          ------------------------------------------------------------  0
Sobic.001G481800.1   ------------------------------------------------------------  0
Sobic.003G260200.1   --------------------------------------------MA--------------  2
Sobic.009G224800.1   ------------------------------------------------------------  0
Sobic.009G224800.2   ------------------------------------------------------------  0
AT1G66760.2          ------------------------------------------------------------  0
AT1G64820.1          --------------------------------------------ME--------------  2
AT1G66780.1          --------------------------------------------ME--------------  2
Manes.02G072400.1    ------------------------------------------------------------  0
Manes.01G113500.1    ------------------------------------------------------------  0
Manes.01G113500.2    ------------------------------------------------------------  0
AT1G15150.1          --------------------------------------------MQ--------------  2
AT1G15160.1          --------------------------------------------ME--------------  2
AT1G15170.1          --------------------------------------------MG--------------  2
AT1G15180.1          --------------------------------------------MG--------------  2
AT1G71140.1          ------------------------------------------------------------  0
Manes.06G063000.1    ------------------------------------------------------------  0
Manes.14G109700.1    --------------------------------------------ME--------------  2
Manes.14G109800.1    --------------------------------------------MG--------------  2
Sobic.008G171600.1   ------------------------------------------------------------  0
Sobic.008G171600.2   T-------------------------------------------YREQT-----------  14
Sobic.008G171600.3   ------------------------------------------------------------  0
AT3G59030.1          ------------------------------------------------------------  0
Manes.01G182000.1    ------------------------------------------------------------  0
Manes.02G142000.1    ------------------------------------------------------------  0
AT4G21903.2          --------------------------------------------MNGSN-----------  5
AT4G21910.4          --------------------------------------------MEVPS-----------  5
AT1G11670.1          ----------------------------------------------MGS-----------  3
AT1G61890.1          ------------------------------------------------------------  0
Manes.16G008000.1    ------------------------------------------------------------  0
Manes.17G038200.1    ------------------------------------------------------------  0
Sobic.002G318300.1   ------------------------------------------------------------  0
Manes.17G038300.1    ------------------------------------------------------------  0
Manes.16G007900.1    ------------------------------------------------------------  0
Manes.17G038400.1    ------------------------------------------------------------  0
Sobic.001G012600.1   ------------------------------------------------------------  0
Sobic.001G012600.2   ------------------------------------------------------------  0
Sobic.001G185600.1   ------------------------------------------------------------  0
Sobic.001G185400.1   ------------------------------------------------------------  0
AT3G21690.1          ------------------------------------------------------------  0
Sobic.001G185800.1   ------------------------------------------------------------  0
Sobic.001G185500.1   ------------------------------------------------------------  0
Sobic.001G185500.2   ------------------------------------------------------------  0
Sobic.007G165500.2   ---MQG------------------------------------------RRDGQRADADTD  15
AT4G00350.1          --------------------------------------------MEIPVREERRSSSSSA  16
Manes.01G255000.1    GWNQKGAKIANN-------------------------------KENDPLRAEGRSSSSSA  78
Sobic.004G349550.1   ------------------------------------------------------------  0
Sobic.004G349600.1   ------------------------------------------------------------  0
Sobic.007G176000.1   ------------------------------------------------------------  0
Sobic.007G176100.1   ------------------------------------------------------------  0
AT1G47530.1          ------------------------------------------------------------  0
Manes.05G164500.1    ------------------------------------------------------------  0
Manes.05G164500.2    ------------------------------------------------------------  0
Manes.18G030900.1    ------------------------------------------------------------  0
AT1G23300.1          ------------------------------------------------------------  0
AT3G26590.1          ------------------------------------------------------------  0
AT5G38030.1          ------------------------------------------------------------  0
Manes.12G023400.1    ------------------------------------------------------------  0
Manes.12G023500.1    ------------------------------------------------------------  0
Manes.12G023600.1    ------------------------------------------------------------  0
Manes.13G025200.1    ------------------------------------------------------------  0
Manes.13G025200.2    ------------------------------------------------------------  0
Sobic.001G273100.1   ------------------------------------------------------------  0
Sobic.001G273100.2   ------------------------------------------------------------  0
Sobic.001G273000.2   ------------------------------------------------------------  0
```

```
Sobic.001G273000.1      ------------------------------------------------------------  0
Sobic.001G273000.3      ------------------------------------------------------------  0
Manes.15G147800.1       ------------------------------------------------------------  0
Manes.15G147900.1       ------------------------------------------------------------  0
Manes.17G098800.1       ------------------------------------------------------------  0
Manes.17G098900.1       ------------------------------------------------------------  0
Manes.17G098900.2       ------------------------------------------------------------  0
Sobic.001G162400.1      ------------------------------------------------------------  0
Sobic.003G307600.2      ------------------------------------------------------------  0
Sobic.002G232200.1      ------------------------------------------------------------  0
Sobic.002G232500.1      ------------------------------------------------------------  0
Sobic.002G232600.1      ------------------------------------------------------------  0
Sobic.007G160700.1      ------------------------------------------------------------  0
Sobic.003G126200.2      --------------------------------------------MA--HGR---------  5
Sobic.001G476700.1      --------------------------------------------MA--STA---------  5
Sobic.001G476700.2      --------------------------------------------MA--STA---------  5
Manes.18G062800.1       ------------------------------------------------MVS---------  3
AT5G44050.1             --------------------------------------------MG--ERD---------  5
AT5G10420.1             --------------------------------------------MD--KKS---------  5
AT5G65380.1             ------------------------------------------------MRG---------  3
Manes.12G129000.1       ------------------------------------------------MPG---------  3
Manes.13G097900.1       ------------------------------------------------MPS---------  3
Sobic.005G020700.1      --------------------------------------------MDE-------------  3
Sobic.008G019400.1      --------------------------------------------MEKSTGG---------  7
Manes.09G135300.1       ------------------------------------------------------------  0
AT1G33080.1             ------------------------------------------------------------  0
AT1G33090.1             ------------------------------------------------------------  0
AT1G33100.1             ------------------------------------------------------------  0
AT1G33110.1             ------------------------------------------------------------  0
AT3G03620.1             ------------------------------------------------------------  0
AT5G17700.1             ------------------------------------------------------------  0
Manes.09G135200.1       ------------------------------------------------------------  0
Manes.09G134800.1       ------------------------------------------------------------  0
Manes.08G150400.1       ------------------------------------------------------------  0
Manes.09G134900.1       ------------------------------------------------------------  0
Manes.09G135000.1       ------------------------------------------------------------  0
Manes.09G134900.2       ------------------------------------------------------------  0
Manes.09G135000.2       ------------------------------------------------------------  0


Sobic.002G099300.1      ------------------------------------------------------------  0
AT4G39030.1             -RVTRNCVRIDR------------------------------------------------  74
Manes.04G084700.1       -QFRTACTAGSA-----------SEVTDDELGNV------SEY------GGSEALVSGSE  94
Manes.11G091900.1       -RLVTPSISPSQ-----------ELFDN----------VLS--END-ANTASDSF-CDEE  92
Sobic.004G019800.2      ------------------------------------------------------------  0
Sobic.004G019800.3      ------------------------------------------------------------  0
Sobic.004G019800.4      ------------------------------------------------------------  0
AT2G21340.1             -RFLRNCASTNQ-----------ELVVDGETGNG----SISELQGDAANGSISP--VEVE  96
Manes.11G092000.1       -RFMRNCINSNH-----------DVVYNDQEIER---------ENDSKIASVSSF-QEGE  94
Sobic.001G476700.3      ----------------------------------------MA-STAERADEEEE-ACRVA  18
Sobic.009G077000.1      ------------------------------------------------------------  0
Sobic.001G454900.1      -----------A--------------IPDDAAAA------AA-ATNATAGEENK-VE-EE  51
Sobic.003G403000.1      -----------A--------------APAENGDV------AA-AGAAENGTAAS-AA--N  69
Sobic.003G403000.2      -----------A--------------APAENGDV------AA-AGAAENGTAAS-AA--N  69
Sobic.003G403000.3      -----------A--------------APAENGDV------AA-AGAAENGTAAS-AA--N  69
Sobic.007G020600.1      ----------------------------------------------MMRGESES-PLLLH  13
Sobic.007G020600.2      ----------------------------------------------MMRGESES-PLLLH  13
AT3G08040.1             ------------------------------------------------------------  0
Manes.09G027700.1       -----------------------------------------------------------M  1
Manes.09G027800.1       ------------------------------------------------------------  0
Manes.09G027900.1       ------------------------------------------------------------  0
Manes.09G026900.1       -----------------------------------------------------------M  1
Manes.09G026900.2       -----------------------------------------------------------M  1
Manes.09G027000.1       ------------------------------------------------------------  0
Manes.07G006000.1       -----------------------------------------------------------M  1
Manes.07G006000.2       -----------------------------------------------------------M  1
Manes.10G143000.1       -----------------------------------------------------------M  1
AT1G51340.2             ----------------------------------------------------------MM  2
Manes.06G164500.1       -------------------------------------------------------MIDVA  5
Manes.06G164500.2       -------------------------------------------------------MIDVA  5
Manes.06G164500.3       -------------------------------------------------------MIDVA  5
```

```
Manes.06G164500.4       -------------------------------------------------------MIDVA  5
Manes.06G164500.5       -------------------------------------------------------MIDVA  5
Manes.14G002600.1       ------------------------------------------------------------  0
Sobic.008G006100.1      -RSPRCTRGG-G--------------KP----VV------TD------VVDEAA-PDKE-  71
Sobic.005G005400.1      --PPRCSGKQ-A-----------------------------A------VVEEET-PSPQ-  61
Sobic.005G005400.3      --PPRCSGKQ-A-----------------------------A------VVEEET-PSPQ-  61
Sobic.005G005400.2      --PPRCSGKQ-A-----------------------------A------VVEEET-PSPQ-  61
AT2G38330.1             ---PAVSTSS-Q--------------RPE----------------------------KQ-  53
Manes.08G096600.1       ---TDITSLE-N--------------SPPKPKPK------SS------LVDSPE-PSAP-  76
Manes.08G096600.2       ---TDITSLE-N--------------SPPKPKPK------SS------LVDSPE-PSAP-  76
AT4G38380.1             -----------Q--------------TNPDCGVV------K-------LGEEDD-SCSSL  86
Sobic.002G286800.1      ---VR------S--------------TGDGGGRV------GF------RGEDAE-GDRSP  100
Sobic.003G149300.2      ---IT------T--------------VASGQSVP------GY------TPDDGD-QCLET  84
Sobic.003G149300.1      ---IT------T--------------VASGQSVP------GY------TPDDGD-QCLET  84
Sobic.003G149300.3      ---IT------T--------------VASGQSVP------GY------TPDDGD-QCLET  84
Sobic.003G149300.4      ------------------------------------------------------------  0
Manes.01G153400.1       ------------------------------------------------------------  0
Manes.04G064900.1       ---PRLGVAFNQ--------------LSSGYGVE------SA------NVEE----RSSL  99
Manes.04G064900.2       ---PRLGVAFNQ--------------LSSGYGVE------SA------NVEE----RSSL  99
Sobic.001G003700.1      ------------------M--TTPPPP---------------------------------  7
AT4G22790.1             ------------------K--SESLDP---------------------------------  12
Manes.02G032300.1       TSKLEEN------PVVIAS--VSPSSP---------------------------------  24
Manes.10G000400.1       ------------------------------------------------------------  0
AT2G38510.1             ------------------------------------------------------------  0
Manes.08G172300.1       AQVLQG---------------LLRR-----------------------------------  18
Manes.09G117000.1       ------------------------------------------------------------  0
Sobic.010G167800.1      ------------------------AAVHGG------------------------------  41
AT5G19700.1             ---------------------HTNLLSKID------------------------------  17
AT4G29140.1             ---------------------TKRNLR-HC------------------------------  42
Manes.03G026500.1       ---------------------NLDNLK-TT------------------------------  44
Manes.16G109300.1       ---------------------NLDNLK-N-------------------------------  40
Sobic.007G181100.1      ---TKAC------PAL--------------------------------------------  27
Manes.13G127800.1       NSRKER-----------AK--TVPLDNQDS------------------------------  36
Sobic.001G446800.1      ------F------KAAAHQ--LLHPVD---------------------------------  34
AT5G52050.1             ---------------------TLPLLQ---------------------------------  17
AT1G58340.1             ---LETC------DTDNPH--YSEFRD---------------------------------  40
Manes.05G186700.1       ------C------FAKFNN--PMDLLS---------------------------------  31
Manes.18G054300.1       ------C------FAKSNK--TMDLCN---------------------------------  31
Sobic.001G320900.1      TTLTTTC------KDPP---------P---------------------------------  61
Sobic.001G320900.2      TTLTTTC------KDPP---------P---------------------------------  61
Sobic.004G283500.1      RAVKEAW------SVSL---------S---------------------------------  61
Sobic.006G184400.1      TAVKEAL------FVFL---------G---------------------------------  74
AT4G23030.1             --IIKN---------QT---------D---------------------------------  14
Manes.01G067000.1       ----GN---------L--------------------------------------------  35
Manes.02G027800.1       ----KN---------ST---------F---------------------------------  39
Manes.06G143600.1       R-QLEH---------KL---------P---------------------------------  47
Manes.14G029100.1       QQQEEH---------EL---------P---------------------------------  51
AT5G49130.1             ----------------------MVV-EEDS------------------------------  7
Manes.03G198000.1       ------------------------------------------------------------  0
Manes.15G011000.1       ------------------------------------------------------------  0
Sobic.001G019700.1      ----KG----------------GGQ-ADAD------------------------------  25
AT1G71870.1             ------------------------M-EDKI------------------------------  5
Manes.02G189800.1       ---------------------------MAD------------------------------  3
Manes.18G098800.1       ---------------------------MAD------------------------------  3
Sobic.010G256932.1      ----------------PSR--NGGA-SAGGAAEA--------------------------  15
Sobic.010G256700.1      ---------------------MGGA-GAGGEA----------------------------  10
Sobic.010G256700.5      ---------------------MGGA-GAGGEA----------------------------  10
Sobic.010G256700.6      ---------------------MGGA-GAGGEA----------------------------  10
Sobic.010G256700.4      ---------------------MGGA-GAGGEA----------------------------  10
Sobic.009G106900.1      ------------------------------------------------------------  0
Manes.S031500.1         -MD----------AQEPSL--KSPLISQEETLEE--------------------------  21
Manes.14G109900.1       MER----------GEKSGL--ESSLIPSQELDEE--------------------------  22
Manes.14G109900.2       ------------------------------------------------------------  0
Manes.14G109900.3       ------------------------------------------------------------  0
AT1G73700.1             ---------------------------------M--------------------------  1
AT2G34360.1             ------------------------------MREE--------------------------  4
AT5G52450.1             ---------------------------MRDDRER--------------------------  7
Sobic.009G106960.1      ------------------------------------------------------------  0
Manes.14G060800.1       DAE----------AFKSSL--ESPLISN--------------------------------  17
Sobic.009G106800.1      --M----------DNKAAT--EEPLLVPR-------------------------------  15
Sobic.007G074300.2      ------------------------------------------------------------  0
```

```
Sobic.009G106700.1      --M----------DGHAAV--HEPLLPSP-------------------------------  15
Sobic.010G138400.4      ------------------------------------------------------------  0
Sobic.010G138400.1      -------------------------MAKRPVEEA--------------------------  9
Sobic.010G138400.5      ------------------------------------------------------------  0
Sobic.004G129900.1      ------------------M--ENQ------------------------------------  4
Sobic.004G129900.2      ------------------------------------------------------------  0
Sobic.006G042200.1      --------------MMPSM--DEPLLGN--------------------------------  12
Sobic.004G129700.1      ------------------M--EEPLLSAPSTTEK--------------------------  14
Sobic.004G129800.1      ------------------M--E----------AA--------------------------  4
Sobic.004G129800.2      ------------------------------------------------------------  0
AT3G23550.1             DP-TSK------------------------------------------------------  7
AT3G23560.1             DP-ATS---------------SPLLDDH--------------------------------  14
Manes.15G088100.1       HPGEEE---------------IKLLSNASS------------------------------  25
Manes.03G109100.1       SPKETN---------------LE-------------------------------------  9
Manes.03G109100.2       ------------------------------------------------------------  0
Sobic.002G311200.1      TPQRAGGASGPSSDGGGSMSSSAPLLGAEE------------------------------  119
Sobic.002G006500.3      DPRRLNLTLAIHGDD-------DEWKAAGE------------------------------  41
Sobic.002G006500.4      DPRRLNLTLAIHGDD-------DEWKAAGE------------------------------  41
AT2G04090.1             ------------------M--EDPLLLGDDQ-----------------------------  11
AT2G04100.1             ------------------M--EDPLLLGDNQ-----------------------------  11
AT2G04066.1             ------------------------------------------------------------  0
AT2G04040.1             ------------------M--EEPFLLRDEL-----------------------------  11
AT2G04080.1             ------------------M--EEPFLPRDEQ-----------------------------  11
AT2G04050.1             ------------------M--EEPFLLQDEH-----------------------------  11
AT2G04070.1             ------------------M--EEPFLPQDEQ-----------------------------  11
Sobic.001G481800.1      --------------MGSSE--APLLLAHPGEGKE--------------------------  18
Sobic.003G260200.1      AAREE--------DEATQA--RPLLLPR--------------------------------  20
Sobic.009G224800.1      ----------------MEE--RVPLLPQYTLR-N--------------------------  15
Sobic.009G224800.2      ----------------MEE--RVPLLPQYTLR-N--------------------------  15
AT1G66760.2             -------------------------MKKSIETPL--------------------------  9
AT1G64820.1             TDF--------------------SLVRKEEEEEE--------------------------  16
AT1G66780.1             NGF--------------------SLVPKEEEEEE--------------------------  16
Manes.02G072400.1       ------------------M--EE--------GAG--------------------------  6
Manes.01G113500.1       ---------------------------------M--------------------------  1
Manes.01G113500.2       ---------------------------------M--------------------------  1
AT1G15150.1             DAE----------RT--------T------NDPV--------------------------  12
AT1G15160.1             DAE----------ST--------T------KDPV--------------------------  12
AT1G15170.1             DAE----------ST--K---DRL------LLPV--------------------------  15
AT1G15180.1             DAE----------ST-SK---TSL------LLPV--------------------------  16
AT1G71140.1             --------------M-DSA--EKGLLVVS-------------------------------  12
Manes.06G063000.1       ------------------M--EKQILLGE--AGE--------------------------  12
Manes.14G109700.1       --D----------SQ-KSL--QESLLPEQIEDKA--------------------------  21
Manes.14G109800.1       DPS----------EN-RNM--EETLLIKQ-NREV--------------------------  22
Sobic.008G171600.1      ---------MGEGS--IN--VIAPLLD----------------------I---------D  16
Sobic.008G171600.2      ---------MGEGS--IN--VIAPLLD----------------------I---------D  30
Sobic.008G171600.3      ---------MGEGS--IN--VIAPLLD----------------------I---------D  16
AT3G59030.1             ----------MSSTETYE--PLLTRLH----------------------SDSQI----TE  22
Manes.01G182000.1       ---------MGSVEPQYQ--SLLPTLD----------------------SHTRI----TH  23
Manes.02G142000.1       ---------MGSAEAEYQ--PLLPTLD----------------------SHSRI----PD  23
AT4G21903.2             ---------ET-VERRIE--LRRPLVD----------------------TEK------KL  25
AT4G21910.4             ---------E--TTNLAD--LRRPLVVP-------------------VVSERKP----PA  29
AT1G11670.1             ---------EA-TTAVNN--LQQPLLE-------------------------ST----KS  22
AT1G61890.1             ---------MN-SESLEN--LHRPLIE-------------------------SS----KS  19
Manes.16G008000.1       ---------ME-T--RSE--LHEPILGS-------------------L-HESTE----HY  22
Manes.17G038200.1       ---------ME-S--QDE--LQEPISQS-------------------PSFDPPA----HS  23
Sobic.002G318300.1      ----------------------------------------------MGRTGDDD----HR  10
Manes.17G038300.1       -----------------------------------------------MNSQSHC----FE  9
Manes.16G007900.1       --------------MMLS--HSQPL----------------------LQSFRSG----HE  18
Manes.17G038400.1       --------------MMLS--HSQPL----------------------LQSFRSG----HE  18
Sobic.001G012600.1      ----------------MD--STTPLLQP--------------------APH-------GG  15
Sobic.001G012600.2      ----------------MD--STTPLLQP--------------------APH-------GG  15
Sobic.001G185600.1      ---------MG-TA--SD--QNTPLLPAD--GGNDDGEGVAVLPGEAAGGAGGG----HG  40
Sobic.001G185400.1      ------------------------------MAGGDE---------AHG------------  9
AT3G21690.1             ---------MD-SSPNDG--VHQPLLH--------------------PQPSPSP----PE  24
Sobic.001G185800.1      -----------------------------MAGGGDG---------RQGGTSSSS----SH  18
Sobic.001G185500.1      ---------MG----ESK--LESPLLGLSAAASGGP---------SPGSGHGHG----GE  32
Sobic.001G185500.2      ------------------------------------------------------------  0
Sobic.007G165500.2      TPR-TPC-CIASASTGGA--GRTA-----------------AMGGD---AS---------  42
AT4G00350.1             GPL-QQTISLAADDAIDS--G----------PSSPLVVKVSVFETEHE-TT---------  53
Manes.01G255000.1       SSR-----SFPRENNVSR--GSVPLIDGVILSDSPVQDHISIFD-----TT---------  117
Sobic.004G349550.1      ------------------------------------------------------------  0
```

```
Sobic.004G349600.1      -------------------------------------MVVAVCDDDD--DD---------  12
Sobic.007G176000.1      ----------------------MAVSGG---------------GDDEM-LREAL----LA  18
Sobic.007G176100.1      ----------------------MSVTSGG----------HGEEEADDL-LKEAL----LA  23
AT1G47530.1             --------------MGKD--KTLPLLDPR-------------------------------  13
Manes.05G164500.1       --------------MAID----APLLDGY-------------------------------  11
Manes.05G164500.2       --------------MAID----APLLDGY-------------------------------  11
Manes.18G030900.1       --------------MGID----APLLDNH-------------------------------  11
AT1G23300.1             ------------------------------------------METLNV-DHEDT----IS  13
AT3G26590.1             ------------------------------------------MAKDKD-ITETL----LT  13
AT5G38030.1             ------------------------------------------MEEDKI-LTETL----LS  13
Manes.12G023400.1       ---------------------------------------MEDSRNQHQ-LHSLA----MS  16
Manes.12G023500.1       ---------------------------------------MEDSRNQHQ-LHSLA----IS  16
Manes.12G023600.1       ---------------MED--STKPLLSPSEDQN---QNHSHHHEDLDS-LLSLP----RS  35
Manes.13G025200.1       ---------------MAD--SREPLLSPKHDQN---YIV-QHHEDQDQ-LHSLL----IS  34
Manes.13G025200.2       ---------------MAD--SREPLLSPKHDQN---YIV-QHHEDQDQ-LHSLL----IS  34
Sobic.001G273100.1      ------------------------------------------MTS---------------  3
Sobic.001G273100.2      ------------------------------------------MTS---------------  3
Sobic.001G273000.2      ------------------------------------------------------------  0
Sobic.001G273000.1      ------------------------------------------------------------  0
Sobic.001G273000.3      ------------------------------------------------------------  0
Manes.15G147800.1       ------------------------------------------MEK---------------  3
Manes.15G147900.1       -------------------------------------------ME---------------  2
Manes.17G098800.1       ------------------------------------------MEE---------------  3
Manes.17G098900.1       ------------------------------------------MEE---------------  3
Manes.17G098900.2       ------------------------------------------------------------  0
Sobic.001G162400.1      ------------------------------------------------------------  0
Sobic.003G307600.2      ------------------------------------------------------------  0
Sobic.002G232200.1      -------------ME-SQ--SDVPLLPT-LPR----------------------------  15
Sobic.002G232500.1      -------------MEENR--SDIPLISG---S----------------------------  14
Sobic.002G232600.1      -------------MESQS--AVVPLI-----A----------------------------  12
Sobic.007G160700.1      -----------MEEEDQA--RAAPLLQQPLPLG---------------------------  20
Sobic.003G126200.2      ---------RKDDEEDCT--CTAALLLRGDDAE---------------------------  27
Sobic.001G476700.1      ---------ERADEEEEA--CRVALLDAGVKKE---------------------------  27
Sobic.001G476700.2      ---------ERADEEEEA--CRVALLDAGVKKE---------------------------  27
Manes.18G062800.1       ---------SNGGG----------------NLN---------------------------  11
AT5G44050.1             ---------DEAEG----------------ILE---------------------------  13
AT5G10420.1             ---------G-GTK----------------AIE---------------------------  12
AT5G65380.1             ---------G-DGE----------------EGS---------------------------  10
Manes.12G129000.1       ---------E-GAV----------------E-----------------------------  8
Manes.13G097900.1       ---------E-VAV----------------E-----------------------------  8
Sobic.005G020700.1      ---------QSSGGGEAA--AKVPLLEPRAAAAAEEEEHHHHHNG-------------AG  39
Sobic.008G019400.1      ---------ANGGEAAEE--AKVPLLQPRAAAVG--------------------------  30
Manes.09G135300.1       ------------------------------------------------------------  0
AT1G33080.1             ----------------------------------------------------------MA  2
AT1G33090.1             ----------------------------------------------------------MA  2
AT1G33100.1             ----------------------------------------------------------MA  2
AT1G33110.1             ----------------------------------------------------------MA  2
AT3G03620.1             -----------------------------------------------------------M  1
AT5G17700.1             ----------------------------------------------------------MS  2
Manes.09G135200.1       ------------------------------------------------------------  0
Manes.09G134800.1       ------------------------------------------------------------  0
Manes.08G150400.1       ------------------------------------------------------------  0
Manes.09G134900.1       ------------------------------------------------------------  0
Manes.09G135000.1       ------------------------------------------------------------  0
Manes.09G134900.2       ------------------------------------------------------------  0
Manes.09G135000.2       ------------------------------------------------------------  0


Sobic.002G099300.1      ------------------------------------------------------------  0
AT4G39030.1             EIDEEEE--EEE--------------KERGDLVKQSIWEQMKE----------IVKFTGP  108
Manes.04G084700.1       REYEKVE--VVR--------------SKRKELAGKSFWKQIKE----------IMMFSGP  128
Manes.11G091900.1       EKEEEMK--MEI--------------SSREGLENQSIWYQIKE----------IVKFTAP  126
Sobic.004G019800.2      ------------------------------------------------------------  0
Sobic.004G019800.3      ------------------------------------------------------------  0
Sobic.004G019800.4      ------------------------------------------------------------  0
AT2G21340.1             AEV-------EE--------------VKVDDLATQSIWGQMKE----------IVMFTGP  125
Manes.11G092000.1       ETE-------VE--------------VKREALENQTMWNQMRE----------IVMFTGP  123
Sobic.001G476700.3      LLDAGVKKEEWQVV---VGGGGDGDGGNNKQQLAARVWEESRK----------LWDIVAP  65
Sobic.009G077000.1      -------------M---VGG-QANI-FGSSDRVWAKLASRNYL----------L------  26
Sobic.001G454900.1      DAPEPAL--LACGP---RKT-GLHL-FVMNIRSVFKLDDLGSE----------VLRIAVP  94
Sobic.003G403000.1      GDGGGSE--LLGGP---RWT-GLHL-FVMNIRSVFKLDELGAE----------VLGIAVP  112
```

```
Sobic.003G403000.2      GDGGGSE--LLGGP---RWT-GLHL-FVMNIRSVFKLDELGAE----------VLGIAVP  112
Sobic.003G403000.3      GDGGGSE--LLGGP---RWT-GLHL-FVMNIRSVFKLDELGAE----------VLGIAVP  112
Sobic.007G020600.1      RSADAME--RGGEH---HHH-PLSV-FLRDARLAFRWDELGQE----------IMKIAVP  56
Sobic.007G020600.2      RSADAME--RGGEH---HHH-PLSV-FLRDARLAFRWDELGQE----------IMKIAVP  56
AT3G08040.1             MTETGDD--LATVK---KPI-PFLV-IFKDLRHVFSRDTTGRE----------ILGIAFP  43
Manes.09G027700.1       ATETLLN--LWENL---AKL-PLVV-LLKDTRNVFNMDELAVE----------IAQIAVP  44
Manes.09G027800.1       ------------------------------------------------------------  0
Manes.09G027900.1       ------------------------------------------------------------  0
Manes.09G026900.1       ATDTLVN--LWESL---AKL-PLVM-LFKDTRNVFNVDELAVE----------IAQIAVP  44
Manes.09G026900.2       ATDTLVN--LWESL---AKL-PLVM-LFKDTRNVFNVDELAVE----------IAQIAVP  44
Manes.09G027000.1       ------------------------------------------------------------  0
Manes.07G006000.1       AEDSALQ--LTERK---WKM-PLMV-FFRDARLIFKMDELGSE----------ILRVAVP  44
Manes.07G006000.2       AEDSALQ--LTERK---WKM-PLMV-FFRDARLIFKMDELGSE----------ILRVAVP  44
Manes.10G143000.1       AEDRALQ--QTERK---WKM-PLLV-FFRDARLVFKMDELGSE----------ILRIAVP  44
AT1G51340.2             SEDG-----YNTDF---PRN-PLYI-FFSDFRSVLKFDELGLE----------IARIALP  42
Manes.06G164500.1       QGDD-----PSMEK---KRT-PACI-FFNDFRHVLKLDELGLE----------IARIALP  45
Manes.06G164500.2       QGDD-----PSMEK---KRT-PACI-FFNDFRHVLKLDELGLE----------IARIALP  45
Manes.06G164500.3       QGDD-----PSMEK---KRT-PACI-FFNDFRHVLKLDELGLE----------IARIALP  45
Manes.06G164500.4       QGDD-----PSMEK---KRT-PACI-FFNDFRHVLKLDELGLE----------IARIALP  45
Manes.06G164500.5       QGDD-----PSMEK---KRT-PACI-FFNDFRHVLKLDELGLE----------IARIALP  45
Manes.14G002600.1       -----------------------------MSRHVLKLDEIGRD----------IARIALP  21
Sobic.008G006100.1      --PG-----IGIKG---EEE-KEDV-AGRGAQGWLRIDGVAAD----------ILAIAAP  109
Sobic.005G005400.1      -------------E---AKN-GEGE-QGRGPQGWFALDTIGLD----------ILSIALP  93
Sobic.005G005400.3      -------------E---AKN-GEGE-QGRGPQGWFALDTIGLD----------ILSIALP  93
Sobic.005G005400.2      -------------E---AKN-GEGE-QGRGPQGWFALDTIGLD----------ILSIALP  93
AT2G38330.1             --QN-----P--LT---SQN-K--P--DHDHKPDPGIGKIGME----------IMSIALP  86
Manes.08G096600.1       --SS-----PSLLN---SFS-G--L--AGRLRNGFKIDELGFE----------ILSIALP  111
Manes.08G096600.2       --SS-----PSLLN---SFS-G--L--AGRLRNGFKIDELGFE----------ILSIALP  111
AT4G38380.1             -D-K-----LPE-----VNG-VH--------TGVARPVDIKRE----------LVMLSLP  115
Sobic.002G286800.1      -AAR-----ASP-----LDG-AKGA-TPAPNVVRDHPGGIRKD----------LMNLAVP  137
Sobic.003G149300.2      -GNK-----------------ISFT-TVKDAVISLNSVGVRSE----------LILLALP  115
Sobic.003G149300.1      -GNK-----------------ISFT-TVKDAVISLNSVGVRSE----------LILLALP  115
Sobic.003G149300.3      -GNK-----------------ISFT-TVKDAVISLNSVGVRSE----------LILLALP  115
Sobic.003G149300.4      ------------------------------------------------------------  0
Manes.01G153400.1       ------------------------------------------------------------  0
Manes.04G064900.1       -EEE-----YSLIN---SRD-EHLD-STGVPIDQSHSSDVKRE----------LIMLSLP  138
Manes.04G064900.2       -EEE-----YSLIN---SRD-EHLD-STGVPIDQSHSSDVKRE----------LIMLSLP  138
Sobic.001G003700.1      ----------------------K-----HEPHGLAAVAAEVRA----------QRGIALP  30
AT4G22790.1             ----------------------EVSEGLCSKTLMQSIVHELKL----------QMRIGLP  40
Manes.02G032300.1       ----------------------SQNTQKWPTNLMQVLLLELKI----------QRGITLP  52
Manes.10G000400.1       ----------------------------------MQVVEELIL----------LGKIACP  16
AT2G38510.1             ----------------------------------MQVGEEMAS----------LTKIACP  16
Manes.08G172300.1       ----------------------LPLSSNIGRPPLNEVGEEVLA----------LGKIACP  46
Manes.09G117000.1       ----------------------------------MQVGEEMLA----------LGKIAGP  16
Sobic.010G167800.1      ---------------------RQLKCRTVPLPTAGETFREAVA----------LCRLAFP  70
AT5G19700.1             ---------------------LEKQNPAPIFPTITELKSEARS----------LFSLAFP  46
AT4G29140.1             ---------------------EN-----RGSPLMAEAVTEAKS----------LFTLAFP  66
Manes.03G026500.1       ---------------------QSSRPQPEIFPSVSDLISETKS----------LFKLAFP  73
Manes.16G109300.1       -----------------------SSPPPEIYPSVSDLISETKS----------LFKLAFP  67
Sobic.007G181100.1      ----------------------GDRPRPARRGDGGAPTEVIAS----------ILRLAVP  55
Manes.13G127800.1       ---------------------AAYCQYQAWSPSLSQAVDEIKQ----------LYTIAFP  65
Sobic.001G446800.1      ----------------------GDEGSGGHALQLSKVAGEARA----------IGRVSVP  62
AT5G52050.1             ----------------------KTSHLKNHSSVLSVFLNEAIS----------ICKISYP  45
AT1G58340.1             ----------------------TDSLDLKRWPSFLEGLEEVKA----------IGKISGP  68
Manes.05G186700.1       ----------------------DDEEELHRLPTPSEVLEEIKA----------LGKISGP  59
Manes.18G054300.1       ----------------------DDEEELHRWPTLSEVLEEIKA----------IGKISGP  59
Sobic.001G320900.1      ----------------------PVLD--DRPSLTRGAASEAAS----------ILSLSLP  87
Sobic.001G320900.2      ----------------------PVLD--DRPSLTRGAASEAAS----------ILSLSLP  87
Sobic.004G283500.1      ----------------------VTFPM-MPSMSAGAAGAEARS----------ILGLALP  88
Sobic.006G184400.1      ----------------------MAFPK-TPVVSSSDARGEARS----------ILGLALP  101
AT4G23030.1             ----------------------HRQDPNPNPTHLSSSIQEAKS----------IAKISLP  42
Manes.01G067000.1       ----------------------TSQPSEKTTTHLSLAINEAKS----------IANIALP  63
Manes.02G027800.1       ----------------------HSLKQNPEKTHLSLAINEAKC----------IANIALP  67
Manes.06G143600.1       ----------------------PQFQEPTHKTLFYLAIREAIC----------IAKIALP  75
Manes.14G029100.1       ----------------------PELQELPHKTHISLAIEEAMS----------IAKIALP  79
AT5G49130.1             ---------------------RLINLQHKYNPTMPEVVEELKR----------IWDISFP  36
Manes.03G198000.1       ---------------------MLSIENSQNYPTMPEVVDELKK----------MADIGFP  29
Manes.15G011000.1       ---------------------MLSTESSQKYPTMPEVVGELKK----------MTDLGFP  29
Sobic.001G019700.1      ---------------------DAAAWPGG-DDDQPSVVAELRA----------LWGMAAP  53
AT1G71870.1             ---------------------QSDDFTSHKNPTLPQVIEELKE----------LWAMVLP  34
Manes.02G189800.1       ---------------------KDPDFPSQKLPSASQVLEELKE----------LWGMALP  32
```

```
Manes.18G098800.1       ---------------------KDSDFPSQKLPSASQVLEELKE----------LWGMALP  32
Sobic.010G256932.1      -S-AASSPPI-LLSR---------------SAPRAAVGAEVRR----------QVGLAAP  47
Sobic.010G256700.1      ----EASSPL-LLPR---------------SAPRPAVGVEVRR----------QVGLAAP  40
Sobic.010G256700.5      ----EASSPL-LLPR---------------SAPRPAVGVEVRR----------QVGLAAP  40
Sobic.010G256700.6      ----EASSPL-LLPR---------------SAPRPAVGVEVRR----------QVGLAAP  40
Sobic.010G256700.4      ----EASSPL-LLPR---------------SAPRPAVGVEVRR----------QVGLAAP  40
Sobic.009G106900.1      ------------------------------------------------------------  0
Manes.S031500.1         ------------SRH---------------RFTKNEILEEVNK----------LLILAGP  44
Manes.14G109900.1       -EEEEEEEEL--CKQ---------------NCCRGDFIEEAKK----------QLWLAGP  54
Manes.14G109900.2       ------------------------------------------------------------  0
Manes.14G109900.3       ------------------------------------------------------------  0
AT1G73700.1             -EDGVTPPLL-ITEK---------------DTTMIRVKEEVKK----------QLWLSAP  34
AT2G34360.1             -REDMLSWPL-IGEK---------------EKRSRFVKEEVEK----------QLLLSGP  37
AT5G52450.1             -GEGDLSWPL-IGEK---------------SS----VKEEVKK----------QLWLSGP  36
Sobic.009G106960.1      ------------------------------------------------------------  0
Manes.14G060800.1       -----SENGIEPAKK---------------CYEKAEIISELKK----------QMNLAGP  47
Sobic.009G106800.1      ------------------------------P--EHTATSEAKR----------LLSLAGP  33
Sobic.007G074300.2      ------------------------------------------------------------  0
Sobic.009G106700.1      ------------------------------PLQGKAASAESKR----------LMRLAGP  35
Sobic.010G138400.4      ------------------------------------------------------------  0
Sobic.010G138400.1      -LLAA-----AGEPE---------------EEEILSVREELKK----------QLWLAGP  38
Sobic.010G138400.5      ------------------------------------------------------------  0
Sobic.004G129900.1      -HHAA---VARGGEE---------------SISSSSVWIETKK----------QLRLAAP  35
Sobic.004G129900.2      ------------------------------------------------------------  0
Sobic.006G042200.1      --------GVL-KTN---------------GVRESLVVAEVRK----------QLYLAGP  38
Sobic.004G129700.1      -LLHGVGGGKEE-EE---------------EESLALAVRETKQ----------QLYLAGP  47
Sobic.004G129800.1      -LLDATGTGKHGSNN---------------EAEAGAVVREVKK----------QLYLAGP  38
Sobic.004G129800.2      ------------------------------------------------------------  0
AT3G23550.1             ----DDHDGEGGRDK------------SSTFVQKLIDVEEAKT----------QIIYSLP  41
AT3G23560.1             ---VGGEDERGRRSR------------SSTLVQKVIDVEEAKA----------QMIYSLP  49
Manes.15G088100.1       --DGAGNEQEEEESR------------RKYWWKKVLDVKEAKK----------QILFSLP  61
Manes.03G109100.1       --ATPLLEPKDSGRC------------RRRWWKNVLDVEEAKK----------QMLVSLP  45
Manes.03G109100.2       ------------------------------------------------------MLVSLP  6
Sobic.002G311200.1      --AGGGEPAPAPRPS------------SWSWVERVVDTAEARA----------QLRFAVP  155
Sobic.002G006500.3      --AAASSARHHHALP------------GWDWA-------EVRG----------QLAFAAP  70
Sobic.002G006500.4      --AAASSARHHHALP------------GWDWA-------EVRG----------QLAFAAP  70
AT2G04090.1             -------------------LITRNLKSTPTW-WM-NFTAELKN----------VSSMAAP  40
AT2G04100.1             -------------------IITGSLKPTPTW-RM-NFTAELKN----------LSRMALP  40
AT2G04066.1             ------------------------------------------------------------  0
AT2G04040.1             -------------------LVPS----QVTW-HTNPLTVELKR----------VSRLAAP  37
AT2G04080.1             -------------------LVSC----KSTW-QSGQVTVELKK----------VSRLAAP  37
AT2G04050.1             -------------------LVPC----KDTW-KSGQVTVELKK----------VSSLAAP  37
AT2G04070.1             -------------------IVPC----KATW-KSGQLNVELKK----------VSRLAVP  37
Sobic.001G481800.1      -DPGADVGDRRRLRC-CWWWRRCGGASSEGW-WAE-VTAEAGR----------LAALAAP  64
Sobic.003G260200.1      ----------------------RPAQEDQKW-WRR-WAREAGW----------VGYLALP  46
Sobic.009G224800.1      -DD--------------GREEKCGGGGGVRW-WRELLAREAGK----------VGCVALP  49
Sobic.009G224800.2      -DD--------------GREEKCGGGGGVRW-WRELLAREAGK----------VGCVALP  49
AT1G66760.2             -LL--------NTKQ-SQDED-----------KEKIRWEKMKK----------VASMAAP  38
AT1G64820.1             -DN--------RN-------G-----------MSYLSMEMMKK----------VSSMAAP  39
AT1G66780.1             -DY--------SNEK-SEDQT-----------SYYLSTEMMKK----------VSFMAAP  45
Manes.02G072400.1       -RS--------REER-KW-AI-----------TRDGFVKEVKK----------TSCIAAP  34
Manes.01G113500.1       -ED--------REAK-------------------GWFMKELKK----------VSLLAAP  23
Manes.01G113500.2       -ED--------REAK-------------------GWFMKELKK----------VSLLAAP  23
AT1G15150.1             -DR--------IEKV-TWRDL-----------QDGSFTAELKR----------LICFAAP  41
AT1G15160.1             -DR--------VEKV-TWRDL-----------QDGSFTAELKK----------LICFAAP  41
AT1G15170.1             -ER--------VENV-TWSDL-----------RDGSFTVELKR----------LIFFAAP  44
AT1G15180.1             -ER--------VENV-TWRDL-----------RDGLFTAELKR----------LICFAAP  45
AT1G71140.1             -DR--------EEVN-KK----------------DGFLRETKK----------LSYIAGP  36
Manes.06G063000.1       -EK--------PQRI-SW----------------RVLTQEAKR----------TAYIAGP  36
Manes.14G109700.1       -QV--------SAFL-TW----------------DVFTEEGKK----------LFYIAGP  45
Manes.14G109800.1       -DH--------SSAL-TW----------------QVFFQEVKK----------LGFIAGP  46
Sobic.008G171600.1      ESSGASEVLLQQE------------PVPWGVLARLAA-WEAGN----------LWRISWA  53
Sobic.008G171600.2      ESSGASEVLLQQE------------PVPWGVLARLAA-WEAGN----------LWRISWA  67
Sobic.008G171600.3      ESSGASEVLLQQE------------PVPWGVLARLAA-WEAGN----------LWRISWA  53
AT3G59030.1             RSSPEIEEFLRRR----------GSTVTPRWWLKLAV-WESKL----------LWTLSGA  61
Manes.01G182000.1       LSSQAIEEFLQQR------------TVPLRWWPRLVA-WESRL----------LWLLSWA  60
Manes.02G142000.1       LSSQAIEEFLEQT------------TVPLRWWPRLVA-WESRL----------LWLLSWA  60
AT4G21903.2             PLEVGLESVLT------------ESSLPYRRRVYLGMCIELKL----------LLRLALP  63
AT4G21910.4             DVGLGLESVLT------------ERSLPYRRRVYLGACIEMKL----------LFRLALP  67
AT1G11670.1             EADFRMESVLT------------DTHLSYFRRIYLASLIEMKY----------LFHLAAP  60
AT1G61890.1             FVDYRLETVLT------------DRELPYFRRIYLAMMIEMKF----------LFHLAAP  57
```

```
Manes.16G008000.1       EVKSELEKVLT------------DTQLPYFKRLRIASWIELKL----------LFPLAGP  60
Manes.17G038200.1       EIDSRLENVLN------------DDKLPYFSRLRLASWIELKQ----------LFHLAAP  61
Sobic.002G318300.1      VVDCRLEALLS----------GAASEAPWLRRMASATALELRL----------LAPLAAP  50
Manes.17G038300.1       GVGSELEEILT------------DTHSSTVKRVGSATWVELKL----------LFKLAAP  47
Manes.16G007900.1       AVTSELEDILN------------DTRTSTFKRLRSATWVELKL----------LFQLAAP  56
Manes.17G038400.1       AVSSELEDILN------------DTHTPTFKRLRSASCVELKL----------LFKLAAP  56
Sobic.001G012600.1      GGSRELEAILE------------DASVPWARRALRGAGVELPL----------LLRIALP  53
Sobic.001G012600.2      GGSRELEAILE------------DASVPWARRALRGAGVELPL----------LLRIALP  53
Sobic.001G185600.1      GVSAQLERILA------------DESVPRARRLARAARVELRL----------LVALAAP  78
Sobic.001G185400.1      GASGRLESILTAEA-------DASSPWPWARRAWAAASIELRL----------LTRLAAP  52
AT3G21690.1             STNGELETVLS------------DVETPLFLRLRKATIIESKL----------LFNLAAP  62
Sobic.001G185800.1      ELSGQLEGILA------------DREAPWARRASKAAMIELRL----------LAPIAAP  56
Sobic.001G185500.1      AASGQLESILS------------DTSLPWGRRMAAASVVEMRL----------LVRLAAP  70
Sobic.001G185500.2      ------------------------------------------------------------  0
Sobic.007G165500.2      TIEGA-PLLGGRQASHEHDTP----PVRTAGDAARMVWDESKR----------LWGIGLP  87
AT4G00350.1             KLIHA-PSTLLGETTGDADFP----PIQSFRDAKLVCVVETSK----------LWEIAAP  98
Manes.01G255000.1       DLHPA-PSALVHNE--VGDYP----PIQSFEDAKYICLLESSK----------LWAIAAP  160
Sobic.004G349550.1      ------------------------------------------------------------  0
Sobic.004G349600.1      DDEEE-AGALVAAIGSTRDAP----AVHSPRAAWAVFVKESRR----------LWSIAAP  57
Sobic.007G176000.1      AGNGN-GNGSFSKGG--EDLE----EIRSVGSFLRHAAAENRK----------LWYLAGP  61
Sobic.007G176100.1      AGNGS-GSDGGIDGE--EDLE----EIRSVGSFLRHAAEENRK----------LWYLAGP  66
AT1G47530.1             ------------EPP--ELTG----TKSASKVWAKEFGEESKR----------LWELAGP  45
Manes.05G164500.1       ------------AKS--DH-Q----ENK-YISVVRDFNEESKR----------LWKLAGP  41
Manes.05G164500.2       ------------AKS--DH-Q----ENK-YISVVRDFNEESKR----------LWKLAGP  41
Manes.18G030900.1       ------------PNG--SD-Q----EQKFSISVVREFWEESKR----------LWKLAGP  42
AT1G23300.1             SE--Q-EHRAHTKSD--TDMP----PISGGRDFIRQFAAESKK----------LWWLAGP  54
AT3G26590.1             AAE-E-RSDLPFLSV--DDIP----PITTVGGFVREFNVETKK----------LWYLAGP  55
AT5G38030.1             AAE-E-PPALPFSSV--EDIP----PITTVGGFVKEFNVEVKK----------LWYLAGP  55
Manes.12G023400.1       TTN-S-NPSVPDSHR--RDIP----PINTISVFFREFYRESKK----------LWCLACP  58
Manes.12G023500.1       KTN-S-KSSVPDAHG--HDIP----PINSISVFFREFYREFKK----------LWCLACP  58
Manes.12G023600.1       TTS-T-ISFVPDA----DDIS----PINGVSDFFREFYVESKK----------LWYLAGP  75
Manes.13G025200.1       TTN-T-ISFVPDA----DDIP----PINGIRDFLREFYIEFKK----------LWYLAGP  74
Manes.13G025200.2       TTN-T-ISFVPDA----DDIP----PINGIRDFLREFYIEFKK----------LWYLAGP  74
Sobic.001G273100.1      -SSGS-SSRRLEQHEAEKPAHPHGLDALTMKMLVRRSWEESRL----------LLRLAFP  51
Sobic.001G273100.2      -SSGS-SSRRLEQHEAEKPAHPHGLDALTMKMLVRRSWEESRL----------LLRLAFP  51
Sobic.001G273000.2      --------------------------MLIMKRLVSQSWEES-------RLLWRLT---FP  24
Sobic.001G273000.1      -------------------------MHLRYKDKAQTPWKKMNKFEIPFCVVWSLVDGALV  35
Sobic.001G273000.3      -------------------------MHLRYKDKAQTPWKKMNKFEIPFCVVWSLVDGALV  35
Manes.15G147800.1       -TDD-----SVQEKHVDYET-------TSWKNIVKKSWIESKK----------MWEIAAP  40
Manes.15G147900.1       -END-----DLQENHFEMDM-----QMMGRKTTVKKSWDESKK----------MWEIAAP  41
Manes.17G098800.1       -KDD-----SVQEKSLGIEMQVI-NGVMGKKKMVKKSWDESKK----------IWEIAAP  46
Manes.17G098900.1       -KDD-----SVQEKRFGIEMQVI-NGVMGKKKMVKKSWDESKK----------MWEIAAP  46
Manes.17G098900.2       ------------------------------------------------------------  0
Sobic.001G162400.1      -------------------MSSSSMAMGKGKAAVSKSLQESKL----------LWHIAFP  31
Sobic.003G307600.2      ------------------------------------------------------------  0
Sobic.002G232200.1      ---------PPPE-----KR--GGGKIQFQGRLGREVWEESKK----------LGAVVGP  49
Sobic.002G232500.1      ---------ELPD-----RR--GGGKIS---ELAKEVWGESKK----------LWVVAGP  45
Sobic.002G232600.1      ---------ELPE-----KR--GGKT------LVEEVWEESKK----------LWEVTGP  40
Sobic.007G160700.1      ----DF----VDD-----EVGGGAATTKAAGRAAVEWWVESKK----------LWHIVGP  57
Sobic.003G126200.2      ----AWKEEEWHH-----HHHHHHASSSDLWRWMRRVREESRK----------LWEVVGP  68
Sobic.001G476700.1      ----EWQVVVGGG-----GDGDG---GNNKQQLAARVWEESRK----------LWDIVAP  65
Sobic.001G476700.2      ----EWQVVVGGG-----GDGDG---GNNKQQLAARVWEESRK----------LWDIVAP  65
Manes.18G062800.1       ----EALLPHG---------GVDKHDQENARELASRVWIETKK----------LWQIVGP  48
AT5G44050.1             ----KAKIPLLKD---------QNVAEEENGEIKKEIWLETKK----------LWRIVGP  50
AT5G10420.1             ----EATVPLL----------ECHNAAEEGGGMKREIWIETKK----------IWYIVGP  48
AT5G65380.1             ----ESRVALLK--------SP-HTAEEDGEGLKDRILVETKK----------LWQIVGP  47
Manes.12G129000.1       ----ESKVLLLGD-----HFGPRLEEDDQDQSLTKRVWIESKK----------LWQIVGP  49
Manes.13G097900.1       ----ESKVPLLGD-----HFTPKPVEDDQDQRLTKRVWIESKK----------LWQIVGP  49
Sobic.005G020700.1      GGGGGAAAA--------VVVGKSDEAEWSAQPLRRRAWEENKR----------LWVVAGP  81
Sobic.008G019400.1      GGGGDSKAAAEEED---RAAVAAEDASWSTLPLRRRAWEENKK----------LWVVAGP  77
Manes.09G135300.1       -MEEEIKQ---K------LLIEAEKPEHDEVPFKDKLWTETKK----------MWIVAGP  40
AT1G33080.1             RREGEVTETLLK------KSTENRGEDRDGLGMKEKVWRESKK----------LWVVAGP  46
AT1G33090.1             GEGGELTAALLK------KTTENGGEENDELGLKEKVWIESKK----------LWVVAAP  46
AT1G33100.1             GRGGELTEALVK------KT---GREEEDELGMKEKVWIESKK----------LWVVAAP  43
AT1G33110.1             GGGGELTAALLK------KTAENGGEEKDELGLKQKVWIESKK----------LWIVAAP  46
AT3G03620.1             STQEEMEERLLREG----SDAEGQSNNRESIYLRTKVWSEVNK----------MWRIALP  47
AT5G17700.1             GGGGEMEERLLNGS--------ETEQRRESLYLRKKIWSEVRK----------MWRIALP  44
Manes.09G135200.1       MDEAIMQERLLR------------TEELETANLTSRIWTESKK----------TWRVAFP  38
Manes.09G134800.1       ----MEEVEPSNS-----------QEKDSNEELKKRVWEESKK----------IWRIAFP  35
Manes.08G150400.1       ------MEERLIE-----------SEAKDINDLKKRIWAENKK----------IWKVGFP  33
Manes.09G134900.1       -----MEEGLLRS-----------QERDHNNDLKGRIWEENKK----------IWKVGFP  34
```

```
Manes.09G135000.1      -----MEERLLRS-----------EEKDHNNDLKGRIWEENKK----------IWKVGFP  34
Manes.09G134900.2      ------MEGWI-----------------------SFYISKSNT----------IWNVCSN  21
Manes.09G135000.2      ------MEGRI-----------------------SCNVSKSHT----------IWNVCGN  21


Sobic.002G099300.1     ------------------------------------------------------------  0
AT4G39030.1            AMGMWICGPLMSLIDTVVIGQGSS-I-EL-AALGPGTVLCDHMSYVFMF--LSVATSNMV  163
Manes.04G084700.1      ATGLWICGPLMSLISTAVIGRGSS-T-EL-AALGPGTVFCDNMNLLFMF--LSIATSNMV  183
Manes.11G091900.1      ATGLWICGPLMSLIDTAVIGQGSS-L-EL-AALGPGTVLCDNMSYVFMF--LSISTSNMV  181
Sobic.004G019800.2     ----------MSLIDTMVIGQTSA-L-QL-AALGPGTVFCDYLSYIFMF--LSVATSNMV  45
Sobic.004G019800.3     ----------MSLIDTMVIGQTSA-L-QL-AALGPGTVFCDYLSYIFMF--LSVATSNMV  45
Sobic.004G019800.4     ----------MSLIDTMVIGQTSA-L-QL-AALGPGTVFCDYLSYIFMF--LSVATSNMV  45
AT2G21340.1            AAGLWLCGPLMSLIDTAVIGQGSS-L-EL-AALGPATVICDYLCYTFMF--LSVATSNLV  180
Manes.11G092000.1      ATGLWLCGPLMSLIDTAVIGQGSS-I-EL-AALGPGTVVCDYMSYVFMF--LSVATSNLV  178
Sobic.001G476700.3     AIFSRVVTYSMNVITQAFAGHLGD-L-EL-AAI----SIANTVVVGFSFGLMVSRCQVS-  117
Sobic.009G077000.1     -------------LRQ---NQIGM-A-WLNIELVASIAVYNQVPRIAIFPLDSATTSFVV  68
Sobic.001G454900.1     ASLALAADPLASLVDTAFIGRLGS-V-EI-AAVGVSIAIFNQVSKVCIYPLVSVTTSFVA  151
Sobic.003G403000.1     ASLALTADPLASLIDTAFIGRLGS-V-EI-AAVGVAIAVFNQVMKVCIYPLVSVTTSFVA  169
Sobic.003G403000.2     ASLALTADPLASLIDTAFIGRLGS-V-EI-AAVGVAIAVFNQVMKVCIYPLVSVTTSFVA  169
Sobic.003G403000.3     ASLALTADPLASLIDTAFIGRLGS-V-EI-AAVGVAIAVFNQVMKVCIYPLVSVTTSFVA  169
Sobic.007G020600.1     GALALMADPVASLVDTAFIGHIGP-V-EL-GAVGVSIAVFNQVSRIAVFPLVSVTTSFVA  113
Sobic.007G020600.2     GALALMADPVASLVDTAFIGHIGP-V-EL-GAVGVSIAVFNQVSRIAVFPLVSVTTSFVA  113
AT3G08040.1            AALALAADPIASLIDTAFVGRLGA-V-QL-AAVGVSIAIFNQASRITIFPLVSLTTSFVA  100
Manes.09G027700.1      AALALAADPVASLIDTAFIGHLGP-V-EL-AAVGVSIAIFNQVSKIAIFPLVSVTTSFVA  101
Manes.09G027800.1      ------------------------------------------------------------  0
Manes.09G027900.1      ------------------------------------------------------------  0
Manes.09G026900.1      AALALAADPVASLIDTAFIGHLGP-V-EL-AAVGVSIAIFNQMSKIAIFPLVSVTTSFVA  101
Manes.09G026900.2      AALALAADPVASLIDTAFIGHLGP-V-EL-AAVGVSIAIFNQMSKIAIFPLVSVTTSFVA  101
Manes.09G027000.1      ------------------------------------------------------------  0
Manes.07G006000.1      AAMALAADPIASLIDTAFIGHLGP-V-EI-AAVGVSIAIFNQASKVTIFPLVSITTSFVA  101
Manes.07G006000.2      AAMALAADPIASLIDTAFIGHLGP-V-EI-AAVGVSIAIFNQASKVTIFPLVSITTSFVA  101
Manes.10G143000.1      AAMALAADPIASLIDTAFIGHLGP-V-EI-AAVGVSIAIFNQASKVTIFPLVSITTSFVA  101
AT1G51340.2            AALALTADPIASLVDTAFIGQIGP-V-EL-AAVGVSIALFNQVSRIAIFPLVSITTSFVA  99
Manes.06G164500.1      AALALTADPIASLVDTAFIGQIGP-V-EL-AAVGVSIALFNQVSRIAIFPLVSITTSFVA  102
Manes.06G164500.2      AALALTADPIASLVDTAFIGQIGP-V-EL-AAVGVSIALFNQVSRIAIFPLVSITTSFVA  102
Manes.06G164500.3      AALALTADPIASLVDTAFIGQIGP-V-EL-AAVGVSIALFNQVSRIAIFPLVSITTSFVA  102
Manes.06G164500.4      AALALTADPIASLVDTAFIGQIGP-V-EL-AAVGVSIALFNQVSRIAIFPLVSITTSFVA  102
Manes.06G164500.5      AALALTADPIASLVDTAFIGQIGP-V-EL-AAVGVSIALFNQVSRIAIFPLVSITTSFVA  102
Manes.14G002600.1      AALALTADPIASMMDTAFIGQIGP-V-EL-AAVGVSIALFNQVSRIAIFPLVSVTTSFVA  78
Sobic.008G006100.1     AVLALAADPITALVDTAFVGHIGS-A-QL-AAVGASTSIFNLVSKLFNVPLLNVTTSFVA  166
Sobic.005G005400.1     AALALAADPIAALVDTAFVGHIGS-A-EL-AAVGVSISVFNLVSKLFNVPLLNVTTSFVA  150
Sobic.005G005400.3     AALALAADPIAALVDTAFVGHIGS-A-EL-AAVGVSISVFNLVSKLFNVPLLNVTTSFVA  150
Sobic.005G005400.2     AALALAADPIAALVDTAFVGHIGS-A-EL-AAVGVSISVFNLVSKLFNVPLLNVTTSFVA  150
AT2G38330.1            AALALAADPITSLVDTAFVGHIGS-A-EL-AAVGVSVSVFNLVSKLFNVPLLNVTTSFVA  143
Manes.08G096600.1      AALALAADPITSLVDTAFVGHIGP-V-EL-AAVGVSVSAFNLVSKLFNVPLLNVTTSFVA  168
Manes.08G096600.2      AALALAADPITSLVDTAFVGHIGP-V-EL-AAVGVSVSAFNLVSKLFNVPLLNVTTSFVA  168
AT4G38380.1            AIAGQAIDPLTLLMETAYIGRLGS-V-EL-GSAGVSMAIFNTISKLFNIPLLSVATSFVA  172
Sobic.002G286800.1     AIVGQAIDPVAQLLETAYVGRLGP-V-EL-GSAAVGMSVFNIISKLFNIPLLSITTSFVA  194
Sobic.003G149300.2     AVLGQAIDPMAQLMETAYIGRLGA-L-EL-ASAGIGISIFNIVSKIFNIPLLSIATSFVA  172
Sobic.003G149300.1     AVLGQAIDPMAQLMETAYIGRLGA-L-EL-ASAGIGISIFNIVSKIFNIPLLSIATSFVA  172
Sobic.003G149300.3     AVLGQAIDPMAQLMETAYIGRLGA-L-EL-ASAGIGISIFNIVSKIFNIPLLSIATSFVA  172
Sobic.003G149300.4     ---------MAQLMETAYIGRLGA-L-EL-ASAGIGISIFNIVSKIFNIPLLSIATSFVA  48
Manes.01G153400.1      ------------------------------------------------------------  0
Manes.04G064900.1      AIAGQAIDPLSQLMETAYIGRLGP-V-EL-GSAGVSITIFNNISKLFNIPLLSVATSFVA  195
Manes.04G064900.2      AIAGQAIDPLSQLMETAYIGRLGP-V-EL-GSAGVSITIFNNISKLFNIPLLSVATSFVA  195
Sobic.001G003700.1     LIGMNLTWFAKQAVTTAFLGRLGD-L-QL-AAGTLGYSFANVTGFAVL------------  75
AT4G22790.1            LVVMNLLWFGKMTTTSVFLGRQGE-L-NL-AGGSLGFSFANVTGFSVL------------  85
Manes.02G032300.1      LLAMNLTWFAKIAITTAFLGRLGQ-L-PL-AGGTLGFTFANVTGFSVL------------  97
Manes.10G000400.1      MAISSLVLHSKSIISMLFLGHLGD-I-EL-AGGSLAIGFANITGYSVI------------  61
AT2G38510.1            IVMTSLLIFSRSIISMWFLSHLGK-V-EL-AGGALAMGFGNITGVSVL------------  61
Manes.08G172300.1      IILTTILIYSRSVISMLFLSRMGK-I-EL-AGGSLALGFANITGLSVM------------  91
Manes.09G117000.1      IILTTMLIYSRSVISMLFLSRMGK-K-EL-AGGSLALGFANITGLSVM------------  61
Sobic.010G167800.1     IALTALLLYSRTALSMLFLGSIGD-L-PL-AAGSLAVGFANITGYSVL------------  115
AT5G19700.1            TILAALILYARSAISMLFLGHIGE-L-EL-AGGSLAIAFANITGYSVL------------  91
AT4G29140.1            IAVTALVLYLRSAVSMFFLGQLGD-L-EL-AAGSLAIAFANITGYSVL------------  111
Manes.03G026500.1      IALTSLILYSRSILSMLFLGHLGD-I-EL-AAGSLAIAFANITGYSVL------------  118
Manes.16G109300.1      IALTAIILYARSIVSMLFLGRLGD-L-EL-AAGSLAIAFANITGYSVL------------  112
Sobic.007G181100.1     MVGAGLLMYMRSLVSMLFLGTLGR-L-PL-AGGSLALGFANITGYSVL------------  100
Manes.13G127800.1      MIITGLLIYGKSAISMFFMGKLGK-D-VL-AGGSLSIGIANISGYSVI------------  110
Sobic.001G446800.1     MAVTGLVMYSRALISMLFLGRLGE-L-AL-AGGSLALGFANITGYSVL------------  107
AT5G52050.1            LVLTGLFLYVRSFVSLSFLGGLGD-A-TL-AGGSLAAAFANITGYSLF------------  90
```

```
AT1G58340.1          TAMTGLLMYSRAMISMLFLGYLGE-L-EL-AGGSLSIGFANITGYSVI------------  113
Manes.05G186700.1    TAITGVILYSRAMISMLFLGYLGE-L-EL-AGGSLSIGFANITGYSVI------------  104
Manes.18G054300.1    TVLTGLILYSRAMISMLFLGYLGE-L-EL-AGGSLSIGFANITGYSVI------------  104
Sobic.001G320900.1   MIMTGLILYVRPMISMLFLGRLGE-L-AL-AGGSLAIGFGNITGYSVL------------  132
Sobic.001G320900.2   MIMTGLILYVRPMISMLFLGRLGE-L-AL-AGGSLAIGFGNITGYSVL------------  132
Sobic.004G283500.1   MILTGLLLYLRSMISMLFLGRLGG-L-AL-AGGSLAIGFANITGYSVL------------  133
Sobic.006G184400.1   MILTGLLLYLRSMISMLFLGRLGG-L-AL-AGGSLAIGFANITGYSVL------------  146
AT4G23030.1          LILTGLLLYSRSMISMLFLGRLND-LSAL-SGGSLALGFANITGYSLL------------  88
Manes.01G067000.1    MILTGLLLYSRSMISMLFLGRLGE-L-AL-AGGSLAIGFANITGYSIL------------  108
Manes.02G027800.1    MILTGLLLYSRSMISMLFLGRLGD-L-AL-AGGSLAIGFANITGYSIL------------  112
Manes.06G143600.1    MILTGLLLYSRSMISMLFLGRLGE-L-AL-AGGSLAIGFANITGYSIL------------  120
Manes.14G029100.1    LILTGLVLYSRSMISMMFLGRLGD-L-AL-AGGSLAIGFANITGYSVL------------  124
AT5G49130.1          VAAMSILNYLKNMTSVVCMGRLGS-L-EL-AGGALAIGFTNITGYSVL------------  81
Manes.03G198000.1    IAAMSLVGYLKNMILVACMGRLGS-L-EL-AGGALAIGFTNITGYSVL------------  74
Manes.15G011000.1    IAAMSLVGYLKNMILVVCMGKLGS-L-EL-AGGALAIGFTNITGYSVL------------  74
Sobic.001G019700.1   ITALNCVVYLRAMVSVLCLGRLGP-L-DL-AGGALAIGLTNITGHSVL------------  98
AT1G71870.1          ITAMNCLVYVRAVVSVLFLGRLGS-L-EL-AGGALSIGFTNITGYSVM------------  79
Manes.02G189800.1    ITAAHLMAFFRAVVSVMFLGRLGS-L-EL-AGGALSIGFTNITGYSVL------------  77
Manes.18G098800.1    ITAAHLMAFFRAVVSSIFLGRLGS-L-EL-AGGALSIGFTNITGYSVL------------  77
Sobic.010G256932.1   LVACSLLQYSLQL---------RE-L-SL-SGASIAPSFANVTGFSVL------------  83
Sobic.010G256700.1   LVACSLLQYSLQVVSVMFAGHLGE-L-SL-SSASVAASFANVTGFSVL------------  85
Sobic.010G256700.5   LVACSLLQYSLQVVSVMFAGHLGE-L-SL-SSASVAASFANVTGFSVL------------  85
Sobic.010G256700.6   LVACSLLQYSLQVVSVMFAGHLGE-L-SL-SSASVAASFANVTGFSVL------------  85
Sobic.010G256700.4   LVACSLLQYSLQVVSVMFAGHLGE-L-SL-SSASVAASFANVTGFSVL------------  85
Sobic.009G106900.1   ------------------------------------------------------------  0
Manes.S031500.1      LISASFFTFLLQTISVMFVGHLGE-L-AL-SGASMATSFASMTGFSIL------------  89
Manes.14G109900.1    LIAVSMLQYCLQVISVMFVGHLGE-L-AL-SSASMASSFGSVTGFSVL------------  99
Manes.14G109900.2    ------------------------------------------------------------  0
Manes.14G109900.3    ------------------------------------------------------------  0
AT1G73700.1          LIGVSLLQYSLQVISVMFVGHLGS-L-PL-SAASIATSFASVTGFTFL------------  79
AT2G34360.1          LIAVSLLQFCLQIISVMFVGHLGS-L-PL-SAASIATSFASVTGFTFL------------  82
AT5G52450.1          LIAVSLLQFCLQVISVMFVGHLGS-L-PL-SAASIATSFASVTGFSFL------------  81
Sobic.009G106960.1   ------------------------------------------------------------  0
Manes.14G060800.1    LVLVSFLQYSLQMISVMFVGHLGE-L-SL-SSASMATSFAGVTGFALM------------  92
Sobic.009G106800.1   LVASCILQNVVQLVSVMFVGHLGE-L-PL-AGASLASSLANVTGFSLL------------  78
Sobic.007G074300.2   ------------------------------------------------------------  0
Sobic.009G106700.1   IVASCVLQNVVNMASVMFVGHLGE-L-PL-AGASLATSLANVTGYSLL------------  80
Sobic.010G138400.4   -------------------------------------------MRKAK------------  5
Sobic.010G138400.1   MIGGALLQNVIQMISVMYVGHLGE-L-PL-AGASMANSFATVTGLSLL------------  83
Sobic.010G138400.5   ----------------------------------------------MQ------------  2
Sobic.004G129900.1   LAAGFLLQKVIQTISIMFVGRLGE-L-PL-AGASLATSFASVTGFSLL------------  80
Sobic.004G129900.2   ------------------------------------------------------------  0
Sobic.006G042200.1   LIAAWILQNIVQMISVMFVGHLGE-L-AL-SSASIATSFAGVTGFSLL------------  83
Sobic.004G129700.1   LVVGFLLQNLVQMVSVMFVGHLGE-L-AL-ASASLATSFAGVTGFSLL------------  92
Sobic.004G129800.1   LVVGFLLQNMVQMVSVMFVGHLGE-L-AL-ASASLATSFAGVTGFSLL------------  83
Sobic.004G129800.2   ------------------------------------------------------------  0
AT3G23550.1          MIFTNLFYYCIPLTSVMFASQLGQ-L-EL-AGATLANSWATVTGFAFM------------  86
AT3G23560.1          MILTNVFYYCIPITSVMFASHLGQ-L-EL-AGATLANSWATVSGFAFM------------  94
Manes.15G088100.1    MILTNVFYYLIPLVSVMFAGHLGE-L-EL-AGATLANSWATVTGFAFM------------  106
Manes.03G109100.1    MILTTFFYYSISLVSVMFAGRLGD-L-EL-AAATLAFSWANITGYNFT------------  90
Manes.03G109100.2    MILTTFFYYSISLVSVMFAGRLGD-L-EL-AAATLAFSWANITGYNFT------------  51
Sobic.002G311200.1   MVVTSMAYYGIPLVSVMFSGHLGDDV-HL-AGATLGNSWATVTGYAFV------------  201
Sobic.002G006500.3   MVTTNMAYYAIPLVSVMYAGRLGD-L-QL-AAATLGNSWGTVTGIALM------------  115
Sobic.002G006500.4   MVTTNMAYYAIPLVSVMYAGRLGD-L-QL-AAATLGNSWGTVTGIALM------------  115
AT2G04090.1          MATVTVSQYLLPVISVMVAGHCGE-L-QL-SGVTLATAFANVSGFGIM------------  85
AT2G04100.1          MATVTVAQYLLPVISVMVAGHRSE-L-QL-SGVALATSFTNVSGFSVM------------  85
AT2G04066.1          ------------------------------------------------------------  0
AT2G04040.1          MATVTIAQYLLPVISVMVAGHNGE-L-QL-SGVALANSFTNVTGFSIM------------  82
AT2G04080.1          MATVTIAQYLLPVISVMVAGHIGE-L-EL-AGVALATSFTNVSGFSIM------------  82
AT2G04050.1          MAAVTIAQYLLPVISVMVAGHNGE-L-QL-SGVALATSFTNVSGFSIL------------  82
AT2G04070.1          MATVTIAQYLLPVISVMVAGHNGE-L-QL-SGVALATSFTNVSGFSIM------------  82
Sobic.001G481800.1   MIAVALLQLMMQLISTIMVGHL-GEV-PL-AGAAIANSLTNVSGFSVL------------  109
Sobic.003G260200.1   MVVVNLSQYAVQVSSNMMVGHLPGVL-PL-SSAAIATSLANVSGFSLL------------  92
Sobic.009G224800.1   MAAVSVSQYAVQVASNMMVGHLPGVL-PL-SASAIATSLATVSGFSLL------------  95
Sobic.009G224800.2   MAAVSVSQYAVQVASNMMVGHLPGVL-PL-SASAIATSLATVSGFSLL------------  95
AT1G66760.2          MVAVNMSQYLLQATSTMIVGHRSE-L-AL-AGIALGSSFANVTGFGVL------------  83
AT1G64820.1          MVAVSVSQFLLQVISMVMAGHLDE-L-SL-SAVAIATSLTNVTGFSLI------------  84
AT1G66780.1          MVAVAASQYLLQVISIVMAGHLDE-L-SL-SAVAIATSLTNVTGFSLI------------  90
Manes.02G072400.1    MVAVSVLQYLLQVVSVIMVGHLGDQL-AL-SSVAIATSLTNVVGFSLL------------  80
Manes.01G113500.1    MVVVSVSQNLLPPISLMMAGHLGE-L-HL-SAVSVATSFTNATGFALL------------  68
Manes.01G113500.2    MVVVSVSQNLLPPISLMMAGHLGE-L-HL-SAVSVATSFTNATGFALL------------  68
```

| | | |
|---|---|---|
| AT1G15150.1 | MAAVVIIQFMIQIISMVMVGHLGR-L-SL-ASASFAVSFCNVTGFSFI------------ | 86 |
| AT1G15160.1 | MAAVVITQSMLQIITMVIVGHLGR-L-SL-ASASFAISFCNVTGFSFI------------ | 86 |
| AT1G15170.1 | MAAVVIAQFMLQIVSMMMVGHLGN-L-SL-ASASLASSFCNVTGFSFI------------ | 89 |
| AT1G15180.1 | MAAVVIAQFMLQIISMVMVGHLGN-L-SL-ASASLASSFCNVTGFSFI------------ | 90 |
| AT1G71140.1 | MIAVNSSMYVLQVISIMMVGHLGE-L-FL-SSTAIAVSFCSVTGFSVV------------ | 81 |
| Manes.06G063000.1 | MVAVNLSQYFLQIISIMMVGHLGQ-L-SL-SSTAIAISFCGVTGFSLL------------ | 81 |
| Manes.14G109700.1 | MVTVTLSLYLINVISMMMVGHLGE-L-AL-SSSAIAISLSAVTGFSLM------------ | 90 |
| Manes.14G109800.1 | MVAVILSQFLVQFISMTMVGHLSE-L-AL-SSTAIAISLSSVTGLSPL------------ | 91 |
| Sobic.008G171600.1 | SILTTLFSFTLSLVTQMFVGHLGE-L-EL-AGASITNIGIQGLAYGVM------------ | 98 |
| Sobic.008G171600.2 | SILTTLFSFTLSLVTQMFVGHLGE-L-EL-AGASITNIGIQGLAYGVM------------ | 112 |
| Sobic.008G171600.3 | SILTTLFSFTLSLVTQMFVGHLGE-L-EL-AGASITNIGIQGLAYGVM------------ | 98 |
| AT3G59030.1 | SIVVSVLNYMLSFVTVMFTGHLGS-L-QL-AGASIATVGIQGLAYGIM------------ | 106 |
| Manes.01G182000.1 | SIIVSLFNFMLSFVTQMFSGHLGA-V-EL-AGASIANVGIQGLAYGIM------------ | 105 |
| Manes.02G142000.1 | SIVVSIFNFMLSFVTQMFAGHLGA-V-EL-AGASIANVGIQGLAYGIM------------ | 105 |
| AT4G21903.2 | AILVYLINGGMGISARIFAGHLGS-T-QL-AAASIGNSSFS-LVYALM------------ | 107 |
| AT4G21910.4 | AILVYLVNSGMGISARIFAGHLGK-N-EL-AAASIGNSCFS-LVYGLM------------ | 111 |
| AT1G11670.1 | AIFVYVINNGMSMLTRIFAGRLGS-M-QL-AAASLGNSGFNMFTLGLM------------ | 105 |
| AT1G61890.1 | AIFVYVINNGMSILTRIFAGHVGS-F-EL-AAASLGNSGFNMFTYGLL------------ | 102 |
| Manes.16G008000.1 | AVFVYMINNFMSQSTRVFAGHLGN-L-EL-AAASLGNGGIQLFAYGLM------------ | 105 |
| Manes.17G038200.1 | AVFVYMINNLMSLSTRAFAGHLGN-L-EL-AAASLGNSGVQLFAYGLM------------ | 106 |
| Sobic.002G318300.1 | AVVVYMLIIVMSSTTQIVCGQLGN-V-QL-AAASLGNNGIQVFAYGLM------------ | 95 |
| Manes.17G038300.1 | AVIVYLLNNVVSMSTQIFCGHLGN-L-QL-AAVSLGNTGIQVFAYGVM------------ | 92 |
| Manes.16G007900.1 | AVIVYLLNNVVSMSTQIFCGHLGT-L-QL-AAVSLGNTGIQVFAYGLM------------ | 101 |
| Manes.17G038400.1 | AVIVYLLNNVVSMSTQIFCGHLGT-L-QL-AAVSLGNTGIQVFAYGLM------------ | 101 |
| Sobic.001G012600.1 | AVAVYMINYLMSMSTQIFCGQLGN-L-EL-AAVSLGNTGIQVFAYGLM------------ | 98 |
| Sobic.001G012600.2 | AVAVYMINYLMSMSTQIFCGQLGN-L-EL-AAVSLGNTGIQVFAYGLM------------ | 98 |
| Sobic.001G185600.1 | AVAVYMINYSMSLSTRIFCGQLGT-L-EL-AAASLGNVGIQVFAYGLM------------ | 123 |
| Sobic.001G185400.1 | AVVMYMINYLMSMSTQIFSGHLGN-L-EL-AAASLGNTGIQIFAYGLM------------ | 97 |
| AT3G21690.1 | AVIVYMINYLMSMSTQIFSGHLGN-L-EL-AAASLGNTGIQVFAYGLM------------ | 107 |
| Sobic.001G185800.1 | AVVVYVLNNVLSISTQIFSGHLGN-L-EL-AASSLGNNGIQVFAYGLM------------ | 101 |
| Sobic.001G185500.1 | AVLVYMINYLMSMSTQIFSGHLGT-L-EL-AAASLGNTGIQVFAYGLM------------ | 115 |
| Sobic.001G185500.2 | ------------------------------------------------------------ | 0 |
| Sobic.007G165500.2 | IAVGMLSMYAISSITQMFIGHLGN-L-PL-AAASIGLSVFSTFALGFL------------ | 132 |
| AT4G00350.1 | IAFNILCNYGVNSFTSIFVGHIGD-L-EL-SAVAIALSVVSNFSFGFL------------ | 143 |
| Manes.01G255000.1 | IAFNILCNYGVNSFTNIFVGHIGN-V-EL-SAVAIALSVIANFSFGFL------------ | 205 |
| Sobic.004G349550.1 | ------------------------------------------------------------ | 0 |
| Sobic.004G349600.1 | IAFNIMCMYGTNSTTQIFAGHIGN-R-EL-SAVAIGLSVVSNFSFGFL------------ | 102 |
| Sobic.007G176000.1 | AIFTSIAQYSLGAITLVFAGHLTT-L-EL-DAFSTENNVIAGLALGIT------------ | 106 |
| Sobic.007G176100.1 | AILTSIAQYSLGAITQVFAGHLTT-L-EL-DAISTENNVVAGLAFGIM------------ | 111 |
| AT1G47530.1 | AIFTAISQYSLGALTQTFSGRLGE-L-EL-AAVSVENSVISGLAFGVM------------ | 90 |
| Manes.05G164500.1 | AIFTAICQYSLGALTQTFAGLVGE-L-EL-AAVSVENSVIAGLAFGVM------------ | 86 |
| Manes.05G164500.2 | AIFTAICQYSLGALTQTFAGLVGE-L-EL-AAVSVENSVIAGLAFGVM------------ | 86 |
| Manes.18G030900.1 | AIFTALCQYSLGALTQTFAGLVGE-I-EL-AAVSVENSVIAGLAFGVM------------ | 87 |
| AT1G23300.1 | AIFTSFCQYSLGAVTQILAGHVNT-L-AL-AAVSIQNSVISGFSVGIM------------ | 99 |
| AT3G26590.1 | AIFTSVNQYSLGAITQVFAGHIST-I-AL-AAVSVENSVVAGFSFGIM------------ | 100 |
| AT5G38030.1 | AIFMSITQYSLGAATQVFAGHIST-I-AL-AAVSVENSVIAGFSFGVM------------ | 100 |
| Manes.12G023400.1 | AIFTTICQYSLGAITQVFSGQVST-L-AL-AAVSVGNSVIAGFSFGVM------------ | 103 |
| Manes.12G023500.1 | AIFTSICQYSLGAITQVFSGQVST-L-AL-AAVSIENSLIAGFSFGVM------------ | 103 |
| Manes.12G023600.1 | AIFTSICQYSLGAITQVFSGQVGT-L-AL-AAVSVENSVIAGFSFGAM------------ | 120 |
| Manes.13G025200.1 | AIFTCICQYSLGAVTQIFSGQVGT-L-AL-AAVSVENSVISGFSFGAM------------ | 119 |
| Manes.13G025200.2 | AIFTCICQYSLGAVTQIFSGQVGT-L-AL-AAVSVENSVISGFSFGAM------------ | 119 |
| Sobic.001G273100.1 | ALLTEVFQFSIGFVTTAFVGHLGE-V-EL-AAVSVVENILDSSAYGVL------------ | 96 |
| Sobic.001G273100.2 | ALLTEVFQFSIGFVTTAFVGHLGE-V-EL-AAVSVVENILDSSAYGVL------------ | 96 |
| Sobic.001G273000.2 | VLLAEVFQFSIGFVTTAFVGHLGD-V-EL-AAVTVAENILDTSAYGLL------------ | 69 |
| Sobic.001G273000.1 | LHCTEVFQFSIGFVTTAFVGHLGD-V-EL-AAVTVAENILDTSAYGLL------------ | 80 |
| Sobic.001G273000.3 | LHCTEVFQFSIGFVTTAFVGHLGD-V-EL-AAVTVAENILDTSAYGLL------------ | 80 |
| Manes.15G147800.1 | AMITAVTHFSIGFVTCAFVGHLGE-L-EL-AAVSIVENVIETFADGIM------------ | 85 |
| Manes.15G147900.1 | AMITTVTQFSIGFVTSAFVGHLGE-M-EL-AAVSIVHNVIEGFVDGVM------------ | 86 |
| Manes.17G098800.1 | AMLTAVAQFSIGFVTSAFVGHLGE-V-EL-AAVSIVQNVIEGFVYGVM------------ | 91 |
| Manes.17G098900.1 | ALLTAVAQFSIGFVTSAFVGHLGE-V-EL-AAVSIVQNVIEGFVYGVM------------ | 91 |
| Manes.17G098900.2 | ------------------------------------------------------------ | 0 |
| Sobic.001G162400.1 | AILTAVFQFSIGFVTVAFVGHIGS-V-EL-AAVTVVENVIEGFAYGVL------------ | 76 |
| Sobic.003G307600.2 | ------------------------------------------------------------ | 0 |
| Sobic.002G232200.1 | AVFMNLVFSSMNLVSQSFAGHLGD-L-DL-AAFSIANTVVDGFNFAML------------ | 94 |
| Sobic.002G232500.1 | AAFTRLTFYGMTVVSQAFAGHIGD-L-EL-AAFSIATTVISGLSFGFF------------ | 90 |
| Sobic.002G232600.1 | AAFTGMVLYSMTIVSQAFAGHLGD-R-HL-AAFSIANTVISGLNFGIL------------ | 85 |
| Sobic.007G160700.1 | AIFQRIALYGVNVVSQAFIGHLGD-L-EL-AAFSIAATVVAGFNFGFL------------ | 102 |
| Sobic.003G126200.2 | AIFTRAAIYSLNVVMQAVAGHLGD-L-EL-ASVSFACTVLTGFNYGLM------------ | 113 |
| Sobic.001G476700.1 | AIFSRVVTYSMNVITQAFAGHLGD-L-EL-AAISIANTVVVGFSFGLM------------ | 110 |
| Sobic.001G476700.2 | AIFSRVVTYSMNVITQAFAGHLGD-L-EL-AAISIANTVVVGFSFGLMG----HDSPVAA | 118 |
| Manes.18G062800.1 | AIFSRIASFSMNIITQAFAGHLGD-V-QL-ASISIANTVIVGFNFGLL------------ | 93 |

```
AT5G44050.1             AIFTRVTTNLIFVITQAFAGHLGE-L-EL-AAISIVNNVIIGFNYSLF------------  95
AT5G10420.1             SIFTGLATYSILIITQAFAGHLGD-L-EL-AAISIINNFTLGFNYGLL------------  93
AT5G65380.1             AIFSRVTTTYSMLVITQAFAGHLGD-L-EL-AAISIVNNVTVGFNFGLL------------  92
Manes.12G129000.1       AIFSRLTSYSMLVITQAFAGHLGD-L-EL-AAISIANNVIVGFDFGLL------------  94
Manes.13G097900.1       AIFSRLTSYSMLVITQAFAGHLGD-L-EL-AAISIANNVIVGLDFGLL------------  94
Sobic.005G020700.1      SICARFASFGVTVISQAFIGHIGA-T-EL-AAYALVSTVLMRFSNGVL------------  126
Sobic.008G019400.1      SIFTRFASFGVTVISQAFIGHIGA-T-EL-AAYALVSTVLMRFSNGIL------------  122
Manes.09G135300.1       AIFTRFSTFGISVISQAFVGHIGS-T-EL-AGYSLVVTVLLRFANGIL------------  85
AT1G33080.1             AIFTRFSTSGLSLISQAFIGHLGS-T-EL-AAYSITLTVLLRFSNGIL------------  91
AT1G33090.1             SIFTKFSTYGVSLVTQGFVGHIGP-T-EL-AAYSITFTVLLRFSNGIL------------  91
AT1G33100.1             AIFTRYSTFGVSMVTQAFIGHLGP-T-EL-AAYSITFTILLRFSNGIL------------  88
AT1G33110.1             AIFTRFSTFGVSIISQSFIGHLGP-I-EL-AAYSITFTVLLRFSNGIL------------  91
AT3G03620.1             SSLFRMTSFGSIIVAQAFIGHSSE-L-GL-AAYALLQSTFIRFLYGLM------------  92
AT5G17700.1             STLFRVMSFGCVVVAQAFIGHSSE-T-GL-AAYALLQSTFIRFIYGIM------------  89
Manes.09G135200.1       AMITKVTYFGMIVVTQSFIGHISE-L-QL-AAYALEQTFFVRFVNGIL------------  83
Manes.09G134800.1       SMLTKITQFGMLVVTQGFIGHIGE-V-EL-AAFALVQIISVRFVQGIV------------  80
Manes.08G150400.1       ATLARVSQYGMFVVTQSFIGHVGE-L-EL-AGYALIQIITVRFANGIL------------  78
Manes.09G134900.1       STLARVTQFGMFVVTQAFIGHVGE-L-QL-AGYALIQIITVRFANGIL------------  79
Manes.09G135000.1       ATLARVTQYGMFVVTQAFIGHFGE-L-QL-AGYALIQIVIIRFANGIL------------  79
Manes.09G134900.2       TSIH-WTRWGTA---TCWL---------------CTHPDHHRSICKWD------------  50
Manes.09G135000.2       TSIH-WTFWGTT---ACRL---------------CTHPDRHHSICKWD------------  50


Sobic.002G099300.1      ------------------------------------------------------------  0
AT4G39030.1             ATSLAKQD-KK-------------------------------------------------  173
Manes.04G084700.1       ATSLAKRD-KK-------------------------------------------------  193
Manes.11G091900.1       ATSLAKQD-KN-------------------------------------------------  191
Sobic.004G019800.2      ATSLAKKD-EE-------------------------------------------------  55
Sobic.004G019800.3      ATSLAKKD-EE-------------------------------------------------  55
Sobic.004G019800.4      ATSLAKKD-EE-------------------------------------------------  55
AT2G21340.1             ATSLARQD-KD-------------------------------------------------  190
Manes.11G092000.1       ATSLARRD-KN-------------------------------------------------  188
Sobic.001G476700.3      ----SVLKPSRERE--KGVERTQDKRRCGCGRS--------------------HA-----  146
Sobic.009G077000.1      -DEDLLSPFYISN*----------------------------------------------  80
Sobic.001G454900.1      -EEDAIISKAIEEKSSQDLE-----------------------KASH-----------VD  176
Sobic.003G403000.1      -EEDAVLSKGGAKVIDNGEEEEE-------------------LEAGQVGPEKHTAAAGAD  209
Sobic.003G403000.2      -EEDAVLSKGGAKVIDNGEEEEE-------------------LEAGQVGPEKHTAAAGAD  209
Sobic.003G403000.3      -EEDAVLSKGGAKVIDNGEEEEE-------------------LEAGQVGPEKHTAAAGAD  209
Sobic.007G020600.1      -EEDAMSNCRDNDKINQENE-------CS-------------------VSV--------S  138
Sobic.007G020600.2      -EEDAMSNCRDNDKINQENE-------CS-------------------VSV--------S  138
AT3G08040.1             -EEDTMEKMKEEANKANLVHAE-------------TILVQDSLEKGISSPT--------S  138
Manes.09G027700.1       -EEESAGKSSNDENAS--------------------------LEDGLL-VN--------K  125
Manes.09G027800.1       ------------------------------------------------------------  0
Manes.09G027900.1       ------------------------------------------------------------  0
Manes.09G026900.1       -EEDSAGKSSTKEDAS--------------------------LEDGSV-VN--------K  125
Manes.09G026900.2       -EEDSAGKSSTKEDAS--------------------------LEDGSV-VN--------K  125
Manes.09G027000.1       ------------------------------------------------------------  0
Manes.07G006000.1       -EEDTAQKMSNEPQKGEEFEKKDSAKTCEMKELVPEDVMLENLEKGSAEDT--------E  152
Manes.07G006000.2       -EEDTAQKMSNEPQKGEEFEKKDSAKTCEMKELVPEDVMLENLEKGSAEDT--------E  152
Manes.10G143000.1       -EEDTVQRVSKEPQEVENLEKKDSGKTS----VVKEDVMLENLEKGSATDT--------E  148
AT1G51340.2             -EEDACSSQQDTVRDHKEC-----------------------IEIGINNPT--------E  127
Manes.06G164500.1       -EEDTIGKVSTEAQE-SES-----------------------LEAGSL-VN--------S  128
Manes.06G164500.2       -EEDTIGKVSTEAQE-SES-----------------------LEAGSL-VN--------S  128
Manes.06G164500.3       -EEDTIGKVSTEAQE-SES-----------------------LEAGSL-VN--------S  128
Manes.06G164500.4       -EEDTIGKVSTEAQE-SES-----------------------LEAGSL-VN--------S  128
Manes.06G164500.5       -EEDTIGKVSTEAQE-SES-----------------------LEAGSL-VN--------S  128
Manes.14G002600.1       -EEDTIGKLSPEAQE-SES-----------------------LETGSH-VN--------S  104
Sobic.008G006100.1      -EQQAMDGNSNITRERDEF-----------------------------------------  184
Sobic.005G005400.1      -EQQAVDDDSAGSREGYDF-----------------------------------------  168
Sobic.005G005400.3      -EQQAVDDDSAGSREGYDF-----------------------------------------  168
Sobic.005G005400.2      -EQQAVDDDSAGSREGYDF-----------------------------------------  168
AT2G38330.1             -EEQAIAAKDDN------------------------------------------------  154
Manes.08G096600.1       -EEQALISKEKD------------------------------------------------  179
Manes.08G096600.2       -EEQALISKEKD------------------------------------------------  179
AT4G38380.1             -EDIAKIAAQDLASEDS-------------------------------------------  188
Sobic.002G286800.1      -EDVSKHDSSKSAS----------------------------------------------  207
Sobic.003G149300.2      -EDISRSATKHPSS----------------------------------------------  185
Sobic.003G149300.1      -EDISRSATKHPSS----------------------------------------------  185
Sobic.003G149300.3      -EDISRSATKHPSS----------------------------------------------  185
Sobic.003G149300.4      -EDISRSATKHPSS----------------------------------------------  61
Manes.01G153400.1       ------------------------------------------------------------  0
```

```
Manes.04G064900.1       -EDISRNEIKDSPSEQNVQEN---------------------------------------  215
Manes.04G064900.2       -EDISRNEIKDSPSEQNVQEN---------------------------------------  215
Sobic.001G003700.1      --------------------------------------------TGLC------------  79
AT4G22790.1             --------------------------------------------YGIS------------  89
Manes.02G032300.1       --------------------------------------------NGLC------------  101
Manes.10G000400.1       --------------------------------------------KGLA------------  65
AT2G38510.1             --------------------------------------------KGLS------------  65
Manes.08G172300.1       --------------------------------------------KGLA------------  95
Manes.09G117000.1       --------------------------------------------KGLA------------  65
Sobic.010G167800.1      --------------------------------------------SGLS------------  119
AT5G19700.1             --------------------------------------------AGLA------------  95
AT4G29140.1             --------------------------------------------SGLA------------  115
Manes.03G026500.1       --------------------------------------------SGLA------------  122
Manes.16G109300.1       --------------------------------------------SGLS------------  116
Sobic.007G181100.1      --------------------------------------------SGLA------------  104
Manes.13G127800.1       --------------------------------------------SGLA------------  114
Sobic.001G446800.1      --------------------------------------------SGLA------------  111
AT5G52050.1             --------------------------------------------SGLT------------  94
AT1G58340.1             --------------------------------------------SGLS------------  117
Manes.05G186700.1       --------------------------------------------SGLA------------  108
Manes.18G054300.1       --------------------------------------------SGLA------------  108
Sobic.001G320900.1      --------------------------------------------SGLA------------  136
Sobic.001G320900.2      --------------------------------------------SGLA------------  136
Sobic.004G283500.1      --------------------------------------------SGLA------------  137
Sobic.006G184400.1      --------------------------------------------SGLA------------  150
AT4G23030.1             --------------------------------------------SGLS------------  92
Manes.01G067000.1       --------------------------------------------SGLA------------  112
Manes.02G027800.1       --------------------------------------------SGLA------------  116
Manes.06G143600.1       --------------------------------------------SGLA------------  124
Manes.14G029100.1       --------------------------------------------SGLA------------  128
AT5G49130.1             --------------------------------------------SGLA------------  85
Manes.03G198000.1       --------------------------------------------SGLA------------  78
Manes.15G011000.1       --------------------------------------------SGLA------------  78
Sobic.001G019700.1      --------------------------------------------FGLA------------  102
AT1G71870.1             --------------------------------------------VGLA------------  83
Manes.02G189800.1       --------------------------------------------VGLA------------  81
Manes.18G098800.1       --------------------------------------------VGLA------------  81
Sobic.010G256932.1      --------------------------------------------G---------------  84
Sobic.010G256700.1      --------------------------------------------LGMG------------  89
Sobic.010G256700.5      --------------------------------------------LGMG------------  89
Sobic.010G256700.6      --------------------------------------------LGMG------------  89
Sobic.010G256700.4      --------------------------------------------LGMG------------  89
Sobic.009G106900.1      ------------------------------------------------------------  0
Manes.S031500.1         --------------------------------------------RGTG------------  93
Manes.14G109900.1       --------------------------------------------LGMG------------  103
Manes.14G109900.2       ----------------------------------------------MG------------  2
Manes.14G109900.3       ----------------------------------------------MG------------  2
AT1G73700.1             --------------------------------------------LGTA------------  83
AT2G34360.1             --------------------------------------------MGTA------------  86
AT5G52450.1             --------------------------------------------MGTA------------  85
Sobic.009G106960.1      ------------------------------------------------------------  0
Manes.14G060800.1       --------------------------------------------LGMG------------  96
Sobic.009G106800.1      --------------------------------------------AGMA------------  82
Sobic.007G074300.2      ------------------------------------------------------------  0
Sobic.009G106700.1      --------------------------------------------TGMA------------  84
Sobic.010G138400.4      --------------------------------------------LGMA------------  9
Sobic.010G138400.1      --------------------------------------------LGMA------------  87
Sobic.010G138400.5      --------------------------------------------LGMA------------  6
Sobic.004G129900.1      --------------------------------------------TGMA------------  84
Sobic.004G129900.2      ----------------------------------------------MA------------  2
Sobic.006G042200.1      --------------------------------------------SGMA------------  87
Sobic.004G129700.1      --------------------------------------------AGMA------------  96
Sobic.004G129800.1      --------------------------------------------AGMA------------  87
Sobic.004G129800.2      ----------------------------------------------MA------------  2
AT3G23550.1             --------------------------------------------TGLS------------  90
AT3G23560.1             --------------------------------------------VGLS------------  98
Manes.15G088100.1       --------------------------------------------TGLS------------  110
Manes.03G109100.1       --------------------------------------------AGLS------------  94
Manes.03G109100.2       --------------------------------------------AGLS------------  55
Sobic.002G311200.1      --------------------------------------------TGLS------------  205
Sobic.002G006500.3      --------------------------------------------TGLS------------  119
```

| | | |
|---|---|---|
| Sobic.002G006500.4 | --------------------------------------------TGLS------------ | 119 |
| AT2G04090.1 | --------------------------------------------YGLV------------ | 89 |
| AT2G04100.1 | --------------------------------------------FGLA------------ | 89 |
| AT2G04066.1 | ------------------------------------------------------------ | 0 |
| AT2G04040.1 | --------------------------------------------CGLV------------ | 86 |
| AT2G04080.1 | --------------------------------------------FGLV------------ | 86 |
| AT2G04050.1 | --------------------------------------------FGLA------------ | 86 |
| AT2G04070.1 | --------------------------------------------FGLV------------ | 86 |
| Sobic.001G481800.1 | --------------------------------------------MGLA------------ | 113 |
| Sobic.003G260200.1 | --------------------------------------------IGLA------------ | 96 |
| Sobic.009G224800.1 | --------------------------------------------IGMA------------ | 99 |
| Sobic.009G224800.2 | --------------------------------------------IGMA------------ | 99 |
| AT1G66760.2 | --------------------------------------------FGLS------------ | 87 |
| AT1G64820.1 | --------------------------------------------VGFA------------ | 88 |
| AT1G66780.1 | --------------------------------------------FGLA------------ | 94 |
| Manes.02G072400.1 | --------------------------------------------SGMA------------ | 84 |
| Manes.01G113500.1 | --------------------------------------------FGLA------------ | 72 |
| Manes.01G113500.2 | --------------------------------------------FGLA------------ | 72 |
| AT1G15150.1 | --------------------------------------------IGLS------------ | 90 |
| AT1G15160.1 | --------------------------------------------MGLS------------ | 90 |
| AT1G15170.1 | --------------------------------------------IGLS------------ | 93 |
| AT1G15180.1 | --------------------------------------------VGLS------------ | 94 |
| AT1G71140.1 | --------------------------------------------FGLA------------ | 85 |
| Manes.06G063000.1 | --------------------------------------------FGMS------------ | 85 |
| Manes.14G109700.1 | --------------------------------------------CGMA------------ | 94 |
| Manes.14G109800.1 | --------------------------------------------MGMA------------ | 95 |
| Sobic.008G171600.1 | --------------------------------------------IGMA------------ | 102 |
| Sobic.008G171600.2 | --------------------------------------------IGMA------------ | 116 |
| Sobic.008G171600.3 | --------------------------------------------IGMA------------ | 102 |
| AT3G59030.1 | --------------------------------------------LGMA------------ | 110 |
| Manes.01G182000.1 | --------------------------------------------LGMA------------ | 109 |
| Manes.02G142000.1 | --------------------------------------------LGMA------------ | 109 |
| AT4G21903.2 | --------------------------------------------LGMG------------ | 111 |
| AT4G21910.4 | --------------------------------------------LGMG------------ | 115 |
| AT1G11670.1 | --------------------------------------------LGMG------------ | 109 |
| AT1G61890.1 | --------------------------------------------LGMG------------ | 106 |
| Manes.16G008000.1 | --------------------------------------------LGMG------------ | 109 |
| Manes.17G038200.1 | --------------------------------------------LGMG------------ | 110 |
| Sobic.002G318300.1 | --------------------------------------------LGMG------------ | 99 |
| Manes.17G038300.1 | --------------------------------------------LGMG------------ | 96 |
| Manes.16G007900.1 | --------------------------------------------LGMG------------ | 105 |
| Manes.17G038400.1 | --------------------------------------------LGMG------------ | 105 |
| Sobic.001G012600.1 | --------------------------------------------LGMG------------ | 102 |
| Sobic.001G012600.2 | --------------------------------------------LGMG------------ | 102 |
| Sobic.001G185600.1 | --------------------------------------------LGMG------------ | 127 |
| Sobic.001G185400.1 | --------------------------------------------LGMG------------ | 101 |
| AT3G21690.1 | --------------------------------------------LGMG------------ | 111 |
| Sobic.001G185800.1 | --------------------------------------------LGMG------------ | 105 |
| Sobic.001G185500.1 | --------------------------------------------LGMG------------ | 119 |
| Sobic.001G185500.2 | ----------------------------------------------MG------------ | 2 |
| Sobic.007G165500.2 | --------------------------------------------LGMG------------ | 136 |
| AT4G00350.1 | --------------------------------------------LGMA------------ | 147 |
| Manes.01G255000.1 | --------------------------------------------FGMG------------ | 209 |
| Sobic.004G349550.1 | ----------------------------------------------MA------------ | 2 |
| Sobic.004G349600.1 | --------------------------------------------LGMG------------ | 106 |
| Sobic.007G176000.1 | --------------------------------------------LGMG------------ | 110 |
| Sobic.007G176100.1 | --------------------------------------------LGMG------------ | 115 |
| AT1G47530.1 | --------------------------------------------LGMG------------ | 94 |
| Manes.05G164500.1 | --------------------------------------------LGMG------------ | 90 |
| Manes.05G164500.2 | --------------------------------------------LGMG------------ | 90 |
| Manes.18G030900.1 | --------------------------------------------LGMG------------ | 91 |
| AT1G23300.1 | --------------------------------------------LGMG------------ | 103 |
| AT3G26590.1 | --------------------------------------------LGMG------------ | 104 |
| AT5G38030.1 | --------------------------------------------LGMG------------ | 104 |
| Manes.12G023400.1 | --------------------------------------------LGMG------------ | 107 |
| Manes.12G023500.1 | --------------------------------------------LGMG------------ | 107 |
| Manes.12G023600.1 | --------------------------------------------LGMG------------ | 124 |
| Manes.13G025200.1 | -------------------------------------------QLGMG------------ | 124 |
| Manes.13G025200.2 | --------------------------------------------LGMG------------ | 123 |
| Sobic.001G273100.1 | --------------------------------------------YGMG------------ | 100 |
| Sobic.001G273100.2 | --------------------------------------------YGMG------------ | 100 |

```
Sobic.001G273000.2      --------------------------------------------FGMG------------  73
Sobic.001G273000.1      --------------------------------------------FGMG------------  84
Sobic.001G273000.3      --------------------------------------------FGMG------------  84
Manes.15G147800.1       --------------------------------------------LGMG------------  89
Manes.15G147900.1       --------------------------------------------LGMG------------  90
Manes.17G098800.1       --------------------------------------------LGMG------------  95
Manes.17G098900.1       --------------------------------------------LGMG------------  95
Manes.17G098900.2       ----------------------------------------------MG------------  2
Sobic.001G162400.1      --------------------------------------------LGMG------------  80
Sobic.003G307600.2      ----------------------------------------------MG------------  2
Sobic.002G232200.1      --------------------------------------------LGMA------------  98
Sobic.002G232500.1      --------------------------------------------VGMA------------  94
Sobic.002G232600.1      --------------------------------------------LGMA------------  89
Sobic.007G160700.1      --------------------------------------------LGMA------------  106
Sobic.003G126200.2      --------------------------------------------LGMA------------  117
Sobic.001G476700.1      --------------------------------------------LGMA------------  114
Sobic.001G476700.2      -------------------------WICLVPQLLPR----RRIWLGMA------------  137
Manes.18G062800.1       --------------------------------------------LGMA------------  97
AT5G44050.1             --------------------------------------------IGMA------------  99
AT5G10420.1             --------------------------------------------LGMA------------  97
AT5G65380.1             --------------------------------------------LGMA------------  96
Manes.12G129000.1       --------------------------------------------LGMA------------  98
Manes.13G097900.1       --------------------------------------------LGMA------------  98
Sobic.005G020700.1      --------------------------------------------LGMA------------  130
Sobic.008G019400.1      --------------------------------------------LGMA------------  126
Manes.09G135300.1       --------------------------------------------LGMA------------  89
AT1G33080.1             --------------------------------------------LGMA------------  95
AT1G33090.1             --------------------------------------------LGMA------------  95
AT1G33100.1             --------------------------------------------LGMA------------  92
AT1G33110.1             --------------------------------------------LGMA------------  95
AT3G03620.1             --------------------------------------------GGMS------------  96
AT5G17700.1             --------------------------------------------AGMS------------  93
Manes.09G135200.1       --------------------------------------------IGMS------------  87
Manes.09G134800.1       --------------------------------------------LGMS------------  84
Manes.08G150400.1       --------------------------------------------LGMS------------  82
Manes.09G134900.1       --------------------------------------------LGMS------------  83
Manes.09G135000.1       --------------------------------------------LGMS------------  83
Manes.09G134900.2       --------------------------------------------TGMS------------  54
Manes.09G135000.2       --------------------------------------------TGMS------------  54


Sobic.002G099300.1      ------------------------------------------------------GIWIGM  6
AT4G39030.1             ------------------------------------------EAQHQISVLLFIGLVCGL  191
Manes.04G084700.1       ------------------------------------------EVQHQISILLLVGLICGI  211
Manes.11G091900.1       ------------------------------------------EVQHQLSVLVFIGLTCGF  209
Sobic.004G019800.2      ------------------------------------------LAQHQVSMLLFLALACGI  73
Sobic.004G019800.3      ------------------------------------------LAQHQVSMLLFLALACGI  73
Sobic.004G019800.4      ------------------------------------------LAQHQVSMLLFLALACGI  73
AT2G21340.1             ------------------------------------------EVQHQISILLFIGLACGV  208
Manes.11G092000.1       ------------------------------------------EVQHQISILLFVGLACGV  206
Sobic.001G476700.3      -----LAGEA----------------------------ACHRRRTRSHQCR*--------  164
Sobic.009G077000.1      ------------------------------------------------------------  80
Sobic.001G454900.1      SETNNLPASGPDLAECVNSCIPTE-----CT--DLPNQGCKKRYIPSVTSALIVGSILGL  229
Sobic.003G403000.1      PEKQQQPADEEAAKNGGEGCAPAV-----VAGRSSGKKSGNRRFVPSVTSALIVGALLGL  264
Sobic.003G403000.2      PEKQQQPADEEAAKNGGEGCAPAV-----VAGRSSGKKSGNRRFVPSVTSALIVGALLGL  264
Sobic.003G403000.3      PEKQQQPADEEAAKNGGEGCAPAV-----VAGRSSGKKSGNRRFVPSVTSALIVGALLGL  264
Sobic.007G020600.1      EMEELISPEGASA-----TTSISSFE----TDSCEVSVEQKRKNIPSVSTALLLGGVLGL  189
Sobic.007G020600.2      EMEELISPEGASA-----TTSISSFE----TDSCEVSVEQKRKNIPSVSTALLLGGVLGL  189
AT3G08040.1             NDTN-QPQQPPAPD--------------TKSNSGNKSNKKEKRTIRTASTAMILGLILGL  183
Manes.09G027700.1       ETEELLPKSGSIS---------------------------TKRHIPSASSALVIACVL--  156
Manes.09G027800.1       --------------------------------------------------------MV--  2
Manes.09G027900.1       --------------------------------------------------------MV--  2
Manes.09G026900.1       EMEELLPKSESTHKSSS-------------ASSISTKRDYERRHIPSASSALAIACVLGV  172
Manes.09G026900.2       EMEELLPKSESTHKSSS-------------ASSISTKRDYERRHIPSASSALAIACVLGV  172
Manes.09G027000.1       ------------------------------------------------------------  0
Manes.07G006000.1       E-KDPIPDDADCKAVA---CKSSTFTEGKAFKEKPKNK-KERRHIPSASTALIVGGILGL  207
Manes.07G006000.2       E-KDPIPDDADCKAVA---CKSSTFTEGKAFKEKPKNK-KERRHIPSASTALIVGGILGL  207
Manes.10G143000.1       KNKDSIPEDAT--------CKSPTFTEGKGVNEKSNNKKKGRRHIPSASTALIVGGILGL  200
AT1G51340.2             ETIELIPEKHKDSLSDEFKTSSSIFS--------ISKPPAKKRNIPSASSALIIGGVLGL  179
Manes.06G164500.1       ESKELIPQNG--------KSLITSFD--------IAKIENERRHIPSASSAVVIGAILGF  172
Manes.06G164500.2       ESKELIPQNG--------KSLITSFD--------IAKIENERRHIPSASSAVVIGAILGF  172
```

```
Manes.06G164500.3    ESKELIPQNG--------KSLITSFD--------IAKIENERRHIPSASSAVVIGAILGF  172
Manes.06G164500.4    ESKELIPQNG--------KSLITSFD--------IAKIENERRHIPSASSAVVIGAILGF  172
Manes.06G164500.5    ESKELIPQNG--------KSLITSFD--------IAKIENERRHIPSASSAVVIGAILGF  172
Manes.14G002600.1    ESKELIPQNDPVEGASKSKSLISIFE--------VSKIENERRHIPSASSALVIGAILGF  156
Sobic.008G006100.1   -------------------------------LTPIEKARQQKKVLPAVSTSLALAAGIGL  213
Sobic.005G005400.1   -------------------------------PRSPEELTQKRKFLPAVSTSLALAAGIGL  197
Sobic.005G005400.3   -------------------------------PRSPEELTQKRKFLPAVSTSLALAAGIGL  197
Sobic.005G005400.2   -------------------------------PRSPEELTQKRKFLPAVSTSLALAAGIGL  197
AT2G38330.1          ----------------------------------DS-IETSKKVLPSVSTSLVLAAGVGI  179
Manes.08G096600.1    ----------------------------------CVNEPQGKTYLPAVSTSLALAAAVGI  205
Manes.08G096600.2    ----------------------------------CVNEPQGKTYLPAVSTSLALAAAVGI  205
AT4G38380.1          ------------------------------QSDIPSQGLPERKQLSSVSTALVLAIGIGI  218
Sobic.002G286800.1   --------------------------------GNISDKIGERKRLPSISSALLLAAAIGV  235
Sobic.003G149300.2   --------------------------------GK--------LELTSVSSALILAAGIGI  205
Sobic.003G149300.1   --------------------------------GK--------LELTSVSSALILAAGIGI  205
Sobic.003G149300.3   --------------------------------GK--------LELTSVSSALILAAGIGI  205
Sobic.003G149300.4   --------------------------------GK--------LELTSVSSALILAAGIGI  81
Manes.01G153400.1    ------------------------------------------------------------  0
Manes.04G064900.1    -----------------------------ITNGKPTADVAERKQLSSVSTALLLAVGIGI  246
Manes.04G064900.2    -----------------------------ITNGKPTADVAERKQLSSVSTALLLAVGIGI  246
Sobic.001G003700.1   GAMEPICGQA------------------------HGAGNVA-LLRGTLLRATLMLLAASV  114
AT4G22790.1          AAMEPICGQA------------------------FGAKNFK-LLHKTLFMAVLLLLLISV  124
Manes.02G032300.1    AAMEPICGQA------------------------YGAKNFR-LLHKTLLMTIFLLLLTTL  136
Manes.10G000400.1    MGMEPICCQA------------------------YGAKKWS-ILSQTYKKTLCILFLATI  100
AT2G38510.1          VGMDPICGQA------------------------FGAKRWT-VLSHTFQKMFCLLIVVSV  100
Manes.08G172300.1    MGMDPICGQA------------------------YGAKRWS-VISQTYLRTLCLLLVVAL  130
Manes.09G117000.1    MGMDPICGQA------------------------YGAKRWS-VISQTYLRTLCLLLLVAL  100
Sobic.010G167800.1   LGMDPLCTQA------------------------FGANQPR-LLGLTLYRSVLFLLCCSL  154
AT5G19700.1          LGMDPLCSQA------------------------FGAGRPK-LLSLTLQRTVLFLLTSSV  130
AT4G29140.1          LGMEPLCSQA------------------------FGAHRFK-LLSLTLHRTVVFLLVCCV  150
Manes.03G026500.1    LGMEPLCSQA------------------------FGAQRTK-LLSVTLHRSVIFLLVSSL  157
Manes.16G109300.1    LGMEPLCSQA------------------------FGAQRPK-LLSVTLHRSVIFLLVSSI  151
Sobic.007G181100.1   AGMDPVCGQA------------------------FGAGRTS-VLTAALRRTVVLLLAASV  139
Manes.13G127800.1    MGTEAISSQA------------------------CGAKQWP-LMGQTLQRTIAILILTCV  149
Sobic.001G446800.1   LGMEPICGQA------------------------FGARRGK-LLALALHRTVLLLLAVAL  146
AT5G52050.1          MGVESICSQA------------------------FGARRYN-YVCASVKRGIILLLVTSL  129
AT1G58340.1          MGMEPICGQA------------------------YGAKQMK-LLGLTLQRTVLLLLSCSV  152
Manes.05G186700.1    MGMEPICGQA------------------------YGAKQWK-LLGLTLQRTVLLLLSTSI  143
Manes.18G054300.1    MGMEPICGQA------------------------YGAKQWK-LLGLTLQRTVLLLLSTSV  143
Sobic.001G320900.1   MGMEPVCGQA------------------------VGAKNLP-LVGATMQRMVLLLLALSV  171
Sobic.001G320900.2   MGMEPVCGQA------------------------VGAKNLP-LVGATMQRMVLLLLALSV  171
Sobic.004G283500.1   MGMEPICGQA------------------------FGAGHYE-LLGVTTQRAVLMLLAAAV  172
Sobic.006G184400.1   MGMEPICGQA------------------------FGAGNFS-LLGITMQRTVLLLIAAAV  185
AT4G23030.1          IGMEPICVQA------------------------FGAKRFK-LLGLALQRTTLLLLLCSL  127
Manes.01G067000.1    MGMEPICGQA------------------------FGAKRYK-LLGLTMQRTILLLILISL  147
Manes.02G027800.1    MGMEPICGQA------------------------FGAKRYK-LLGLTMQRTILLLILISF  151
Manes.06G143600.1    MGMEPICGQA------------------------FGAQKHS-LLGLTLQRTILLLIFTSL  159
Manes.14G029100.1    MGMEPICGQS------------------------FGAQKHT-LLGLTLQRTILLLLMTSL  163
AT5G49130.1          TGMEPLCGQA------------------------IGSKNPS-LASLTLKRTIFLLLLASL  120
Manes.03G198000.1    MGMEPLCSQA------------------------FGSRNLS-VASMTLQRTILMLLVVSL  113
Manes.15G011000.1    MGMEPLCSQA------------------------FGSRNLS-VASLTLQRTIVMLLLASL  113
Sobic.001G019700.1   SGLEPLCAQA------------------------FGSRNHE-LLTLSVQRAVLLLFLAAV  137
AT1G71870.1          SGLEPVCSQA------------------------YGSKNWD-LLTLSLHRMVVILLMASL  118
Manes.02G189800.1    SGLEPVCSQA------------------------YGSKNWD-LLSLSLQRMILILFIAII  116
Manes.18G098800.1    SGLEPVCSQA------------------------YGSQNWD-LLSLSLQRMILILLIAII  116
Sobic.010G256932.1   ---------------------------------------------------YNCFMLTGI  93
Sobic.010G256700.1   SALDTFCGQS------------------------YGARQYD-MLGTHMQRAIIVLMLTGV  124
Sobic.010G256700.5   SALDTFCGQS------------------------YGARQYD-MLGTHMQRAIIVLMLTGV  124
Sobic.010G256700.6   SALDTFCGQS------------------------YGARQYD-MLGTHMQRAIIVLMLTGV  124
Sobic.010G256700.4   SALDTFCGQS------------------------YGARQYD-MLGTHMQRAIIVLMLTGV  124
Sobic.009G106900.1   ------------------------------------------------------------  0
Manes.S031500.1      SALETFCGQA------------------------YGAKQYH-MLGIHLQRAVIVLLLVSV  128
Manes.14G109900.1    SALETLCGQA------------------------YGAKQYH-MLGIHTQRAMLTLLALSI  138
Manes.14G109900.2    SALETLCGQA------------------------YGAKQYH-MLGIHTQRAMLTLLALSI  37
Manes.14G109900.3    SALETLCGQA------------------------YGAKQYH-MLGIHTQRAMLTLLALSI  37
AT1G73700.1          SALETLCGQA------------------------YGAKLYG-KLGIQMQRAMFVLLILSV  118
AT2G34360.1          SAMDTVCGQS------------------------YGAKMYG-MLGIQMQRAMLVLTLLSV  121
AT5G52450.1          SALDTLCGQA------------------------YGAKKYG-MLGIQMQRAMFVLTLASI  120
Sobic.009G106960.1   ------------------------------------------------------------  0
Manes.14G060800.1    SALETFCGQA------------------------YGARQFH-MLGVHLQRAMLVLVLTSI  131
Sobic.009G106800.1   SALDTLCGQA------------------------FGARQYG-LLGVYKQRAMLVLALACL  117
```

```
Sobic.007G074300.2      ------------------------------------------------------------  0
Sobic.009G106700.1      TALDTLCGQA------------------------FGARQHR-LLGVYKQRAMVVLGLACV  119
Sobic.010G138400.4      SALDTLCGQA------------------------FGARQYY-LLGIYKQRAMFLLTLVSL  44
Sobic.010G138400.1      SALDTLCGQA------------------------FGARQYY-LLGIYKQRAMFLLTLVSL  122
Sobic.010G138400.5      SALDTLCGQA------------------------FGARQYY-LLGIYKQRAMFLLTLVSL  41
Sobic.004G129900.1      SSLDTLCGQA------------------------FGAGQHH-LLGIYKQRAMLVLAAVSV  119
Sobic.004G129900.2      SSLDTLCGQA------------------------FGAGQHH-LLGIYKQRAMLVLAAVSV  37
Sobic.006G042200.1      SSLDTLCGQS------------------------FGAKQYY-LLGIYKQRAILVLTLVSL  122
Sobic.004G129700.1      CSLDTLCGQA------------------------FGAKQYY-QLSVYKQRAMVVLTLVSI  131
Sobic.004G129800.1      CSLDTLCGQA------------------------FGAGQHH-QLGVYKQRAMVVLALVSV  122
Sobic.004G129800.2      CSLDTLCGQA------------------------FGAGQHH-QLGVYKQRAMVVLALVSV  37
AT3G23550.1             GALETLCGQG------------------------FGAKSYR-MLGIHLQSSCIVSLVFTI  125
AT3G23560.1             GSLETLCGQG------------------------FGAKRYR-MLGVHLQSSCIVSLVFSI  133
Manes.15G088100.1       GALETLCGQG------------------------FGAKLYR-MLGIHLQASCIISFFLSI  145
Manes.03G109100.1       EALETFCGQG------------------------FGAKAYN-MLGLYLQAFCIISFFFSI  129
Manes.03G109100.2       EALETFCGQG------------------------FGAKAYN-MLGLYLQAFCIISFFFSI  90
Sobic.002G311200.1      GALETLCGQA------------------------YGAGLYR-MLGLYLQSSLIMSAAVSA  240
Sobic.002G006500.3      GSLETLCGQG------------------------YGARAYR-TMGVHLQASLLTSALASA  154
Sobic.002G006500.4      GSLETLCGQG------------------------YGARAYR-TMGVHLQASLLTSALASA  154
AT2G04090.1             GALETLCGQA------------------------YGAKQYT-KIGTYTFSAIVSNVPIVV  124
AT2G04100.1             GALETLCGQA------------------------YGAKQYA-KIGTYTFSAIVSNVPIVV  124
AT2G04066.1             ------------------------------------------------------------  0
AT2G04040.1             GALETLCGQA------------------------YGAKQYE-KIGTYAYSAIASNIPICF  121
AT2G04080.1             GALETLCGQA------------------------YGAEQYE-KIGTYTYSAMASNIPICF  121
AT2G04050.1             GALETLCGQA------------------------YGAKQYE-KIGTYTYSATASNIPICV  121
AT2G04070.1             GSLETLSGQA------------------------YGAKQYE-KMGTYTYSAISSNIPICV  121
Sobic.001G481800.1      SGLETICGQA------------------------FGAEQYH-KVALYTYRSIIVLLIASV  148
Sobic.003G260200.1      SALETLCGQA------------------------YGAKQYH-KLGLDTYRAIVTLLVVCI  131
Sobic.009G224800.1      SGLETLCGQA------------------------YGAKQYD-KLGMHTYRAIVTLIVVSI  134
Sobic.009G224800.2      SGLETLCGQA------------------------YGAKQYD-KLGMHTYRAIVTLIVVSI  134
AT1G66760.2             GSLETLCGQA------------------------YGAKQYH-KLGSYTFTSIVFLLIISV  122
AT1G64820.1             GALDTLCGQA------------------------FGAEQFG-KIGAYTYSSMLCLLVFCF  123
AT1G66780.1             GALETLCGQA------------------------FGAGQFR-NISAYTYGSMLCLLLVCF  129
Manes.02G072400.1       GALETLCGQA------------------------YGAEEHH-KLGTYTYSAIISLVMICP  119
Manes.01G113500.1       GALETLCGQA------------------------YGAGQYH-KLGSYTYCAIISLLPVCV  107
Manes.01G113500.2       GALETLCGQA------------------------YGAGQYH-KLGSYTYCAIISLLPVCV  107
AT1G15150.1             CALDTLSGQA------------------------YGAKLYR-KLGVQAYTAMFCLTLVCL  125
AT1G15160.1             CALDTLSGQA------------------------YGAKLYR-KLGVQAYTAMFCLTLVCL  125
AT1G15170.1             CALDTLSGQA------------------------YGAKLYR-KLGVQTYTAMFCLALVCL  128
AT1G15180.1             CALDTLSGQA------------------------YGAKLYR-KVGVQTYTAMFCLALVCL  129
AT1G71140.1             SALETLCGQA------------------------NGAKQYE-KLGVHTYTGIVSLFLVCI  120
Manes.06G063000.1       SALETLCGQA------------------------HGAKQYR-QFGVQIYTAIFSLNIVCI  120
Manes.14G109700.1       SGLETLCGQA------------------------YGAEQYR-KLGSQTYSAIFSLILVAF  129
Manes.14G109800.1       SALETLCGQA------------------------YGAKQYK-KLGIQTQTAIFCLILVCI  130
Sobic.008G171600.1      SAVQTVCGQA------------------------YGARRYA-AMGIVCQRALVLQLATAI  137
Sobic.008G171600.2      SAVQTVCGQA------------------------YGARRYA-AMGIVCQRALVLQLATAI  151
Sobic.008G171600.3      SAVQTVCGQA------------------------YGARRYA-AMGIVCQRALVLQLATAI  137
AT3G59030.1             SAVQTVCGQA------------------------YGARQYS-SMGIICQRAMVLHLAAAV  145
Manes.01G182000.1       SAVQTACGQA------------------------YGAKRYS-AMGVICQRAIVLHLGAAV  144
Manes.02G142000.1       SAVQTVCGQA------------------------YGAKQYS-AMGIICQRAIVLHLGAAV  144
AT4G21903.2             SAVETLCGQA------------------------YGAHRYE-MLGIYLQRATIVLALVGF  146
AT4G21910.4             SAVETLCGQA------------------------YGAHRYE-MLGIYLQRATIVLALVGL  150
AT1G11670.1             SAVETLCGQA------------------------HGAHRYD-MLGVYLQRSTIVLVITGL  144
AT1G61890.1             SAVETLCGQA------------------------HGAHRYE-MLGVYLQRSTVVLILTCL  141
Manes.16G008000.1       SAVETLCGQA------------------------YGAHRNE-MLGIYLQRAIVVLTLTAI  144
Manes.17G038200.1       SAVETLCGQA------------------------YGAQKYA-MLGTYLQRATVVLTLTGI  145
Sobic.002G318300.1      SAVETLCGQA------------------------YGAEKYE-MLGVYLQRSTVLLMATGV  134
Manes.17G038300.1       SAVETLCGQA------------------------YGANRFE-MLGIYLQRSIILLVATGI  131
Manes.16G007900.1       SAVETLCGQA------------------------FGAHKYE-MLGVYLQRSTILLMAAGI  140
Manes.17G038400.1       SAVETLCGQA------------------------YGAHKFE-MLGIYLQRSTILLMATGI  140
Sobic.001G012600.1      SAVETLCGQA------------------------YGAHKPG-MLGVYLQRSTVLLTATGV  137
Sobic.001G012600.2      SAVETLCGQA------------------------YGAHKPG-MLGVYLQRSTVLLTATGV  137
Sobic.001G185600.1      SAVETLCGQA------------------------YGAHKYD-MLGIYMQRSIVLLTATGV  162
Sobic.001G185400.1      SAVETLCGQA------------------------YGAQKYD-MLGIYLQRSAVLLCATGV  136
AT3G21690.1             SAVETLCGQA------------------------YGGRKYE-MLGVYLQRSTVLLTLTGL  146
Sobic.001G185800.1      SAVETLCGQA------------------------YGAHRYE-MLGIYLQRSTILLVAVGV  140
Sobic.001G185500.1      SAVETLCGQA------------------------YGAHKYD-MLGIYLQRSTILLMATGV  154
Sobic.001G185500.2      SAVETLCGQA------------------------YGAHKYD-MLGIYLQRSTILLMATGV  37
Sobic.007G165500.2      SALETLCGQA------------------------FGAGQVA-MLGVYLQRSWLILVAACV  171
AT4G00350.1             SALETLCGQA------------------------FGAGQMD-MLGVYMQRSWLILLGTSV  182
Manes.01G255000.1       SALETLCGQA------------------------FGAGQTE-LLGVYMQRSWIILFSACF  244
```

```
Sobic.004G349550.1      SALETLCGQA------------------------YGAGHLH-LLGVYAQRSVVILAASAL  37
Sobic.004G349600.1      SALETLCGQA------------------------YGAGQVA-MLGVYMQRSWIVLAASAA  141
Sobic.007G176000.1      SALETLCGQA------------------------YGAKQLH-MLGVYLQRSWIILTAMAV  145
Sobic.007G176100.1      SALETLCGQA------------------------YGAKQLH-MLGVYLQRSWIILTAMAV  150
AT1G47530.1             SALETLCGQA------------------------YGAGQIR-MMGIYMQRSWVILFTTAL  129
Manes.05G164500.1       SALETLCGQA------------------------YGAGQLR-MLGIYMQRSWVILLTTAC  125
Manes.05G164500.2       SALETLCGQA------------------------YGAGQLR-MLGIYMQRSWVILLTTAC  125
Manes.18G030900.1       SALETLCGQA------------------------FGAGQLR-MLGIYMQRSWVILLTTAC  126
AT1G23300.1             SALATLCGQA------------------------YGAGQLE-MMGIYLQRSWIILNSCAL  138
AT3G26590.1             SALETLCGQA------------------------FGAGKLS-MLGVYLQRSWVILNVTAL  139
AT5G38030.1             SALETLCGQA------------------------FGAGKLS-MLGVYLQRSWVILNVTAV  139
Manes.12G023400.1       SALETLCGQA------------------------FGAGQLE-MLGIYLQRSWVILGTTAS  142
Manes.12G023500.1       SALETLCGQA------------------------FGAGQLE-MLGIYLQRSWVILGTTAS  142
Manes.12G023600.1       SALETLCGQA------------------------FGAGQLD-MLGIYLQRSWIILCTTAS  159
Manes.13G025200.1       SALETLCGQA------------------------FGAGKID-MLGIYLQRSWIILCTTAS  159
Manes.13G025200.2       SALETLCGQA------------------------FGAGKID-MLGIYLQRSWIILCTTAS  158
Sobic.001G273100.1      SALETLSGQA------------------------VGAGQPE-KLGVYTQQSWIISVATAV  135
Sobic.001G273100.2      SALETLSGQA------------------------VGAGQPE-KLGVYTQQSWIISVATAV  135
Sobic.001G273000.2      SALNTLIGQA------------------------VGAGQLD-RLGTYTQQSLIICGTTAL  108
Sobic.001G273000.1      SALNTLIGQA------------------------VGAGQLD-RLGTYTQQSLIICGTTAL  119
Sobic.001G273000.3      SALNTLIGQA------------------------VGAGQLD-RLGTYTQQSLIICGTTAL  119
Manes.15G147800.1       SALETLCGQA------------------------VGAGQLN-MLGIYMQRSWIITGVTAL  124
Manes.15G147900.1       SALETLCGQA------------------------VGAGQMN-LLGIYMQRSWIITGVTAL  125
Manes.17G098800.1       SALEALCGQA------------------------VGAGQLN-MLGIYMQRSWIITGITAL  130
Manes.17G098900.1       SALETLCGQA------------------------VGAGQLN-MLGIYMQRSWIITGITAL  130
Manes.17G098900.2       SALETLCGQA------------------------VGAGQLN-MLGIYMQRSWIITGITAL  37
Sobic.001G162400.1      SALETLCGQA------------------------VGAGQVS-MLGVYIQRSWLICGATAV  115
Sobic.003G307600.2      SALETLCGQA------------------------VGAGQLQ-MLGVYMQRSWIICVATSL  37
Sobic.002G232200.1      SATETLCGQA------------------------YGAKQYH-MLGIYLQRSWLVLLAFAV  133
Sobic.002G232500.1      SAMETLCGQA------------------------YGAKQYH-MMGIYLQRSWLILLSFAV  129
Sobic.002G232600.1      SALETLCGQA------------------------YGAKQYS-MMGTYLQRSWLVLLAFAV  124
Sobic.007G160700.1      SALETLCGQA------------------------FGAKKHH-MLGVYLQRSWIVLLIFAA  141
Sobic.003G126200.2      SALETLCGQA------------------------YGAKKYH-MMGVYMQRSWIVLFVCAL  152
Sobic.001G476700.1      SALETLCGQA------------------------FGAKKFH-MMGVYMQRSWIVLFLCAV  149
Sobic.001G476700.2      SALETLCGQA------------------------FGAKKFH-MMGVYMQRSWIVLFLCAV  172
Manes.18G062800.1       SALETLCGQA------------------------FGAKRYQ-MLGIYMQRSWIVLFFCCF  132
AT5G44050.1             TALETLCGQA------------------------FGAKKYD-MFGVYLQRSWIVLFLFSI  134
AT5G10420.1             SALETLCGQA------------------------FGAREYY-MLGVYMQRYWIILFLCCI  132
AT5G65380.1             SALETLCGQA------------------------FGAKKYH-MLGVYMQRSWIVLFFCCV  131
Manes.12G129000.1       SALETLCGQA------------------------FGAKKYY-MLGVYMQRSWIVLFVCCI  133
Manes.13G097900.1       SALETLCGQA------------------------FGAKKYY-MLGVYMQRSWIVLFLCCV  133
Sobic.005G020700.1      SALETLCGQS------------------------YGAKQYH-MLGIYLQRSWIILFACSV  165
Sobic.008G019400.1      SALETLCGQS------------------------YGAKQYH-MLGIYLQRSWIILFACSV  161
Manes.09G135300.1       SALETLCGQS------------------------YGAKQYH-MLGIYLQRSWIVLTMCAI  124
AT1G33080.1             SALETLCGQA------------------------YGAKQYH-MLGIYLQRSWIVLTGCTI  130
AT1G33090.1             SALGTLCGQA------------------------YGAKQYH-MLGIHLQRSWIVLTGCTI  130
AT1G33100.1             GALGTLCGQA------------------------YGAKQYQ-MLGIYLQRSWIVLTGGTI  127
AT1G33110.1             SALETLCGQA------------------------YGAKQNH-MLGIYLQRSWIVLTGCTI  130
AT3G03620.1             SATETLCGQA------------------------YGAEQYH-TMGIYLQRSWIVDMAVTT  131
AT5G17700.1             SATETLCGQA------------------------YGAEQYH-MMGIYLQRSWIVDTFIAT  128
Manes.09G135200.1       SATETLCGQA------------------------FGAAQYH-MLGIYLQRSWIVDHILTT  122
Manes.09G134800.1       SAMETLCGQA------------------------FGAKQYH-MMGIYLQRSWIINFSTAT  119
Manes.08G150400.1       SATDTLCGQA------------------------FGARQYH-MMGVYLQRSWIINFLTAT  117
Manes.09G134900.1       SATETLCGQA------------------------FGARQYH-MMGIYLQRSCIINVVTAT  118
Manes.09G135000.1       SATETLCGQA------------------------FGARQYH-MMGIYLQRSCIINLITAT  118
Manes.09G134900.2       SATETLCGQA------------------------FGARQYH-MMGIYLQRSCIINVVTAT  89
Manes.09G135000.2       SATETLCGQA------------------------FGARQYH-MMGIYLQRSCIINLITAT  89


Sobic.002G099300.1      LTGTFLQMSILLA-I-IFTTKWD------------------KQAAL-----AEVR-----  36
AT4G39030.1             MMLLLTRLFGPWAVT-AFTRGKN------------------IEIVPAANKYIQIRGLAWP  232
Manes.04G084700.1       LMLLFTQFLGSWALT-AFAGPKN------------------LHIVPAASKYVQIRGLAWP  252
Manes.11G091900.1       LMILFTKFFAASVLA-AFAGSNN------------------LHIVPAANTYVQIRGLAWP  250
Sobic.004G019800.2      GMFLFTKVFGTQVLT-AFTGSGN------------------YELISSANTYAQIRGFAWP  114
Sobic.004G019800.3      GMFLFTKVFGTQVLT-AFTGSGN------------------YELISSANTYAQIRGFAWP  114
Sobic.004G019800.4      GMFLFTKVFGTQVLT-AFTGSGN------------------YELISSANTYAQIRGFAWP  114
AT2G21340.1             TMMVLTRLFGSWALT-AFTGVKN------------------ADIVPAANKYVQIRGLAWP  249
Manes.11G092000.1       LMFLFTRFFGSWALT-AFTGPKN------------------VHIVPAANTYVQIRGYAWP  247
Sobic.001G476700.3      ------------------------------------------------------------  164
Sobic.009G077000.1      ------------------------------------------------------------  80
Sobic.001G454900.1      LQAVFLVFSAKFVLN-IMGVKSG------------------SPMQKPAVRYLTIRSLGAP  270
```

```
Sobic.003G403000.1      FQTVFLVAAGKPLLR-LMGVKPG------------------SPMVMPALRYLTLRALGAP  305
Sobic.003G403000.2      FQTVFLVAAGKPLLR-LMGVKPG------------------SPMVMPALRYLTLRALGAP  305
Sobic.003G403000.3      FQTVFLVAAGKPLLR-LMGVKPG------------------SPMVMPALRYLTLRALGAP  305
Sobic.007G020600.1      LETVLLVFSAKPILG-YMGVTPD------------------SAMMKPALQYLVLRSLGAP  230
Sobic.007G020600.2      LETVLLVFSAKPILG-YMGVTPD------------------SAMMKPALQYLVLRSLGAP  230
AT3G08040.1             VQAIFLIFSSKLLLG-VMGVKPN------------------SPMLSPAHKYLSIRALGAP  224
Manes.09G027700.1       ------------------------------------------------------------  156
Manes.09G027800.1       ------------------------------------------------------------  2
Manes.09G027900.1       ------------------------------------------------------------  2
Manes.09G026900.1       IQALFLILAAKPVLS-YMGVQSD------------------SPMLIPAQQYLTLRSLGAP  213
Manes.09G026900.2       IQALFLILAAKPVLS-YMGVQSD------------------SPMLIPAQQYLTLRSLGAP  213
Manes.09G027000.1       ------------------------------------------------------------  0
Manes.07G006000.1       VQAIFLIFCAKPLLS-IMGVKSG------------------SPMLTPARKYLTLRALGSP  248
Manes.07G006000.2       VQAIFLIFCAKPLLS-IMGVKSG------------------SPMLTPARKYLTLRALGSP  248
Manes.10G143000.1       VQAIFLIFCAKPLLN-IMGVKSN------------------SPMLTPARKYLTLRSLGSP  241
AT1G51340.2             FQAVFLISAAKPLLS-FMGVKHD------------------SPMMRPSQRYLSLRSLGAP  220
Manes.06G164500.1       IQAIFLISGAKPLLN-FMGVGSD------------------SPMLIPAQQYLTLRSLGAP  213
Manes.06G164500.2       IQAIFLISGAKPLLN-FMGVGSD------------------SPMLIPAQQYLTLRSLGAP  213
Manes.06G164500.3       IQAIFLISGAKPLLN-FMGVGSD------------------SPMLIPAQQYLTLRSLGAP  213
Manes.06G164500.4       IQAIFLISGAKPLLN-FMGVGSD------------------SPMLIPAQQYLTLRSLGAP  213
Manes.06G164500.5       IQAIFLISGAKPLLN-FMGVGSD------------------SPMLIPAQQYLTLRSLGAP  213
Manes.14G002600.1       VQAIFLISAAKPLLN-FMGVSSD------------------SPMLIPAQQYLTLRSLGAP  197
Sobic.008G006100.1      LEMVALIVGSGTLIN-IIGIPVD------------------SPMRAPAEQFLTLRALGAP  254
Sobic.005G005400.1      METVALNFGSGTLMD-MIGIPID------------------SPMRIPAEQFLTFRAYGAP  238
Sobic.005G005400.3      METVALNFGSGTLMD-MIGIPID------------------SPMRIPAEQFLTFRAYGAP  238
Sobic.005G005400.2      METVALNFGSGTLMD-MIGIPID------------------SPMRIPAEQFLTFRAYGAP  238
AT2G38330.1             AEAIALSLGSDFLMD-VMAIPFD------------------SPMRIPAEQFLRLRAYGAP  220
Manes.08G096600.1       AEAVALFLGSGFLMN-IMGITAD------------------SPMRVPAENFLSWRAFGAP  246
Manes.08G096600.2       AEAVALFLGSGFLMN-IMGITAD------------------SPMRVPAENFLSWRAFGAP  246
AT4G38380.1             FEALALSLASGPFLR-LMGIQSM------------------SEMFIPARQFLVLRALGAP  259
Sobic.002G286800.1      IEALALILGSGILLN-IMGVSHA------------------SAMHNPARLFLSVRALGAP  276
Sobic.003G149300.2      MEALALFLGSGLFLK-LMGVSPV------------------SPMHRPAKLFLSLRALGAP  246
Sobic.003G149300.1      MEALALFLGSGLFLK-LMGVSPV------------------SPMHRPAKLFLSLRALGAP  246
Sobic.003G149300.3      MEALALFLGSGLFLK-LMGVSPV------------------SPMHRPAKLFLSLRALGAP  246
Sobic.003G149300.4      MEALALFLGSGLFLK-LMGVSPV------------------SPMHRPAKLFLSLRALGAP  122
Manes.01G153400.1       -----MKLGSFPF--------------------------------------L--------  9
Manes.04G064900.1       FEAVALSLGCGPFLN-LMGIKLD------------------SPMRVPAERFLLLRAVGAP  287
Manes.04G064900.2       FEAVALSLGCGPFLN-LMGIKLD------------------SPMRVPAERFLLLRAVGAP  287
Sobic.001G003700.1      PIALLWTRVDA-VLL-RFGQQPDIADTARTYVLCLLPDLAVTSVLNPLKAYLSAQEVTLP  172
AT4G22790.1             PISFLWLNVHK-ILT-GFGQREDISFIAKKYLLYLLPELPILSFLCPLKAYLSSQGVTLP  182
Manes.02G032300.1       PVSFLWLNVDK-ILI-HFGQQEDISHIAKTYLFYLLPDLVITSLLCPLKAYLSSQGVTVP  194
Manes.10G000400.1       PISLLWLYMEP-ILL-GFGQDQNITSIARVYITYSIPELLSQAHLHPLRIFIRIQNLTKP  158
AT2G38510.1             PIAVTWLNIEP-IFL-RLGQDPDITKVAKTYMLFFVPELLAQAMLHPLRTFLRTQGLTSP  158
Manes.08G172300.1       PISLLWLNVEP-IFL-RLGQDPDITSVAKVYMVFCIPELIAQAVLHPMRSFLRTQGLTAP  188
Manes.09G117000.1       PISLLWLNVEP-ILL-RLGQDPDITNVAKVYMVFCIPELIAQAVLHPIRSFLRIQGLTAP  158
Sobic.010G167800.1      PLSALWLNMSK-ILV-FLGQDMEITALAQDYILFSLPDLFSFSVIHPLRVYLRSQGITWP  212
AT5G19700.1             VIVALWLNLGK-IMI-YLHQDPSISSLAQTYILCSIPDLLTNSFLHPLRIYLRAQGITSP  188
AT4G29140.1             PISVLWFNVGK-ISV-YLHQDPDIAKLAQTYLIFSLPDLLTNTLLHPIRIYLRAQGIIHP  208
Manes.03G026500.1       PISLLWLNMSK-ILL-YLHQDPKITRLSHTYLLFSLPDLLTNAFIHPIRIYLRAQGITHP  215
Manes.16G109300.1       PISLLWLNMSK-ILL-YLHQDPNITRLAHTYLLFSLPDLLTNSFVHPIRIYLRAQGITHP  209
Sobic.007G181100.1      PITLLWLAMHR-ALV-ATGQDPDIAAAAYDFILCSLPDLVVQSFLHPLRVYLRAQSVTLP  197
Manes.13G127800.1       PISLLWLNVEP-ILI-FCGQDPAILSVASTYLAFSLPDLFLQSFINPLKIYLRTQNITLP  207
Sobic.001G446800.1      PISALWVTSTGYVLK-LLGQDEGVADAAQTFAAYASADLAVLAVLHPLRVYLRSQNLTLP  205
AT5G52050.1             PVTLLWMNMEK-ILL-ILKQDKKLASEAHIFLLYSVPDLVAQSFLHPLRVYLRTQSKTLP  187
AT1G58340.1             PISFSWLNMRR-ILL-WCGQDEEISSVAQQFLLFAIPDLFLLSLLHPLRIYLRTQNITLP  210
Manes.05G186700.1       PISFMWLNMKR-ILL-WCGQDQEISSMAQTFILFSIPDLFFLSLLHPLRIYLRTQGITLP  201
Manes.18G054300.1       PISFMWLNMKR-ILL-WCGQDQEISSVAHTFILFSIPDLFFLSLLHPLRIYLRTQSITLP  201
Sobic.001G320900.1      PVAFLWAHMEP-LLL-LCGQDAAISAAAQRYILLCLPDLLFQSFLHPLRIYLRTQSINFP  229
Sobic.001G320900.2      PVAFLWAHMEP-LLL-LCGQDAAISAAAQRYILLCLPDLLFQSFLHPLRIYLRTQSINFP  229
Sobic.004G283500.1      PIGGLWVHIRP-LLL-LCGQDAGIAAVAETYILASLPDLLLQAFLHPVRIYLRTQSINLP  230
Sobic.006G184400.1      PIGGLWMHMRP-LLL-LCGQDTGIAAVAETYILASLPDLVLQAFIHPVRIYLRAQSINLP  243
AT4G23030.1             PISILWLNIKK-ILL-FFGQDEEISNQAEIFILFSLPDLILQSFLHPIRIYLRSQSITLP  185
Manes.01G067000.1       PISFLWFNMKN-ILL-FCGQEDGIATEAQTYILFSLPDLILQSLLHPLRIYLRTQSITLP  205
Manes.02G027800.1       PIAFSWFNMKK-ILL-FCGQEEDIAAEAQTYILYSLPDLILQSFLHPLRIYLRTQSITLP  209
Manes.06G143600.1       PISLLWLNMKR-ILL-FCGQDMAIATEAQSFLLYSLPDLLAQSFLHPLRIYLRTQSITLP  217
Manes.14G029100.1       PISFLWLNMKR-ILL-FCGQDLAIATEAQSFLLYSLPDLLAQSLLHPLRIYLRTQSITLP  221
AT5G49130.1             PISLLWLNLAP-LML-MLRQQHDITRVASLYCSFSLPDLLANSFLHPLRIYLRCKGTTWP  178
Manes.03G198000.1       PIGLLWLNLEP-LML-ALHQDPEITRLASLYCRFCIPDLVANSLLHPLRIYLRSKGTTWP  171
Manes.15G011000.1       PIGLLWVNLEP-LML-ALHQDPDITRIASLYCRFSIPDLIANGLLHPLRIYLRSKGTTWP  171
Sobic.001G019700.1      PIALLWLNVGP-ILV-ALGQDPTISAHAASYAAFALPDLAAGAVLQPLRVYLRSQGITRP  195
AT1G71870.1             PISLLWINLGP-IML-FMGQNPEITATAAEYCLYALPDLLTNTLLQPLRVYLRSQRVTKP  176
```

| | | |
|---|---|---|
| Manes.02G189800.1 | PISLLWLNLES-IMN-FMGQDRDITAMAATYCIYSLPDLLTNTLLQPLRVFLRSQKVTKP | 174 |
| Manes.18G098800.1 | PISLLWLNLET-IMN-SMGQDRNITAMASTYCIYSLPDLLTNTLLQPLRVFLRSQKVTKP | 174 |
| Sobic.010G256932.1 | HLAFVLGFAGE-ILI-ALSQNPEISFQARLYAQWLIPGLFAYGLLQCLTRFLLTQNIVQI | 151 |
| Sobic.010G256700.1 | PLAFVLAFAGQ-ILI-ALGQNPEISSEAGLYAQWLIPGLFAYGLLQCLTRFLQTQNIVQI | 182 |
| Sobic.010G256700.5 | PLAFVLAFAGQ-ILI-ALGQNPEISSEAGLYAQWLIPGLFAYGLLQCLTRFLQTQNIVQI | 182 |
| Sobic.010G256700.6 | PLAFVLAFAGQ-ILI-ALGQNPEISSEAGLYAQWLIPGLFAYGLLQCLTRFLQTQNIVQI | 182 |
| Sobic.010G256700.4 | PLAFVLAFAGQ-ILI-ALGQNPEISSEAGLYAQWLIPGLFAYGLLQCLTRFLQTQNIVQI | 182 |
| Sobic.009G106900.1 | ------------------------------------------------------------ | 0 |
| Manes.S031500.1 | LLAFVWANAGE-ILL-FFRQDPEIAHEAGQYARYMIPSIFGFAIQDCLIRFLQTQNNVIP | 186 |
| Manes.14G109900.1 | PLAIIWFYTST-ILI-FVGQDHEISTGAGIFNRWMIPSLFAFALLQCLNRFLQTQNNVFP | 196 |
| Manes.14G109900.2 | PLAIIWFYTST-ILI-FVGQDHEISTGAGIFNRWMIPSLFAFALLQCLNRFLQTQNNVFP | 95 |
| Manes.14G109900.3 | PLAIIWFYTST-ILI-FVGQDHEISTGAGIFNRWMIPSLFAFALLQCLNRFLQTQNNVFP | 95 |
| AT1G73700.1 | PLSIIWANTEQ-ILV-LVHQDKSIASVAGSYAKYMIPSLFAYGLLQCINRFLQAQNNVFP | 176 |
| AT2G34360.1 | PLSIVWANTEH-FLV-FFGQDKSIAHLSGSYARFMIPSIFAYGLLQCLNRFLQAQNNVIP | 179 |
| AT5G52450.1 | PLSIIWANTEH-LLV-FFGQNKSIATLAGSYAKFMIPSIFAYGLLQCFNRFLQAQNNVFP | 178 |
| Sobic.009G106960.1 | ------------------------------------------------------------ | 0 |
| Manes.14G060800.1 | PISFIWVFTGQ-IFI-GLNQDPQISIHSGIYARWLIPAIVPYGLLQCQSRFLQTQNIVLP | 189 |
| Sobic.009G106800.1 | PIAVVWANAGR-ILV-LLGQDRDIAAEAGAYSRWLILGLVPYVPLACHIRFLQTQSVVVP | 175 |
| Sobic.007G074300.2 | ------------------------------------------------------------ | 0 |
| Sobic.009G106700.1 | PIALVWACAGR-ILL-FLGQDPEIAAEAGAYARWLIPSLAAYVPLQCHVRFLQTQSVVLP | 177 |
| Sobic.010G138400.4 | PLAVVWFYTGE-ILL-LFGQDADIAAEAGTYARWMIPLLFAYGLLQCHVRFLQTQNIVVP | 102 |
| Sobic.010G138400.1 | PLAVVWFYTGE-ILL-LFGQDADIAAEAGTYARWMIPLLFAYGLLQCHVRFLQTQNIVVP | 180 |
| Sobic.010G138400.5 | PLAVVWFYTGE-ILL-LFGQDADIAAEAGTYARWMIPLLFAYGLLQCHVRFLQTQNIVVP | 99 |
| Sobic.004G129900.1 | PVSLVWAYTGD-ILV-WFRQDREIAAGAGSYIRCMVPALFLFGQLQCHVRFLQPQNVVVP | 177 |
| Sobic.004G129900.2 | PVSLVWAYTGD-ILV-WFRQDREIAAGAGSYIRCMVPALFLFGQLQCHVRFLQPQNVVVP | 95 |
| Sobic.006G042200.1 | VVAIIWSYTGQ-ILL-LFGQDPEIAAGAGSYIRWMIPALFVYGPLQCHVRFLQTQNIVLP | 180 |
| Sobic.004G129700.1 | PVSVVWAYTGE-ILA-WCGQDPEIAAAAGIYIRWLIPALFLFGALQCHVRFLQTQNLVVP | 189 |
| Sobic.004G129800.1 | PVAVVWTYTGE-ILA-WCGQDPEIAAGAGSYIRWLIPALFAYGALQCHVRFLQTQNLVVP | 180 |
| Sobic.004G129800.2 | PVAVVWTYTGE-ILA-WCGQDPEIAAGAGSYIRWLIPALFAYGALQCHVRFLQTQNLVVP | 95 |
| AT3G23550.1 | LITILWFFTES-VFL-LLRQDPSISKQAALYMKYLAPGLLAYGFLQNILRFCQTQCIVTP | 183 |
| AT3G23560.1 | LITIFWFFTES-IFG-LLRQDPSISKQAALYMKYQAPGLLAYGFLQNILRFCQTQSIIAP | 191 |
| Manes.15G088100.1 | IISVIWFYTES-ILV-LLHQDPQISATAALYMKNLIPGLFAYGFLQNMLRFLQTQSTVMP | 203 |
| Manes.03G109100.1 | TISILWLYTEK-ILI-ILNQDPQISKEAALYIKYLIPGLFAYGFLQNIMRFLQTQSVVVP | 187 |
| Manes.03G109100.2 | TISILWLYTEK-ILI-ILNQDPQISKEAALYIKYLIPGLFAYGFLQNIMRFLQTQSVVVP | 148 |
| Sobic.002G311200.1 | AVSALWWFTEP-VLL-FLRQEPEVSRAAAAFVQAQVPGLFAFAFVQCLLRYLQTQSVVLP | 298 |
| Sobic.002G006500.3 | AVSLLWLYSEP-LLV-FLRQDPGTSRLAADFLRHSVPALFAYGFIQCALRFLQAQSVVAP | 212 |
| Sobic.002G006500.4 | AVSLLWLYSEP-LLV-FLRQDPGTSRLAADFLRHSVPALFAYGFIQCALRFLQAQSVVAP | 212 |
| AT2G04090.1 | LISILWFYMDK-LFV-SLGQDPDISKVAGSYAVCLIPALLAQAVQQPLTRFLQTQGLVLP | 182 |
| AT2G04100.1 | LISILWFYMDK-LFV-SLGQDPDISKVAGSYAVCLIPALLAQAVQQPLTRFLQTQGLVLP | 182 |
| AT2G04066.1 | ------------------------------------------------------------ | 0 |
| AT2G04040.1 | LISILWLYIEK-ILI-SLGQDPEISRIAGSYAFWLIPALFGQAIVIPLSRFLLTQGLVIP | 179 |
| AT2G04080.1 | IISILWIYIEK-LLI-TLGQEPDISRVAGSYSLWLVPALFAHAIFLPLTRFLLAQGLVIS | 179 |
| AT2G04050.1 | LISVLWIYIEK-LLI-SLGQDPDISRVAGSYALWLIPALFAHAFFIPLTRFLLAQGLVLP | 179 |
| AT2G04070.1 | LISILWIYMEK-LLI-SLGQDPDISRVAGSYALRLIPTLFAHAIVLPLTRFLLAQGLVLP | 179 |
| Sobic.001G481800.1 | PMAITWVFIPD-VLP-LIGQDPQIASEAGRYALWLIPGLFAFSVAQCLSKFLQSQSLIFP | 206 |
| Sobic.003G260200.1 | PLSLLWVFMDK-ILV-LIGQDPLISQGAGRYMIWMIPGLFANAVIQPLTKFLQTQSLIYP | 189 |
| Sobic.009G224800.1 | PISLLWAFIGK-LLM-LIGQDPLISKEAGRYIAWLIPGLFAYAISQPLTKFLQSQSLIIP | 192 |
| Sobic.009G224800.2 | PISLLWAFIGK-LLM-LIGQDPLISKEAGRYIAWLIPGLFAYAISQPLTKFLQSQSLIIP | 192 |
| AT1G66760.2 | PISILWMFMNQ-ILL-LLHQDPQIAELAGVYCLWLVPALFGYSVLESLVRYFQSQSLIYP | 180 |
| AT1G64820.1 | SISIVWFFMDK-LLE-IFHQDPLISQLACRYSIWLIPALFGFTLLQPMTRYFQSQGITLP | 181 |
| AT1G66780.1 | PISLLWVFMDK-LLE-LFHQDPLISQLACRYSIWLIPALFGYSVLQSMTRFFQSQGLVLP | 187 |
| Manes.02G072400.1 | PICLLWIFLDR-LLP-LIGQDPLISREACKYSIWLIPALFGSAILKPLTRFLQTQSVILP | 177 |
| Manes.01G113500.1 | PVSILWIFMAR-LLI-LVGLDPHISMAACKYSIGLIPALFGYAILQSLFRYFQSQSLILP | 165 |
| Manes.01G113500.2 | PVSILWIFMAR-LLI-LVGLDPHISMAACKYSIGLIPALFGYAILQSLFRYFQSQSLILP | 165 |
| AT1G15150.1 | PLSLLWFNMGK-LIV-ILGQDPAIAHEAGRYAAWLIPGLFAYAVLQPLIRYFKNQSLITP | 183 |
| AT1G15160.1 | PLSLLWFNMGK-LLV-ILGQDPSIAHEAGRFAAWLIPGLFAYAVLQPLTRYFKNQSLITP | 183 |
| AT1G15170.1 | PLSLIWFNMEK-LLL-ILGQDPSIAHEAGKYATWLIPGLFAYAVLQPLTRYFQNQSLITP | 186 |
| AT1G15180.1 | PLTLIWLNMET-LLV-FLGQDPSIAHEAGRYAACLIPGLFAYAVLQPLTRYFQNQSMITP | 187 |
| AT1G71140.1 | PLSLLWTYIGD-ILS-LIGQDAMVAQEAGKFATWLIPALFGYATLQPLVRFFQAQSLILP | 178 |
| Manes.06G063000.1 | PLSVLWIYMGK-ILV-FMGQDTLISQQAAKFSSCLIPALFGYANLQAVVRYFQMQSLIFP | 178 |
| Manes.14G109700.1 | AVSIIWFNMEK-LLL-LLGQDPLIAHEAGKFTSMLVPALFAYAIFQPLTKYFQTQSLTIP | 187 |
| Manes.14G109800.1 | PLSVIWINMGK-ILI-FIGQDPRISHEAGKFTMWLVPQLFAYATLQPLIRYFQTQSLIFP | 188 |
| Sobic.008G171600.1 | PIAFLYWYAGP-FLR-LIGQEADVAAAGQLYARGLMPQLLAFTLFSPMQRFLQAQNIVNP | 195 |
| Sobic.008G171600.2 | PIAFLYWYAGP-FLR-LIGQEADVAAAGQLYARGLMPQLLAFTLFSPMQRFLQAQNIVNP | 209 |
| Sobic.008G171600.3 | PIAFLYWYAGP-FLR-LIGQEADVAAAGQLYARGLMPQLLAFTLFSPMQRFLQAQNIVNP | 195 |
| AT3G59030.1 | FLTFLYWYSGP-ILK-TMGQSVAIAHEGQIFARGMIPQIYAFALACPMQRFLQAQNIVNP | 203 |
| Manes.01G182000.1 | LLTFLYWFSGS-VFV-AMGQSTAIAEQGQIFARGLIPQIYAFAICCPLQRFLQAQNIVNP | 202 |
| Manes.02G142000.1 | LLTFLYWFSGP-VLI-AMGQSEDIAEQGEIFARGLIPQLYAFAMSCPMQRFLQAQNVVNP | 202 |
| AT4G21903.2 | PMTILYTFSYP-ILL-LLGEPKTVSYMGSLYIAGLIPQIFAYAVYFTAQKFLQAQSVVAP | 204 |
| AT4G21910.4 | PMTLLYTFSYP-ILI-LLGEPKTVSYMGSKYIAGLIPQIFAYAVNFTAQKFLQAQSVVAP | 208 |
| AT1G11670.1 | PMTLLFIFSKP-LLI-SLGEPADVASVASVFVYGMIPMIFAYAVNFPIQKFLQSQSIVTP | 202 |

```
AT1G61890.1            PMSFLFLFSNP-ILT-ALGEPEQVATLASVFVYGMIPVIFAYAVNFPIQKFLQSQSIVTP  199
Manes.16G008000.1      PMTVIYLLSKP-ALL-LLGEPKKVASAAAVFVYGLIPQIFAYAVNFPIQKFLQAQSIVYP  202
Manes.17G038200.1      PMTVIYLLSKP-ILL-LLGEPPKMASAAAVFVYGLIPQIFAYAVNFPIQKFLQAQSIVNP  203
Sobic.002G318300.1     PLAAMYALSEP-LLL-LLGQSPEIAGAAAEFAYGLVPQIFAYAANFPIQKFLQAQSIVAP  192
Manes.17G038300.1      PLMLIYVFCKP-ILL-LLGEPNNIASAAEIFVFGLIPQIFAYAVNFPIQKFLQAQSIIAP  189
Manes.16G007900.1      PLTLIYIFSKP-ILV-LLGEPTDIATAAAVFVYGLIPQIFAYAANFPIQKFLQSQSIIAP  198
Manes.17G038400.1      PLTLVYIFSKP-ILI-LLGEPNDIAAAAAVFVYGLIPQIFAYAANFPIQKFLQSQSIIAP  198
Sobic.001G012600.1     PLAVAYGFSER-ILV-FLGESERIAHAAAVFVYGLIPQIFAYAANFPIQKFLQAQSIVAP  195
Sobic.001G012600.2     PLAVAYGFSER-ILV-FLGESERIAHAAAVFVYGLIPQIFAYAANFPIQKFLQAQSIVAP  195
Sobic.001G185600.1     PLAVVYVFSKQ-ILL-LLGESERIAEAAWVFVLGLIPQIFAYAFNFPIQKFLQAQSIVAP  220
Sobic.001G185400.1     PLAVVYAFSEP-ILV-FLGQSPEIARAASIFVYGLIPQIFAYAINFPIQKFMQAQSIVLP  194
AT3G21690.1            LLTLIYVFSEP-ILL-FLGESPAIASAASLFVYGLIPQIFAYAANFPIQKFLQSQSIVAP  204
Sobic.001G185800.1     PLSVIYAFSEP-ILV-FLGESPEIAKAAAVFVYGLIPQVFAYAVNFPIQKFLQAQSIVSP  198
Sobic.001G185500.1     PLAVIYAFSRP-ILV-LLGESPAIASAAAVFVYGLIPQIFAYAANFPIQKFMQAQSIMAP  212
Sobic.001G185500.2     PLAVIYAFSRP-ILV-LLGESPAIASAAAVFVYGLIPQIFAYAANFPIQKFMQAQSIMAP  95
Sobic.007G165500.2     IMTPLFVFAEP-LLL-LLGQDADVAREAARFSIYIIPSIYAMAINFGASKFLQAQSKVTV  229
AT4G00350.1            CLLPLYIYATP-LLI-LLGQEPEIAEISGKFTTQIIPQMFALAINFPTQKFLQSQSKVGI  240
Manes.01G255000.1      FLLPLYLYATP-ILK-ILGIEADIAVIAGRFTMQVIPQMFSLAINFPTQKFLQAQSKVGV  302
Sobic.004G349550.1     LLSPLYLLAAP-ILR-ALGQDEAVAAAAGDFTLRTLPQLLSLPLAFPTQKLLQAQGEVGA  95
Sobic.004G349600.1     LLTPLYVYAAP-VLR-LLGQDEGIAGAAGTFTRGIIPQMFALAVNFPAQKFLQAQSKVGV  199
Sobic.007G176000.1     LMLPLYLFATP-ILR-LFHQDAEIADLAGRLALYMIPQLFAYAFNFPIQKFLQAQSKVMA  203
Sobic.007G176100.1     LMLPLYLFATP-ILR-LFHQDAEIADLAGRLALYMIPQLFAYAFNFPIQKFLQAQSKVMA  208
AT1G47530.1            FLLPVYIWAPP-ILS-FFGEAPHISKAAGKFALWMIPQLFAYAANFPIQKFLQSQRKVLV  187
Manes.05G164500.1      LLVPIYVWSPP-ILE-LIGQTTQISTAAGKFALWMLPQLFAYAMNFPIQKFLQSQRKVFV  183
Manes.05G164500.2      LLVPIYVWSPP-ILE-LIGQTTQISTAAGKFALWMLPQLFAYAMNFPIQKFLQSQRKVFV  183
Manes.18G030900.1      LLVPIYVWSPP-ILE-LIGETTEISTAAGKFALWMLPQLFAYALNFPIQKFLQSQSKVFV  184
AT1G23300.1            LLCLFYVFATP-LLS-LLGQSPEISKAAGKFSLWMIPQLFAYAVNFATAKFLQAQSKVIA  196
AT3G26590.1            ILSLLYIFAAP-ILA-SIGQTAAISSAAGIFSIYMIPQIFAYAINFPTAKFLQSQSKIMV  197
AT5G38030.1            ILSLLYIFAAP-ILA-FIGQTPAISSATGIFSIYMIPQIFAYAVNYPTAKFLQSQSKIMV  197
Manes.12G023400.1      LMSFFYIFATQ-ILK-LIGQQAAISKAAGIYAIWMIPQLFAYAMNFPMAKFLQAQRKMIV  200
Manes.12G023500.1      LLSFLYIFATQ-ILK-LIGQQAAISKAAGIFSIWMIPQLFAYAMNFPMAKFLQAQRKMIV  200
Manes.12G023600.1      LLSLIYIFAAQ-LLK-LIGQTEAISKTAGVFAIWMIPQLFAYAMNFPMAKFLQAQSKIMV  217
Manes.13G025200.1      LLSLIYIFAAQ-ILK-LIGQTEAISKAAGIFSIWMIPQLFAYAVNFPMGKFLQAQSKVMV  217
Manes.13G025200.2      LLSLIYIFAAQ-ILK-LIGQTEAISKAAGIFSIWMIPQLFAYAVNFPMGKFLQAQSKVMV  216
Sobic.001G273100.1     ALAPAYVLAAP-LLHRSLHQPGAVSRAAGPYARWAVPRLLAHALNIPLLMFFQAQSRIWA  194
Sobic.001G273100.2     ALAPAYVLAAP-LLHRSLHQPGAVSRAAGPYARWAVPRLLAHALNIPLLMFFQAQSRIWA  194
Sobic.001G273000.2     ALAPVYIFATP-ILQFFLHQPVDVSRAAGQYARWAIPRLFANAMDIPLLMFFRGQSRVWT  167
Sobic.001G273000.1     ALAPVYIFATP-ILQFFLHQPVDVSRAAGQYARWAIPRLFANAMDIPLLMFFRGQSRVWT  178
Sobic.001G273000.3     ALAPVYIFATP-ILQFFLHQPVDVSRAAGQYARWAIPRLFANAMDIPLLMFFRGQSRVWT  178
Manes.15G147800.1      VLTPFYVFASP-LLQ-FLHQDKDISKLAGKYSIWVIPQLFAYAINYPIQKFLQAQSRVWI  182
Manes.15G147900.1      VFTPFYVFASP-LLQ-LLHQDKDISNLAGKYSIWVIPQLFAYAVNYPIQKFLQAQSRVWV  183
Manes.17G098800.1      FLAPFYVFASP-ILQ-LLHQDKGISKLAGKYSIWVIPQLFAYALNFPIQKFLQAQSRVWI  188
Manes.17G098900.1      FLAPFYVFASP-ILQ-LLHQDKGISKLAGRYSIWVIPQLFAYALNFPIQKFLQAQSRVWI  188
Manes.17G098900.2      FLAPFYVFASP-ILQ-LLHQDKGISKLAGRYSIWVIPQLFAYALNFPIQKFLQAQSRVWI  95
Sobic.001G162400.1     VLTPTYLFTAP-ILR-ALRQPGDVARVAGTYARWVAPQLFAYAANFPLQKFFQSQSRVWV  173
Sobic.003G307600.2     VLLPLYVFTSP-ILR-LLRQSADISAVSGRYARWCVPQLFAYAVNFPMQKFYQAQSRVWV  95
Sobic.002G232200.1     LLAPVYVFSGQ-LLA-AFGQPAELARAAGSVSVYFLPSHFMYAIHLPVMTFLQCQRKNWV  191
Sobic.002G232500.1     LLTPTYIFSEQ-LLT-ALGQPAELSRQAGLVSLYMLPLHFVYAIVLPLNKFLQCQRKNWV  187
Sobic.002G232600.1     LLAPTYIFSGQ-LLM-VLGQPAELSREAGLLGMYLLPLHLMFAIQLPLNKFLQCQRKNWV  182
Sobic.007G160700.1     ALTPTYIFMED-LLL-LIGQTPELSRLAGQMSVWLLPQHFAMAMLLPLTRFLQSQLKNWV  199
Sobic.003G126200.2     LLTPMYFFTED-LLL-VTGQPPKLSAMAGRVSMWFIPLHFSQAFLFPLQLFLQCQRKNLA  210
Sobic.001G476700.1     LLLPMYFFAED-VLL-LTGQSPELSAMAGKVSVWFIPLHFSFAFLFPLQRFLQCQMKNFA  207
Sobic.001G476700.2     LLLPMYFFAED-VLL-LTGQSPELSAMAGKVSVWFIPLHFSFAFLFPLQRFLQCQMKNFA  230
Manes.18G062800.1      LLLPVYVFASP-ILE-WLGQPANVAEETCVVAMWLLPLHFSFAFIFPLQRFLQSQLKNQV  190
AT5G44050.1            LLLPMYIFATP-ILK-FMGQPDDIAELSGIISVWAIPTHFSFAFFFPINRFLQCQLKNSV  192
AT5G10420.1            LLLPMYLFATP-ILK-FIGQSDDIAELTGTIALWVIPVHFAFAFFFPLNRFLQCQLKNKV  190
AT5G65380.1            LLLPTYIFTTP-VLK-FLGQPDDIAELSGVVAIWVIPLHFAFTLSFPLQRFLQCQLKNRV  189
Manes.12G129000.1      LLLPLYLFASP-ALK-LLGQPKDVAELSGIVSVWMIPLHFSFAFQFPLQRFLQSQLKNTI  191
Manes.13G097900.1      LLLPMYIFASP-VLK-LLGQPKDVAELSGVVSVCMIPFHFSFAFQFPLQRFLQSQLKNTV  191
Sobic.005G020700.1     VLLPIYLFTAP-LLV-ALGQDPEIAVVAGTISLWYIPVMFSYVWAFTLQMYLQAQSKNII  223
Sobic.008G019400.1     VLLPVYLFTEP-LLV-ALGQDPEISAVAGTISLWYIPVMFSYVWAFTLQMYLQAQSKNMI  219
Manes.09G135300.1      CLLPLFIFTTY-ILR-ALGQDESIVEVAGNISLWLIPVMFSFIPSFTCQMYLQAQSKNMI  182
AT1G33080.1            CLMPIYIFAGP-ILL-ALGQEERLVRVARIIALWVIGINISFVPSFTCQMFLQAQSKNKI  188
AT1G33090.1            CIMPIFIFSGP-ILL-ALGQEDHIVRVARVIALWLIAINFTFVPAFTCQIFLQSQSKNKI  188
AT1G33100.1            CLMPVFIFAGP-ILL-ALGQEERIVRVARVLALWVIGINFSFVPSFTCQMFLQAQSKNKI  185
AT1G33110.1            CLTPVYIFSGP-ILL-ALGQEERIVRVARIIALWVIGINFSFVPSFTCQMFLQAQSKNKI  188
AT3G03620.1            LFLPFIVLAGP-ILR-LLGQNVEITKTVDEIYPWMIPYVYSLIFTMTIQMYLQAQMRNAI  189
AT5G17700.1            LFVPFIVLAGP-ILR-LLGQNVVISETVDEIYPWVIPYLYSIVFTMTMQMYLQAQMKNAI  186
Manes.09G135200.1      IVVPLFIFATP-ILR-LIGQVEEIAVVAGKISLWFIPFLYYFVFSLTMQMYLQTQMKNQI  180
Manes.09G134800.1      VLLPVFIFSSQ-VLK-LLGQEEDTARKAGIISLWFIPSIYSSILNFTMEKFLQAQLKNIV  177
Manes.08G150400.1      MLLPVFIFSGK-IFR-LLGEEEEIADSAGYISVLFIPMLYFFALAFSLQKFLQTQLKNMI  175
```

```
Manes.09G134900.1       ILLPVFIFSGK-IFR-LLGEEEGIANTAGYISLWFIPMLYFFSLAFAIQKYLQTQLKNMI  176
Manes.09G135000.1       ILLPVFIFSGK-IFR-LLGEEEDIANTAGYISLWFIPMLYFFSLAFAIQKYLQTQLKNMI  176
Manes.09G134900.2       ILLPVFIFSGK-IFR-LLGEEEGIANTAGYISLWFIPMLYFFSLAFAIQKYLQTQLKNMI  147
Manes.09G135000.2       ILLPVFIFSGK-IFR-LLGEEEDIANTAGYISLWFIPMLYFFSLAFAIQKYLQTQLKNMI  147


Sobic.002G099300.1      ---------MAEWGGKNENLPLMETTHTADFSADT-NDHCLTLYFGCGIVKVS-------  79
AT4G39030.1             FILVGLVAQSASLGMKNSWGPLKA-LAAATIINGL-GDTILCLFLGQGIAGAAWATTASQ  290
Manes.04G084700.1       AILYGLVCQSSSLGMKDSLGPLNA-LVVASAVNAI-GHLVLCSWLGYGIIGAAWATMTSQ  310
Manes.11G091900.1       AILIGWVAQSASLGMKDSLGPLKA-LIVASAINGI-GDIFLCRFLDYGIAGAAWATMVSQ  308
Sobic.004G019800.2      AVLVGLVAQSASLGMKDSWGPLKA-LAAASVINGV-GDIFLCSVCGYGIAGAAWATMVSQ  172
Sobic.004G019800.3      AVLVGLVAQSASLGMKDSWGPLKA-LAAASVINGV-GDIFLCSVCGYGIAGAAWATMVSQ  172
Sobic.004G019800.4      AVLVGLVAQSASLGMKDSWGPLKA-LAAASVINGV-GDIFLCSVCGYGIAGAAWATMVSQ  172
AT2G21340.1             AVLIGWVAQSASLGMKDSWGPLKA-LAVASAINGV-GDVVLCTFLGYGIAGAAWATMVSQ  307
Manes.11G092000.1       AVIVGWVAQSASLGMKDSWGPLKA-LAVSSIFNFV-GDVVLCSFLGYGIAGAAWATMVSQ  305
Sobic.001G476700.3      ------------------------------------------------------------  164
Sobic.009G077000.1      ------------------------------------------------------------  80
Sobic.001G454900.1      AVLLSLAMQGVFRGFKDTKTPLYA-TVVGDAANII-LDPILMFVCHMGVTGAAIAHVVSQ  328
Sobic.003G403000.1      AVLLSLAMQGVFRGFKDAKTPLYA-IVAGDAANIV-LDPILIFGCRLGVIGAAIAHVLSQ  363
Sobic.003G403000.2      AVLLSLAMQGVFRGFKDAKTPLYA-IVAGDAANIV-LDPILIFGCRLGVIGAAIAHVLSQ  363
Sobic.003G403000.3      AVLLSLAMQGVFRGFKDAKTPLYA-IVAGDAANIV-LDPILIFGCRLGVIGAAIAHVLSQ  363
Sobic.007G020600.1      AVLLSLATQGVFRGFKDTKTPLYA-TVAGDAINIV-LDPIFIFVFQYGVSGAAIAHVISQ  288
Sobic.007G020600.2      AVLLSLATQGVFRGFKDTKTPLYA-TVAGDAINIV-LDPIFIFVFQYGVSGAAIAHVISQ  288
AT3G08040.1             ALLLSLAMQGIFRGFKDTKTPLFA-TVVADVINIV-LDPIFIFVLRLGIIGAAIAHVISQ  282
Manes.09G027700.1       ------------------------------------------------------------  156
Manes.09G027800.1       ------------------------------------------------------------  2
Manes.09G027900.1       ------------------------------------------------------------  2
Manes.09G026900.1       AVLLSLAMQGVFRGIKDTKTPLFA-TVVGDVANII-LDPIFIFVFRLNVSGAAIAHVISQ  271
Manes.09G026900.2       AVLLSLAMQGVFRGIKDTKTPLFA-TVVGDVANII-LDPIFIFVFRLNVSGAAIAHVISQ  271
Manes.09G027000.1       ------------------------------------------------------------  0
Manes.07G006000.1       AVLLSLAMQGVFRGFKDTKTPLYA-TVAGDVANII-LDPIFIFACRLGVSGAAIAHVLSQ  306
Manes.07G006000.2       AVLLSLAMQGVFRGFKDTKTPLYA-TVAGDVANII-LDPIFIFACRLGVSGAAIAHVLSQ  306
Manes.10G143000.1       AVLLSLAMQGVFRGFKDTKTPLYA-TVAGDVTNII-LDPIFIFVCRLGVSGAAIAHVLSQ  299
AT1G51340.2             AVLLSLAAQGVFRGFKDTTTPLFA-TVIGDVTNII-LDPIFIFVFRLGVTGAATAHVISQ  278
Manes.06G164500.1       AVLLSLAMQGVFRGFKDTKTPLYA-TVAGDVTNII-LDPLFMFVFRLGVSGAAIAHVISQ  271
Manes.06G164500.2       AVLLSLAMQGVFRGFKDTKTPLYA-TVAGDVTNII-LDPLFMFVFRLGVSGAAIAHVISQ  271
Manes.06G164500.3       AVLLSLAMQGVFRGFKDTKTPLYA-TVAGDVTNII-LDPLFMFVFRLGVSGAAIAHVISQ  271
Manes.06G164500.4       AVLLSLAMQGVFRGFKDTKTPLYA-TVAGDVTNII-LDPLFMFVFRLGVSGAAIAHVISQ  271
Manes.06G164500.5       AVLLSLAMQGVFRGFKDTKTPLYA-TVAGDVTNII-LDPLFMFVFRLGVSGAAIAHVISQ  271
Manes.14G002600.1       AVLLSLAMQGVFRGFKDTKTPLYA-TVAGDITNII-LDPIFMFIFRLGVSGAAIAHVISQ  255
Sobic.008G006100.1      PIIVALASQGAFRGFLDTRTPLYA-VGAGNLLNAV-LDALLIFPLGLGVSGAALATVTSE  312
Sobic.005G005400.1      PIVIALAAQGAFRGLMDTKTPLYA-VGVGNLINVI-LDAILVFPLGLGVRGAALATVTSE  296
Sobic.005G005400.3      PIVIALAAQGAFRGLMDTKTPLYA-VGVGNLINVI-LDAILVFPLGLGVRGAALATVTSE  296
Sobic.005G005400.2      PIVIALAAQGAFRGLMDTKTPLYA-VGVGNLINVI-LDAILVFPLGLGVRGAALATVTSE  296
AT2G38330.1             PIVVALAAQGAFRGFKDTTTPLYA-VVAGNVLNAV-LDPILIFVLGFGISGAAAATVISE  278
Manes.08G096600.1       PIVIALAAQGTFRGFKDTKTPLYA-IGAGNLLNAV-LDPILIFVFGFGIGGAAISTVISE  304
Manes.08G096600.2       PIVIALAAQGTFRGFKDTKTPLYA-IGAGNLLNAV-LDPILIFVFGFGIGGAAISTVISE  304
AT4G38380.1             AYVVSLALQGIFRGFKDTKTPVYC-LGIGNFLAVF-LFPLFIYKFRMGVAGAAISSVISQ  317
Sobic.002G286800.1      AVVVSLAIQGVFRGLKDTKTPLLY-SGLGNISAVV-LLPFFVYYLNLGLTGAALATIASQ  334
Sobic.003G149300.2      ANVLMLAVQGIFRGFKDTKTPVFY-IGLGNLSAVA-LLPLLIYGFQLGITGAAISTVVSQ  304
Sobic.003G149300.1      ANVLMLAVQGIFRGFKDTKTPVFY-IGLGNLSAVA-LLPLLIYGFQLGITGAAISTVVSQ  304
Sobic.003G149300.3      ANVLMLAVQGIFRGFKDTKTPVFY-IGLGNLSAVA-LLPLLIYGFQLGITGAAISTVVSQ  304
Sobic.003G149300.4      ANVLMLAVQGIFRGFKDTKTPVFY-IGLGNLSAVA-LLPLLIYGFQLGITGAAISTVVSQ  180
Manes.01G153400.1       -------------LFLFHLLLLLL-LGLSNLSALF-LFPVLMHYYHLGVTGAAISTVVSQ  54
Manes.04G064900.1       AVVVSLALQGIFRGFKDTKSPVYC-LALGNLSAIF-LFPILMYYFKLGVTGAAISTVVSQ  345
Manes.04G064900.2       AVVVSLALQGIFRGFKDTKSPVYC-LALGNLSAIF-LFPILMYYFKLGVTGAAISTVVSQ  345
Sobic.001G003700.1      TLFA---------------------AALALALHIP-LTVSLSA--RMGVRGVAAAVWLSD  208
AT4G22790.1             IMFT---------------------TAAATSLHIP-INIVLSK--ARGIEGVAMAVWITD  218
Manes.02G032300.1       IMFS---------------------SAVGLALHIP-INIFLAK--AKGLEGISMAIWITD  230
Manes.10G000400.1       IAIV---------------------VIFSMILHFP-INYLLVIHLNLGVKGVALASFWYT  196
AT2G38510.1             LTIS---------------------AIVSILLHPL-FNYVFVVRMRLGVKGVAIAMAFNT  196
Manes.08G172300.1       LTIC---------------------AVGAVILHTP-FNYFFAIYLNLGVKGVALAIACNT  226
Manes.09G117000.1       LTVS---------------------AVAAVILHAP-INYFFAIYLKLGVKGVALAIAFNT  196
Sobic.010G167800.1      LAAA---------------------AGAAVLFHAP-TNYVLVARLELGAPGVAAAASASN  250
AT5G19700.1             LTLA---------------------TLAGTIFHIP-MNFFLVSYLGWGFMGVSMAAAASN  226
AT4G29140.1             VTLA---------------------SLSGAVFHLP-ANLFLVSYLRLGLTGVAVASSITN  246
Manes.03G026500.1       LTLA---------------------SLTGTVLHLP-VNLLLVNHLKLGVSGVATAAAASN  253
Manes.16G109300.1       LTLA---------------------SLIGTILHLP-INFLLVNHLKLGVSGVAAAATVSN  247
Sobic.007G181100.1      LTYA---------------------AAAALLLHVP-VNCLLVHSLRLGIRGVALGAVCTN  235
Manes.13G127800.1       LMLS---------------------AAFALALHAL-INQILVCHLGLGIQGIAVAVTITD  245
Sobic.001G446800.1      ITAC---------------------SLFSVLLHGP-INYLLVVRLQMGVAGVALAVALTD  243
```

```
AT5G52050.1          LSIC---------------------TVIASFLHLP-ITFFLVSYLGLGIKGIALSGVVSN  225
AT1G58340.1          VTYS---------------------TAVSVLLHVP-LNYLLVVKLEMGVAGVAIAMVLTN  248
Manes.05G186700.1    LTYC---------------------SAISVLLHIP-LNFLLVVHFKLGIAGVAIAMVWTN  239
Manes.18G054300.1    LTYC---------------------SAISVLLHVP-LNFLLVVHFKMGIAGVAIAMVWTN  239
Sobic.001G320900.1   LTAC---------------------AVLAVAMHLP-INYLLVSVLGLGVEGVALASALAN  267
Sobic.001G320900.2   LTAC---------------------AVLAVAMHLP-INYLLVSVLGLGVEGVALASALAN  267
Sobic.004G283500.1   LTLC---------------------AALAIALHLP-INYVLVSVLGLGISGVALASVLAN  268
Sobic.006G184400.1   LTVC---------------------AALAIAIHLP-INYVLVTVLGLGIRGVAFASVLAN  281
AT4G23030.1          LTYS---------------------AFFAVLLHIP-INYLLVSSLGLGLKGVALGAIWTN  223
Manes.01G067000.1    LTFC---------------------AALSILFHIP-INYFLVSVLNLGIKGVALSGIWTN  243
Manes.02G027800.1    LTYC---------------------AAISILLHIP-INYFLVSVLNLGTKGVALSGVWTN  247
Manes.06G143600.1    LTFC---------------------ATLSMVLHIP-INYLLVTHLDLGIKGVALSGVWTN  255
Manes.14G029100.1    LTFC---------------------AILSILLHIP-INYLLVTHLNLGIKGVALSGVWTN  259
AT5G49130.1          LMWC---------------------TLVSVLLHLP-ITAFFTFYISLGVPGVAVSSFLTN  216
Manes.03G198000.1    LMCC---------------------TSVSIVLHLP-ITIFLAFTLRLGVPGIAISTFISN  209
Manes.15G011000.1    LMWC---------------------TLISIVLHLP-ITIFLAFTLQHGVPGIAISTCISN  209
Sobic.001G019700.1   MAAC---------------------SAIAVVLHVP-LNVALVFGMGLGVRGVAAAQALTN  233
AT1G71870.1          MMWC---------------------TLAAVAFHVP-LNYWLVMVKHWGVPGVAIASVVTN  214
Manes.02G189800.1    IMYC---------------------SLLAVIFHVP-LNYILVVVMGLGVPGVAMASVVTN  212
Manes.18G098800.1    IMYC---------------------SLMAVVFHVP-LNYMLVVMMGLGVPGVAMASVVTN  212
Sobic.010G256932.1   LVAC---------------------SGLTLLLHVM-LCWLLVQIFGIGHKGAALATSISY  189
Sobic.010G256700.1   LVAC---------------------SGLTLLLHVM-LCWLLVQIFGIGHKGAALATSISY  220
Sobic.010G256700.5   LVAC---------------------SGLTLLLHVM-LCWLLVQIFGIGHKGAALATSISY  220
Sobic.010G256700.6   LVAC---------------------SGLTLLLHVM-LCWLLVQIFGIGHKGAALATSISY  220
Sobic.010G256700.4   LVAC---------------------SGLTLLLHVM-LCWLLVQIFGIGHKGAALATSISY  220
Sobic.009G106900.1   -MAS---------------------SAVTALAHVPRVLRALVYRVGMGSKGAALSAAVSY  38
Manes.S031500.1      MMII---------------------SGNTTLLHIF-ICWVLVFKSGLGNKGAAMANAISY  224
Manes.14G109900.1    MMVS---------------------SGITASLHIL-ICWVLVFKSGLGSKGAAMAITISY  234
Manes.14G109900.2    MMVS---------------------SGITASLHIL-ICWVLVFKSGLGSKGAAMAITISY  133
Manes.14G109900.3    MMVS---------------------SGITASLHIL-ICWVLVFKSGLGSKGAAMAITISY  133
AT1G73700.1          VFVC---------------------SGITTCLHLL-LCWLFVLKTGLGYRGAALAISVSY  214
AT2G34360.1          VVIC---------------------SGVTTSLHVI-ICWVLVLKSGLGFRGAAVANAISY  217
AT5G52450.1          VVFC---------------------SGVTTSLHVL-LCWVLVFKSGLGFQGAALANSISY  216
Sobic.009G106960.1   ------------------------------------------------------------  0
Manes.14G060800.1    LVLS---------------------TGITSLVHVL-ICWTFIFRFGFGNKAAALSIAISY  227
Sobic.009G106800.1   VMVS---------------------SGVTALGHAV-VCWALVFKAGMGSKGAALSIAISY  213
Sobic.007G074300.2   -MAS---------------------SGATALCHLA-VCWALVHRAGMGSKGAALSNAVSY  37
Sobic.009G106700.1   VTAS---------------------SGATALCHLL-VCWALVYRAGMGSKGAALSNAVSY  215
Sobic.010G138400.4   VMAS---------------------AGATAACHLV-VCWVLVYPLGMGSKGAALSNAVSY  140
Sobic.010G138400.1   VMAS---------------------AGATAACHLV-VCWVLVYPLGMGSKGAALSNAVSY  218
Sobic.010G138400.5   VMAS---------------------AGATAACHLV-VCWVLVYPLGMGSKGAALSNAVSY  137
Sobic.004G129900.1   VMLS---------------------SGATVAVHVA-VCWLLVRRLGLGADGAALANAVSN  215
Sobic.004G129900.2   VMLS---------------------SGATVAVHVA-VCWLLVRRLGLGADGAALANAVSN  133
Sobic.006G042200.1   VMLS---------------------SGATALNHLL-VCWLLVYKIGMGNKGAALANAISY  218
Sobic.004G129700.1   VMLS---------------------SGATALCHPA-VCWLLVRALGLGSNGAALANAISY  227
Sobic.004G129800.1   VMLS---------------------AGATAVCHPA-VCWLLVRALGLGRNGAALANAVSY  218
Sobic.004G129800.2   VMLS---------------------AGATAVCHPA-VCWLLVRALGLGRNGAALANAVSY  133
AT3G23550.1          LVLF---------------------SFLPLVINIG-TTYALVHLAGLGFIGAPIATSISL  221
AT3G23560.1          LVIF---------------------SFVPLVINIA-TAYVLVYVAGLGFIGAPIATSISL  229
Manes.15G088100.1    LVLF---------------------SFIPMCIHIG-IAYALIYCTTLGFKGAPLAVSISL  241
Manes.03G109100.1    LVLC---------------------SAIPMFIHIG-ITYGLVHCANFGFKGAPLAASISL  225
Manes.03G109100.2    LVLC---------------------SAIPMFIHIG-ITYGLVHCANFGFKGAPLAASISL  186
Sobic.002G311200.1   LVVC---------------------SVAPFALHVA-LTHLLVNVLGLGLAGAGAAVSATF  336
Sobic.002G006500.3   LVAF---------------------SLLPLAAHIG-VAHALVNALGMGFAGAAVATSVSL  250
Sobic.002G006500.4   LVAF---------------------SLLPLAAHIG-VAHALVNALGMGFAGAAVATSVSL  250
AT2G04090.1          LLYC---------------------AITTLLFHIP-VCLILVYAFGLGSNGAALAIGLSY  220
AT2G04100.1          LLYC---------------------AITTLLFHIP-VCLILVYAFGLGSNGAALAIGLSY  220
AT2G04066.1          ------------------------------------------------------------  0
AT2G04040.1          LLFT---------------------AVTTLLFHVL-VCWTLVFLFGLGCNGPAMATSVSF  217
AT2G04080.1          LLYS---------------------AMTTLLFHIA-VCWTLVFALGLGSNGAAIAISLSF  217
AT2G04050.1          LLYC---------------------TLTTLLFHIP-VCWAFVYAFGLGSNGAAMAISVSF  217
AT2G04070.1          LLYF---------------------ALTTLLFHIA-VCWTLVSALGLGSNGAALAISVSF  217
Sobic.001G481800.1   MVLS---------------------SFTTLAVFIP-LCWFMVYKVGMGNAGAAFAVSICD  244
Sobic.003G260200.1   LLLS---------------------SVATAAIHIP-LCYVMVFKTGLGYTGAALTISISY  227
Sobic.009G224800.1   MLWS---------------------SIATLLLHIP-ICWLLVFKTSLGYIGASLAISLSY  230
Sobic.009G224800.2   MLWS---------------------SIATLLLHIP-ICWLLVFKTSLGYIGASLAISLSY  230
AT1G66760.2          MVLS---------------------SLAALSFHVP-LCWLMVHKFDFGAKGAAASIGISY  218
AT1G64820.1          LFVS---------------------SLGALCFHIP-FCWLLVYKLKFGIVGAALSIGFSY  219
AT1G66780.1          LFLS---------------------SLGALFFHVP-FSWLLVYKLRFGIVGAALSIGFSY  225
Manes.02G072400.1    MLLS---------------------SLLILFFHST-ACWTFVYKLGLGYKGTALAFGLSI  215
Manes.01G113500.1    MLLS---------------------SCAALCFHVP-FCWVLIYKWELGNIGGAIAIDVAY  203
```

```
Manes.01G113500.2       MLLS---------------------SCAALCFHVP-FCWVLIYKWELGNIGGAIAIDVAY  203
AT1G15150.1             LLVT---------------------SSVVFCIHVP-LCWLLVYKSGLGHIGGALALSLSY  221
AT1G15160.1             LLIT---------------------SCVVFCLHVP-LCWLLVYKSGLDHIGGALALSLSY  221
AT1G15170.1             LLIT---------------------SYVVFCIHVP-LCWFLVYNSGLGNLGGALAISLSN  224
AT1G15180.1             LLIT---------------------SCFVFCLHVP-LCWLLVYKSGLGNLGGALALSFSN  225
AT1G71140.1             LVMS---------------------SVSSLCIHIV-LCWSLVFKFGLGSLGAAIAIGVSY  216
Manes.06G063000.1       LIIS---------------------SLFAVCFHVV-VCWILVFNSGLGNLGAAFSIGISY  216
Manes.14G109700.1       MLIS---------------------SCVTLCLHIP-LCWTLVFKSGLRNLGGALAISVSH  225
Manes.14G109800.1       MVLS---------------------SCGALCFHIP-LCWVLVFKSGLDNLGAAVAMCISN  226
Sobic.008G171600.1      VAYI---------------------TLAVLIFHTL-ASWLGVFVLGFGLLGAALILSFSW  233
Sobic.008G171600.2      VAYI---------------------TLAVLIFHTL-ASWLGVFVLGFGLLGAALILSFSW  247
Sobic.008G171600.3      VAYI---------------------TLAVLIFHTL-ASWLGVFVLGFGLLGAALILSFSW  233
AT3G59030.1             LAYM---------------------SLGVFLLHTL-LTWLVTNVLDFGLLGAALILSFSW  241
Manes.01G182000.1       LACN---------------------AVGVFFVHVF-LSWLVIYKLDYGLLGAALTLSLSW  240
Manes.02G142000.1       LAYM---------------------SVGVFFVHIL-LSWLVIYKLEYGLLGAALTLSFSW  240
AT4G21903.2             SAYI---------------------SAAALVLQIS-LTWITVYAMGQGLMGIAYVLTISW  242
AT4G21910.4             SAFI---------------------SAAALILQIL-LTWITVYVMDMGFMGIAYVLTISW  246
AT1G11670.1             SAYI---------------------SAATLVIHLI-LSWLSVFKFGWGLLGLSVVHSLSW  240
AT1G61890.1             SAYI---------------------SAATLVIHLI-LSWIAVYRLGYGLLALSLIHSFSW  237
Manes.16G008000.1       SAII---------------------SAATLGVHLL-VTWVALFKPGMGLVGASLALSLSW  240
Manes.17G038200.1       SAII---------------------SAATLVLHLL-LSWIAVFQLGMGLLGASLILSLSW  241
Sobic.002G318300.1      SAYI---------------------LAASFALHVP-LSWLAVYGLGLGLLGASLTLSLTW  230
Manes.17G038300.1       SAYI---------------------SLAGSVLHII-FTWLAVYKFNWGLLGAALVLSFSW  227
Manes.16G007900.1       SAYI---------------------SLVALAVHIL-FTWLAVFKWNWGLLGAALILSLSW  236
Manes.17G038400.1       SAYI---------------------SLVALGVHVL-FTWLGVFKWNWGLLGAALILSLSW  236
Sobic.001G012600.1      SAYI---------------------STATLALHLA-LTWLAVDRLGMGLLGGALVLSLSW  233
Sobic.001G012600.2      SAYI---------------------STATLALHLA-LTWLAVDRLGMGLLGGALVLSLSW  233
Sobic.001G185600.1      SAYI---------------------STAALAGHLV-LSWLAVYRMGLGLLGASLILSLSW  258
Sobic.001G185400.1      SAYI---------------------STATLALHLL-LSWVVVYKAGLGLLGASLVLSLSW  232
AT3G21690.1             SAYI---------------------STATLFVHLL-LSWLAVYKLGMGLLGASLVLSLSW  242
Sobic.001G185800.1      SAYI---------------------SAATLAVHLV-LGWLVVYRFGMGLLGASLVLSLSW  236
Sobic.001G185500.1      SAYI---------------------SAATLAVHVA-LSYLAVYRWGMGLLGASLILSLSW  250
Sobic.001G185500.2      SAYI---------------------SAATLAVHVA-LSYLAVYRWGMGLLGASLILSLSW  133
Sobic.007G165500.2      PAYI---------------------GFGALLINVL-LNYLFVYVLGWGLPGAAAAYDVAH  267
AT4G00350.1             MAWI---------------------GFFALTLHIF-ILYLFINVFKWGLNGAAAAFDVSA  278
Manes.01G255000.1       LAWI---------------------GLAALIIHIG-LLYLFINIFKWGLAGAAAAYDVSA  340
Sobic.004G349550.1      LAWI---------------------GAAALAAHVA-MLALFVPVLGWGLRGAAAAYDVTS  133
Sobic.004G349600.1      MAWI---------------------GLAALLAHVA-LLALLVSVLGWGVAGAALAYDTSS  237
Sobic.007G176000.1      MAAV---------------------SAAALAFHVA-LSWFLVGPMRMGLVGLAVALNASW  241
Sobic.007G176100.1      MAAV---------------------SAAALAFHVA-LSWFLVGPMRMGLVGLAVALNASW  246
AT1G47530.1             MAWI---------------------SGVVLVIHAV-FSWLFILYFKWGLVGAAITLNTSW  225
Manes.05G164500.1       MAWI---------------------SAVVLLLHAF-FSWLLILKLGWGLTGAAITLNTSW  221
Manes.05G164500.2       MAWI---------------------SAVVLLLHAF-FSWLLILKLGWGLTGAAITLNTSW  221
Manes.18G030900.1       MAWV---------------------SAVVLVLHAI-FSWLLILKLGWGLTGAAITLNTSW  222
AT1G23300.1             MAVI---------------------AATVLLQHTL-LSWLLMLKLRWGMAGGAVVLNMSW  234
AT3G26590.1             MAVI---------------------SAVALVIHVP-LTWFVIVKLQWGMPGLAVVLNASW  235
AT5G38030.1             MAAI---------------------SAVALVLHVL-LTWFVIEGLQWGTAGLAVVLNASW  235
Manes.12G023400.1       MAMI---------------------SAAALVLHTF-FSWLLMLKLGWGLVGAVVVLNASW  238
Manes.12G023500.1       MAVI---------------------SAAALVLHTF-FSWLLMLKLGWGLVGAAVVLNASW  238
Manes.12G023600.1       MALI---------------------SAAAFVLHTL-FSWLLMLKLGWGLVGAAVVLNASW  255
Manes.13G025200.1       MALI---------------------AAAALVLHTV-FSWLLMLKLGWSLVGAAVVLNASW  255
Manes.13G025200.2       MALI---------------------AAAALVLHTV-FSWLLMLKLGWSLVGAAVVLNASW  254
Sobic.001G273100.1      VAAI---------------------SGAALCAHAV-LTYVAVARLGYGLPGAAVAGDVSH  232
Sobic.001G273100.2      VAAI---------------------SGAALCAHAV-LTYVAVARLGYGLPGAAVAGDVSH  232
Sobic.001G273000.2      LAAI---------------------SGVALAVHTV-LTYIAVRQLGYGLPGAAVAGDISQ  205
Sobic.001G273000.1      LAAI---------------------SGVALAVHTV-LTYIAVRQLGYGLPGAAVAGDISQ  216
Sobic.001G273000.3      LAAI---------------------SGVALAVHTV-LTYIAVRQLGYGLPGAAVAGDISQ  216
Manes.15G147800.1       MTII---------------------SIVALAFHVL-LNWVLVTKLNHGLLGAAIAGNISW  220
Manes.15G147900.1       MTII---------------------SVVALAFHVL-LNWVLVTKLDHGLAGAAIAGNISW  221
Manes.17G098800.1       MTII---------------------SIAALAIHVL-LNWVLVTKLDHGLVGAAVAGNISW  226
Manes.17G098900.1       MTII---------------------SIAALAIHVL-LNWVLVTKLDHGLVGAAVAGNISW  226
Manes.17G098900.2       MTII---------------------SIAALAIHVL-LNWVLVTKLDHGLVGAAVAGNISW  133
Sobic.001G162400.1      VTAV---------------------SGAGLAVHVV-LNYVVVARLGHGLLGAAVVGNVTW  211
Sobic.003G307600.2      MTAI---------------------SGAVLAVHAL-LNWLVVSRLGRGLVGAAVVGDVSW  133
Sobic.002G232200.1      PTVA---------------------TAAVFAVHVA-ATWLLVNCLGLGVFGVAMAFNISW  229
Sobic.002G232500.1      AAVT---------------------TAAAFPVHVV-ATWLLVRCFRLGVFGAAMALTLSW  225
Sobic.002G232600.1      IALS---------------------SVLGFPVHVV-ATWLLAQRFQLGVLGAAMSLNLSW  220
Sobic.007G160700.1      TAIT---------------------AAVALAIHVV-VTYVLVQVLGYGIVGAVASADMAW  237
Sobic.003G126200.2      NATA---------------------AAAALGIHLF-VSWLFVARLKLGLVGVALTLSVSW  248
Sobic.001G476700.1      NAAA---------------------SAVALVIHIF-VSWLFVSRFQFGLAGIALTLNFSW  245
Sobic.001G476700.2      NAAA---------------------SAVALVIHIF-VSWLFVSRFQFGLAGIALTLNFSW  268
```

```
Manes.18G062800.1      IAWV---------------------TLAALVINAF-TSWVFVYVLDFGVIGAAIALDVSW  228
AT5G44050.1            IAIS---------------------SGVSLVVHIF-VCWLFVYVLELGVIGTIATANVSW  230
AT5G10420.1            IAIS---------------------AGVSLAVHIL-VCWFFVYGYKLGIIGTMASVNVPW  228
AT5G65380.1            TAYA---------------------AAVALVVHIL-VCWLFVDGLKLGVVGTVATISISW  227
Manes.12G129000.1      IAWI---------------------SLVALLVHVM-VSWLLVSKLQLGVIGTAMTLNFSW  229
Manes.13G097900.1      IAWV---------------------SFVALLVHVM-VSWLFVYKLQLGVVGTAMTQLF--  227
Sobic.005G020700.1     ITYL---------------------ALLNLGLHLL-LSWLMTVKFQLGVAGVMGSMVIAM  261
Sobic.008G019400.1     ITYL---------------------AVLNLGLHLV-LSWLLAVRLQLGLAGVMGSMVIAM  257
Manes.09G135300.1      IAYL---------------------AAFSLSIHIL-LSWLLTVKYKFGIPGAMASTILAY  220
AT1G33080.1            IAYV---------------------AAVSLGVHVF-LSWLLVVHFDFGIAGAMTSSLVAH  226
AT1G33090.1            IAYV---------------------SAVTLGLHVF-FSWLLVVHFNFGITGAMTSTLVAF  226
AT1G33100.1            ISYV---------------------TAVSLGLHVF-FSWLLVAHFNFGITGAMTSMLIAF  223
AT1G33110.1            IAYV---------------------AAVSLGVHVF-LSWLLMVHFNFGITGAMTSTLVAF  226
AT3G03620.1            VGVL---------------------STLSLALDLV-VTWWCVSVMGMGIGGALLGLNVGS  227
AT5G17700.1            IGIL---------------------STLALVLDIA-ATWWCVSVMGMGIHGALLGLNISS  224
Manes.09G135200.1      IGWL---------------------SAISFVIHVL-LSWVFVTKLDWGVSGAMAALNISA  218
Manes.09G134800.1      VGCL---------------------SATSFVLHLL-LSWIFLVKLNLGIPGAMGAMIISN  215
Manes.08G150400.1      VGWL---------------------SAASFVVHVI-LSWIFVSKLNWGIPGAMTAMNISS  213
Manes.09G134900.1      VGWV---------------------SAASFVLHVL-LSWLFVSKLNWGIPGAMSAMSISS  214
Manes.09G135000.1      IGWV---------------------SAASFVLHVL-LSWLFVSKLNWGIPGAMSAMSISS  214
Manes.09G134900.2      VGWV---------------------SAASFVLHVL-LSWLFVSKLNWGIPGAMSAMSISS  185
Manes.09G135000.2      IGWV---------------------SAASFVLHVL-LSWLFVSKLNWGIPGAMSAMSISS  185


Sobic.002G099300.1     ------------------------------------------------------------  79
AT4G39030.1            IVSAYMMMDSLNKE-GYN---------AYSFA----------------------------  312
Manes.04G084700.1      VIAAYMMTEALNKK-GYN---------AFAIS----------------------------  332
Manes.11G091900.1      VVAAYMMIDSLNKK-GYN---------AYAIS----------------------------  330
Sobic.004G019800.2     VVAAFMMMQNLSNK-GFR---------AFSFT----------------------------  194
Sobic.004G019800.3     VVAAFMMMQNLSNK-GFR---------AFSFT----------------------------  194
Sobic.004G019800.4     VVAAFMMMQNLSNK-GFR---------AFSFT----------------------------  194
AT2G21340.1            VVAAYMMMDALNKK-GYS---------AFSFC----------------------------  329
Manes.11G092000.1      VVAAYMMIEALNKK-GYN---------AFAFS----------------------------  327
Sobic.001G476700.3     ------------------------------------------------------------  164
Sobic.009G077000.1     ------------------------------------------------------------  80
Sobic.001G454900.1     YLITLILLCRLVQQ-VDV-------------I----------------------------  346
Sobic.003G403000.1     YLITLIMLSKLVRK-VDV-------------V----------------------------  381
Sobic.003G403000.2     YLITLIMLSKLVRK-VDV-------------V----------------------------  381
Sobic.003G403000.3     YLITLIMLSKLVRK-VDV-------------V----------------------------  381
Sobic.007G020600.1     YFIASILLWRLRLH-VDL-------------L----------------------------  306
Sobic.007G020600.2     YFIASILLWRLRLH-VDL-------------L----------------------------  306
AT3G08040.1            YFMTLILFVFLAKK-VNL-------------I----------------------------  300
Manes.09G027700.1      ------------------------------------------------------------  156
Manes.09G027800.1      ------------------------------------------------------------  2
Manes.09G027900.1      ------------------------------------------------------------  2
Manes.09G026900.1      YLISLILLWKLIEH-VDL-------------L----------------------------  289
Manes.09G026900.2      YLISLILLWKLIEH-VDL-------------L----------------------------  289
Manes.09G027000.1      ------------------------------------------------------------  0
Manes.07G006000.1      YLISLILLWSLMKK-VDL-------------L----------------------------  324
Manes.07G006000.2      YLISLILLWSLMKK-VDL-------------L----------------------------  324
Manes.10G143000.1      YLISLILLWRLMKK-VDL-------------L----------------------------  317
AT1G51340.2            YLMCGILLWKLMGQ-VDI-------------F----------------------------  296
Manes.06G164500.1      YLISIILLRRLMEQ-VDL-------------L----------------------------  289
Manes.06G164500.2      YLISIILLRRLMEQ-VDL-------------L----------------------------  289
Manes.06G164500.3      YLISIILLRRLMEQ-VDL-------------L----------------------------  289
Manes.06G164500.4      YLISIILLRRLMEQ-VDL-------------L----------------------------  289
Manes.06G164500.5      YLISIILLRRLMEQ-VDL-------------L----------------------------  289
Manes.14G002600.1      YLISIILLWRLMEQ-VDL-------------L----------------------------  273
Sobic.008G006100.1     YLTAFILLWKLNNE-VDL-------------F----------------------------  330
Sobic.005G005400.1     YVIACILLWKLNSK-VVL-------------F----------------------------  314
Sobic.005G005400.3     YVIACILLWKLNSK-VVL-------------F----------------------------  314
Sobic.005G005400.2     YVIACILLWKLNSK-VVL-------------F----------------------------  314
AT2G38330.1            YLIAFILLWKLNEN-VVL-------------L----------------------------  296
Manes.08G096600.1      YFIAFILLWKLNSE-VTL-------------I----------------------------  322
Manes.08G096600.2      YFIAFILLWKLNSE-VTL-------------I----------------------------  322
AT4G38380.1            YTVAILMLILLNKR-VIL-------------L----------------------------  335
Sobic.002G286800.1     YVGMFLLLWSLSKR-AVL-------------L----------------------------  352
Sobic.003G149300.2     YIITVLLLRSLSKK-AVL-------------L----------------------------  322
Sobic.003G149300.1     YIITVLLLRSLSKK-AVL-------------L----------------------------  322
Sobic.003G149300.3     YIITVLLLRSLSKK-AVL-------------L----------------------------  322
Sobic.003G149300.4     YIITVLLLRSLSKK-AVL-------------L----------------------------  198
```

```
Manes.01G153400.1       YLVCFLMIWNLNKR-TIL-------------S----------------------------  72
Manes.04G064900.1       YIVAFLMIWNLNKR-VVL-------------L----------------------------  363
Manes.04G064900.2       YIVAFLMIWNLNKR-VVL-------------L----------------------------  363
Sobic.001G003700.1      LALALMLAAYVLAHELRRG-TT-TKPPTPTP-----------------------------  237
AT4G22790.1             FIVVILLTGYVIVVERMKE-NK--WKQGGWLN----------------------------  247
Manes.02G032300.1       AIVVILLASCVLLKENGKG-GN--WKEGGWLD----------------------------  259
Manes.10G000400.1       INLSLGLLAYLILSRTAMK-P---WKNKQLVDMHNEYTSNVNYNETYMGSVVINTRNKIV  252
AT2G38510.1             MNIDVGLLVYTCFSDSLIK-P---WEGL--------------------------------  220
Manes.08G172300.1       INMNIGLLIYVAVSKKPLK-P---WHGI--------------------------------  250
Manes.09G117000.1       INMNIGLLIYVAVSKKPLK-P---WHGI--------------------------------  220
Sobic.010G167800.1      LVLLVVLLACVLGRRDSAL-RA--AG-PPTAE----------------------------  278
AT5G19700.1             LLVVIFLVAHVWIA-GLHQ-PT--WT-RPSSE----------------------------  253
AT4G29140.1             IFVVAFLVCYVWAS-GLHA-PT--WT-DPTRD----------------------------  273
Manes.03G026500.1       FFVLLSLVSYVWFF-GLHE-PT--WT-RPSRE----------------------------  280
Manes.16G109300.1       FFVLLSLVSYVWIF-GLHE-PT--WT-RPSRE----------------------------  274
Sobic.007G181100.1      LNFLLFLVAYVYLT-GLMR-HGDDGGNGKADA---------------------------L  266
Manes.13G127800.1       LNLLAALLIYLCFS-GICR-ES--WQ-GWSLQ----------------------------  272
Sobic.001G446800.1      LNLLLALLCFLAIS-GAHR-DS--WV-GPTLD----------------------------  270
AT5G52050.1             FNLVAFLFLYICFF-EDKL-SVNEDEKITEET----------------------------  255
AT1G58340.1             LNLVVLLSSFVYFT-SVHS-DT--WV-PITID----------------------------  275
Manes.05G186700.1       LNVFLLLFSFVYFS-GVYK-DS--WV-SPSMD----------------------------  266
Manes.18G054300.1       LNVFIFLFSFVYFS-GVYK-DS--WV-SPSMD----------------------------  266
Sobic.001G320900.1      LNLVLLLLAYIYFS-GVHR-AT--GGFTLSEK----------------------------  295
Sobic.001G320900.2      LNLVLLLLAYIYFS-GVHR-AT--GGFTLSEK----------------------------  295
Sobic.004G283500.1      LNLLLFLLAYILFK-GVHK-RT--GGLALSAE----------------------------  296
Sobic.006G184400.1      LNLLLFLLGYIFFM-GVHR-RT--GGFVLSRE----------------------------  309
AT4G23030.1             VNLLGFLIIYIVFS-GVYQ-KT--WG-GFSMD----------------------------  250
Manes.01G067000.1       FNLVSSLIIYVILS-GAHR-KT--WG-GISLE----------------------------  270
Manes.02G027800.1       FNLVTSLIIYVMVS-GVHK-KT--WG-GISLE----------------------------  274
Manes.06G143600.1       LNLIVSLIIYILIS-GVHK-KT--WG-GFSTE----------------------------  282
Manes.14G029100.1       FNFVGSLIIYIFIS-GVQE-KT--WG-GFSRE----------------------------  286
AT5G49130.1             FISLSLLLCYIYLENNNND-KT--TSKSLCLD----------------T--------PLM  249
Manes.03G198000.1       FNTLLYLLCYMYFTRVPEE-PL--YTPLSPLP------------QLP-P--------S--  243
Manes.15G011000.1       FNTLLFLLCYMYFTRVPEE-PL--YTPLSPPP------------QPS-Q--------PSQ  245
Sobic.001G019700.1      TNMLLFLLAYIRWARAC-E-GT--WKGWA-------------------------------  258
AT1G71870.1             LIMVVLLVGYVWVS-GMLQ-KR--VSGDGDGG------------STTMV--------AVV  250
Manes.02G189800.1       MNMVALMVGYVWWVSGRWE-MK--WSG---------------------------------  236
Manes.18G098800.1       MNMVVLMVGYMWWVRGWWE-MK--WTG---------------------------------  236
Sobic.010G256932.1      WFNVALLVVYVKVS-EAGR-RS--WH-GWSRE----------------------------  216
Sobic.010G256700.1      WFNVALLVVYVKVS-EAGR-RS--WH-GWSRE----------------------------  247
Sobic.010G256700.5      WFNVALLVVYVKVS-EAGR-RS--WH-GWSRE----------------------------  247
Sobic.010G256700.6      WFNVALLVVYVKVS-EAGR-RS--WH-GWSRE----------------------------  247
Sobic.010G256700.4      WFNVALLVVYVKVS-EAGR-RS--WH-GWSRE----------------------------  247
Sobic.009G106900.1      GVNLTILALYVRLS-GACN-AT--WT-GFSTD----------------------------  65
Manes.S031500.1         WVNAISLMLYVKKS-PTCK-ET--WA-GFSKE----------------------------  251
Manes.14G109900.1       WINVFLLALYIKFS-PACM-KT--WT-GFSRE----------------------------  261
Manes.14G109900.2       WINVFLLALYIKFS-PACM-KT--WT-GFSRE----------------------------  160
Manes.14G109900.3       WINVFLLALYIKFS-PACM-KT--WT-GFSRE----------------------------  160
AT1G73700.1             WFNVILLSCYVKFS-PSCS-HS--WT-GFSKE----------------------------  241
AT2G34360.1             WLNVILLSCYVKFS-PSCS-LT--WT-GFSKE----------------------------  244
AT5G52450.1             WLNVVLLFCYVKFS-PSCS-LT--WT-GFSKE----------------------------  243
Sobic.009G106960.1      ------------------------------------------------------------  0
Manes.14G060800.1       CINVFILAIYIKFS-PTCK-HT--WT-GFSRA----------------------------  254
Sobic.009G106800.1      SFNLAMLALYVRFS-SACK-RT--WT-GFSTE----------------------------  240
Sobic.007G074300.2      GLNLAILALYVRLS-SACG-RT--WN-GFSVE----------------------------  64
Sobic.009G106700.1      AINLVILALYVRLS-DACK-ET--WN-GFSWE----------------------------  242
Sobic.010G138400.4      WVNVAILAVYVRVS-SACK-ET--WT-GFSTE----------------------------  167
Sobic.010G138400.1      WVNVAILAVYVRVS-SACK-ET--WT-GFSTE----------------------------  245
Sobic.010G138400.5      WVNVAILAVYVRVS-SACK-ET--WT-GFSTE----------------------------  164
Sobic.004G129900.1      LVNLSALALYIRLS-PSCK-AT--WL-GFSRQ----------------------------  242
Sobic.004G129900.2      LVNLSALALYIRLS-PSCK-AT--WL-GFSRQ----------------------------  160
Sobic.006G042200.1      LTNVSILAIYVRLA-PACR-NT--WR-GFSKE----------------------------  245
Sobic.004G129700.1      LANLAFLALYVRLS-PSCK-YS--WT-GFSAE----------------------------  254
Sobic.004G129800.1      LANLSFLAVYVRAS-PACK-ST--WT-CFSAE----------------------------  245
Sobic.004G129800.2      LANLSFLAVYVRAS-PACK-ST--WT-CFSAE----------------------------  160
AT3G23550.1             WIAFVSLGFYVICS-DKFK-ET--WT-GFSME----------------------------  248
AT3G23560.1             WIAFLSLGTYVMCS-EKFK-ET--WT-GFSLE----------------------------  256
Manes.15G088100.1       WLSVLMVAMYVIIA-KKFE-HT--WH-GFSFE----------------------------  268
Manes.03G109100.1       WISFFVLAMYVLFA-KKIE-HT--WC-GFSFE----------------------------  252
Manes.03G109100.2       WISFFVLAMYVLFA-KKIE-HT--WC-GFSFE----------------------------  213
Sobic.002G311200.1      WVSCLLLRAYVLRS-GAFS-ET--WE-GFSAE----------------------------  363
```

```
Sobic.002G006500.3      WLSFLMLAAYVMAS-DRFR-ET--WP-GLTTE---------------------------  277
Sobic.002G006500.4      WLSFLMLAAYVMAS-DRFR-ET--WP-GLTTE---------------------------  277
AT2G04090.1             WFNVLILALYVRFS-SACE-KT--RG-FVSDD---------------------------  247
AT2G04100.1             WFNVLILALYVRFS-SSCE-KT--RG-FVSDD---------------------------  247
AT2G04066.1             -----------------------------------------------------------  0
AT2G04040.1             WFYAVILSCYVRFS-SSCE-KT--RG-FVSRD---------------------------  244
AT2G04080.1             WFYAVILSCHVRFF-SSCE-KT--RG-FVSND---------------------------  244
AT2G04050.1             WFYVVILSCYVRYS-SSCD-KT--RV-FVSSD---------------------------  244
AT2G04070.1             WFFAMTLSCYVRFS-SSCE-KT--RR-FVSQD---------------------------  244
Sobic.001G481800.1      WVEVTVLGLYIKFS-PSCE-KT--RA-PFTWE---------------------------  271
Sobic.003G260200.1      WLNVAMLVGYIVFS-SSCK-ET--RA-RPTIE---------------------------  254
Sobic.009G224800.1      WLNVIMLAAYIRYS-NSCK-ET--RS-PPTVE---------------------------  257
Sobic.009G224800.2      WLNVIMLAAYIRYS-NSCK-ET--RS-PPTVE---------------------------  257
AT1G66760.2             WLNAVFLWVYMKRS-SRCV-ET--RI-YMSKD---------------------------  245
AT1G64820.1             WLNVFLLWIFMRYS-ALHR-EM--KN-LGLQE---------------------------  246
AT1G66780.1             WLNVGLLWAFMRDS-ALYR-KN--WN-LRAQE---------------------------  252
Manes.02G072400.1       WLNVFLLGFYVKCS-SACQ-KT--RT-PISKD---------------------------  242
Manes.01G113500.1       WLNVIFLVSYFLFS-SSCE-KT--RI-LCWRD---------------------------  230
Manes.01G113500.2       WLNVIFLVSYFLFS-SSCE-KT--RI-LCWRD---------------------------  230
AT1G15150.1             WLYAIFLGSFMYYS-SACS-ET--RA-PLTME---------------------------  248
AT1G15160.1             WLYAIFLGSFMYFS-SACS-ET--RA-PLTME---------------------------  248
AT1G15170.1             WLYAIFLGSFMYYS-SACS-ET--RA-PLSME---------------------------  251
AT1G15180.1             CLYTIILGSLMCFS-SACS-ET--RA-PLSME---------------------------  252
AT1G71140.1             WLNVTVLGLYMTFS-SSCS-KS--RA-TISMS---------------------------  243
Manes.06G063000.1       WVNWILLALYMRFS-SSCE-KT--RV-SVSME---------------------------  243
Manes.14G109700.1       WLNVIFLASYMTFS-PACS-KT--RV-PISME---------------------------  252
Manes.14G109800.1       WLNVIILALYMKFA-SACA-KT--RA-PISME---------------------------  253
Sobic.008G171600.1      WVLVVLTWGYIVWS-PACK-ET--WT-GLSLL---------------------------  260
Sobic.008G171600.2      WVLVVLTWGYIVWS-PACK-ET--WT-GLSLL---------------------------  274
Sobic.008G171600.3      WVLVVLTWGYIVWS-PACK-ET--WT-GLSLL---------------------------  260
AT3G59030.1             WLLVAVNGMYILMS-PNCK-ET--WT-GFSTR---------------------------  268
Manes.01G182000.1       WLLVILNGLYIVLS-PKCK-ET--WT-GLSIS---------------------------  267
Manes.02G142000.1       WLLVILNGLYIVLS-PKCK-ET--WT-GLSIN---------------------------  267
AT4G21903.2             WFIVGAQTFYVITS-VRFK-DT--WT-GFSWK---------------------------  269
AT4G21910.4             WVIVGSQCFYIAVS-PKFR-HT--WT-GLSWR---------------------------  273
AT1G11670.1             WIIVLAQIIYIKIS-PRCR-RT--WD-GFSWK---------------------------  267
AT1G61890.1             WIIVVAQIVYIKMS-PRCR-RT--WE-GFSWK---------------------------  264
Manes.16G008000.1       WIIVVAQFVYIVKS-DRCK-ET--WT-GFSLQ---------------------------  267
Manes.17G038200.1       WVIVAAQFIYIVKT-SRCK-QT--WN-GFTLQ---------------------------  268
Sobic.002G318300.1      WVLVAGQFAYIVWS-PRCR-AT--WT-GFTWA---------------------------  257
Manes.17G038300.1       WFMVLAQFVYIVTS-ERCK-QS--WT-GFSWE---------------------------  254
Manes.16G007900.1       WLIVIAQFLYIVMS-RKCR-KT--WA-GFSVQ---------------------------  263
Manes.17G038400.1       WFIVIAQFLYIVKS-KKCR-KT--WA-GFSVQ---------------------------  263
Sobic.001G012600.1      WIIVLAQFGYIVTS-PRCR-ET--WT-GFTSQ---------------------------  260
Sobic.001G012600.2      WIIVLAQFGYIVTS-PRCR-ET--WT-GFTSQ---------------------------  260
Sobic.001G185600.1      WVIVVAQFVYIVRS-QRCRRRT--WT-GFSCR---------------------------  286
Sobic.001G185400.1      WLIVAAQFAYIVVS-PKCR-HT--WT-GFTFQ---------------------------  259
AT3G21690.1             WIIVVAQFVYIVTS-ERCR-ET--WR-GFSVQ---------------------------  269
Sobic.001G185800.1      WIIVAAQFLYIVTS-ERCR-RT--WT-GLSCR---------------------------  263
Sobic.001G185500.1      WVIVVAQFVYIVTS-ERCR-LT--WT-GFSWE---------------------------  277
Sobic.001G185500.2      WVIVVAQFVYIVTS-ERCR-LT--WT-GFSWE---------------------------  160
Sobic.007G165500.2      WVIALGQMAYII-G-WCK--DG--WR-GWSAA---------------------------  292
AT4G00350.1             WGIAIAQVVYVV-G-WCK--DG--WK-GLSWL---------------------------  303
Manes.01G255000.1       WGISLAQVAYVV-G-WCK--DG--WR-GLSCN---------------------------  365
Sobic.004G349550.1      WVVALAQVAYVT-R-RCRG-RG--WD-GLSWD---------------------------  159
Sobic.004G349600.1      WLTSLAQVAYVV-G-WCP--DG--WT-GLSRA---------------------------  262
Sobic.007G176000.1      WLVVLGQLAYIL-M-GYCP-GA--WN-GFDCL---------------------------  267
Sobic.007G176100.1      WLVVLGQLAYIL-M-GYCP-GA--WN-GFDCL---------------------------  272
AT1G47530.1             WLIVIGQLLYIL-I-TKSD-GA--WT-GFSML---------------------------  251
Manes.05G164500.1       WIIVIAQLLYIF-I-TKSD-GA--WS-GFSWL---------------------------  247
Manes.05G164500.2       WIIVIAQLLYIF-I-TKSD-GA--WS-GFSWL---------------------------  247
Manes.18G030900.1       WLIVIGQLLYIF-I-TKSD-GA--WS-GFSWL---------------------------  248
AT1G23300.1             WLIDVTQIVYIC-G-GSSG-RA--WS-GLSWM---------------------------  260
AT3G26590.1             CFIDMAQLVYIF-S-GTCG-EA--WS-GFSWE---------------------------  261
AT5G38030.1             WFIVVAQLVYIF-S-GTCG-EA--WS-GFSWE---------------------------  261
Manes.12G023400.1       WLIDLSQFLYIV-S-GACR-QA--WN-GFSCQ---------------------------  264
Manes.12G023500.1       WLIDLSQFLYIV-S-GACR-QS--WN-GFSSQ---------------------------  264
Manes.12G023600.1       WFIDLAQFFYII-S-GTCG-RT--WN-GFSWK---------------------------  281
Manes.13G025200.1       WFIDLAQFLYII-S-GSCG-RA--WN-GFSWK---------------------------  281
Manes.13G025200.2       WFIDLAQFLYII-S-GSCG-RA--WN-GFSWK---------------------------  280
Sobic.001G273100.1      WLVVAAQFAYMTTG-ERFP-DA--WN-GFTVR---------------------------  259
```

```
Sobic.001G273100.2      WLVVAAQFAYMTTG-ERFP-DA--WN-GFTVR----------------------------  259
Sobic.001G273000.2      WLIVAAQFAYMI-G-GRFP-DT--WK-GFTMC----------------------------  231
Sobic.001G273000.1      WLIVAAQFAYMI-G-GRFP-DT--WK-GFTMC----------------------------  242
Sobic.001G273000.3      WLIVAAQFAYMI-G-GRFP-DT--WK-GFTMC----------------------------  242
Manes.15G147800.1       WLVVLGQMVYLV-C-GCFP-EA--RT-GFSWL----------------------------  246
Manes.15G147900.1       WLVVLGQIVYVV-C-GYFP-EA--WT-GFSWS----------------------------  247
Manes.17G098800.1       WLVVLGQIVYVF-C-GCFP-EA--WT-GFSWS----------------------------  252
Manes.17G098900.1       WLVVLGQIIYVV-C-GCFP-EA--WT-GFSWS----------------------------  252
Manes.17G098900.2       WLVVLGQIIYVV-C-GCFP-EA--WT-GFSWS----------------------------  159
Sobic.001G162400.1      WLVIAAQVGYLV-S-GCFP-EA--WQ-GFSML----------------------------  237
Sobic.003G307600.2      WLVNVAQFVYLV-G-GSFP-GA--WT-GFSRK----------------------------  159
Sobic.002G232200.1      AVLAALLLSYAL-G-GGCP-ET--WS-GFSTS----------------------------  255
Sobic.002G232500.1      ALATVGLLSYAL-G-GGCP-ET--WR-GFSAS----------------------------  251
Sobic.002G232600.1      ALITGLQLAYAV-G-GGCP-ET--WR-GFSSS----------------------------  246
Sobic.007G160700.1      WLVVLGQYVYVV-G-GGCP-LS--WK-GFTME----------------------------  263
Sobic.003G126200.2      WTITAMLFVYVT-C-GGCP-ET--WH-GFTAE----------------------------  274
Sobic.001G476700.1      WATGAMLFAYVS-C-GGCP-DT--WH-GFSLE----------------------------  271
Sobic.001G476700.2      WATGAMLFAYVS-C-GGCP-DT--WH-GFSLE----------------------------  294
Manes.18G062800.1       WFMVFALLGYVL---YWCP-LT--WT-GFSVQ----------------------------  253
AT5G44050.1             WLNVFILFTYTT-C-GGCP-LT--WT-GFSME----------------------------  256
AT5G10420.1             WLNIFILFLYST-R-GGCT-LT--WT-GFSSE----------------------------  254
AT5G65380.1             WVNVLILLVYST-C-GGCP-LT--WT-GLSSE----------------------------  253
Manes.12G129000.1       WVLVFGHLGYTV-C-GGCP-LT--WN-GFSIE----------------------------  255
Manes.13G097900.1       ------------------------------LV----------------------------  229
Sobic.005G020700.1      WIPVFGQLAFVF-F-GGCP-HT--WT-GFSSA----------------------------  287
Sobic.008G019400.1      WIPVFGQLAFVF-F-GGCP-LT--WT-GFSSA----------------------------  283
Manes.09G135300.1       WIPNIGQLAFVT-C-GGCP-ET--WK-GFSFL----------------------------  246
AT1G33080.1             WLPNIAQVLFVT-C-GGCT-ET--WR-GFSWL----------------------------  252
AT1G33090.1             WMPNIVQLLYVT-S-GGCK-DT--WR-GFTML----------------------------  252
AT1G33100.1             WLPIIVQLLYVT-C-GGCK-DT--WR-GFSML----------------------------  249
AT1G33110.1             WLPNIAQLLFVT-C-GGCK-DT--WR-GFSMM----------------------------  252
AT3G03620.1             WAMVLAEFVYIF-G-GWCP-FT--WT-GFSIA----------------------------  253
AT5G17700.1             WSVAIAEFVYVF-G-GWCP-HT--WT-GFSTA----------------------------  250
Manes.09G135200.1       WLTVVELFLYVL-G-GWCP-NT--WK-GFTRA----------------------------  244
Manes.09G134800.1       WLVVIGDLVYIF-G-GWCP-NT--WK-GFTLA----------------------------  241
Manes.08G150400.1       WLVIIGELVYVF-G-GWCP-DT--WR-GFTSA----------------------------  239
Manes.09G134900.1       WLIVIGLLVYVF-G-GWCP-DT--WR-GFTLA----------------------------  240
Manes.09G135000.1       WLIVIGQLVYVF-G-GWCP-ET--WK-GFTLA----------------------------  240
Manes.09G134900.2       WLIVIGLLVYVF-G-GWCP-DT--WR-GFTLA----------------------------  211
Manes.09G135000.2       WLIVIGQLVYVF-G-GWCP-ET--WK-GFTLA----------------------------  211


Sobic.002G099300.1      ------------------------------------------------------------  79
AT4G39030.1             -I--PSP---QELWKISALAAPVFISIFS---KIAF------YSFIIYCAT--SMGTHVL  355
Manes.04G084700.1       -I--PST---DEFSQIFSIAAPVFVTMFS---KVAF------YSLMTYFAT--AKGTFTV  375
Manes.11G091900.1       -I--PSP---SDLMTIFGLAAPVFVMMMS---KVAF------YSLLVYFAT--SMGTLSL  373
Sobic.004G019800.2      -I--PSV---RELLQIFEIAAPVFVTMTS---KVAF------YALLTYSAT--SMGAITL  237
Sobic.004G019800.3      -I--PSV---RELLQIFEIAAPVFVTMTS---KVAF------YALLTYSAT--SMGAITL  237
Sobic.004G019800.4      -I--PSV---RELLQIFEIAAPVFVTMTS---KVAF------YALLTYSAT--SMGAITL  237
AT2G21340.1             -V--PSP---SELLTIFGLAAPVFITMMS---KVLF------YTLLVYFAT--SMGTNII  372
Manes.11G092000.1       -V--PTL---DEILTIVRLAAPVFVTMMS---KVVF------YSLLIYFAT--SMGTYSV  370
Sobic.001G476700.3      ------------------------------------------------------------  164
Sobic.009G077000.1      ------------------------------------------------------------  80
Sobic.001G454900.1      -P--PSI----KSLKFGRFLGCGFLLLARVVAVTFC------VTLAASLAA--RHGPTIM  391
Sobic.003G403000.1      -P--PSL----KCLKFRRFLGCGFLLLARVVAVTFC------VTLAASLAA--RHGPTAM  426
Sobic.003G403000.2      -P--PSL----KCLKFRRFLGCGFLLLARVVAVTFC------VTLAASLAA--RHGPTAM  426
Sobic.003G403000.3      -P--PSL----KCLKFRRFLGCGFLLLARVVAVTFC------VTLAASLAA--RHGPTAM  426
Sobic.007G020600.1      -P--PSF----KHLQFGRFLKNGFLLLARVIAATFC------VTLSASMAA--RQGSTPM  351
Sobic.007G020600.2      -P--PSF----KHLQFGRFLKNGFLLLARVIAATFC------VTLSASMAA--RQGSTPM  351
AT3G08040.1             -P--PNF----GDLQFGRFLKNGLLLLARTIAVTFC------QTLAAAMAA--RLGTTPM  345
Manes.09G027700.1       ------------------------------------------------------------  156
Manes.09G027800.1       ------------------------------------------------------------  2
Manes.09G027900.1       ------------------------------------------------------------  2
Manes.09G026900.1       -P--PNI----KDLQFSRFLKNGFMLLMRVIAATFC------VTLAASLAA--RYGSTSM  334
Manes.09G026900.2       -P--PNI----KDLQFSRFLKNGFMLLMRVIAATFC------VTLAASLAA--RYGSTSM  334
Manes.09G027000.1       -----------------------------------------------------------M  1
Manes.07G006000.1       -P--PSA----KDLQFGRFLKNGFLLLARVIAATIC------VTLAASRAA--RLGSTPM  369
Manes.07G006000.2       -P--PSA----KDLQFGRFLKNGFLLLARVIAATIC------VTLAASRAA--RLGSTPM  369
Manes.10G143000.1       -P--PSL----KDLQFGRFLKNGFLLLARVIAATIC------VTLAASRAA--RLGSTPM  362
AT1G51340.2             -N--MST----KHLQFCRFMKNGFLLLMRVIAVTFC------VTLSASLAA--REGSTSM  341
Manes.06G164500.1       -P--PSA----KNLQFSKFLKNGFLLLMRVVAVTFC------VTLSASLAA--RQGSTSM  334
```

```
Manes.06G164500.2       -P--PSA----KNLQFSKFLKNGFLLLMRVVAVTFC------VTLSASLAA--RQGSTSM  334
Manes.06G164500.3       -P--PSA----KNLQFSKFLKNGFLLLMRVVAVTFC------VTLSASLAA--RQGSTSM  334
Manes.06G164500.4       -P--PSA----KNLQFSKFLKNGFLLLMRVVAVTFC------VTLSASLAA--RQGSTSM  334
Manes.06G164500.5       -P--PSA----KNLQFSKFLKNGFLLLMRVVAVTFC------VTLSASLAA--RQGSTSM  334
Manes.14G002600.1       -P--PSF----RHLQFGKFLKNGLLLLMRVIAVTFC------VTLSASLAA--RQGATSM  318
Sobic.008G006100.1      -S--WNI---IEDGGVIRYLKSGGLLIGRTIAVFLT------LTLSTSLAA--REGPVPM  376
Sobic.005G005400.1      -S--RNV---I-GGGIIRYLKSGGLLIGRTIAVLLT------MTLSTSLAA--RQGPVPM  359
Sobic.005G005400.3      -S--RNV---I-GGGIIRYLKSGGLLIGRTIAVLLT------MTLSTSLAA--RQGPVPM  359
Sobic.005G005400.2      -S--RNV---I-GGGIIRYLKSGGLLIGRTIAVLLT------MTLSTSLAA--RQGPVPM  359
AT2G38330.1             -S--PQI----KVGRANQYLKSGGLLIGRTVALLVP------FTLATSLAA--QNGPTQM  341
Manes.08G096600.1       -S--PNI----DGSRVVRYLNSGGLLIGRTIAVLLT------MTLATSMAA--REGPIPM  367
Manes.08G096600.2       -S--PNI----DGSRVVRYLNSGGLLIGRTIAVLLT------MTLATSMAA--REGPIPM  367
AT4G38380.1             -P--PKI----GSLKFGDYLKSGGFVLGRTLSVLVT------MTVATSMAA--RQGVFAM  380
Sobic.002G286800.1      -P--PKI----KDLEFVGYIKSGGMLLGRTLSVLIT------MTLGTAMAA--RQGTVAM  397
Sobic.003G149300.2      -P--PGI----DQLEFGGYLKSGGMLLGRTLSILLT------MTIGTSMAA--RQGPTAM  367
Sobic.003G149300.1      -P--PGI----DQLEFGGYLKSGGMLLGRTLSILLT------MTIGTSMAA--RQGPTAM  367
Sobic.003G149300.3      -P--PGI----DQLEFGGYLKSGGMLLGRTLSILLT------MTIGTSMAA--RQGPTAM  367
Sobic.003G149300.4      -P--PGI----DQLEFGGYLKSGGMLLGRTLSILLT------MTIGTSMAA--RQGPTAM  243
Manes.01G153400.1       -F--LSI----KGLHFGGYLKSGGFLLGRTLAAVLT------ITLSTSMAA--HQGALAM  117
Manes.04G064900.1       -P--PKM----GSLQFGVYLKSGGFLIGRTLAVLTT------MTLATSMAA--RQGPLAM  408
Manes.04G064900.2       -P--PKM----GSLQFGVYLKSGGFLIGRTLAVLTT------MTLATSMAA--RQGPLAM  408
Sobic.001G003700.1      -------TPASPTSSL--------LRLAVPCCLNTC-LEWWSYEILVLLTGRLPDARRMV  281
AT4G22790.1             -------QSAQDWLTL--------IKLSGPCCLTVC-LEWWCYEILVLLTGRLPNPVQAV  291
Manes.02G032300.1       -------QGIYDWLRL--------LKLSGPCCLTSC-LEWWCWEILVLLTGRLANAKQAV  303
Manes.10G000400.1       LHVTSFF---QGWQPL--------VSLMLPSVLSVC-LEWWWYEIMLLLCGLQDNPQASV  300
AT2G38510.1             -ALRSLF---RGWWPL--------LSLAAPSAISVC-LEYWWYEIMLFLCGLLGNPKASV  267
Manes.08G172300.1       -TATSML---QGWKPL--------LSLSLPSVVSVC-LEWWWYEIMLFLCGLLSNPKASV  297
Manes.09G117000.1       -TASSIF---YGWRPL--------LSLALPSVISVC-LEWWWYEIMLFLCGLLTNPKANV  267
Sobic.010G167800.1      -C----L---AGWGPL--------ARLAAPSCVSVC-LEWWWYEVMILLCGLLPDPKPAV  321
AT5G19700.1             -C----F---KDWGPV--------VTLAIPSCIGVC-LEWWWYEIMTVLCGLLIDPSTPV  296
AT4G29140.1             -C----F---RGWAPL--------LRLAGPSCVSVC-LEWWWYEIMIVLCGLLVNPRSTV  316
Manes.03G026500.1       -C----F---TGWKPL--------IKLAAPSCVSVC-LEWWWYEIMIVLCGLLANPKSAV  323
Manes.16G109300.1       -C----F---TGWKPL--------IRLAAPSCVSVC-LEWWWYEIMIILCGLLANPKSTI  317
Sobic.007G181100.1      MCATPPPAEDVEWGCL--------LRLSLHSCMSVC-LEWWWYEIMVLLCGVLADPKAAV  317
Manes.13G127800.1       -C----F---DEWKPI--------LGLAIPSCISVC-LEWWWYELMIVLSGLLTNASEAV  315
Sobic.001G446800.1      -C----L---RGWPEM--------LRLAVPTATAVC-LEWWWYELMIVLSGLLANPRATV  313
AT5G52050.1             -C----EDSVREWKKL--------LCLAIPSCISVC-LEWWCYEIMILLCGFLLDPKASV  301
AT1G58340.1             -S----L---KGWSAL--------LSLAIPTCVSVC-LEWWWYEFMIILCGLLANPRATV  318
Manes.05G186700.1       -C----L---RGWSSL--------LSLAVPTCVSVC-LEWWWYEFMIMLCGLLANPKATI  309
Manes.18G054300.1       -C----L---RGWSSL--------LSLAVPTCVSVC-LEWWWYEFMIMLCGLLVNPRSTI  309
Sobic.001G320900.1      -L----FKDVTGWMRL--------ARLAVASCASVC-LEWWWYEIMILLCGLLADPKATV  341
Sobic.001G320900.2      -L----FKDVTGWMRL--------ARLAVASCASVC-LEWWWYEIMILLCGLLADPKATV  341
Sobic.004G283500.1      -S----F---RGWGEL--------VSLALPSCVSVC-LEWWWYEIMILLCGLLANPQATV  339
Sobic.006G184400.1      -S----F---RGWGEL--------ASLALPSCVSVC-LEWWWYEIMILLCGLLLNPQATV  352
AT4G23030.1             -C----F---KGWRSL--------MKLAIPSCVSVC-LEWWWYEIMILLCGLLLNPQATV  293
Manes.01G067000.1       -C----L---RGWKSL--------LNLAIPSCISVC-LEWWWYEIMILLCGLLLNPRATV  313
Manes.02G027800.1       -C----L---RGWKSL--------LNLAIPSCISVC-LEWWWYEIMIMLCGLLLNPRATV  317
Manes.06G143600.1       -C----F---KEWKTL--------LNLAIPSCISVC-LEWWWYEIMILLCGLLLNPRATV  325
Manes.14G029100.1       -C----L---KEWKTL--------LNLAIPSCISVC-LEWWWYEIMILLCGLLLNPRATV  329
AT5G49130.1             LYGSRDSGENDVWSTL--------VKFAVPSCIAVC-LEWWWYEFMTVLAGYLPEPKVAL  300
Manes.03G198000.1       -------LGVREWGIL--------IRLAVPSCVAVC-LEWWWYEFMTIAAGYLSKPRVAL  287
Manes.15G011000.1       PPYSTSFSLGREWGIL--------LRLAIPSCLAVC-LEWWWYEFMTILAGYLSKPRVAL  296
Sobic.001G019700.1      ----RPAAVASGLPAL--------ASLAVPSCVGVC-LEWWWYEVVTVLAGYLPNPAAAV  305
AT1G71870.1             AQSSSVMELVGGLGPL--------MRVAVPSCLGIC-LEWWWYEIVIVMGGYLENPKLAV  301
Manes.02G189800.1       ----RIGGVCGGVGPL--------LKLAVPSCLGIC-LEWWWYEIVIVMAGYLPNPTLAV  283
Manes.18G098800.1       ----GIGGVCDGVGPL--------LKLAVPSCLGIC-LEWWWYEIVIVMAGYLPNPTLAV  283
Sobic.010G256932.1      -A----L-KLKDAKVY--------LKLAIPTTFMIC-LEYWAFEMVVLLAGFLPDPKLET  261
Sobic.010G256700.1      -A----L-KLKDAKVY--------LKLAIPSTFMTC-LEYWAFEMVVLLAGFLPDPKLET  292
Sobic.010G256700.5      -A----L-KLKDAKVY--------LKLAIPSTFMTC-LEYWAFEMVVLLAGFLPDPKLET  292
Sobic.010G256700.6      -A----L-KLKDAKVY--------LKLAIPSTFMTC-LEYWAFEMVVLLAGFLPDPKLET  292
Sobic.010G256700.4      -A----L-KLKDAKVY--------LKLAIPSTFMTC-LEYWAFEMVVLLAGFLPDPKLET  292
Sobic.009G106900.1      -A----L-PLSGLREF--------DSQSSPSRRR*-------------------------  85
Manes.S031500.1         -A----L-H--GIPKF--------LMLAIPSAAMLS-LEIWSFEMMVLLSGLLPNPKLET  294
Manes.14G109900.1       -A----L-H--DILSF--------VKLAVPSAIMIC-LEYWSFEMVVLLSGLLPNPKLET  304
Manes.14G109900.2       -A----L-H--DILSF--------VKLAVPSAIMIC-LEYWSFEMVVLLSGLLPNPKLET  203
Manes.14G109900.3       -A----L-H--DILSF--------VKLAVPSAIMIC-LEYWSFEMVVLLSGLLPNPKLET  203
AT1G73700.1             -A----F-Q--ELYDF--------SKIAFPSAVMVC-LELWSFELLVLASGLLPNPVLET  284
AT2G34360.1             -A----R-R--DIIPF--------MKLVIPSAFMVCSLEMWSFELLVLSSGLLPNPVLET  288
AT5G52450.1             -A----L-R--DILPF--------LRLAVPSALMVC-LEMWSFELLVLLSGLLPNPVLET  286
Sobic.009G106960.1      ------------------------------------------------------------  0
Manes.14G060800.1       -G----T-K--DLLSF--------LKLGIPSALMVC-LEFWSYEFLVIISGLLPNPKLQL  297
```

| | | |
|---|---|---|
| Sobic.009G106800.1 | -A----F-K--DLHRF--------TELAIPSAMMVC-LEWWSFELLVLLSGLLPNPKLET | 283 |
| Sobic.007G074300.2 | -G----F-K--ELRQF--------ANLAVPSAFMIC-VEFWAFEIIVLLSGLLPNPQLET | 107 |
| Sobic.009G106700.1 | -A----F-K--DLWRF--------TELAWPSAIMIC-LEWWSFEVLVLLSGLLPNPQLET | 285 |
| Sobic.010G138400.4 | -A----F-H--DALSF--------FRLAIPSALMVC-LEMWSFELIVLLSGLLPNPQLET | 210 |
| Sobic.010G138400.1 | -A----F-H--DALSF--------FRLAIPSALMVC-LEMWSFELIVLLSGLLPNPQLET | 288 |
| Sobic.010G138400.5 | -A----F-H--DALSF--------FRLAIPSALMVC-LEMWSFELIVLLSGLLPNPQLET | 207 |
| Sobic.004G129900.1 | -A----F-H--GILGF--------LKLAMPSAAMVC-MEWWSFEFLVLLSGLLPNPKLET | 285 |
| Sobic.004G129900.2 | -A----F-H--GILGF--------LKLAMPSAAMVC-MEWWSFEFLVLLSGLLPNPKLET | 203 |
| Sobic.006G042200.1 | -A----F-H--DIPSF--------LRLGVPSALMVC-LEWWSFELLVLLSGLLPNPKLET | 288 |
| Sobic.004G129700.1 | -A----F-R--GVPDF--------LKLAVPSAVMVC-MKWWSFEFLVMFSGLLPNPKLET | 297 |
| Sobic.004G129800.1 | -A----F-R--GVPDF--------LKLAVPSAVMVC-MEWWSFELLVLLSGLLPNPKLET | 288 |
| Sobic.004G129800.2 | -A----F-R--GVPDF--------LKLAVPSAVMVC-MEWWSFELLVLLSGLLPNPKLET | 203 |
| AT3G23550.1 | -S----F-H--HVVLN--------LTLSIPSAAMVC-LEYWAFEILVFLAGLMRNPEITT | 291 |
| AT3G23560.1 | -S----F-R--YIVIN--------LTLSLPSAAMVC-LEYWAFEILVFLAGVMPNPEINT | 299 |
| Manes.15G088100.1 | -S----F-N--YILTN--------IKLALPSAAMVC-LEYWAFEILVFLAGTMPNSKITT | 311 |
| Manes.03G109100.1 | -S----F-H--YLHIT--------LKLALPSAAMVC-LDDWATEILVFLAGLMPDSQIST | 295 |
| Manes.03G109100.2 | -S----F-H--YLHIT--------LKLALPSAAMVC-LDDWATEILVFLAGLMPDSQIST | 256 |
| Sobic.002G311200.1 | -A----F-K--YVAPT--------VKLATPSAVMVC-LEYWAFELLVLIAGLLPNSTVST | 406 |
| Sobic.002G006500.3 | -A----F-R--HVLPG--------MKLAIPSAVMVC-FEYWSFEFLVLFAGLMPESQVST | 320 |
| Sobic.002G006500.4 | -A----F-R--HVLPG--------MKLAIPSAVMVC-FEYWSFEFLVLFAGLMPESQVST | 320 |
| AT2G04090.1 | -F----V-L--SVKQF--------FQYGIPSAAMTT-IEWSLFELLILSSGLLPNPKLET | 290 |
| AT2G04100.1 | -F----V-L--SVKQF--------FQYGIPSAAMTT-IEWSLFEFLILSSGLLPNPKLET | 290 |
| AT2G04066.1 | ---------------------------------------------------------MEA | 3 |
| AT2G04040.1 | -F----V-S--SIKQF--------FQYGIPSAAMIC-LEWWLFEILILCSGLLPNPKLET | 287 |
| AT2G04080.1 | -F----M-S--SIKQY--------FQYGVPSAGLIC-LEWWLFELLILCSGLLPNPKLET | 287 |
| AT2G04050.1 | -F----V-S--CIKQF--------FHFGVPSAAMVC-LEWWLFELLILCSGLLPNPKLET | 287 |
| AT2G04070.1 | -F----L-S--SVKQF--------FRYGVPSAAMLC-LEWWLFELLILCSGLLQNPKLET | 287 |
| Sobic.001G481800.1 | -A----F-R--GIGNF--------MRLAVPSALMIC-LEWWSYELLVLLCGVLPNAALET | 314 |
| Sobic.003G260200.1 | -V----F-R--GVDAF--------LRLALPSALMMC-FEWWSFELLTLMSGLLPNPELQT | 297 |
| Sobic.009G224800.1 | -A----F-K--GVGVF--------LRLALPSALMLC-FEWWSFEILILVSGILPNPELQT | 300 |
| Sobic.009G224800.2 | -A----F-K--GVGVF--------LRLALPSALMLC-FEWWSFEILILVSGILPNPELQT | 300 |
| AT1G66760.2 | -V----F-V--HTNIF--------FQFAIPSAMMCC-LEWLAFEVITLLSGLLPNSKLET | 288 |
| AT1G64820.1 | -L----I-S--SMKQF--------IALAIPSAMMIC-LEWWSFEILLLMSGLLPNSKLET | 289 |
| AT1G66780.1 | -I----F-L--SMKQF--------ITLAIPTAMMTC-LEWWSFELLILMSGLLPNSKLET | 295 |
| Manes.02G072400.1 | -A----F-L--GIGEI--------FRLGVPSAVMVC-LKWWSMELLTLLSGLLPNAKLET | 285 |
| Manes.01G113500.1 | -I----F-S--SISEF--------WRFAVSSSVMVC-LEWWTFELLVLLAGLLKNSKLET | 273 |
| Manes.01G113500.2 | -I----F-S--SISEF--------WRFAVSSSVMVC-LEWWTFELLVLLAGLLKNSKLET | 273 |
| AT1G15150.1 | -I----F-E--GVREF--------IKYALPSAAMLC-LEWWSYELIIILLSGLLPNPQLET | 291 |
| AT1G15160.1 | -I----F-E--GVREF--------IKYALPSAAMLC-LEWWSYELIIILLSGLLPNPQLET | 291 |
| AT1G15170.1 | -I----F-D--GIGEF--------FKYALPSAAMIC-LEWWSYELIIILLSGLLPNPQLET | 294 |
| AT1G15180.1 | -I----F-D--GIGEF--------FRYALPSAAMIC-LEWWSYELIIILLSGLLPNPQLET | 295 |
| AT1G71140.1 | -L----F-E--GMGEF--------FRFGIPSASMIC-LEWWSFEFLVLLSGILPNPKLEA | 286 |
| Manes.06G063000.1 | -L----F-Q--GVGQF--------FRLAIPSAGMIC-LEWWSFEFLTMLSGILPNPRLET | 286 |
| Manes.14G109700.1 | -L----F-H--GIGEF--------FRFALPSAVMIC-LQWWSYELVILLSGLLPNPQLET | 295 |
| Manes.14G109800.1 | -L----F-H--GIGEF--------FRFAIPSAVMIC-LEWWSFELLVLLSGLLPNPELET | 296 |
| Sobic.008G171600.1 | -A----F---RGLWGY--------AKLAFASAVMLA-LEIWYVQGFVLLTGFLPNSEIAL | 303 |
| Sobic.008G171600.2 | -A----F---RGLWGY--------AKLAFASAVMLA-LEIWYVQGFVLLTGFLPNSEIAL | 317 |
| Sobic.008G171600.3 | -A----F---RGLWGY--------AKLAFASAVMLA-LEIWYVQGFVLLTGFLPNSEIAL | 303 |
| AT3G59030.1 | -A----F---RGIWPY--------FKLTVASAVMLC-LEIWYNQGLVIISGLLSNPTISL | 311 |
| Manes.01G182000.1 | -A----F---QDIWPY--------FKLTAASAVMLC-LEIWYNQGMVLISGLLPNPTISL | 310 |
| Manes.02G142000.1 | -A----F---RGIWPY--------FKLTAASAVMLC-LEIWYSQGMVLISGLLSNPTIAL | 310 |
| AT4G21903.2 | -S----L---HGLWSF--------FKLSAGSAVMIC-LELWYTQILVLLAGLLKDPALSL | 312 |
| AT4G21910.4 | -S----L---QGLWSF--------FKLSAGSAVMIC-LEMWYSQILVLLAGLLENPARSL | 316 |
| AT1G11670.1 | -A----F---DGLWDF--------FQLSAASAVMLC-LESWYSQILVLLAGLLKDPELAL | 310 |
| AT1G61890.1 | -A----F---EGLWDF--------FRLSAASAVMLC-LESWYSQILVLLAGLLKNPELAL | 307 |
| Manes.16G008000.1 | -A----F---SGLWPF--------VKLSAGSAVMLC-LETWYSQILILIAGLLDNPEIAL | 310 |
| Manes.17G038200.1 | -A----F---SGLWEF--------VKLSIASAVMLC-LETWYFQILVLIAGLLDDPEIAL | 311 |
| Sobic.002G318300.1 | -A----F---ADLPGF--------AGLSAASAVMLA-LEVWYFQVLILLAGMLPDPQVAL | 300 |
| Manes.17G038300.1 | -A----I---SGLWSF--------FKLSAASALMLC-LETWYYEVLVLIAGLLQNAEIAL | 297 |
| Manes.16G007900.1 | -A----F---FGLWSF--------FKLSAASAVMLC-LETWYFQVLVLIAGLLENAEIAL | 306 |
| Manes.17G038400.1 | -A----F---FGLWGF--------FKLSAASAVMLC-LETWYFQVLVLIAGLLENAEIAL | 306 |
| Sobic.001G012600.1 | -A----F---HSLGSF--------FKLSAASAVMLC-LETWYFQILVLIAGLLKNPELSL | 303 |
| Sobic.001G012600.2 | -A----F---HSLGSF--------FKLSAASAVMLC-LETWYFQILVLIAGLLKNPELSL | 303 |
| Sobic.001G185600.1 | -A----F---SGLPEF--------LKLSFASAVMLC-LETWYTQITVLVAGLLKDPEIAL | 329 |
| Sobic.001G185400.1 | -A----F---SGLWDF--------LKLSAASAVMLC-LETWYFQVLVLIAGLLPNPELAL | 302 |
| AT3G21690.1 | -A----F---SGLWSF--------FKLSAASAVMLC-LETWYFQILVLLAGLLENPELAL | 312 |
| Sobic.001G185800.1 | -A----F---SGLPEF--------LKLSTASAVMLC-LETWYFQVLVLIAGLLDNPQLAL | 306 |
| Sobic.001G185500.1 | -A----F---SGLPSF--------FKLSVASAVMLC-LETWYFQILVLIAGLLKDPELAL | 320 |
| Sobic.001G185500.2 | -A----F---SGLPSF--------FKLSVASAVMLC-LETWYFQILVLIAGLLKDPELAL | 203 |
| Sobic.007G165500.2 | -A----F---RDIWAF--------VRLSFESAVMLC-LEIWYMSTITVLTGDLEDAQIAV | 335 |
| AT4G00350.1 | -A----F---QDVWPF--------LKLSFASAVMLC-LEIWYFMTIIVLTGHLEDPVIAV | 346 |

```
Manes.01G255000.1       -A----F---KDIWGF--------VKLSIASAVMIC-LEIWYFMTIIVLTGHLADPVIAV  408
Sobic.004G349550.1      -A----L---RGLWPF--------ARLSLASAVMLC-LEVWYMTVLVVLTGRLDDAEIAV  202
Sobic.004G349600.1      -A----F---TDLWAF--------VKLSLASAVMLC-LEMWYMMLLVVLTGHLDDAEIAV  305
Sobic.007G176000.1      -A----F---SDLVGF--------ARLSLGSAVMLC-LEFWFYMFLIVIVGNLENAQVAV  310
Sobic.007G176100.1      -A----F---SDLVGF--------ARLSLGSAVMLC-LEFWFYMFLIVIVGNLENAQVAV  315
AT1G47530.1             -A----F---RDLYGF--------VKLSLASALMLC-LEFWYLMVLVVVTGLLPNPLIPV  294
Manes.05G164500.1       -A----F---SDLWGF--------VKLSLASAVMLC-LEFWYLMVLVVITGRLPNPLVPV  290
Manes.05G164500.2       -A----F---SDLWGF--------VKLSLASAVMLC-LEFWYLMVLVVITGRLPNPLVPV  290
Manes.18G030900.1       -A----F---ADLWGF--------VKLSLASAVMLC-LEFWYLMILVVVTGRLPNPLIPV  291
AT1G23300.1             -A----F---KNLRGF--------ARLSLASAVMVC-LEVWYFMALILFAGYLKNPQVSV  303
AT3G26590.1             -A----F---HNLWSF--------VRLSLASAVMLC-LEVWYFMAIILFAGYLKNAEISV  304
AT5G38030.1             -A----F---HNLWSF--------VRLSLASAVMLC-LEVWYLMAVILFAGYLKNAEISV  304
Manes.12G023400.1       -A----F---KNLWGF--------VRLSLSSAVMLC-LEVWYYMALIIILAGYLKDAEVSV  307
Manes.12G023500.1       -A----F---KNLWGF--------VRLSLSSAVMLC-LEIWYYMALILFAGYLKYAEVSV  307
Manes.12G023600.1       -A----F---QNLWGF--------VRLSLASAVMLC-LEVWYFMALILFAGYLKNAEVAV  324
Manes.13G025200.1       -A----F---QSLWGF--------VRLSLASAVMIC-LEIWYLYALILFAGYLKNAEVSV  324
Manes.13G025200.2       -A----F---QSLWGF--------VRLSLASAVMIC-LEIWYLYALILFAGYLKNAEVSV  323
Sobic.001G273100.1      -A----F---SNLGAF--------VKLSLGSAVMIC-LEFWYYTTLLILVGLLPQAKLQI  302
Sobic.001G273100.2      -A----F---SNLGAF--------VKLSLGSAVMIC-LEFWYYTTLLILVGLLPQAKLQI  302
Sobic.001G273000.2      -A----F---NNIGAF--------VKLSLGSAVMIC-LEFWYNTTLLILVGLLKHAKFQL  274
Sobic.001G273000.1      -A----F---NNIGAF--------VKLSLGSAVMIC-LEFWYNTTLLILVGLLKHAKFQL  285
Sobic.001G273000.3      -A----F---NNIGAF--------VKLSLGSAVMIC-------------------------  262
Manes.15G147800.1       -A----L---KSLASF--------LKLSIASAVMLC-LEEWYFTTVILMVGWLDNPEIAV  289
Manes.15G147900.1       -A----L---KSLASF--------LKLSIASAVMLC-LEVWYFTTLILMVGRLDNPEIAV  290
Manes.17G098800.1       -A----L---KSISSF--------LKLSLASAVMLC-LELWYVTAVIFMVGRLHNPAIAV  295
Manes.17G098900.1       -A----L---KSISSF--------LKLSLASAVMLC-LELWYFTAVILMVGWLHNPEIAV  295
Manes.17G098900.2       -A----L---KSISSF--------LKLSLASAVMLC-LELWYFTAVILMVGWLHNPEIAV  202
Sobic.001G162400.1      -A----F---SNLAAF--------VKLSLASAVMLC-LELWYYTAVLILVGFLKNAQLQI  280
Sobic.003G307600.2      -A----F---ASLGGF--------VRLSIASAVMLC-LEMWYYTAVLILVGCLKNPEIQV  202
Sobic.002G232200.1      -A----F---VDLKEF--------VMLSASSGVMVC-LENWYYRILIFLTAYMKSAELAV  298
Sobic.002G232500.1      -A----F---VDLKDF--------IKLSAASGVMLC-LENWYYRILVFLTGYVKNAELAV  294
Sobic.002G232600.1      -A----F---MGLKDF--------VSLSVASGVMTC-LESWYYRLLIFLTAYAKNAELAV  289
Sobic.007G160700.1      -A----F---ADFWEF--------IKLSSASGVMLC-LENWYYRLLVLLTGYLKNAEIAV  306
Sobic.003G126200.2      -A----F---AGLGEF--------IKLSAASGVMLC-LENWYYRILILLTGNLKNAAVAV  317
Sobic.001G476700.1      -A----F---AGMWEF--------VKLSSASGVMLC-LENWYYRILVLLTGNLKDAAIAV  314
Sobic.001G476700.2      -A----F---AGMWEF--------VKLSSASGVMLC-LENWYYRILVLLTGNLKDAAIAV  337
Manes.18G062800.1       -A----F---SGLWEF--------FGLSAASGVMLC-LENWYYRILILMTGHLKNATLAV  296
AT5G44050.1             -S----F---TRLWEF--------TKLSASSGIMVC-LENWYYRMLIVMTGNLEDARIDV  299
AT5G10420.1             -A----F---TGLLEL--------TKLSASSGIMLC-LENWYYKILMLMTGNLVNAKIAV  297
AT5G65380.1             -A----L---TGLWEF--------LKLSASSGVMLC-LENWYYRILIIMTGNLQNARIAV  296
Manes.12G129000.1       -A----F---SGLWEF--------TKLSAASGVMLC-LENWYYRILILMTGNLKNAEIAV  298
Manes.13G097900.1       -A----F---SGLWEF--------TKLSAASGVMLC-LGNWYYKILILMTGNMKNAEIAV  272
Sobic.005G020700.1      -A----F---ADLGAI--------VKLSLSSGVMLC-LELWYNTILVLLTGYMKNAEVAL  330
Sobic.008G019400.1      -A----F---ADLGAI--------IKLSLSSGVMLC-LELWYNTILVLLTGYMKNAEIAL  326
Manes.09G135300.1       -A----F---KDLFPI--------IKLSLSSGAMLC-LELWYSTVLILLTGNMKNAEVSI  289
AT1G33080.1             -A----F---KDLWPV--------FKLSVSSGGMIC-LELWYNSILILLTGNLKNAEVAL  295
AT1G33090.1             -A----F---KDLWPV--------FKLSLSSGGMVC-LELWYNSILVLLTGNLKNAEVAI  295
AT1G33100.1             -A----F---KDLWPV--------LKLSLSSGGMLC-LELWYNSVLVLLTGNLKNAEVAL  292
AT1G33110.1             -A----F---KDLWPV--------FKLSMSSGGMLC-LELWYNSILVLLTGNLKNAEVAL  295
AT3G03620.1             -A----F---VDLIPM--------LKLSISSGFMIC-LEYWYMSILVLMAGYTKDAKIAI  296
AT5G17700.1             -A----F---LDLIPM--------LKLSISSGFMLC-LEYWYMSIIVLMSGYAKDANIAI  293
Manes.09G135200.1       -A----F---SDLLPV--------VKLSVSSGFMFC-LELWYNSILVLIAGYMKNAATAI  287
Manes.09G134800.1       -A----F---SDLVPL--------LKLSLSSGLMLC-LQLWYTAVLVLLAGYMKNAKTAI  284
Manes.08G150400.1       -A----F---TDLVPV--------IKLSISSGLMLC-LEFWYNALLVLLAGYMENAATQV  282
Manes.09G134900.1       -A----F---SDLIPV--------IKLSISSGLMLC-LELWYTASLVLLAGYMKDATTQV  283
Manes.09G135000.1       -A----F---FNLIPV--------IKLSISSGLMLC-LELWYTASLVLLAGYMKNATTQV  283
Manes.09G134900.2       -A----F---SDLIPV--------IKLSISSGLMLC-LELWYTASLVLLAGYMKDATTQV  254
Manes.09G135000.2       -A----F---FNLIPV--------IKLSISSGLMLC-LELWYTASLVLLAGYMKNATTQV  254


Sobic.002G099300.1      ------------------------------------------------------------  79
AT4G39030.1             AAHQVMAQTYRMCNVWGEPLSQTAQSF------MPEM-LYGANRNLPKARTLL--KSL-M  405
Manes.04G084700.1       AAHQVMIQMYGMCVVFGEPLSQTAQSF------MPEL-LYGVERSLEKAQMLL--KSL-M  425
Manes.11G091900.1       AAHQVMIQAFMTCSVWGEPLSQTAQSF------MPEL-MYGTKRSLVKARMLM--KSL-T  423
Sobic.004G019800.2      AAHQVMINVLCMCTVWGEPLSQTAQSF------MPEL-IYGANQNLTKARMLL--KSL-V  287
Sobic.004G019800.3      AAHQVMINVLCMCTVWGEPLSQTAQSF------MPEL-IYGANQNLTKARMLL--KSL-V  287
Sobic.004G019800.4      AAHQVMINVLCMCTVWGEPLSQTAQSF------MPEL-IYGANQNLTKARMLL--KSL-V  287
AT2G21340.1             AAHQVMLQIYTMSTVWGEPLSQTAQSF------MPEL-LFGINRNLPKARVLL--KSL-V  422
Manes.11G092000.1       AAHQVMLQTYSMCTVWGEPLSQTAQSF------MPEL-IYGFNRSLEKARMLQ--KSL-V  420
Sobic.001G476700.3      ------------------------------------------------------------  164
Sobic.009G077000.1      ------------------------------------------------------------  80
```

```
Sobic.001G454900.1      AGFQICCQLWLATSLLADGLAVAGQAV---------LASAFAKNDKKKVVAAT--SRV-L  439
Sobic.003G403000.1      AAFQICTQVWLATSLLADGLAVAGQAM---------IASAFAKEDRYKVAATA--ARV-L  474
Sobic.003G403000.2      AAFQICTQVWLATSLLADGLAVAGQAM---------IASAFAKEDRYKVAATA--ARV-L  474
Sobic.003G403000.3      AAFQICTQVWLATSLLADGLAVAGQAM---------IASAFAKEDRYKVAATA--ARV-L  474
Sobic.007G020600.1      AAFQICLQTWLACSLLADGLAFAGQAI---------LASAFARKDYPKATATA--SRI-L  399
Sobic.007G020600.2      AAFQICLQTWLACSLLADGLAFAGQAI---------LASAFARKDYPKATATA--SRI-L  399
AT3G08040.1             AAFQICLQVWLTSSLLNDGLAVAGQAI---------LACSFAEKDYNKVTAVA--SRV-L  393
Manes.09G027700.1       ------------------------------------------------------------  156
Manes.09G027800.1       ------------------------------------------------------------  2
Manes.09G027900.1       ------------------------------------------------------------  2
Manes.09G026900.1       AAFQVCLQIWMATSLLADGLAVAGQAI---------LASAFANKDHDKAKAIT--SRV-F  382
Manes.09G026900.2       AAFQVCLQIWMATSLLADGLAVAGQAI---------LASAFANKDHDKAKAIT--SRV-F  382
Manes.09G027000.1       AAFQVCLQIWMATSLLADGLAVAGQAM---------LASAFANKDHDRAKAIA--SRV-F  49
Manes.07G006000.1       AAFQVCLQVWLTSSLLADGLAVAGQAI---------IACAFAEKDYQKATTAA--TRV-L  417
Manes.07G006000.2       AAFQVCLQVWLTSSLLADGLAVAGQAI---------IACAFAEKDYQKATTAA--TRV-L  417
Manes.10G143000.1       AAFQVCLQVWLTSSLLADGLAVAGQAI---------IACAFAEKDYQKATTAA--TRV-L  410
AT1G51340.2             AAFQVCLQVWLATSLLADGYAVAGQAI---------LASAFAKKDYKRAAATA--SRV-L  389
Manes.06G164500.1       AAFQVCLQVWLTTSLLADALAVAGQAI---------LASAFAKRDYDKATATA--SRI-L  382
Manes.06G164500.2       AAFQVCLQVWLTTSLLADALAVAGQAI---------LASAFAKRDYDKATATA--SRI-L  382
Manes.06G164500.3       AAFQVCLQVWLTTSLLADALAVAGQAI---------LASAFAKRDYDKATATA--SRI-L  382
Manes.06G164500.4       AAFQVCLQVWLTTSLLADALAVAGQAI---------LASAFAKRDYDKATATA--SRI-L  382
Manes.06G164500.5       AAFQVCLQVWLTTSLLADALAVAGQAI---------LASAFAKRDYDKATATA--SRI-L  382
Manes.14G002600.1       AAFQVCLQVWLTTSLLADGLAVSGQAI---------LASAFAKKDYEKVTATA--SRV-L  366
Sobic.008G006100.1      AGYEICLQVWLTISLLNDALALAGQAL---------LATEYAKGNYKQARTVL--YRV-L  424
Sobic.005G005400.1      AGHQLCLQVWLTISLLNDALALAGQAL---------LASEYAKGNYKQARLVL--YRV-L  407
Sobic.005G005400.3      AGHQLCLQVWLTISLLNDALALAGQAL---------LASEYAKGNYKQARLVL--YRV-L  407
Sobic.005G005400.2      AGHQLCLQVWLTISLLNDALALAGQAL---------LASEYAKGNYKQARLVL--YRV-L  407
AT2G38330.1             AGHQIVLEIWLAVSLLTDALAIAAQSL---------LATTYSQGEYKQAREVL--FGV-L  389
Manes.08G096600.1       AGHQICMQVWLAVSLLNDALALAGQAL---------LAGGYSQGNYEQARQVT--YRV-L  415
Manes.08G096600.2       AGHQICMQVWLAVSLLNDALALAGQAL---------LAGGYSQGNYEQARQVT--YRV-L  415
AT4G38380.1             AAHQICMQVWLAVSLLTDALASSGQAL---------IASSASKRDFEGVKEVT--TFV-L  428
Sobic.002G286800.1      AAHQICLQVWLAVSLLSDALAVSAQAL---------IASSFAKLDYEKVEEVT--FYV-L  445
Sobic.003G149300.2      AAHQICLQVWLAVSLLADSLAVSAQAL---------IASSYAILDYKRVQKIT--MFA-L  415
Sobic.003G149300.1      AAHQICLQVWLAVSLLADSLAVSAQAL---------IASSYAILDYKRVQKIT--MFA-L  415
Sobic.003G149300.3      AAHQICLQVWLAVSLLADSLAVSAQAL---------IASSYAILDYKRVQKIT--MFA-L  415
Sobic.003G149300.4      AAHQICLQVWLAVSLLADSLAVSAQAL---------IASSYAILDYKRVQKIT--MFA-L  291
Manes.01G153400.1       AAHQICLQVWLSVSLLVDAQAASCQAPSLFNHNLPLLLSSAAKGDHSRVKEIA--LCS-L  174
Manes.04G064900.1       AAHQICMQVWLAVSLLTDALAASGQAL---------TASYLSKGDYKNVKEVA--NFV-L  456
Manes.04G064900.2       AAHQICMQVWLAVSLLTDALAASGQAL---------TASYLSKGDYKNVKEVA--NFV-L  456
Sobic.001G003700.1      AVVAVTLNFDYLLFAAMLSLSVSASVR---------VSNELGAGDAALARRAA--RVS-V  329
AT4G22790.1             SILIIVFNFDYLLYAVMLSLGTCVATR---------VSNELGANNPKGAYRAA--YTT-L  339
Manes.02G032300.1       GVLAIVLNFDYLLFSVMLSLATCASIR---------VSNELGANEACPAYQAA--YVS-L  351
Manes.10G000400.1       AAMGILIQTTGLLYVVPYSLNLGLSAR---------VGQELGAGQPSQAKRAT--TVG-L  348
AT2G38510.1             AAMGILIQTTGILYVVPFAISSAIATR---------VGHALGGGQPTRAQCTT--VIG-L  315
Manes.08G172300.1       AATGILIQTAGLIYSFPFSLSCSLSTR---------VGHALGAGQPARAQWTA--IIG-I  345
Manes.09G117000.1       AATGILIQTAGLIYSFPFSLSCSLSTR---------VGHALGAGEPARAQWTA--IIG-V  315
Sobic.010G167800.1      ASMGVLMQTTALVYVFPSSLGLGASTR---------VGNELGANRPGRARAAA--HVA-V  369
AT5G19700.1             ASMGILIQTTSLLYIFPSSLGLAVSTR---------VGNELGSNRPNKARLSA--IVA-V  344
AT4G29140.1             AAMGVLIQTTSFLYVFPSSLSFAVSTR---------VGNELGANRPKTAKLTA--TVA-I  364
Manes.03G026500.1       ASMGILIQTTSLLYVFPSSLGFAVSTR---------VGNELGANRPDKAKLSA--AVA-V  371
Manes.16G109300.1       ASMGILIQTTSLLYVFPSSLGFAVSTR---------VGNELGANRPHKARLSA--VVA-V  365
Sobic.007G181100.1      AAMGILIQTTSLLYIFPHSLSCAVSTR---------VGHELGAGRPERARLVA--RVG-L  365
Manes.13G127800.1       ATMGILIQATSLVYIFPSSLSLAVSTR---------VGNELGANHPSKAKTSS--IVA-L  363
Sobic.001G446800.1      ASMGILIQATSLVYVFPSSLGQGASTR---------VSHQLGAGRPAGARRAA--GAA-L  361
AT5G52050.1             ASMGILIQITSLVYIFPHSLSLGVSTR---------VGNELGSNQPKRARRAA--IVG-L  349
AT1G58340.1             ASMGILIQTTALVYVFPSSLSLGVSTR---------ISNELGAKRPAKARVSM--IIS-L  366
Manes.05G186700.1       ASMGILIQTTSLVYVFPSSLSLGVSTR---------VGNELGANRPAKARISM--IVS-L  357
Manes.18G054300.1       ASMGILIQTTSLVYVFPSSLSLGVSTR---------VGNELGANRPAKARISM--IVS-L  357
Sobic.001G320900.1      ASMGVLIQTTSLLYIFPSSLSFGVSTR---------VSNELGANRPGAARAAA--RAG-L  389
Sobic.001G320900.2      ASMGVLIQTTSLLYIFPSSLSFGVSTR---------V-----------------------  369
Sobic.004G283500.1      ASMGILIQTTSLIYIFPSSLGFGVSTR---------VSNELGANRPDHAGRAA--TVG-L  387
Sobic.006G184400.1      ASMGILIQTTSLIYIFPSSLSFGVSTR---------VSNELGANRPEDASRAA--TVG-L  400
AT4G23030.1             ASMGILIQTTALIYIFPSSLSISVSTR---------VGNELGANQPDKARIAA--RTG-L  341
Manes.01G067000.1       ASMGILIQTTALIYIFPSSLSFGVSTR---------VGNELGANNPQKAKLAA--IIG-L  361
Manes.02G027800.1       ASMGILIQTTALIYIFPSSLSFSVSTR---------VGNELGANHPQKAKLAA--IIG-L  365
Manes.06G143600.1       ASMGILIQTTSLIYIFPSSLSFSVSTR---------VGNELGANQPKKAKLAA--IVG-L  373
Manes.14G029100.1       ASIGILIQTTSLIYIFPSSLGFSVSTR---------VSNEMGGNQPKKAKLAA--IVG-L  377
AT5G49130.1             AAAAIVIQTTSLMYTIPTALSAAVSTR---------VSNELGAGRPEKAKTAA--TVA-V  348
Manes.03G198000.1       ATSAIVIQTTSLLYTLPTALSSSVATR---------VGNELGAGRPVKARLAT--AVA-I  335
Manes.15G011000.1       ATSAIVIQTTSLMYTLPTALSASVSTR---------VGNELGAGRPGKARLAT--AVA-I  344
Sobic.001G019700.1      GAAGVLIQTTSLMYTVPMALAACVSTR---------VGNELGAGKPRRARMAA--LVA-L  353
```

```
AT1G71870.1           AATGILIQTTSLMYTVPMALAGCVSAR---------VGNELGAGRPYKARLAA--NVA-L  349
Manes.02G189800.1     AATGILIQTTSMMYTVPMALAGCVSAR---------VGNELGAGKPYKAKLAA--MVA-L  331
Manes.18G098800.1     AATGILIQTTSMMYTVPMALAGCVSAR---------VGNELGAGRPYKARLAA--MVA-L  331
Sobic.010G256932.1    SILSVSLNTMWMVHAIPCGLSSAISIR---------VF-------PDAARLSV--YVS-G  302
Sobic.010G256700.1    SILSVSLNTMWMVYTIPSGLSSAISIR---------VSNELGAGNPHAARLSV--YVS-G  340
Sobic.010G256700.5    SILSVSLNTMWMVYTIPSGLSSAISIR---------VSNELGAGNPHAARLSV--YVS-G  340
Sobic.010G256700.6    SILSVSLNTMWMVYTIPSGLSSAISIR---------VSNELGAGNPHAARLSV--YVS-G  340
Sobic.010G256700.4    SILSVSLNTMWMVYTIPSGLSSAISIR---------VSNELGAGNPHAARLSV--YVS-G  340
Sobic.009G106900.1    ------------------------------------------------------------  85
Manes.S031500.1       SALSISLNTSAMLYMIPLGLSAATSTR---------VSNELGAGRPQAASLTV--CVA-T  342
Manes.14G109900.1     SVLSISLNTCWMVYMISVGLGGAISTR---------VSNELGAGHPQGARLAL--CVM-I  352
Manes.14G109900.2     SVLSISLNTCWMVYMISVGLGGAISTR---------VSNELGAGHPQGARLAL--CVM-I  251
Manes.14G109900.3     SVLSISLNTCWMVYMISVGLGGAISTR---------VSNELGAGHPQGARLAL--CVM-I  251
AT1G73700.1           SVLSICLNTSLTIWQISVGLGGAASIR---------VSNELGAGNPQVAKLAV--YVI-V  332
AT2G34360.1           SC-------PRTVWMIPFGLSGAASTR---------VSNELGSGNPKGAKLAV--RVV-L  329
AT5G52450.1           SVLSICLNTSGTMWMIPFGLSGAASTR---------ISNELGAGNPKVAKLAV--RVV-I  334
Sobic.009G106960.1    --------------MVPLGLGTSTSTC---------VSNELGAGQPGAARLAA--RVVVV  35
Manes.14G060800.1     SMMSISLNTSSVVFRIPFGLGSAVSTR---------VSNELGAGRPDAACLAV--RIV-I  345
Sobic.009G106800.1    SVLSICLNTGALLFMVPFGLCTAISTR---------VSNELGAGEPQAAKLAT--RVV-M  331
Sobic.007G074300.2    SVLSICLNTSILLFMVPLGLSYSVSTR---------VSNELGAGQPQAAKLAM--RVV-M  155
Sobic.009G106700.1    SVLSICLNTGALLYMIPLGLTYSISTR---------VSNELGAGQPQAAKMAT--KVV-M  333
Sobic.010G138400.4    SVLSISLNTATIVWMIPFGLGSAISTR---------VSNELGAGRPQAARLAV--RVV-V  258
Sobic.010G138400.1    SVLSISLNTATIVWMIPFGLGSAISTR---------VSNELGAGRPQAARLAV--RVV-V  336
Sobic.010G138400.5    SVLSISLNTATIVWMIPFGLGSAISTR---------VSNELGAGRPQAARLAV--RVV-V  255
Sobic.004G129900.1    AVMSICFNTYVFAFMFPMGLGAAASIR---------VSNELGAGRPPAARLAT--RVV-M  333
Sobic.004G129900.2    AVMSICFNTYVFAFMFPMGLGAAASIR---------VSNELGAGRPPAARLAT--RVV-M  251
Sobic.006G042200.1    SVLSISLNTGSLAFMIPFGLSAAISTR---------VSNELGAGRPHAAHLST--RVV-M  336
Sobic.004G129700.1    AVLSICLNTNSFAFMVPLGLGGAVSTR---------VSNELGAGHPRAARLAT--RVV-A  345
Sobic.004G129800.1    AVLSICLNTSSLAFMAPLGLGAAVSTR---------VSNELGAGRPQAARLAA--RVV-V  336
Sobic.004G129800.2    AVLSICLNTSSLAFMAPLGLGAAVSTR---------VSNELGAGRPQAARLAA--RVV-V  251
AT3G23550.1           SLVAICVNTESISYMLTCGLSAATSTR---------VSNELGAGNVKGAKKAT--SVS-V  339
AT3G23560.1           SLVAICVNTEAISYMLTYGLSAAASTR---------VSNELGAGNVKGAKKAT--SVS-V  347
Manes.15G088100.1     SLIAICVNTESVAYMLTYGLSAAASTR---------VSNELGAGNPGRAKGAM--AVT-L  359
Manes.03G109100.1     SLLSMCVNAETVAFMLADGLSAAASTR---------ISNELGAGNPDRAKNAV--AVT-L  343
Manes.03G109100.2     SLLSMCVNAETVAFMLADGLSAAASTR---------ISNELGAGNPDRAKNAV--AVT-L  304
Sobic.002G311200.1    SLIAMCSSTEAIAYMITFGFSAAVSTR---------VSNEIGAGNVERAKNAV--SVT-M  454
Sobic.002G006500.3    SIIAMCQNTETISYMITYGFAAVISTR---------VSNELGARNIGKAKKAL--TVS-L  368
Sobic.002G006500.4    SIIAMCT-------------------R---------VSNELGARNIGKAKKAL--TVS-L  349
AT2G04090.1           SVLSICLTTSSLHCVIPMGIGAAGSTR---------ISNELGAGNPEVARLAV--FAG-I  338
AT2G04100.1           SVLSICLTTSSLHYVIPMGIGAAGSIR---------VSNELGAGNPEVARLAV--FAG-I  338
AT2G04066.1           ---------SVHNYSRSPGFY-----------------------VEKILKVSIKRIEK-K  30
AT2G04040.1           SVLSICLTIETLHYVISAGVAAAVSTR---------VSNNLGAGNPQVARVSV--LAG-L  335
AT2G04080.1           SVLSICLTIGTLHYVIPSGVAAAVSTR---------VSNKLGAGNPQVARVSV--LAG-L  335
AT2G04050.1           SVLSICLTTASLHYVIPGGVAAAVSTR---------VSNKLGAGIPQVARVSV--LAG-L  335
AT2G04070.1           SVLSICLTTATLHYVIPVGVAAAVSTR---------VSNKLGAGIPQVARVSV--LAG-L  335
Sobic.001G481800.1    SVLSICISTVVLVYNLPYGIGTAASVR---------VSNELGAGNPDGARLVV--VVA-L  362
Sobic.003G260200.1    SVLSICLTSVTLLFTIPFGLGAAGSTR---------VANELGAGNPDGARSAV--RVV-L  345
Sobic.009G224800.1    SVLSICLTTITLMYTIPYGLGAAASTR---------VANELGGGNPEGARSSV--QVV-M  348
Sobic.009G224800.2    SVLSICLTTITLMYTIPYGLGAAASTR---------VANELGGGNPEGARSSV--QVV-M  348
AT1G66760.2           SVISICLTTSSLHYNLVNGIGDAASTN---------VANELGAGNPRGARDSA--AAA-I  336
AT1G64820.1           SVISICLTTSAVHFVLVNAIGASASTH---------VSNELGAGNHRAARAAV--NSA-I  337
AT1G66780.1           SVLSICLTMSSLHYVIVNAIGAAASTH---------VSNKLGAGNPKAARSAA--NSA-I  343
Manes.02G072400.1     SVLSICLTISTLHFTIPYGFGAAASTR---------VSNEVGAGNPHSARLAV--VVA-M  333
Manes.01G113500.1     SVLSICITTTSLHYFVQYGISVAASTR---------VSNELGSGNPQAARTVV--HVV-L  321
Manes.01G113500.2     SVLSICITTTSLHYFVQYGISVAASTR---------VSNELGSGNPQAARTVV--HVV-L  321
AT1G15150.1           SVLSICFETLSITYSIPLAIAAAASTR---------ISNELGAGNSRAAHIVV--YAA-M  339
AT1G15160.1           SVLSVCLQTLSMTYSIPLAIAAAASTR---------ISNELGAGNSRAAHIVV--YAA-M  339
AT1G15170.1           SVLSVCLQTISTMYSIPLAIAAAASTR---------ISNELGAGNSRAAHIVV--YAA-M  342
AT1G15180.1           SVLSVCLQTTATVYSIHLAIAAAASTR---------ISNELGAGNSRAANIVV--YAA-M  343
AT1G71140.1           SVLSVCLSTQSSLYQIPESLGAAASTR---------VANELGAGNPKQARMAV--YTA-M  334
Manes.06G063000.1     SVLSVCLATISTLYTVPDGLGAAASTR---------VSNELGAGNARAAYVAV--WCS-M  334
Manes.14G109700.1     SVLSICLTTIATLYSFPYGLSAAVSTR---------VSNELGAGKPRVARTAV--CCL-M  343
Manes.14G109800.1     SILSVCLTTISTLYAIPYGFGAAASTR---------VSNELGAGNPQAARVAV--YAV-S  344
Sobic.008G171600.1    DSLSICINYWNWDFNIMLGLSYAASIR---------VGNELGASHPKVARFSV--IVV-V  351
Sobic.008G171600.2    DSLSICINYWNWDFNIMLGLSYAASIR---------VGNELGASHPKVARFSV--IVV-V  365
Sobic.008G171600.3    DSLSICINYWNWDFNIMLGLSYAASIR---------VGNELGASHPKVARFSV--IVV-V  351
AT3G59030.1           DAISICMYYLNWDMQFMLGLSAAISVR---------VSNELGAGNPRVAMLSV--VVV-N  359
Manes.01G182000.1     DSISVCMNYWNWDIEFMLGLAAATSVR---------VSNELGAGHPKVAKFSV--IVV-N  358
Manes.02G142000.1     DSISICMNYLNWDMQFMLGLAAAISVR---------VSNELGAGHPKVAKFSV--FVV-N  358
AT4G21903.2           DSLSICMSISALSFMVSVGFNAAVSVR---------TSNELGAGNPKSALFST--WTA-T  360
AT4G21910.4           DSLSICMSISALSFMVSVGFNAAVSVR---------TSNELGAGNPKSAWFST--WTA-T  364
```

| | | |
|---|---|---|
| AT1G11670.1 | DSLAICMSISAMSFMVSVGFNAAASVR---------VSNELGAGNPRSAAFST--AVT-T | 358 |
| AT1G61890.1 | DSLAICMSISAISFMVSVGFNAAASVR---------VSNELGAGNPRAAAFST--VVT-T | 355 |
| Manes.16G008000.1 | DSLAVCLSINGLMFMVSVGFNAAASVR---------VSNELGAGNPKSAEFSV--FIV-N | 358 |
| Manes.17G038200.1 | DSLAVCMSVSALLFMVSVGFNAAASVR---------VSNELGAGNPKSAAFSV--LMV-N | 359 |
| Sobic.002G318300.1 | DSLTVCTSIQSWVFMISVGFNAAASVR---------VGNELGAGNPRSAAFSA--WMV-T | 348 |
| Manes.17G038300.1 | DSLSICMTVSLWVFMISVGFNAAASVR---------VSNELGAGHPKSAAFSV--IIV-T | 345 |
| Manes.16G007900.1 | DSLSVCMTISGWVYMISVGFNAAASVR---------VSNELGAGHPKSAAFSV--IIV-N | 354 |
| Manes.17G038400.1 | DSLSVCMTISGWVYMISVGFNAAASVR---------VSNELGAGHPKSAAFSV--IMV-N | 354 |
| Sobic.001G012600.1 | DSLSICMTINGWVFMISVGFNAAASVR---------VGNELGAGNPRAAAFSV--VVV-T | 351 |
| Sobic.001G012600.2 | DSLSICMTINGWVFMISVGFNAAASVR---------VGNELGAGNPRAAAFSV--VVV-T | 351 |
| Sobic.001G185600.1 | DSLAVCMSISGWVFMVSVGFNAAASVR---------VSNELGAGHPMATSFSV--KVV-T | 377 |
| Sobic.001G185400.1 | DALSVCMTISGWVFMISVGFNAAASVR---------VSNELGAGHPKSAYFSV--WVV-T | 350 |
| AT3G21690.1 | DSLSICMTISGWVFMISVGFNAAISVR---------VSNELGAGNPKSAAFSV--IIV-N | 360 |
| Sobic.001G185800.1 | DSLTVCMTLAGWVFMISIGFNAAASVR---------VGNELGAGHPRAAAFSV--VVV-T | 354 |
| Sobic.001G185500.1 | SSLSVCMTITGWVFMISVGFNAAASVR---------VSNELGAGNPKAASFSV--VVV-T | 368 |
| Sobic.001G185500.2 | SSLSVCMTITGWVFMISVGFNAAASVR---------VSNELGAGNPKAASFSV--VVV-T | 251 |
| Sobic.007G165500.2 | DSLGICMNINGWEGMIFIGLNAAISVR---------VSNELGSGRPRAAWNAV--MVV-V | 383 |
| AT4G00350.1 | GSLSICMNINGWEGMLFIGINAAISVR---------VSNELGSGHPRAAKYSV--IVT-V | 394 |
| Manes.01G255000.1 | GSLSICMNINGWEGMLFIGINAAISVR---------VSNELGSAHPRAAKYSV--IVT-C | 456 |
| Sobic.004G349550.1 | GSVSICLNVNGWEAMLFIGLNAAISVR---------VSNELGSGRARAAKHAV--AAV-V | 250 |
| Sobic.004G349600.1 | DSIAICMNINGWEGMLFIGLSAAISVR---------VSNELGSGRPRASMYAV--MVV-L | 353 |
| Sobic.007G176000.1 | AAVSICTNLFGWQIMVFFGFNAAISVR---------VSNELGAGRPRAAKLAI--LVV-L | 358 |
| Sobic.007G176100.1 | AAVSICTNLFGWQIMVFFGFNAAISVR---------VSNELGAGRPRAAKLAI--LVV-L | 363 |
| AT1G47530.1 | DAISICMNIEGWTAMISIGFNAAISVR---------VSNELGAGNAALAKFSV--IVV-S | 342 |
| Manes.05G164500.1 | DAISICMNIQGWNAMIAIGFNAAISVR---------VSNELGAGNARLAKYSV--IVV-S | 338 |
| Manes.05G164500.2 | DAISICMNIQGWNAMIAIGFNAAISVR---------VSNELGAGNARLAKYSV--IVV-S | 338 |
| Manes.18G030900.1 | DAISICMNIQGWDAMIAIGFNAAISVR---------VSNELGAGNARLAKFSV--KVV-S | 339 |
| AT1G23300.1 | AALSICMNILGWPIMVAFGFNAAVSVR---------ESNELGAEHPRRAKFLL--IVA-M | 351 |
| AT3G26590.1 | AALSICMNILGWTAMIAIGMNTAVSVR---------VSNELGANHPRTAKFSL--LVA-V | 352 |
| AT5G38030.1 | AALSICMNILGWTAMIAIGMNAAVSVR---------VSNELGAKHPRTAKFSL--LVA-V | 352 |
| Manes.12G023400.1 | DALSICMNIFGWTMMICLGMNAAISVR---------VSNELGAGHPRTAKFSV--AVA-V | 355 |
| Manes.12G023500.1 | DALSICLNIFGWTVMICFGINAAISVR---------VSNELGAGHPRTAKFSV--VVA-V | 355 |
| Manes.12G023600.1 | DALSICMNILGWTVMVALGMNAAISVR---------VSNELGAAHPRTAKFSL--VVA-V | 372 |
| Manes.13G025200.1 | DAMSICMNILGWTMMVAMGMNAAISVR---------ISNELGAGHPRTAKFSL--VVA-V | 372 |
| Manes.13G025200.2 | DAMSICMNILGWTMMVAMGMNAAISVR---------ISNELGAGHPRTAKFSL--VVA-V | 371 |
| Sobic.001G273100.1 | DIMSVCLNFEFMTIMVALGFSTAIGVR---------VSNELGANRPKETKFAV--LVA-V | 350 |
| Sobic.001G273100.2 | DIMSVCLNFEFMTIMVALGFSTAIGVR---------VSNELGANRPKETKFAV--LVA-V | 350 |
| Sobic.001G273000.2 | DIMSVCLNYEFMAILVAMGFSTAIGIR---------VSNELGAKRPMETRFAV--LVA-V | 322 |
| Sobic.001G273000.1 | DIMSVCLNYEFMAILVAMGFSTAIGIR---------VSNELGAKRPMETRFAV--LVA-V | 333 |
| Sobic.001G273000.3 | ------LNYEFMAILVAMGFSTAIGIR---------VSNELGAKRPMETRFAV--LVA-V | 304 |
| Manes.15G147800.1 | DAISICMNLQLWTLTIALGFNVAISVR---------VSKELGAGHPKAAKFSM--VVA-V | 337 |
| Manes.15G147900.1 | DAISICMNLQLWALTIALGFNAAISVR---------VSNELGAGNPKAAKFSA--VVT-L | 338 |
| Manes.17G098800.1 | DAVSICMNLQRWTLMIALGFNAALSVR---------VSNELGAGNPKAAKFPV--VVT-L | 343 |
| Manes.17G098900.1 | DAVSICMNLQLWTLMIALGFNAAISVR---------VSNELGAGNPKAAKFSV--VVT-L | 343 |
| Manes.17G098900.2 | DAVSICMNLQLWTLMIALGFNAAISVR---------VSNELGAGNPKAAKFSV--VVT-L | 250 |
| Sobic.001G162400.1 | DVMSICINYQLWTLMVALGFNAAVSVR---------VSNELGANRPKAAKFSV--IVA-V | 328 |
| Sobic.003G307600.2 | DAISICMNYQLWTLMVAVGFNAAVSVR---------VSNELGANHPKAAKFSV--VVA-T | 250 |
| Sobic.002G232200.1 | DALSICMSLTGWEMMIHMGFLEGTGVR---------VANELGAANAHGARFAT--IVS-T | 346 |
| Sobic.002G232500.1 | DALSICISYAGWEMMIHLGFLAGTGVR---------VANELGAANGARARFAT--IVS-M | 342 |
| Sobic.002G232600.1 | DALSICLSWAGWEMMIHFGFLAGTGVR---------VANELGANNGRAAKFAT--IVS-T | 337 |
| Sobic.007G160700.1 | DALSICQTINGWEMMIPFGFLAATGVR---------VANELGAGSGKGARFAI--VVS-I | 354 |
| Sobic.003G126200.2 | DALSICMNINGWEMTIPLAFFAGTGVR---------VANELGAGNGIGARFAA--IVS-S | 365 |
| Sobic.001G476700.1 | DALSICMTINGWEMMIPLAFFAGTGVR---------VANELGAGNGKGARFAT--IVS-S | 362 |
| Sobic.001G476700.2 | DALSICMTINGWEMMIPLAFFAGTGVR---------VANELGAGNGKGARFAT--IVS-S | 385 |
| Manes.18G062800.1 | DALSICMSINGWEFMIPLAFLAATGVR---------VANELGAGNGKAAKFAT--IVS-M | 344 |
| AT5G44050.1 | DSMSICMSINGLEMMVPLAFFAGTSVR---------VANELGAGNGKRARFAM--IIS-V | 347 |
| AT5G10420.1 | DSLSICMSVNGWEMMIPLAFFAGTGVR---------VANELGAGNGKGARFAT--IVS-I | 345 |
| AT5G65380.1 | DSLSICMAINGWEMMIPLAFFAGTGVR---------VANELGAGNGKGARFAT--IVS-V | 344 |
| Manes.12G129000.1 | DALSICMTINGWEMMIPLAFFAATGVR---------VANELGAGNGKGAKFAT--VVS-V | 346 |
| Manes.13G097900.1 | DALSICMTINGWEMMIPLAFFAATGVR---------VANELGAGNGKGAKFAT--LVS-V | 320 |
| Sobic.005G020700.1 | DALSICLNINGWEMMISFGFLAATGVR---------VANELGAGSARRAKFAI--YNV-V | 378 |
| Sobic.008G019400.1 | DALSICLNINGWEMMVSIGFLAAAGVR---------VANELGAGSARRAKFAI--YNV-V | 374 |
| Manes.09G135300.1 | DALAICLNINGWEMMISFGFLAAASVR---------VSNELGRGSSKDAKFSI--VVT-V | 337 |
| AT1G33080.1 | NALAICININALEMMVAFGFMAAASVR---------VSNEIGSGNSNGAKFAT--MVV-V | 343 |
| AT1G33090.1 | DALAICINVNALQMMIALGFLAAVSVR---------VSNELGRGNPEGAKFAT--IVA-V | 343 |
| AT1G33100.1 | DALAICISINALEMMIALGFLAAVSVR---------VSNELGSGNPKGAKFAT--LIA-V | 340 |
| AT1G33110.1 | DALAICLNINGLEMMIALGFLAAASVR---------VSNELGSGNPKGAKFAT--LTA-V | 343 |
| AT3G03620.1 | SAFSICQYIYTWELNICLGFLGAACVR---------VANELGKGDAHAVRFSI--KVI-L | 344 |
| AT5G17700.1 | SAFSICQYIYSWEMNICFGLMGAACVR---------VANELGKGDADAVRFSI--KVV-L | 341 |
| Manes.09G135200.1 | SAFSICLNVSNWQFMICLGFLIAASVR---------VSNELGRGNAKAANFAI--KVV-L | 335 |
| Manes.09G134800.1 | SAFSICLNITAWDAMLCLGFLAAASVR---------VSNELGKEDAKAAKFSV--KLN-L | 332 |

```
Manes.08G150400.1       SALSICLNITGWEFMLCVGFLTASSVR---------VSNELGRGDAKAAKFSV--KVI-F  330
Manes.09G134900.1       SALSICLNITAWELMLFVGFLTSSSVR---------VSNELGRGDAKAAKFSV--KVI-F  331
Manes.09G135000.1       SALSICLNITAWELVLCVGFMTASSVR---------VSNELGRGDAKAAKFSV--KVI-F  331
Manes.09G134900.2       SALSICLNITAWELMLFVGFLTSSSVR---------VSNELGRGDAKAAKFSV--KVI-F  302
Manes.09G135000.2       SALSICLNITAWELVLCVGFMTASSVR---------VSNELGRGDAKAAKFSV--KVI-F  302


Sobic.002G099300.1      ----------------------------------------------------------YL  81
AT4G39030.1             IIGATLGLVLGVIGTAVPGLFPGVYTHD-KVIISEMHRLLIPFFMALSALPMTVSLEGTL  464
Manes.04G084700.1       IVGAILGVVIASAGAFIPWLFPNIFTHD-LNVIQEMHKVLILFFIALSPTPCTHSLEGTL  484
Manes.11G091900.1       IIGTILGLMLGIVGTSVPWLFPKIFTPD-QQVIQEMHKVLVPFFMALAVTPCILSFEGTL  482
Sobic.004G019800.2      IIGAITGLTLGAVGTLVPWLFPSVFTND-QMVIQQMHRVLAPYFSVLVVTPSIHSLEGTL  346
Sobic.004G019800.3      IIGAITGLTLGAVGTLVPWLFPSVFTND-QMVIQQMHRVLAPYFSVLVVTPSIHSLEGTL  346
Sobic.004G019800.4      IIGAITGLTLGAVGTLVPWLFPSVFTND-QMVIQQMHRVLAPYFSVLVVTPSIHSLEGTL  346
AT2G21340.1             IIGATLGIVVGTIGTAVPWLFPGIFTRD-KVVTSEMHKVIIPYFLALSITPSTHSLEGTL  481
Manes.11G092000.1       IIGATLGLVLGVIGTSVPWLCPNLFTPD-ENVIREMHKVLLPYFMALAITPSTHSLEGTL  479
Sobic.001G476700.3      ------------------------------------------------------------  164
Sobic.009G077000.1      ------------------------------------------------------------  80
Sobic.001G454900.1      QLSIVLGMGLTVVLGLAMRFGAGIFTSD-LPVIEVIHKGIPFVAGTQTINSLAFVFDGIN  498
Sobic.003G403000.1      QLGVVLGAALTALLGLGLQFGAGVFTSD-AAVIKTIRKGVPFVAGTQTLNTLAFVFDGIN  533
Sobic.003G403000.2      QLGVVLGAALTALLGLGLQFGAGVFTSD-AAVIKTIRKGVPFVAGTQTLNTLAFVFDGIN  533
Sobic.003G403000.3      QLGVVLGAALTALLGLGLQFGAGVFTSD-AAVIKTIRKGVPFVAGTQTLNTLAFVFDGIN  533
Sobic.007G020600.1      QLALVLGLILSILLGIGLRIGSRLFTSD-QGVLHHIYIGIPFVCLTQPINALAFVFDGIN  458
Sobic.007G020600.2      ----------------------------------------QFVCLTQPINALAFVFDGIN  419
AT3G08040.1             QMGFVLGLGLSVFVGLGLYFGAGVFSKD-PAVIHLMAIGIPFIAATQPINSLAFVLDGVN  452
Manes.09G027700.1       ------------------------------------------------------------  156
Manes.09G027800.1       ------------------------------------------------------------  2
Manes.09G027900.1       ------------------------------------------------------------  2
Manes.09G026900.1       QYGLILGLVLSNLLLGGLQFASRLFTED-VNVLNLISVGIPFVAATQIVNVLAFVFDGIN  441
Manes.09G026900.2       QYGLILGLVLSNLLLGGLQFASRLFTED-VNVLNLISVGIPFVAATQIVNVLAFVFDGIN  441
Manes.09G027000.1       QYGLLLGLVLSIFLFGGLQFASRLFTED-VNVLNLIAVGIPFVAATQIVNVLAFVFDGIN  108
Manes.07G006000.1       QMSFVLGLGLAVVVGVGLHFGDGIFSKD-PNVLHIISIGIPFVAATQPINSIAFVFDGVN  476
Manes.07G006000.2       QMSFVLGLGLAVVVGVGLHFGDGIFSKD-PNVLHIISIGIPFVAATQPINSIAFVFDGVN  476
Manes.10G143000.1       QMSFVLGLGLAVVVGVGLHFGDGIFSKD-PNVLHIISIGIPFVAATQPINSIAFVFDGVN  469
AT1G51340.2             QLGLVLGFVLAVILGAGLHFGARVFTKD-DKVLHLISIGLPFVAGTQPINALAFVFDGVN  448
Manes.06G164500.1       QLGLLLGLMLAVVLGIGLSFGARLFTTD-VNVLHMISIGIPFVAGTQPINALAFVFDGVN  441
Manes.06G164500.2       QLGLLLGLMLAVVLGIGLSFGARLFTTD-VNVLHMISIGIPFVAGTQPINALAFVFDGVN  441
Manes.06G164500.3       QLGLLLGLMLAVVLGIGLSFGARLFTTD-VNVLHMISIGIPFVAGTQPINALAFVFDGVN  441
Manes.06G164500.4       QLGLLLGLMLAVVLGIGLSFGARLFTTD-VNVLHMISIGIPFVAGTQPINALAFVFDGVN  441
Manes.06G164500.5       QLGLLLGLMLAVVLGIGLSFGARLFTTD-VNVLHMISIGIPFVAGTQPINALAFVFDGVN  441
Manes.14G002600.1       QLGLLLGLMLAVILGLGLSFGARLFTTD-VDVLHVISIGIPFVVGTQPINALAFVFDGVN  425
Sobic.008G006100.1      QVGGVTGVALAASLFVGFGSLSLLFTDD-PAVLDVALSGVWFVTISQPVNAIAFVADGLY  483
Sobic.005G005400.1      QIGGVTGIALAVVLFFGFGSFSVLFTND-RAVLDIAKSGVWFVTVSQPINAIAFVIDGLY  466
Sobic.005G005400.3      QIGGVTGIALAVVLFFGFGSFSVLFTND-RAVLDIAKSGVWFVTVSQPINAIAFVIDGLY  466
Sobic.005G005400.2      QIGGVTGIALAVVLFFGFGSFSVLFTND-RAVLDIAKSGVWFVTVSQPINAIAFVIDGLY  466
AT2G38330.1             QVGLATGTGLAAVLFITFEPFSSLFTTD-SEVLKIALSGTLFVAGSQPVNALAFVLDGLY  448
Manes.08G096600.1       EIGIITGIGLGAILFIGFGAFSSLFSTD-SEVLEIAWSGVLFVAGSQPMNALAFVLDGLY  474
Manes.08G096600.2       EIGIITGIGLGAILFIGFGAFSSLFSTD-SEVLEIAWSGVLFVAGSQPMNALAFVLDGLY  474
AT4G38380.1             KIGVVTGIALAIVLGMSFSSIAGLFSKD-PEVLRIVRKGVLFVAATQPITALAFIFDGLH  487
Sobic.002G286800.1      KAGVFVGIALALLLFASFGRLAEVFSKD-PMVIQIVRGGVLFVSASQPINALAFIFDGLH  504
Sobic.003G149300.2      QIGVFSGLALAIGLYASFGNIARLFTSD-PEVLMVVKSCALFVCASQPINALAFIFDGLL  474
Sobic.003G149300.1      QIGVFSGLALAIGLYASFGNIARLFTSD-PEVLMVVKSCALFVCASQPINALAFIFDGLL  474
Sobic.003G149300.3      QIGVFSGLALAIGLYASFGNIARLFTSD-PEVLMVVKSCALFVCASQPINALAFIFDGLL  474
Sobic.003G149300.4      QIGVFSGLALAIGLYASFGNIARLFTSD-PEVLMVVKSCALFVCASQPINALAFIFDGLL  350
Manes.01G153400.1       KTGLITGICLAIILGVSFSSVATLFTKD-VEVLATVRSGLLFVSGSQPINALAYIFDGLH  233
Manes.04G064900.1       KIGLLTGACLAAILGVSFGSIATLFTKD-DEVLGIVRMGVLFVSVSQPMNALAFIFDGLH  515
Manes.04G064900.2       KIGLLTGACLAAILGVSFGSIATLFTKD-DEVLGIVRMGVLFVSVSQPMNALAFIFDGLH  515
Sobic.001G003700.1      AGGLAAGLAGGLLMLAARRPWPRIYTRS-PEVRDGVGRAMKVMAMLEVVNFPLNVCGGIV  388
AT4G22790.1             IVGIISGCIGALVMIAFRGFWGSLYTHHDQLILNGVKKMMLIMAVIEVVNFPLMVCGEIV  399
Manes.02G032300.1       AVSFISGFIGALVMVAARGIWGPLFSHD-KGIIRGVKKMMLLMALVEVVNFPLAVCGGIV  410
Manes.10G000400.1       IVSIFCGLLAFVFTVSVKDWWGKMYTKE-QQILDLISLVLPIVGVCELGNCPQTAACGVL  407
AT2G38510.1             ILAVAYGLAAAVFVTALRSVWGKMFTDE-PEILGLISAALPILGLCEIGNSPQTAACGVL  374
Manes.08G172300.1       ILGFACGLTAAVFTAVFSSVWGKLYTDE-PQILDLISTGLPLLGLCEIGNSPQTAACGVL  404
Manes.09G117000.1       VLGFACGVMATIFTFFFSSIWGKLYTDE-PQVLELISIGLPLLGLCEIGNSPQTAACGVL  374
Sobic.010G167800.1      AGAAGMGLAAMSFAAGVRHAWGRMFTAD-DEILRLTAAALPVVGLCELGNCPQTVGCGVL  428
AT5G19700.1             SFAGVMGLTASAFAWGVSDVWGWIFTND-VAIIKLTAAALPILGLCELGNCPQTVGCGVV  403
AT4G29140.1             VFAAVTGIIAAAFAYSVRNAWGRIFTGD-KEILQLTAAALPILGLCEIGNCPQTVGCGVV  423
Manes.03G026500.1       LISAIMGLTASTFTSGMRERWGRMFTRD-VEILRLTSAALPILGLCELGNCPQTVGCGVL  430
Manes.16G109300.1       FISAIMGLSASTFASGMSQRWGQMFTSD-GEILRLTAAALPILGLCELGNCPQTVGCGVM  424
Sobic.007G181100.1      CCGAALGLLACAFAASVRGVWARMFTTD-AAILRLASAALPILGAAELGNCPQTAGCGVL  424
Manes.13G127800.1       SCAIFTSFIAMLFMTSMRHAWGQIFTTD-TAILSLTATAMPVVGLCELGNCPQTTGCGVL  422
```

| | | |
|---|---|---|
| Sobic.001G446800.1 | SIGVGVGLAAAAFMVSVRSHWGRMFTSD-ADILRLTAVALPIAGLCELGNCPQTAGCGVL | 420 |
| AT5G52050.1 | GLSIALGFTAFAFTVSVRNTWAMFFTDD-KEIMKLTAMALPIVGLCELGNCPQTTGCGVL | 408 |
| AT1G58340.1 | FCAIALGLMAMVFAVLVRHHWGRLFTTD-AEILQLTSIALPIVGLCELGNCPQTTGCGVL | 425 |
| Manes.05G186700.1 | VCALALGLLAMLFTTLMRHQWGRFFTSD-SEILELTAVALPIAGLCELGNCPQTTGCGVL | 416 |
| Manes.18G054300.1 | VCAFVLGLLAMLFTSLMRHQWGRFFTSD-AEILELTAVALPIAGLCELGNCPQTTGCGVL | 416 |
| Sobic.001G320900.1 | ALSALQGLASFLFAVSVRDVWARMFTSD-TSILALTASVLPILGLCELGNCPQTTGCGVL | 448 |
| Sobic.001G320900.2 | ------------------------------------------------------------ | 369 |
| Sobic.004G283500.1 | MLGFAFGGVASAFAYLVRGAWATMFTAD-PAIVALTASVLPILGACELGNCPQTTGCGVL | 446 |
| Sobic.006G184400.1 | MLGFAFGGLASAFAFAVRNVWASMFTAD-PAIIALTASVLPVLGLCELGNCPQTTGCGVL | 459 |
| AT4G23030.1 | SLSLGLGLLAMFFALMVRNCWARLFTDE-EEIVKLTSMVLPIIGLCELGNCPQTTLCGVL | 400 |
| Manes.01G067000.1 | SSSFGLGFSALFFAVMVRKVWATMFTED-AEIIALTSMVLPIIGLCELGNCPQTTGCGVL | 420 |
| Manes.02G027800.1 | CSSFALGFSALFFAVMVRKVWAIMFTED-AEIIALTSMVLPIIGLCELGNCPQTTGCGVL | 424 |
| Manes.06G143600.1 | SCSFILGFSALSFTVTVRKIWASMFTQD-KEIIALTSLVLPIIGLCELGNCPQTTGCGVL | 432 |
| Manes.14G029100.1 | SCSFVLGFSALSFTITVRKIWASMFTQD-KEIIALTSLVLPIIGLCELGNCPQTTGCGVL | 436 |
| AT5G49130.1 | GAAVAVSVFGLVGTTVGREAWGKVFTAD-KVVLELTAAVIPVIGACELANCPQTISCGIL | 407 |
| Manes.03G198000.1 | GLALLSSLFGLVLTTLGRETWGRVFTED-EQVLELCMIVLPIIGVCELANCPQTTSCGIL | 394 |
| Manes.15G011000.1 | GLALLSSLFGLVLTTAGKEAWGRVFTED-EEVLELSMIVLPIIGLCELANCPQTTSCGIL | 403 |
| Sobic.001G019700.1 | WCALAIGVVHVAWTVALSRRWVELFTTE-AGVVRLASAAMPVVGLCELGNCPQTTGCGVL | 412 |
| AT1G71870.1 | ACAFVVGALNVAWTVILKERWAGLFTGY-EPLKVLVASVMPIVGLCELGNCPQTTGCGIL | 408 |
| Manes.02G189800.1 | LCAFVIGIINVTWTVILRERWAGLFTKD-SLVKGLVASVLPIIGLCELGNCPQTTGCGIL | 390 |
| Manes.18G098800.1 | GCAFVIGIINVSWTVILKERWGSLFTKD-GLVKGLVASVLPIIGLCELGNCPQTTGCGIL | 390 |
| Sobic.010G256932.1 | IICLAEGLFLAIITVLVQDVGGYMYINK-EEVVKHVSIMMPILATYDFMDGIQCMLS--- | 358 |
| Sobic.010G256700.1 | IMCLAEGLFVAIITVLVRDVWGYLYSNE-EDVVKHVSIMMPILATSDFMDGTQCTLSGAA | 399 |
| Sobic.010G256700.5 | IMCLAEGLFVAIITVLVRDVWGYLYSNE-EDVVKHVSIMMPILATSDFMDGTQCTLSGAA | 399 |
| Sobic.010G256700.6 | IMCLAEGLFVAIITVLVRDVWGYLYSNE-EDVVKHVSIMMPILATSDFMDGTQCTLSGAA | 399 |
| Sobic.010G256700.4 | IMCLAEGLFVAIITVLVRDVWGYLYSNE-EDVVKHVSIMMPILATSDFMDGTQCTLSGAA | 399 |
| Sobic.009G106900.1 | ------------------------------------------------------------ | 85 |
| Manes.S031500.1 | FLVATEGIFVALSLILGHNVWGYLYSRE-ERVVKYVGKMLIFIGASHFFDGIQSVLSGTA | 401 |
| Manes.14G109900.1 | IIALSEGTVVGISTILVRREWGKLYSNE-EEVIKYVANMMPLLALSDFLDGFQCVLSGAA | 411 |
| Manes.14G109900.2 | IIALSEGTVVGISTILVRREWGKLYSNE-EEVIKYVANMMPLLALSDFLDGFQCVLSGAA | 310 |
| Manes.14G109900.3 | IIALSEGTVVGISTILVRREWGKLYSNE-EEVLLEDVDGRNYVHLSIL--GLIML*---- | 303 |
| AT1G73700.1 | GIAVAEGIVVVTVLLSIRKILGHAFSSD-PKIIAYAASMIPIVACGNFLDGLQCVLSGVA | 391 |
| AT2G34360.1 | SFSIVESILVGTVLILIRKIWGFAYSSD-PEVVSHVASMLPILALGHSLDSFQTVLSGVA | 388 |
| AT5G52450.1 | CIAVAESIVIGSVLILIRNIWGLAYSSE-LEVVSYVASMMPILALGNFLDSLQCVLSGVA | 393 |
| Sobic.009G106960.1 | CMTLSQGVVLATIMILLRNVWGYAYSSD-TRRW*-------------------------- | 67 |
| Manes.14G060800.1 | FLAVAESLSLGLALVAARNVWGFLYTNE-KEVVRYLASVLPVLALSNFMDGMQAVLSGTA | 404 |
| Sobic.009G106800.1 | CIAF----------------------NE-PEVVTYIARMIPVLAISFFTDGLHSCLSGVV | 368 |
| Sobic.007G074300.2 | CLALSSGFLLTMAMILLRSVWGHMYSNE-KEVVAYIAKMMPVLAISFFIDGIHGSLSGVL | 214 |
| Sobic.009G106700.1 | YMALSEGLVISLTMTLLRNIWGYMYSNE-KEIVTYIAKMLPILGISFFIDGLHSSLSGVL | 392 |
| Sobic.010G138400.4 | LLAVSEGLAVGLILVCVRYIWGHAYSNV-EEVVTYVAKMMLVIAVSNFFDGIQCVLSGVA | 317 |
| Sobic.010G138400.1 | LLAVSEGLAVGLILVCVRYIWGHAYSNV-EEVVTYVAKMMLVIAVSNFFDGIQCVLSGVA | 395 |
| Sobic.010G138400.5 | LLAVSEGLAVGLILVCVRYIWGHAYSNV-EEVVTYVAKMMLVIAVSNFFDGIQCVLSGVA | 314 |
| Sobic.004G129900.1 | LLAFSLGVSEGLVMVLARTLLGYAYTND-KEVVLYTARLMPILAACTLLDCLQCVLSGVV | 392 |
| Sobic.004G129900.2 | LLAFSLGVSEGLVMVLARTLLGYAYTND-KEVVLYTARLMPILAACTLLDCLQCVLSGVV | 310 |
| Sobic.006G042200.1 | VLAIVVGILIGLAMILVRNLWGYAYSNE-EEVVKYISKMMPILAVSFLFDCVQCVLSGVA | 395 |
| Sobic.004G129700.1 | VLALAAGVSEGVVMVLVRHQWGYAYSNE-EEVVRYTARMMPLIAVSLVFDGMQSVLSGVV | 404 |
| Sobic.004G129800.1 | LLALIVGMSEGLVMVLVRDLWGYAYSNE-EEVARYTARMMPVLAVSVMLDSQQCVLSGVV | 395 |
| Sobic.004G129800.2 | LLALIVGMSEGLVMVLVRDLWGYAYSNE-EEVARYTARMMPVLAVSVMLDSQQCVLSGVV | 310 |
| AT3G23550.1 | KLSLVLALGVVIAILVGHDAWVGLFSNS-HVIKEGFASLRFFLAASITLDSIQGVLSGVA | 398 |
| AT3G23560.1 | KLSLVLALGVVIVLLVGHDGWVGLFSDS-YVIKEEFASLRFFLAASITLDSIQGVLSGVA | 406 |
| Manes.15G088100.1 | KLSVFLALIVVLALAFGHNIWAAMFSDS-HAIVEDFASMATLLAISITVDSIQGVLSGVA | 418 |
| Manes.03G109100.1 | KLSLILALILVLSLVLGHKTWTNLFSES-RVITKEFESMLPLLAISITLDSVSGVLSGVA | 402 |
| Manes.03G109100.2 | KLSLILALILVLSLVLGHKTWTNLFSES-RVITKEFESMLPLLAISITLDSVSGVLSGVA | 363 |
| Sobic.002G311200.1 | KLSVFLAVTFVLLLAFGHGPWARLFSGS-ATIVSAFGAIAPLMVVSIVLDSAQGVLSGVA | 513 |
| Sobic.002G006500.3 | ALSLMLGVAFLLLLGLGHDLWVRLFSSS-QAVASAFASMTPLLIGSVVLDSTQGVLSGVA | 427 |
| Sobic.002G006500.4 | ALSLMLGVAFLLLLGLGHDLWVRLFSSS-QAVASAFASMTPLLIGSVVLDSTQGVLSGVA | 408 |
| AT2G04090.1 | FLWFLEATICSTLLFTCKNIFGYAFSNS-KEVVDYVTELSSLLCLSFMVDGFSSVLDGVA | 397 |
| AT2G04100.1 | FLWFLEATICSTLLFICRDIFGYAFSNS-KEVVDYVTELSPLLCISFLVDGFSAVLGGVA | 397 |
| AT2G04066.1 | KENRYVTAFFSTLLFTCRNIIGYTFSNS-KEVVDYVADISPLLCLSFILDGLTAVLNGVA | 89 |
| AT2G04040.1 | CLWIVESAFFSILLFTCRNIIGYAFSNS-KEVLDYVADLTPLLCLSFILDGFTAVLNGVA | 394 |
| AT2G04080.1 | CLWLVESAFFSTLLFTCRNIIGYTFSNS-KEVVDYVADISPLLCLSFILDGLTAVLNGVA | 394 |
| AT2G04050.1 | CLWLVESAFFSTLLFTCRNIIGYAFSNS-KEVVDYVANLTPLLCLSFILDGFTAVLNGVA | 394 |
| AT2G04070.1 | CLWLVESSFFSILLFAFRNIIGYAFSNS-KEVVDYVADLSPLLCLSFVLDGFTAVLNGVA | 394 |
| Sobic.001G481800.1 | SIIICTAVLLSITLLSFRHFVGIAFSNE-EEVVNHVTRMVPLLSISVLTDNLQGVLSGIS | 421 |
| Sobic.003G260200.1 | SMAGIDAVIVSGTLLAARRLVGLAYSSE-EEVISSVAAMVPLVCITVITDCLQGVLSGVA | 404 |
| Sobic.009G224800.1 | CIAVMEAVIITIILLASQHILGYAYSSD-KDVVAYVNAMVPFVCVSVAADSLQGVLSGYI | 407 |
| Sobic.009G224800.2 | CIAVMEAVIITIILLASQHILGYAYSSD-KDVVAYVNAMVPFVCVSVAADSLQGVLSGIA | 407 |
| AT1G66760.2 | IIAAVESVIVSSSLFLSRSVWPYAYSNV-EEVISYVTDITPILCISILMDSFLTVLSGIV | 395 |
| AT1G64820.1 | FLGGVGALITTITLYSYRKSWGYVFSNE-REVVRYATQITPILCLSIFVNSFLAVLSGVA | 396 |
| AT1G66780.1 | FLGMIDAAIVSISLYSYRRNWAYIFSNE-SEVADYVTQITPFLCLSIGVDSFLAVLSGVA | 402 |
| Manes.02G072400.1 | CVAGAEAVIVSTSLFFCRHFLGYAYSND-KQVVDYVSIMTPLICLSVIMDSLQAVLSGVA | 392 |

| | | |
|---|---|---|
| Manes.01G113500.1 | VISITEAAITSTTLFFTRYIFGYAFSND-KEVVDYVTEVAPLLCLSVIVDSLLAVLCGIA | 380 |
| Manes.01G113500.2 | VISITEAAITSTTLFFTRYIFGYAFSND-KEVVDYVTEVAPLLCLSVIVDSLLAVLCG*- | 378 |
| AT1G15150.1 | SLAVMDALMVSMSLLAGRHVFGHVFSSD-KKTIEYVAKMAPLVSISIILDSLQGVLSGVA | 398 |
| AT1G15160.1 | SLAVVDALMVGTSLLAGKNLLGQVFSSD-KNTIDYVAKMAPLVSISLILDSLQGVLSGVA | 398 |
| AT1G15170.1 | SLAVIDALIVSMSLLIGRNLFGHIFSSD-KETIDYVAKMAPLVSISLMLDALQGVLSGIA | 401 |
| AT1G15180.1 | SLAVVEILILSTSLLVGRNVFGHVFSSD-KETIDYVAKMAPLVSISLILDGLQGVLSGIA | 402 |
| AT1G71140.1 | VITGVESIMVGAIVFGARNVFGYLFSSE-TEVVDYVKSMAPLLSLSVIFDALHAALSGVA | 393 |
| Manes.06G063000.1 | FVATTLSIIVSSILFASRHVFGYIFSNE-KEVVDYVTDMAPLVSISVILESFQVTLSGVA | 393 |
| Manes.14G109700.1 | FITAAELILVSGTLFVSRHVFGYSFSSD-KEVVDAVSSMAPLVCLSVIIDGLQGVFSGVA | 402 |
| Manes.14G109800.1 | FLAVVETTIVTATLFASRRIFGYVFSNE-KDVVDYVTTMAPLLCLSVIMDSLQGVLSGVA | 403 |
| Sobic.008G171600.1 | VVSIAFSFLATLTVLILRYPLSTLYTSS-ATVIEAVISLMPLMAISIFLNGIQPILSGVA | 410 |
| Sobic.008G171600.2 | VVSIAFSFLATLTVLILRYPLSTLYTSS-ATVIEAVISLMPLMAISIFLNGIQPILSGVA | 424 |
| Sobic.008G171600.3 | VVSIAFSFLATLTVLILRYPLSTLYTSS-ATVIEAVISLMPLMAISIFLNGIQPILSGVA | 410 |
| AT3G59030.1 | ITTVLISSVLCVIVLVFRVGLSKAFTSD-AEVIAAVSDLFPLLAVSIFLNGIQPILSGVA | 418 |
| Manes.01G182000.1 | GTAIIISTIFSVIVLCFRVELSKLFTSD-SEVIEAISNLTPLLAISVFLNGVQPILSGVA | 417 |
| Manes.02G142000.1 | GTSVCISIIFSVIVLSFRVALSKLFTSD-SEVIEAVSNLTPLLAISVFLNGVQPILSGVA | 417 |
| AT4G21903.2 | FVSFVISVVEALVVIASRDNVSYIFTSD-ADVAKAVSDLCPFLAVTIILNGIQPVLSGVA | 419 |
| AT4G21910.4 | FVSFVISVTEALAVIWFRDYVSYIFTED-ADVAKAVSDLCPFLAITIILNGIQPVLSGVA | 423 |
| AT1G11670.1 | GVSFLLSLFEAIVILSWRHVISYIFTDS-PAVAEAVAELSPFLAITIVLNGVQPVLSGVA | 417 |
| AT1G61890.1 | GVSFLLSVFEAIVVLSWRHVISYAFTDS-PAVAEAVADLSPFLAITIVLNGIQPVLSGVA | 414 |
| Manes.16G008000.1 | LVSFIIAVIEAAVVLALRHLISYAFTSG-ETVADAVAELSPFLGITLILNGIQPVLSGVA | 417 |
| Manes.17G038200.1 | LVSLVISLVEAVIVLALRNVISYAFTSG-ETVAAAVSDLCPLLAITLVLNGVQPVLSGVA | 418 |
| Sobic.002G318300.1 | ALSAFVSAIAGLVTFLLRHKLSYIFTSG-EVVSRAVADLCPLLVGTIVLCGIQPVLSGVA | 407 |
| Manes.17G038300.1 | SSSFVISVIAAILVMIFREVISYGFTEG-EAVAKAVSELSPFLAVTLILNGVQPVLSGVA | 404 |
| Manes.16G007900.1 | LCSFIIAVVAAIVVMILRDYLSYAFTDG-ETVSKAVSDLTPFLAVTLILNGVQPVLSGVA | 413 |
| Manes.17G038400.1 | LCSFVIAVIAAILVMMLRDYLSYAFTEG-EVVSKAVSQLSPFLAVTIILNGVQPVLSGVA | 413 |
| Sobic.001G012600.1 | SLSLAVAVVCAVVVLCIRDQLSYFFTGG-EAVARAVSDLCPLLAVTLVLNGVQPVLSGVA | 410 |
| Sobic.001G012600.2 | SLSLAVAVVCAVVVLCIRDQLSYFFTGG-EAVARAVSDLCPLLAVTLVLNGVQPVLSGVA | 410 |
| Sobic.001G185600.1 | TLSLMVASIIAVIVMCLRDYISYVFTKG-DDVARAVSTMTPLLAVTIVLNGIQPVLSGVA | 436 |
| Sobic.001G185400.1 | AVSTLISVMLSIVILCLRNYISYLFTEG-EVVSNAVADLCPLLAITLILNGIQPVLSGVA | 409 |
| AT3G21690.1 | IYSLITCVILAIVILACRDVLSYAFTEG-KEVSDAVSDLCPLLAVTLVLNGIQPVLSGVA | 419 |
| Sobic.001G185800.1 | ALSFVITLAMAVVFLVFRDYLSYIFTEG-ETVARAVSDLCPLLAATLILNGIQPVLSGVA | 413 |
| Sobic.001G185500.1 | LLSFVLSVLISIVILLCRDYISYIFTEG-EDVSRAVSKLTPLLAVTLILNGIQPVLSGVA | 427 |
| Sobic.001G185500.2 | LLSFVLSVLISIVILLCRDYISYIFTEG-EDVSRAVSKLTPLLAVTLILNGIQPVLSGVA | 310 |
| Sobic.007G165500.2 | GEALLIGIVCMALILIFRDSFSIIFTSD-ATLQRAVAKIAGLLGLTMVLNSVQPVVSGVA | 442 |
| AT4G00350.1 | IESLVIGVVCAIVILITRDDFAVIFTES-EEMRKAVADLAYLLGITMILNSLQPVISGVA | 453 |
| Manes.01G255000.1 | VESLLIGILCAGIILATKNEFSNIFTDS-VEMRKAVAKLAYLLGITMILNSVQPVISGVA | 515 |
| Sobic.004G349550.1 | AQSLAMGLVAMALVLAYRNSFAVLFTGD-RDMQAAVGKVAHLLAATMVLNSVQPVISGVA | 309 |
| Sobic.004G349600.1 | AQSLALGLLAMVLVLATREQFPAIFTGD-RHLQKAVSSIGYLLAVTMVLNSVQPVISGVA | 412 |
| Sobic.007G176000.1 | MSSVAIGLAFFVLVLAFRDVYGAPFTDS-PEVVRAVASLGVVFAFSLLLNSVQPVLSGVA | 417 |
| Sobic.007G176100.1 | MSSVAIGLAFFVLVLAFRDVYGAPFTDS-PEVVRAVASLGVVFAFSLLLNSVQPVLSGVA | 422 |
| AT1G47530.1 | ITSTLIGIVCMIVVLATKDSFPYLFTSS-EAVAAETTRIAVLLGFTVLLNSLQPVLSGVA | 401 |
| Manes.05G164500.1 | VTSIAIGVICMAVVFATRDYFPYLFTTS-EAVAKETTRLSILLGITVLLNSLQPVLSGVA | 397 |
| Manes.05G164500.2 | VTSIAIGVICMAVVFATRDYFPYLFTTS-EAVAKETTRLSILLGITVLLNSLQPVLSGVA | 397 |
| Manes.18G030900.1 | VTSISIGVICMAVVFATRDYFPYLFTTS-EAVANETTRLAILLGITVLLNSLQPVLSGVA | 398 |
| AT1G23300.1 | ITSVSIGIVISVTLIVLRDKYPAMFSDD-EEVRVLVKQLTPLLALTIVINNIQPVLSGVA | 410 |
| AT3G26590.1 | ITSTLIGFIVSMILLIFRDQYPSLFVKD-EKVIILVKELTPILALSIVINNVQPVLSGVA | 411 |
| AT5G38030.1 | ITSTVIGLAISIALLIFRDKYPSLFVGD-EEVIIVVKDLTPILAVSIVINNVQPVLSGVA | 411 |
| Manes.12G023400.1 | ISSFTIGVIISVILILARNQYPSLFSKD-SQVQQLVKKLTPLLATSVIINNVQPVLCGVA | 414 |
| Manes.12G023500.1 | ISSLVIGVIISAILLLTRNQYPSLFSKD-SRVKELVSELTPLLAISVIINNVQPVLCGVA | 414 |
| Manes.12G023600.1 | ISSFIIGLILSLILILTRNIYPSLFSND-SQVQELVDELTPLLALCIVINNVQPVLSGVA | 431 |
| Manes.13G025200.1 | ICSFIIGVSLALILILTTNQYPSLFSSD-SQVRDLVIDLTPLLALCIVTNNVQPVLSGVA | 431 |
| Manes.13G025200.2 | ICSFIIGVSLALILILTTNQYPSLFSSD-SQVRDLVIDLTPLLALCIVTNNVQPVLSGVA | 430 |
| Sobic.001G273100.1 | STSIFMGAIFMGVVLIWRTSLPKLFSDS-EEVIHGASKLGHLLALTVCMSSIWPILSGVA | 409 |
| Sobic.001G273100.2 | STSIFMGAIFMGVVLIWRTSLPKLFSDS-EEVIHGASKLGHLLALTVCMSSIWPILSGVA | 409 |
| Sobic.001G273000.2 | STSIFMGSIFMGVVLIWRTSLPKLFSDS-EEVIHGASKLGLLLALTVWMISICPVLSGVA | 381 |
| Sobic.001G273000.1 | STSIFMGSIFMGVVLIWRTSLPKLFSDS-EEVIHGASKLGLLLALTVWMISICPVLSGVA | 392 |
| Sobic.001G273000.3 | STSIFMGSIFMGVVLIWRTSLPKLFSDS-EEVIHGASKLGLLLALTVWMISICPVLSGVA | 363 |
| Manes.15G147800.1 | LTSIIIGVIFTALVLVTKDDYPKVFTGK-PVVMKEASNLCYFLAATIFLNSIQPVLHGVA | 396 |
| Manes.15G147900.1 | LTSTAIGVVFIALILINKNDFPKVFTEK-PVVMKEASKLSYFLAATIFLDSIQPVLYGVA | 397 |
| Manes.17G098800.1 | LTSTITGVVFTALVLVTKNDFPKVFTDK-PAVMKEASKLGYFLAATIFLNSIQPVLHGVA | 402 |
| Manes.17G098900.1 | LTSTISGVVFTALVLVTKNDFPKVFTGK-AAVIKEASKLGYFLAATIFLNSIQPVLHGVA | 402 |
| Manes.17G098900.2 | LTSTISGVVFTALVLVTKNDFPKVFTGK-AAVIKEASKLGYFLAATIFLNSIQPVLHGVA | 309 |
| Sobic.001G162400.1 | LTSGSIGAVFFAVFLAWRTGLPRFFSED-GDVLREASRLGYLLAGSIFLNSVQPVLSGVA | 387 |
| Sobic.003G307600.2 | ATSAVIGVIFTAVALAARKQMPRLFTGD-DVVLRATAKLGYLLAATIFLNSIQPVLSGVA | 309 |
| Sobic.002G232200.1 | AMSFLISLFASLLALIFHNKLAMIFSSS-EAVIDAVDNISVLLALTILLNGIQPVLSGVA | 405 |
| Sobic.002G232500.1 | TTSFLISLFISLLILIFHDKLGMIFSSS-QAVIDAVDNISFLLALTILLNGIQPVLSGVA | 401 |
| Sobic.002G232600.1 | TTSFLICLLISSLALIFHDKLAILFTSS-EAVIDAVDGISVLLALTILLNGIQPVLSGVA | 396 |
| Sobic.007G160700.1 | TTSVVIGLVFWCLILYFDDKIALLFTSS-AVVLDAVHHLSVLLAFTILLNSVQPVLSGVA | 413 |
| Sobic.003G126200.2 | TTSLVIGLFFWVLIMGLHSKIALIFTTS-AVVLDAVDKLSLLLAFTILLNSVQPVLSGVA | 424 |
| Sobic.001G476700.1 | ITSLVIGLFFWVLIMGLHDKFALIFTSS-SVVLDAVDNLSVLLAFTILLNSIQPVLSGVA | 421 |

```
Sobic.001G476700.2      ITSLVIGLFFWVLIMGLHDKFALIFTSS-SVVLDAVDNLSVLLAFTILLNSIQPVLSGVA  444
Manes.18G062800.1       VQSTIIGLIICLIIVIFHNKFALIFTSS-SDVLEEVDKLSIFLAVTILLNSIQPVLSGVA  403
AT5G44050.1             TQSLIIGIIISVLIYFLLDQIGWMFSSS-ETVLKAVNNLSILLSFAILLNSVQPVLSGVA  406
AT5G10420.1             TLSLMIGLFFTVIIVIFHDQIGSIFSSS-EAVLNAVDNLSILLAFTVLLNSVQPVLSGVA  404
AT5G65380.1             TQSLIIGLFFWVLIMLLHNQIAWIFSSS-VAVLDAVNKLSLLLAFTVLLNSVQPVLSGVA  403
Manes.12G129000.1       TTSVIIGIFFWVLIMIFHNQLALIFTSS-ASVLKAVSHLSILLAFTVLLNSVQPVLSGVA  405
Manes.13G097900.1       TTSIIIGLIFLVLIMIFHNQLALIFTSS-APVLEAVSHLSLLLAFTVLLNSVQPILSGVA  379
Sobic.005G020700.1      ITSFSIGFVLFVLFLFFRGGLAYIFTDS-QAVAESVADLSPLLAFSILLNSVQPVLSGVA  437
Sobic.008G019400.1      IISFSIGFVLFVLFLFFRGSLAYIFTES-QAVAKAVADLSPLLAFSILLNSVQPVLSGVA  433
Manes.09G135300.1       LTSFIIGFILFLVFLSLRGKLAYLFTEN-PKVADAVSDLSPLLAFSILMNSIQPVLSGVA  396
AT1G33080.1             STSLSIGIIFFFIFLFLRERVSYIFTTS-EAVATQVADLSPLLAFSILLNSIQPVLSGVA  402
AT1G33090.1             FTSLSIGLVLFFVFLFLRGRISYIFTTS-EAVAAEVADLSPLLAFSILLNSVQPVLSGVA  402
AT1G33100.1             FTSLSIGIVLFFVFLFLRGRISYIFTTS-EAVAAEVADLSPLLAFSILLNSVQPVLSGVA  399
AT1G33110.1             FTSLSLGIVLFFVFLFLRGRVSYIFTTS-EAVAAEVADLSPLLAFSILMNSVQPVLSGVA  402
AT3G03620.1             TISTLMGVIFSALCLAFCGRISYLFSNS-DEVSDAVNDLSVILAVSILLNSIQPILSGVA  403
AT5G17700.1             VVSAVIGVICSALCLAFGGQISYLFSDS-QAVSDAVADLSIVLSISILFNIIQPILSGVA  400
Manes.09G135200.1       CTSACLGVVFWVLALVFGRKLSYIFTDN-EEVADMVSDLSVLLSFTLLLNSIQPVLSGIA  394
Manes.09G134800.1       ATSFCIGVFLWIMCLLFGQKIAYLFTSK-TAVAEYVSSLSLPLAFSVLLNGVQPIFSGAA  391
Manes.08G150400.1       FTSLCVGVLFFFVCLVFDSQIAKLFTNE-QDVIKAVSSLSLLLALSVLLNSFQTVLTGVA  389
Manes.09G134900.1       FTSVCIGVLFFIVCLALDRQIAKIFTNE-QQVIKAVSSLSLLVAFSVLLNSFQAVLTGVA  390
Manes.09G135000.1       FTSLGIGVLFFIVCLAFDRQIAKIFTNE-QQVIKAVSSLSLLVAFSVLLNSFQAVLTGVA  390
Manes.09G134900.2       FTSVCIGVLFFIVCLALDRQIAKIFTNE-QQVIKAVSSLSLLVAFSVLLNSFQAVLTGVA  361
Manes.09G135000.2       FTSLGIGVLFFIVCLAFDRQIAKIFTNE-QQVIKAVSSLSLLVAFSVLLNSFQAVLTGVA  361


Sobic.002G099300.1      YVIYNFCFYHYSIQGC--L---EI*-----------------------------------  100
AT4G39030.1             LAGRDLKFVSSVMSSS--F---IIGCLTLMFVTRSGYGLLGCWFVLVGFQWGRF----GL  515
Manes.04G084700.1       LAGRDFKFISLSMSGC--F---SLGGLLLLLVSSRGYGLPGCWCALVAFQWGRF----FF  535
Manes.11G091900.1       LAGRDLKYLSLSTGGC--F---SLGAVVLLIVSSRGYGLLGCWCTLLGFQWARF----FL  533
Sobic.004G019800.2      LAGRDLRYLSQSMGVC--F---SIGTVLLMLLRNKG-SLPGCWWVLVLFQWSRF----GS  396
Sobic.004G019800.3      LAGRDLRYLSQSMGVC--F---SIGTVLLMLLRNKG-SLPGCWWVLVLFQWSRF----GS  396
Sobic.004G019800.4      LAGRDLRYLSQSMGVC--F---SIGTVLLMFRQLSR-HQAAATGFSVPMRP---------  391
AT2G21340.1             LAGRDLRYISLSMTGC--L---AVAGLLLMLLSNGGFGLRGCWYALVGFQWARF----SL  532
Manes.11G092000.1       MAGRDLKFLSLSMTGC--L---CVGALVLMLVSSRAYGLAGCWCALVGFQWSRF----FL  530
Sobic.001G476700.3      ------------------------------------------------------------  164
Sobic.009G077000.1      ------------------------------------------------------------  80
Sobic.001G454900.1      FGASDYTYSAYSMVAV--A---SVSIPCLV-YLSVHNGFIGIWIALTIYMSLRT----IA  548
Sobic.003G403000.1      FGASDYAFSAYSMIGV--A---AVSIPSLI-FLSSHGGFVGIWVALTIYMGVRA----LA  583
Sobic.003G403000.2      FGASDYAFSAYSMIGV--A---AVSIPSLI-FLSSHGGFVGIWVALTIYMGVRA----LA  583
Sobic.003G403000.3      FGASDYAFSAYSMIGV--A---AVSIPSLI-FLSSHGGFVGIWVALTIYMGVRA----LA  583
Sobic.007G020600.1      YGASDFGYAAYSMVLV--A---IVSIICIL-TLESYSGFIGIWIALVIYMSLRM----FA  508
Sobic.007G020600.2      YGASDFGYAAYSMVLV--A---IVSIICIL-TLESYSGFIGIWIALVIYMSLRM----FA  469
AT3G08040.1             FGASDFAYTAYSMVGV--A---AISIAAVI-YMAKTNGFIGIWIALTIYMALRA----IT  502
Manes.09G027700.1       ------------GVLV--S---IISILCLF-ALSSSHGFFGIWVALTIFMTFRA----YV  194
Manes.09G027800.1       ------------QVLV--S---IISILCLF-ALSSSHGFFGIWVALTIFMTLRA----FV  40
Manes.09G027900.1       ------------QVLV--S---IISILCLF-ALSSSHGFFGIWVALTIFMTLRA----YV  40
Manes.09G026900.1       YGASDFAYSSYSMVLV--S---IISIVCLF-TLSSSHGFFGIWVALTIFMTLRA----FV  491
Manes.09G026900.2       YGASDFAYSSYSMVLV--S---IISIVCLF-TLSSSHGFFGIWVALTIFMTLRA----FV  491
Manes.09G027000.1       YGASDFAYSSYSMVLV--S---IISILCLF-ALSSSHGFFGIWVALTIFMTLRA----FV  158
Manes.07G006000.1       FGASDFAYSAYSMVLV--A---IASIATIF-VLSNTGGFVGIWIALTIFMGLRT----FA  526
Manes.07G006000.2       FGASDFAYSAYSMVLV--A---IASIATIF-VLSNTGGFVGIWIALTIFMGLRT----FA  526
Manes.10G143000.1       FGASDFAYSAYSMVLV--A---IASIAAIF-VLSKTGGFVGIWVALTIFMGLRT----FA  519
AT1G51340.2             FGASDFGYAAASLVMV--A---IVSILCLL-FLSSTHGFIGLWFGLTIYMSLRA----AV  498
Manes.06G164500.1       FGASDFAYSAYSMVVV--A---IVSILCLV-FLSSSYKFIGIWVALGIYMSLRA----SA  491
Manes.06G164500.2       FGASDFAYSAYSMVVV--A---IVSILCLV-FLSSSYKFIGIWVALGIYMSLRA----SA  491
Manes.06G164500.3       FGASDFAYSAYSMVVV--A---IVSILCLV-FLSSSYKFIGIWVALGIYMSLRA----SA  491
Manes.06G164500.4       FGASDFAYSAYSMVVV--A---IVSILCLV-FLSSSYKFIGIWVALGIYMSLRA----SA  491
Manes.06G164500.5       FGASDFAYSAYSMVVV--A---IVSILCLV-FLSSSYKFIGIWVALGIYMSLRA----SA  491
Manes.14G002600.1       FGASDFAYSAYSMVLV--A---IISIICLL-FLSSSYKFIGIWVALTIYMSLRA----SA  475
Sobic.008G006100.1      YGVSDFAYAAYSTFFA--G---AVSSMFLL-VTAPKFGLSGIWAGLTLFMSLRA----VA  533
Sobic.005G005400.1      YGVSDFAYAAYSMFFV--G---AVSSAFLL-AAAPKLGLGGVWSGLVLFMSLRA----AA  516
Sobic.005G005400.3      YGVSDFAYAAYSMFFV--G---AVSSAFLL-AAAPKLGLGGVWSGLVLFMSLRA----AA  516
Sobic.005G005400.2      YGVSDFAYAAYSMFFV--G---AVSSAFLL-AAAPKLGLGGVWSGLVLFMSLRA----AA  516
AT2G38330.1             YGVSDFGFAAYSMVIV--G---FISSLFML-VAAPTFGLAGIWTGLFLFMALRL----VA  498
Manes.08G096600.1       YGVSDFGFAAYSMLLV--G---LISSLFIL-AAAPVFGLAGVWTGLFLFMTLRV----AA  524
Manes.08G096600.2       YGVSDFGFAAYSMLLV--G---LISSLFIL-AAAPVFGLAGVWTGLFLFMTLRV----AA  524
AT4G38380.1             YGMSDFPYAACSMMVV--G---GISSAFML-YAPAGLGLSGVWVGLSMFMGLRM----VA  537
Sobic.002G286800.1      FGVSDFSYSASSMMVV--G---AISSLFLL-YAPQFFGLPGVWAGLALFMSLRM----TA  554
Sobic.003G149300.2      YGVSDFDYVAQATVC*--------------------------------------------  489
Sobic.003G149300.1      YGVSDFDYVAQATIAV--G---VTSSLVLL-WAPSIFGLAGVWAGLTTLMGLRM----AA  524
Sobic.003G149300.3      YGVSDFDYVAQATIAV--G---VTSSLVLL-WAPSIFGLAGVWAGLTTLMGLRM----AA  524
```

| | | |
|---|---|---|
| Sobic.003G149300.4 | YGVSDFDYVAQATIAV--G---VTSSLVLL-WAPSIFGLAGVWAGLTTLMGLRM----AA | 400 |
| Manes.01G153400.1 | YV-------------A--G---AVSSAFML-YAPSIVGLSGVWSALTLFMGMRT----VA | 270 |
| Manes.04G064900.1 | YGVSDFPYAACSMMLV--G---ALSSIFLL-YAPPIIGIRGVWYGLALFMGLRT----AA | 565 |
| Manes.04G064900.2 | YGVSDFPYAACSMMLV--G---ALSSIFLL-YAPPIIGIRGVWYGLALFMGLRT----AA | 565 |
| Sobic.001G003700.1 | RGTARPLLGMYAVVGG--FYVVALPVGVALGFKA-RLGLEGLLAGFLLGAAVSLAVLVT- | 444 |
| AT4G22790.1 | RGTAKPSLGMYANLSG--FYLLALPLGATLAFKA-KQGLQGFLIGLFVGISLCLSILLI- | 455 |
| Manes.02G032300.1 | RGTARPWLGMYANLGG--FYLVALPTAVLLAFKA-GLGLGGLLLGYLVGIAACVTLLVF- | 466 |
| Manes.10G000400.1 | IGSARPKVGACVNFVA--FYLIGLPVSTLLAFKL-KLGVMGLWFGLAASQASCVCLMIY- | 463 |
| AT2G38510.1 | TGTARPKDGARVNLCA--FYIVGLPVAVTTTFGF-KVGFRGLWFGLLSAQMTCLVMMLY- | 430 |
| Manes.08G172300.1 | TGTARPKDGARINLYA--FYLVGLPVAVLLTFKL-KMGFRGLWFGLFAAQISCVSMMLY- | 460 |
| Manes.09G117000.1 | TGTARTKDGARINLYA--FYLVGLPVAVHLTFKL-KMGFRGLWFGLLAAQISCVSMMLY- | 430 |
| Sobic.010G167800.1 | RGSARPTRAAHVNLGA--FYLVGMPVAVLLAFGL-GVGFVGLWIGLLAAQVCCAGLMLF- | 484 |
| AT5G19700.1 | RGTARPSMAANINLGA--FYLVGTPVAVGLTFWA-AYGFCGLWVGLLAAQICCAAMMLY- | 459 |
| AT4G29140.1 | RGTARPSTAANVNLGA--FYLVGMPVAVGLGFWA-GIGFNGLWVGLLAAQISCAGLMMY- | 479 |
| Manes.03G026500.1 | RGSARPSMAANVNLAS--FYLVGMPMAIGLGFRL-GVGLYGLWLGLLSAQLCCAGLMLY- | 486 |
| Manes.16G109300.1 | RGSARPSNAANVNLGA--FYLVGMPVAIGLGFWL-GVGFCGLWLGLLSAQVCCAGLMLY- | 480 |
| Sobic.007G181100.1 | RGSARPGKAARINVSA--FYGVGMPAALALAFWPARLDFAGMWAGMLAAQLVCAALMLH- | 481 |
| Manes.13G127800.1 | RGSARPSLGANINLGS--FYGIGLPIAILMGFMM-GLGLLGLWLGLLAAQVVCAIIMVV- | 478 |
| Sobic.001G446800.1 | RGSARPASGARINLAS--FYLVGMPVGVALAFGA-RLGFAGLWLGLLAAQAACAVWMAR- | 476 |
| AT5G52050.1 | RGSARPKIGANINGVA--FYAVGIPVGAVLAFWF-GFGFKGLWLGMLAAQITCVIGMMA- | 464 |
| AT1G58340.1 | RGCARPTLGANINLGS--FYFVGMPVAILFGFVF-KQGFPGLWFGLLAAQATCASLMLC- | 481 |
| Manes.05G186700.1 | RGSARPTIGANINLGS--FYLVGMPVAILMGFVA-KMGFSGLWLGLLAAQGSCAILMLY- | 472 |
| Manes.18G054300.1 | RGSARPTIGANINLGS--FYLVGMPVAILMGFVA-KMGFAGLWLGLLAAQASCAILMLY- | 472 |
| Sobic.001G320900.1 | RGSARPKDGAHINLGA--FYGVGTPVAVALAFWA-GQGFRGLWLGLLAAQAACVAVMLV- | 504 |
| Sobic.001G320900.2 | ------------------------------------------------------------ | 369 |
| Sobic.004G283500.1 | RGSARPKDAASINLRS--FYLVGTPVALVLAFWY-HYDFQGLWLGLLAAQATCVVRMLL- | 502 |
| Sobic.006G184400.1 | RGSARPKDAASINLRS--FYLVGTPVALVLAFWL-HYDFKGLWFGLLAAQATCMVRMLL- | 515 |
| AT4G23030.1 | RGSARPKLGANINLCC--FYFVGMPVAVWLSFFS-GFDFKGLWLGLFAAQGSCLISMLV- | 456 |
| Manes.01G067000.1 | RGTARPKMGANINLGC--FYLVGMPVAVWLSFYG-GFDFKGLWLGLLAAQGSCVVTMLF- | 476 |
| Manes.02G027800.1 | RGTARPKMGANINLGC--FYLVGMPVAIWLSFYA-GFDFKGLWLGLLAAQGSCVVTMLF- | 480 |
| Manes.06G143600.1 | RGTARPKVGANINLGC--FYLVGMPVAVWLAFFA-GFDFEGLWLGLLTAQGSCVVTMLV- | 488 |
| Manes.14G029100.1 | RGTARPKVGANINLGC--FYLVGMPVAVWLAFFV-GFDFEGLWLGLLAAQGSCVVTMLV- | 492 |
| AT5G49130.1 | RGSARPGIGAKINFYA--FYVVGAPVAVVLAFVW-GLGFMGLCYGLLGAQLACAISILT- | 463 |
| Manes.03G198000.1 | RGSARPVIGAGINFYS--FYLVGAPVAIYLGFVW-ELGFVGLCYGLLAAQIACVVSILT- | 450 |
| Manes.15G011000.1 | RGSARPGIGAGINFYS--FYLVGAPVAIVLGFVW-KLGFVGLCYGLLAAQIACVVSILM- | 459 |
| Sobic.001G019700.1 | RGTARPAVGARINLLS--FYLVGTPVAVYLAFGAPGVGFRGLWYGLLSAQASCVALVLAA | 470 |
| AT1G71870.1 | RGTGRPAVGAHVNLGS--FYFVGTPVAVGLAFWL-KIGFSGLWFGLLSAQAACVVSILYA | 465 |
| Manes.02G189800.1 | RGTARPAIGARINLGS--FYFVGTPVAVGLAFWL-NIGFAGLWFGLLSAQVACAMSILYV | 447 |
| Manes.18G098800.1 | RGTARPVIGARINLGS--FYFVGTPVAVGLAFGL-NIGFVGLWFGLLSAQVACVVSILYV | 447 |
| Sobic.010G256932.1 | -ICGWQKVCSVINLFAYAYYAIGLPSAVTFSFTL-KIGGVA-----------VQIFALVM | 405 |
| Sobic.010G256700.1 | RGCGWQKVCSVINLFA--YYAIGLPSAVTFAFIL-NIGGPLAGNH--MCYGSANIC---- | 450 |
| Sobic.010G256700.5 | RGCGWQKVCSVINLFA--YYAIGLPSAVTFAFIL-NIGGPLAGNH--MCYGSANIC---- | 450 |
| Sobic.010G256700.6 | RGCGWQKVCSVINLFA--YYAIGLPSAVTFAFIL-NIGGPLAGNH--MCYGSANIC---- | 450 |
| Sobic.010G256700.4 | RGCGWQKVCSVINLFA--YYAIGLPSAVTFAFIL-NIGGKGLWLGI-ICAMAVQIFALVV | 455 |
| Sobic.009G106900.1 | ------------------------------------------------------------ | 85 |
| Manes.S031500.1 | RGCGWQKLGAVINLGA--YYLVGLPCSIVLAFVY-HLGGMGFCIGF-IVGLAVHGLGLLA | 457 |
| Manes.14G109900.1 | RGCGWQKLCAFINLGA--YYVVAIPCALLFAFIL-HIGGMGLWMGI-ICGLLVQVVALVT | 467 |
| Manes.14G109900.2 | RGCGWQKLCAFINLGA--YYVVAIPCALLFAFIL-HIGGMGLWMGI-ICGLLVQVVALVT | 366 |
| Manes.14G109900.3 | ------------------------------------------------------------ | 303 |
| AT1G73700.1 | RGCGWQKIGACVNLGS--YYLVGVPLGLLLGFHF-HIGGRGLWLGI-VTALSVQVLCLSL | 447 |
| AT2G34360.1 | RGCGWQKIGAFVNLGS--YYLVGVPFGLLLGFHF-HVGGRGLWLGI-ICALIVQGVCLSL | 444 |
| AT5G52450.1 | RGCGWQKIGAIINLGS--YYLVGVPSGLLLAFHF-HVGGRGLWLGI-ICALVVQVFGLGL | 449 |
| Sobic.009G106960.1 | ------------------------------------------------------------ | 67 |
| Manes.14G060800.1 | RGCGWQKLGACINLGA--YYLVGLPSALVLTFLF-HFGGMGLWMGI-TCGSSVQALLLLA | 460 |
| Sobic.009G106800.1 | TGCGEQKIGARVNLSA--YYLAGIPMAVFLAFVL-HLNGMGLWLGI-VCGSLTKLVLLLW | 424 |
| Sobic.007G074300.2 | TGCGKQKIGAITNLGA--FYLAGIPMAVLLAFVF-HMNGMGLWLGMVVCGSLTKVLLFAS | 271 |
| Sobic.009G106700.1 | TGCGKQKIGAAVNLGA--FYLLGIPMSVLLAFIF-HLNGMGLWLGI-VCGSVTKLVFLLF | 448 |
| Sobic.010G138400.4 | RGCGWQKIGACINLGA--YYIVGIPSAYLFAFVM-RVGGTGLWLGI-ICGLMVQVLLLMI | 373 |
| Sobic.010G138400.1 | RGCGWQKIGACINLGA--YYIVGIPSAYLFAFVM-RVGGTGLWLGI-ICGLMVQVLLLMI | 451 |
| Sobic.010G138400.5 | RGCGWQKIGACINLGA--YYIVGIPSAYLFAFVM-RVGGTGLWLGI-ICGLMVQVLLLMI | 370 |
| Sobic.004G129900.1 | RGCGRQKIGAFINLAA--FYIVGIPVAAIFAFVC-HLGGMGLWFGI-LIGVAVQMVLLLC | 448 |
| Sobic.004G129900.2 | RGCGRQKIGAFINLAA--FYIVGIPVAAIFAFVC-HLGGMGLWFGI-LIGVAVQMVLLLC | 366 |
| Sobic.006G042200.1 | RGCGWQKIGACVNLGA--YYLIGIPAAFCFAFLY-HLGGMGLWLGI-ICALAIQMLLLLT | 451 |
| Sobic.004G129700.1 | RGCGRQKAGAYINLAA--YYLAGVPSAFVFAFVC-RLGGMGLWLGI-MCGLVVQMLLLS | 460 |
| Sobic.004G129800.1 | RGSGRQKTGAFINLAA--YYLAGIPAAFAFAFVC-HLGGMGLWFGI-LCGLVVQMLSLLS | 451 |
| Sobic.004G129800.2 | RGSGRQKTGAFINLAA--YYLAGIPAAFAFAFVC-HLGGMGLWFGI-LCGLVVQMLSLLS | 366 |
| AT3G23550.1 | RGCGWQRLATVINLGT--FYLIGMPISVLCGFKL-KLHAKGLWIGL-ICGMFCQSASLLL | 454 |
| AT3G23560.1 | RGCGWQRLVTVINLAT--FYLIGMPIAAFCGFKL-KFYAKGLWIGL-ICGIFCQSSSLLL | 462 |
| Manes.15G088100.1 | RGCGWQHLAVYANLAT--FYIIGMPIACLLGFKL-KLYVKGLWIGL-ICGLSCQAATLSL | 474 |
| Manes.03G109100.1 | RGCGWQHLAVWANLAT--FYFIGIPLSYLLGFKL-QLYAKGLWIGL-ICGLSCQAFTFFL | 458 |
| Manes.03G109100.2 | RGCGWQHLAVWANLAT--FYFIGIPLSYLLGFKL-QLYAKGLWIGL-ICGLSCQAFTFFL | 419 |

```
Sobic.002G311200.1      RGCGWQHLAAVTNLVA--FYFVGMPLAVLFAFKL-DLRARGLWAGL-ICGLTCQASTLLV  569
Sobic.002G006500.3      RGCGWQHLAAWTNLVA--FYVIGLPLAILFGFKL-GFQTKGLWMGQ-ICGLLCQNCVLFF  483
Sobic.002G006500.4      RGCGWQHLAAWTNLVA--FYVIGLPLAILFGFKL-GFQTKGLWMGQ-ICGLLCQNCVLFF  464
AT2G04090.1             RGSGWQNIGAWANVVA--YYLLGAPVGFFLGFWG-HMNGKGLWIGV-IVGSTAQGIILAI  453
AT2G04100.1             RGSGWQHIGAWANVVA--YYLLGAPVGLFLGFWC-HMNGKGLWIGV-VVGSTAQGIILAI  453
AT2G04066.1             RGCGWQHIGALINVVA--YYLVGAPVGVYLAFSR-EWNGKGLWCGV-MVGSAVQATLLAI  145
AT2G04040.1             RGSGWQHIGAWNNTVS--YYLVGAPVGIYLAFSR-ELNGKGLWCGV-VVGSTVQATILAI  450
AT2G04080.1             RGCGWQHIGALINVVA--YYLVGAPVGVYLAFSR-EWNGKGLWCGV-MVGSAVQATLLAI  450
AT2G04050.1             RGSGWQHIGALNNVVA--YYLVGAPVGVYLAFNR-ELNGKGLWCGV-VVGSAVQAIILAF  450
AT2G04070.1             RGCGWQHIGALNNVVA--YYLVGAPVGIYLAFSC-ELNGKGLWCGV-VVGSAVQAIILAI  450
Sobic.001G481800.1      RGCGWQHLGAYVNLGA--FYLIGIPVGLVAGFAL-HLGGAGFWIGM-IAGGATQVTLLSV  477
Sobic.003G260200.1      RGCGWQHLGAYVNLGS--FYLLGIPMAILLGFVL-HMGSRGLWMGI-VCGSLSQTTLMSA  460
Sobic.009G224800.1      S*----------------------------------------------------------  408
Sobic.009G224800.2      RGCGWQHLGAYVNLGS--FYLVGIPTALFLGFVL-KMEAKGLWMGI-SCGSIVQFLLLAI  463
AT1G66760.2             RGTGWQKIGAYVNITS--YYVIGIPVGLLLCFHL-HFNGKGLWAGL-VTGSTLQTLILFL  451
AT1G64820.1             RGSGWQRIGGYASLGS--YYLVGIPLGWFLCFVM-KLRGKGLWIGI-LIASTIQLIVFAL  452
AT1G66780.1             RGTGWQHIGAYANIGS--YYLVGIPVGSILCFVV-KLRGKGLWIGI-LVGSTLQTIVLAL  458
Manes.02G072400.1       RGCGWQHIGAYINLAA--FYLCGLPVGAVLGFVA-HLRGKGLWIGI-VAGSMVQSALLSL  448
Manes.01G113500.1       RGCGWQRIGAFINLGA--YYFVGLPLSVVLCFVL-HLRGKGLWIGL-LVGTTVQVAMFAL  436
Manes.01G113500.2       ------------------------------------------------------------  378
AT1G15150.1             SGCGWQHIGAYINFGA--FYLWGIPIAASLAFWV-HLKGVGLWIGI-LAGAVLQTLLLAL  454
AT1G15160.1             SGCGWQHIGAYINFGA--FYLWGIPIAASLAFWV-HLKGVGLWIGI-IAGAVLQTLLLAL  454
AT1G15170.1             RGCGWQHIGAYINLGA--FYLWGIPIAASLAFWI-HLKGVGLWIGI-QAGAVLQTLLLAL  457
AT1G15180.1             RGCGWQHIGAYINLGA--FYLWGIPIAASLAFWI-HLKGVGLWIGI-QAGAVLQTLLLTL  458
AT1G71140.1             RGSGRQDIGAYVNLAA--YYLFGIPTAILLAFGF-KMRGRGLWIGI-TVGSCVQAVLLGL  449
Manes.06G063000.1       RGCGWQNLGAYVNLVA--YYICGIPVAAVLGFWL-KFRGKGLWIGI-QVGSFLQNVMLVI  449
Manes.14G109700.1       RGCGWQHIGAYVNLAS--LYLCGVPAAAILGFWL-QLKGRGLWIGI-NIGALLQTLLLSL  458
Manes.14G109800.1       RGSGWQQIGAYINLGA--YYLCGIPVAAVLAFTV-KLRGMGLWIGI-QVGAFTQTVLLAI  459
Sobic.008G171600.1      IGSGWQATVAYVNVGA--YYLIGLPIGCVLGYKT-SLGAAGIWWGL-IIGVAVQTIALVI  466
Sobic.008G171600.2      IGSGWQATVAYVNVGA--YYLIGLPIGCVLGYKT-SLGAAGIWWGL-IIGVAVQTIALVI  480
Sobic.008G171600.3      IGSGWQATVAYVNVGA--YYLIGLPIGCVLGYKT-SLGAAGIWWGL-IIGVAVQTIALVI  466
AT3G59030.1             IGSGWQAVVAYVNLVT--YYVIGLPIGCVLGFKT-SLGVAGIWWGM-IAGVILQTLTLIV  474
Manes.01G182000.1       IGSGWQAVVAYVNLVT--YYIIGLPIGCVLGFKT-NLGVAGIWWGI-IIGVVFQTLTLII  473
Manes.02G142000.1       IGSGWQAVVAYVNLVT--YYIIGLPIGCVLGFKT-SLGVAGIWWGI-IIGVVFQTITLII  473
AT4G21903.2             VGCGWQTYVAYVNIGC--YYIVGIPIGCILGFTF-NFQAKGIWTGM-IGGTLMQTLILLY  475
AT4G21910.4             VGCGWQTYVAYVNVGC--YYVVGIPVGCILGFTF-DFQAKGIWTGM-IGGTLMQTLILLY  479
AT1G11670.1             VGCGWQAYVAYVNIGC--YYIVGIPIGYVLGFTY-DMGARGIWTGM-IGGTLMQTIILVI  473
AT1G61890.1             VGCGWQAFVAYVNIGC--YYVVGIPVGFVLGFTY-DMGAKGIWTGM-IGGTLMQTIILVI  470
Manes.16G008000.1       VGCGWQAFVAYVNVGC--YYVVGIPLGFLLGFKF-DLGAKGIWCGM-IGGTLMQTMILLW  473
Manes.17G038200.1       VGCGWQAFVAYVNVGC--YYVVGIPVGCLLGFKF-DLGAKGIWAGM-MGGTMMQTIILLW  474
Sobic.002G318300.1      VGCGWQATVAYVNIGC--YYFIGIPLGVLLGFKF-DFGIKGLWGGM-IGGTLIQTLILIW  463
Manes.17G038300.1       VGCGWQAFVAYVNVGC--YYIVGVPVGVLLGFVF-QLGVKGIWSGM-IGGTFLQTLILLW  460
Manes.16G007900.1       VGCGWQAFVAYVNVGC--YYLIGVPLGILLGFKF-NLGAQGIWSGM-IGGTFLQTIILLW  469
Manes.17G038400.1       VGCGWQAFVAYVNVGC--YYLIGVPLGVLLGFKF-KLGAQGIWSGM-IGGTFLQTIILLW  469
Sobic.001G012600.1      VGCGWQAFVAYVNVGC--YYIIGVPLGVFLGFYL-DLGAKGIWSGMVIGGTMMQTLILLW  467
Sobic.001G012600.2      VGCGWQAFVAYVNVGC--YYIIGVPLGVFLGFYL-DLGAKGIWSGMVIGGTMMQTLILLW  467
Sobic.001G185600.1      VGCGWQAFVAYVNIAC--YYGIGIPLGCVLGFYF-DLGAMGIWGGM-IGGLIVQTLVLIW  492
Sobic.001G185400.1      VGCGWQQFVAYVNIGC--YYIVGVPLGAILGFVF-KLGVKGIWGGM-IGGTCMQTAILLW  465
AT3G21690.1             VGCGWQTFVAKVNVGC--YYIIGIPLGALFGFYF-NFGAKGIWTGM-IGGTVIQTFILAW  475
Sobic.001G185800.1      V--GWQKLVAYINVGC--YYFVGIPLGILLGFKF-HLGAKGIWTGM-LGGTCMQTLILFW  467
Sobic.001G185500.1      VGCGWQAFVAYVNVGC--YYIVGIPLGCLLGFYF-DLGAAGIWSGM-IGGTFMQTLILVW  483
Sobic.001G185500.2      VGCGWQAFVAYVNVGC--YYIVGIPLGCLLGFYF-DLGAAGIWSGM-IGGTFMQTLILVW  366
Sobic.007G165500.2      VGGGWQGLVAYINLGC--YYIFGLPLGYLLGYKF-NFGVGGIWSGML-CGVTLQTIILLV  498
AT4G00350.1             VGGGWQAPVAYINLFC--YYAFGLPLGFLLGYKT-SLGVQGIWIGMI-CGTSLQTLILLY  509
Manes.01G255000.1       VGGGWQALVAYINLFC--YYLVGLPLGFLLGYKT-SLHVQGIWMGMI-FGTFLQTLILVY  571
Sobic.004G349550.1      IGGGWQALVAYINLGC--YYALGLPLGFCLGYLL-RLGAPGIWAGML-CGTALQTAILLL  365
Sobic.004G349600.1      VGGGWQAVVAYINLGC--YYAFGLPLGFILGYLF-RFGVKGIWAGML-CGTALQTAILSY  468
Sobic.007G176000.1      VGAGWQWLVAYINLGC--YYLVGIPVGYMIAFPL-RGGVQGMWGGML-TGVGLQTLILIA  473
Sobic.007G176100.1      VGAGWQWLVAYINLGC--YYLVGIPVGYMIAFPL-RGGVQGMWGGML-TGVGLQTLILVA  478
AT1G47530.1             VGAGWQALVAYVNIAC--YYIIGLPAGLVLGFTL-DLGVQGIWGGMV-AGICLQTLILIG  457
Manes.05G164500.1       VGAGWQSLVAYINIGC--YYIVGLPAGILLGFTF-SFGVMGIWSGMA-GGILLQTIILII  453
Manes.05G164500.2       VGAGWQSLVAYINIGC--YYIVGLPAGILLGFTF-SFGVMGIWSGMA-GGILLQTIILII  453
Manes.18G030900.1       VGAGWQSLVAYINIAC--YYIVGFPAGIVLGFTF-KFGAVGIWSGMI-GGICLQTIILII  454
AT1G23300.1             VGAGWQGIVAYVNIGC--YYLCGIPIGLVLGYKM-ELGVKGIWTGML-TGTVVQTSVLLF  466
AT3G26590.1             VGAGWQAVVAYVNIAC--YYVFGIPFGLLLGYKL-NYGVMGIWCGML-TGTVVQTIVLTW  467
AT5G38030.1             VGAGWQAVVAYVNIVC--YYVFGIPFGLLLGYKL-NFGVMGIWCGML-TGTVVQTIVLTW  467
Manes.12G023400.1       IGAGWQAIIAYVNIIC--YYAFGIPLGLILGYKL-DMGVRGIWYGMM-SGSMVQTFALSF  470
Manes.12G023500.1       IGAGWQAIIAYVNIVC--YYVVFGVPLGLALGYKL-DMGVRGIWYGMM-SGSVVQALVLSF  470
Manes.12G023600.1       IGAGWQAAVAYVNIAC--YYVVGIPLGLILGYKL-GMGVRGIWYGMM-SGTALQTLALFF  487
Manes.13G025200.1       IGAGWQAIVAYVNIGC--YYVFGIPMGLILGYKL-DMGVIGIWYGML-SGIAVQTLALFL  487
Manes.13G025200.2       IGAGWQAIVAYVNIGC--YYVFGIPMGLILGYKL-DMGVIGIWYGML-SGIAVQTLALFL  486
```

```
Sobic.001G273100.1      VGAGWQVPVAFINVGC--YYLVGIPMGILFGFKL-KHGTMGIWIGML-TGTFLQMSILLA  465
Sobic.001G273100.2      VGAGWQVPVAFINVGC--YYLVGIPMGILFGFKL-KHGTMGIWIGML-TGTFLQMSILLA  465
Sobic.001G273000.2      VGAGWQVSVAFINIGC--FYLVGIPMGILFGIKL-KHGTMGIWMGML-TGTFLQMAILLA  437
Sobic.001G273000.1      VGAGWQVSVAFINIGC--FYLVGIPMGILFGIKL-KHGTMGIWMGML-TGTFLQMAILLA  448
Sobic.001G273000.3      VGAGWQVSVAFINIGC--FYLVGIPMGILFGIKL-KHGTMGIWMGML-TGTFLQMAILLA  419
Manes.15G147800.1       VGAGWQFLVALINIVC--NILKPIARAVT------SLGMG*-------------------  428
Manes.15G147900.1       VGAGWQVSVALINIGC--YYIIGLPVGAVLGYKL-KLGVKGIWSGML-AGCLLQTVILFC  453
Manes.17G098800.1       VGAGWQFSVAFINVGC--YYIIGLPIGAVLGYKF-DLGVKGIWSGML-AGCLLQIIVLIF  458
Manes.17G098900.1       VGAGWQFLVAFINVGC--YYIIGLPIGAVLGYKF-DLGVKGIWSGML-AGCLLQIIILIF  458
Manes.17G098900.2       VGAGWQFLVAFINVGC--YYIIGLPIGAVLGYKF-DLGVKGIWSGML-AGCLLQIIILIF  365
Sobic.001G162400.1      IGAGWQALVAFVNIGS--YYFVGIPLAALFGFKL-SMDAMGIWLGMT-LGTLLQTAILVF  443
Sobic.003G307600.2      IGAGWQSLVAFVNIGS--YYLVGLPLAAVFGFKL-KLNATGIWVGVL-IGTVLQTVILFV  365
Sobic.002G232200.1      IGSGWQALVAYVNVGS--YYFIGVPLGVLLGWRF-NYGVPGIWAGMI-SGTTMQTLILAV  461
Sobic.002G232500.1      VGSGWQALVAYVNIGS--YYLIGVPFGFLLGWGL-HYGVQGIWVGMI-VGTMVQTLILAY  457
Sobic.002G232600.1      VGSGWQALVAYVNIGS--YYIIGVPFGVLLAWGF-HYGVLGIWVGMI-GGTMVQTLILSF  452
Sobic.007G160700.1      VGSGWQALVAYVNVGT--YYLIGVPLGIILGWPL-QFGVGGIWSGMI-GGTAVQTIILAY  469
Sobic.003G126200.2      VGSGWQSTVAYINIGC--YYIIGIPMGVLLGWLF-NLGVLGIWAGMI-GGTAVQTLILAI  480
Sobic.001G476700.1      VGSGWQSMVAYVNIGS--YYLIGIPLGILLGWLF-NLGVLGIWAGMI-GGTAVQTLILAI  477
Sobic.001G476700.2      VGSGWQSMVAYVNIGS--YYLIGIPLGILLGWLF-NLGVLGIWAGMI-GGTAVQTLILAI  500
Manes.18G062800.1       VGSGWQAMVAYVNLGC--YYVIGLPLGFLMGWVF-KLGVKGIWGGMIFGGTAVQTVILAI  460
AT5G44050.1             VGSGWQSLVAFINLGC--YYFIGLPLGIVMGWMF-KFGVKGIWAGMIFGGTMVQTLILIF  463
AT5G10420.1             VGSGWQSYVAYINLGC--YYLIGLPFGLTMGWIF-KFGVKGIWAGMIFGGTAIQTLILII  461
AT5G65380.1             VGSGWQSYVAYINLGC--YYCIGVPLGFLMGWGF-KLGVMGIWGGMIFGGTAVQTMILSF  460
Manes.12G129000.1       VGSGWQKYVAYINLGC--YYLIGVPLGFLMGWLF-HFGVLGIWAGMIFGGTAIQTLILAI  462
Manes.13G097900.1       VGSGWQKYVAYINLGC--YYLIGIPMGFLMGWLF-HLGVLGIWAGMIFGGTAIQTLILAI  436
Sobic.005G020700.1      VGAGWQSVVAYVNVTS--YYLIGIPLGAVLGYVV-GFEVKGIWIGML-LGTLVQTIVLLF  493
Sobic.008G019400.1      VGAGWQSVVAYVNVTS--YYLIGIPLGAVLGYVV-GLHVKGIWIGML-LGTLVQTIVLLF  489
Manes.09G135300.1       VGAGWQSIVAYVNIAC--YYLVGIPVGVVLGYVI-HLQVKGVWIGML-FGTFVQTIILII  452
AT1G33080.1             VGAGWQKYVTVVNLAC--YYLVGIPSGLFLGYVV-GLQVKGVWLGMI-FGIFVQTCVLTV  458
AT1G33090.1             VGAGWQGYVAYINLAC--YYLLGIPVGLVLGYVV-GLQVKGVWIGML-FGIFVQTCVLTI  458
AT1G33100.1             IGAGWQGYVAYVNLAC--YYLVGIPIGVILGYVV-GLQVKGVWIGML-FGIFVQTCVLTV  455
AT1G33110.1             VGAGWQGYVTYVNLAC--YYLVGIPIGIILGYVV-GLQVKGVWIGML-FGIFVQTCVLTV  458
AT3G03620.1             VGAGMQSIVAVVNLAS--YYAIGIPLGLILTYVF-HLGVKGLWSGML-AGIAIQTIILCY  459
AT5G17700.1             IGAGMQSMVALVNLAS--YYAIGVPLGVLLVYVF-NFGIKGLWSGML-AGVGIQTLILCY  456
Manes.09G135200.1       VGSGQQSIVAFINLGC--YYGVGIPLGIVLAYVA-HLQVKGLWIGLL-SGVLTQTVILSC  450
Manes.09G134800.1       VGAGRQSLVAYVNLFC--YYVVGVPVGVILGFAV-HLQVKGIWIGLI-VGAAMQTLTLAY  447
Manes.08G150400.1       VGAGRQSMVAYVNMSS--YYIVGVPIGVVLGYVA-HLQIKGIWIGMT-IGVVLQVLLLGY  445
Manes.09G134900.1       VGAGRQSMVAYINISC--YYIIGVPIGVILGYVF-HLEIKGIWIGMT-IGVVMQVTVLGY  446
Manes.09G135000.1       VGAGRQSMVAYINISC--YYIIGVPIGVILGYVF-HLEIKGIWVGMI-IGVVMQVMVLGY  446
Manes.09G134900.2       VGAGRQSMVAYINISC--YYIIGVPIGVILGYVF-HLEIKGIWIGMT-IGVVMQVTVLGY  417
Manes.09G135000.2       VGAGRQSMVAYINISC--YYIIGVPIGVILGYVF-HLEIKGIWVGMI-IGVVMQVMVLGY  417


Sobic.002G099300.1      ------------------------------------------------------------  100
AT4G39030.1             YLRRLLSP-----------GGILNSD----GPSPYTVEKIKSI-----------------  543
Manes.04G084700.1       VLQRLLSP-----------RGMLSSN----DTTEFKLGRLKAA--*--------------  563
Manes.11G091900.1       TLQRLLSP-----------NGIFSLK----IQASLNYTS*--------------------  557
Sobic.004G019800.2      ALLRLISP-----------TGMLFNK----NFNQAEYVEAKAT--*--------------  424
Sobic.004G019800.3      ALLRLISP-----------TGMLFNK----NFNQAEYVEAKAT--*--------------  424
Sobic.004G019800.4      --CRSINK-----------SISFHN-----RRARTE*-----------------------  409
AT2G21340.1             SLFRLLSR-----------DGVLYSE----DTSRYAEKVKAA------------------  559
Manes.11G092000.1       ALQRLLSP-----------DGILYSE----DLSRYKIEKQKAA--*--------------  558
Sobic.001G476700.3      ------------------------------------------------------------  164
Sobic.009G077000.1      ------------------------------------------------------------  80
Sobic.001G454900.1      STWRMGAA-----------RGPWKFL----RK*---------------------------  565
Sobic.003G403000.1      STWRMAAA-----------QGPWKFL----RQ*---------------------------  600
Sobic.003G403000.2      STWRMAAA-----------QGPWKFL----RQ*---------------------------  600
Sobic.003G403000.3      STWRMAAA-----------QGPWKFL----RQ*---------------------------  600
Sobic.007G020600.1      GFWRIGTA-----------QGPWAYL----RG*---------------------------  525
Sobic.007G020600.2      GFWRIGTA-----------QGPWAYL----RG*---------------------------  486
AT3G08040.1             GIARMATG-----------TGPWRFL----RGRSSSSS----------------------  526
Manes.09G027700.1       GLLRIGTG-----------TGPWSFL----RK*---------------------------  211
Manes.09G027800.1       GLFRIGTG-----------TGPWSFL----RR*---------------------------  57
Manes.09G027900.1       GLLRIGTG-----------TGPWSFL----RK*---------------------------  57
Manes.09G026900.1       GLLRIGTG-----------MGPWSFL----RS*---------------------------  508
Manes.09G026900.2       GLLRIGTG-----------MGPWSFL----RS*---------------------------  508
Manes.09G027000.1       GLFR*-------------------------------------------------------  162
Manes.07G006000.1       GVWRMGTG-----------TGPWRFL----RGGLLP*-----------------------  547
Manes.07G006000.2       GVWRMGTG-----------TGPWRFL----RGGLLP*-----------------------  547
Manes.10G143000.1       GVWRMGTG-----------TGPWGFL----KGRLLP*-----------------------  540
AT1G51340.2             GFWRIGTG-----------TGPWSFL----RS----------------------------  515
```

```
Manes.06G164500.1       GFWRIGTR-----------TGPWKFI----RMS*--------------------------  509
Manes.06G164500.2       GFWRIGTR-----------TGPWKFI----RMS*--------------------------  509
Manes.06G164500.3       GFWRIGTR-----------TGPWKFI----RMS*--------------------------  509
Manes.06G164500.4       GFWRIGTR-----------TGPWKFI----RMS*--------------------------  509
Manes.06G164500.5       GFWRIGTR-----------TGPWKFI----RMS*--------------------------  509
Manes.14G002600.1       GFWRIGTR-----------TGPWKFL----RSC*--------------------------  493
Sobic.008G006100.1      GLWRLGSK-----------DGPWEVI----WSDSE*------------------------  553
Sobic.005G005400.1      GFWRLGSK-----------GGPWKQI----WSDTKLINDKK*------------------  542
Sobic.005G005400.3      GFWRLGSK-----------GGPWKQI----WSDTKLINDKK*------------------  542
Sobic.005G005400.2      GFWSILLL-----------CAG*-------------------------------------  527
AT2G38330.1             GAWRLGTR-----------TGPWKML----WSAPEKPE----------------------  521
Manes.08G096600.1       GMWRLGTK-----------TGPWKLV----WAKGEQERDLSLTCKG--------------  555
Manes.08G096600.2       GMWRLGTK-----------TGPWKLV----WAKGEQERDLSE*-----------------  551
AT4G38380.1             GFSRLMWR-----------KGPWWFM----HTSDKRLA----------------------  560
Sobic.002G286800.1      GFMRLGWR-----------AGPWWFL----HKKEPKYKLQSRKC*---------------  583
Sobic.003G149300.2      ------------------------------------------------------------  489
Sobic.003G149300.1      GILRLLWK-----------SGPWSFL----HEEP*-------------------------  543
Sobic.003G149300.3      GILRLLWK-----------SGPWSFL----HEEP*-------------------------  543
Sobic.003G149300.4      GILRLLWK-----------SGPWSFL----HEEP*-------------------------  419
Manes.01G153400.1       GYMR*-------------------------------------------------------  274
Manes.04G064900.1       GFIRYIEQ-----------HIS*-------------------------------------  576
Manes.04G064900.2       GFIRYIEQ-----------HIS*-------------------------------------  576
Sobic.001G003700.1      VIVCMDWAAEADKAR--TRAGDAGAG-----R-------VEPSATK--APVESE------  482
AT4G22790.1             FIARIDWEKEAGKAQ--ILTCNTEDE-----QTSQGSGQDSHS-----------------  491
Manes.02G032300.1       FVMRINWEVEADQAQ--KLAIDVQES-----IEGREDRRTSATVTD--ANA*--------  508
Manes.10G000400.1       VLVCTDWKYQAQRAK--ELTQLTDED-----GNN--------------------------  490
AT2G38510.1             TLIRTDWSHQVKRAE--ELTSAAADK-----SHSEDETVHAEVQDD--DDVSSN------  475
Manes.08G172300.1       TVYRTDWKHQAKRAD--ELTSAAGER-----DDLENGLLMTDQ*----------------  496
Manes.09G117000.1       TLFRTDWKYQAERAD--ELTLAAGER-----NDLEKCLLTTDQ*----------------  466
Sobic.010G167800.1      VVGSTDWEAQARRAQ--ELTSGAEAD-----VEKPEAHTSATAVGE--GGRPEN-GEQEG  534
AT5G19700.1             VVATTDWEKEAIRAR--KLTCTEGVD-----VVIT------TTQTN--GDL---------  495
AT4G29140.1             VVGTTDWESEAKKAQ--TLTCAETVE-----NDIIKAVVASTIDGE--CDE---------  521
Manes.03G026500.1       VVGTTHWDLEARRAQ--MLTCIGCEN-----NRKL-----LSDEGE--GEE---------  523
Manes.16G109300.1       VVGSTDWDLEARRAQ--MLTCIGCDT-----KILL-----S--EGD--*-----------  512
Sobic.007G181100.1      AVLRTDWDEQAVRAS--VLTGGGIVV-----VADVKSGHADA---A--KVKANN-GMLVV  528
Manes.13G127800.1       VLMRTDWQVEANRAR--ELTGIDDGV-----GEAEAESKRKN---L--QG-------LIS  519
Sobic.001G446800.1      AVAATDWDVEVGRAK--ELTKASSSS-----SSNSHSECNTT---S--SASA----SDVT  520
AT5G52050.1             ATCRTDWELEAERAK--VLTTAVDCG-----SSDDDAKEDME---A--GMVDK-------  505
AT1G58340.1             ALLRTDWKVQAERAE--ELTSQTPGK-----S--PPLLPIAS---S--KSRSTS-GTE--  524
Manes.05G186700.1       VLCRTDWIVEAERAR--ELTSSSSSS-------------------S--SKPEAI-TNNKK  508
Manes.18G054300.1       VLCRTDWMAQAERAK--ELTKTSSAT-----SNNTSILPISS---S--SPPSNP-ENKIK  519
Sobic.001G320900.1      VITRTDWAKQAELAQ--VLAGVAAPG-----GDAVVNGDDDD---G--GKEK-------D  545
Sobic.001G320900.2      ------------------LAGVAAPG-----GDAVVNGDDDD---G--GKEK-------D  394
Sobic.004G283500.1      VIGRTDWAAEAKRAQ--QLTGAGTVE-----ETKDKESSGKG------SHAS-------K  542
Sobic.006G184400.1      VIGRTDWASEAKRSR--QLTGAKDSD-----DKAGGD------------EKS-------R  549
AT4G23030.1             VLARTDWEVEVHRAK--ELMTRSCDG-----DEDDGNTPFLL---D--S-----------  493
Manes.01G067000.1       VLTRTDWKWQALRAK--ELTGNANTA-----DDVEDAETLQD---H--NTLL--------  516
Manes.02G027800.1       VLTGTDWEWQAHRAK--ELTGNVNGG-----DAFEEYETLKN---D--NDSS--------  520
Manes.06G143600.1       VLGCTDWEFQAQRAK--ELTGTVGVV-----IDDVDANQETE---E--NKQS-------K  529
Manes.14G029100.1       VIYCTDWDFQAQRAK--KLTGNLGVV---------DAGKEIE---E--NNSS-------K  529
AT5G49130.1             VVYNTDWNKESLKAH--DLVGKNVIS------PNVDQIIVKCEEGL--H-----------  502
Manes.03G198000.1       VVYKTDWEKESLKAK--HLVGRSTDA-----VMPHEHHIIKCEEDG--KGNEEE------  495
Manes.15G011000.1       VVYKTDWERESLKAK--HLVGKASDQ-----LAHVNQT------------LKSD------  494
Sobic.001G019700.1      VVWRTDWRVEAMRAK--KLAGLELAH-----PVPTTTAADD--ADD--AAEERK------  513
AT1G71870.1             VLARTDWEGEAVKAM--RLTSLEMRK-----VGQDEESSLL---LL--DDEKLG------  507
Manes.02G189800.1       LLIRTDWEHEAMKSR--KLTSIEMSP-----SNGVRGNEHEKEEEE--DDDESR------  492
Manes.18G098800.1       VLVRTDWEHEAWKSR--KLTSIEMSP-----SDSVEGKEHEEEEE---------R------  486
Sobic.010G256932.1      MMLRTS*-----------------------------------------------------  411
Sobic.010G256700.1      -FGRDDASNQLE*-----------------------------------------------  461
Sobic.010G256700.5      -FGRDDASNQLE*-----------------------------------------------  461
Sobic.010G256700.6      -FGRDDASNQLE*-----------------------------------------------  461
Sobic.010G256700.4      MMLRTNWNEEAEKAQ--ARVQFSDGS-----ITLI*------------------------  483
Sobic.009G106900.1      ------------------------------------------------------------  85
Manes.S031500.1         VTMSTNWKNESMKAR--DRAYDSTAI-----PKDPLA*----------------------  487
Manes.14G109900.1       VNACTNWDQEAEKAM--HRVGETSSA-----DVGVWKN*---------------------  498
Manes.14G109900.2       VNACTNWDQEAEKAM--HRVGETSSA-----DVGVWKN*---------------------  397
Manes.14G109900.3       ------------------------------------------------------------  303
AT1G73700.1             VTIFTNWDKEAKKAT--NRVGSSDDK-----DGDVQ------------------------  476
AT2G34360.1             ITFFTNWDEEVKKAT--SRAKSSSEV-----KEFAVDNGSIL-------V----------  480
AT5G52450.1             VTIFTNWDEEAKKAT--NRIESSSSV-----KDFAVDDRSVV-------VF---------  486
Sobic.009G106960.1      ------------------------------------------------------------  67
```

```
Manes.14G060800.1     ITMHTDWDQEAKKAR--VTVYGSRIP-----GDVPAEVV*--------------------  492
Sobic.009G106800.1    ITLRINWEKEAIKAK--ETVFSSTLP-----IA*--------------------------  450
Sobic.007G074300.2    VAWFIDWNKEAVKAK--DRVFSSSLP-----VT*--------------------------  297
Sobic.009G106700.1    VTCSIDWDNEAVKAK--YRVLSSSLP-----LV*--------------------------  474
Sobic.010G138400.4    ITVCTNWDNEATKAK--SRVFSSSSP-----ASQT*------------------------  401
Sobic.010G138400.1    ITVCTNWDNEATKAK--SRVFSSSSP-----ASQT*------------------------  479
Sobic.010G138400.5    ITVCTNWDNEATKAK--SRVFSSSSP-----ASQT*------------------------  398
Sobic.004G129900.1    ITLYTNWNKEVLKAN--DRVFSCPLP-----VDDTITSGCSEQGNG--CNFVGNKQDDAN  499
Sobic.004G129900.2    ITLYTNWNKEVLKAN--DRVFSCPLP-----VDDTITSGCSEQGNG--CNFVGNKQDDAN  417
Sobic.006G042200.1    ITLCSNWEKEALKAK--DRVYSSSLP-----VDMMT*-----------------------  480
Sobic.004G129700.1    ITLCTNWNNEALKAK--DRVFSSALP-----PLD*-------------------------  487
Sobic.004G129800.1    VTLCTNWNEEALNAK--SRVFSDALP-----VDMET*-----------------------  480
Sobic.004G129800.2    VTLCTNWNEEALNAK--SRVFSDALP-----VDMET*-----------------------  395
AT3G23550.1           MTIFRKWTKLTAATV---------------------------------------------  469
AT3G23560.1           MTIFRKWTKLNVATV---------------------------------------------  477
Manes.15G088100.1     ITIRAKWTRVADLSISRDEENPLVG*----------------------------------  499
Manes.03G109100.1     MTMRTKWSSTELLKGTDEENIVLI*-----------------------------------  482
Manes.03G109100.2     MTMRTKWSSTELLKGTDEENIVLI*-----------------------------------  443
Sobic.002G311200.1    ITVRTKWSKLAEAMQEKKASYVA*------------------------------------  592
Sobic.002G006500.3    ITLRTNWEELDLTMFNKDNDFVC*------------------------------------  506
Sobic.002G006500.4    ITLRTNWEELDLTMFNKDNDFVC*------------------------------------  487
AT2G04090.1           VTACLSWEEQAAKAR--ERIVGRTLE----------------------------------  477
AT2G04100.1           VTACMSWNEQAAKAR--QRIVVRTSS-----FGNGLA-----------------------  483
AT2G04066.1           VTASMNWKEQAEKAR--KRIISTKNG-----LV---------------------------  171
AT2G04040.1           VTASINWKEQAEKAR--KRIVSTENR-----LA---------------------------  476
AT2G04080.1           VTASMNWKEQAEKAR--KRIISTENG-----LV---------------------------  476
AT2G04050.1           VTASINWKEQAEKAR--KRMVSSENR-----LA---------------------------  476
AT2G04070.1           VTASMNWKEQAKKAR--KRLISSENG-----LA---------------------------  476
Sobic.001G481800.1    ITAMTNWQKMADKAR--DRVYEGSLP-----TQAD*------------------------  505
Sobic.003G260200.1    ITFFTDWPKMAEKAR--ERVFSDKAH-----ESAGP*-----------------------  489
Sobic.009G224800.1    ------------------------------------------------------------  408
Sobic.009G224800.2    ITFFSNWQKMSEKAR--ERVFSDEPS-----DKEPLESDG--------SNLF*-------  500
AT1G66760.2           VIGFTNWSKEAIKAR--ERIGDEKVW-----RHD----SL-----------LN-------  482
AT1G64820.1           VTFFTNWEQEATKAR--DRVFEMTPQ-----VKGNQKTQIIVEEDT--QVLLNHIAETV-  502
AT1G66780.1           VTFFTNWEQEVAKAR--DRVIEMIPQ-----EII--------------------------  485
Manes.02G072400.1     ITAFINWEKQVKRAR--ERILQRSSM-----ENDEN*-----------------------  477
Manes.01G113500.1     ITAFTNWQKQLQANMAKDRIFRGQFQ-----QVIDGIEL*--------------------  470
Manes.01G113500.2     ------------------------------------------------------------  378
AT1G15150.1           VTGCTNWKTQAREAR--ERMAVAHES-----ELTESELPI--------------------  487
AT1G15160.1           VTGCINWENQAREAR--KRMAVAHES-----ELTESELPF--------------------  487
AT1G15170.1           VTGCTNWESQADKAR--NRMALAYGT----------------------------------  481
AT1G15180.1           VTGCTNWESQADKAR--NRMALAYGT----------------------------------  482
AT1G71140.1           IVILTNWKKQARKAR--ERVMGDEYE-----EKESEEEHEYIS-----------------  485
Manes.06G063000.1     ITSCTNWEEQARKAR--ERVFERRSI-----SEDGSE*----------------------  479
Manes.14G109700.1     VIIYTDWEKQARKAR--ERIFHGRSS-----VENLLI*----------------------  488
Manes.14G109800.1     VTGCTDWERQASKAR--ERMFEGRSL-----VHNEVM*----------------------  489
Sobic.008G171600.1    LTARTNWDKEVEKAM--QRLQQTAVV-------PVNDV--IA*-----------------  497
Sobic.008G171600.2    LTARTNWDKEVEKAM--QRLQQTAVV-------PVNDV--IA*-----------------  511
Sobic.008G171600.3    LTARTNWDKEVEKAM--QRLQQTAVV-------PVNDV--IA*-----------------  497
AT3G59030.1           LTLKTNWTSEVENAA--QRVKTSATE-------NQEMA--NAGV----------------  507
Manes.01G182000.1     LTSRTNWDAEVEKAA--RRLNESARE-------GLESL--PNP*----------------  505
Manes.02G142000.1     LTSRTNWEAEVQKTV--ERLNESARQ-------ALESG--ANP*----------------  505
AT4G21903.2           VTYQADWDKEVMLHE--IKLKKRESDWICGTTKSLSRI--SNFVMK--------------  517
AT4G21910.4           VTYRTDWDKEVMLHE--IKWKKRGNVWICGTTRRSLSK--TSYNKF-------G--GVID  526
AT1G11670.1           VTFRTDWDKEVEKAS--RRLDQWEDT-----S-PLLKQ----------------------  503
AT1G61890.1           VTLRTDWDKEVEKAS--SRLDQWEES-----REPLLKQ----------------------  501
Manes.16G008000.1     VTIRTNWNEEVEKAR--MRLNCWENN-----EPQEAKI--*-------------------  504
Manes.17G038200.1     VTFRTDWNKEVEKAR--LRLDEWEDK-----KEPLLRT--SK*-----------------  507
Sobic.002G318300.1    ITLRTDWNKEVEEAR--KRLDKWDDT-----RQPLLAS--KE*-----------------  496
Manes.17G038300.1     VTYRTDWNKEVEIAR--SRLNMWDEN-----E-PLLEK--*-------------------  490
Manes.16G007900.1     VTYRTDWNKEVENAK--DRLDIWEEK-----KEPLLED--KRE-------------PLLD  507
Manes.17G038400.1     VTFRTDWNKEVENAK--NRLTMWDEK-----KEPLLED--KRE-------------ESEN  507
Sobic.001G012600.1    VTSRTDWNKEVIDVL--HPCHALLLMLVIHSMHAVFSD----------------------  503
Sobic.001G012600.2    VTSRTDWNKEVEKAR--ARLDKWD-----DKKQPLLED----------------------  498
Sobic.001G185600.1    VTLRTDWNKEVEQAR--MRLNKWEDK-----KKPLLAE--D-------------------  524
Sobic.001G185400.1    VTLRTDWNKEVEEAQ--KRLHKWEDK-----KTTEPLL--A-------------------  497
AT3G21690.1           VTFRTDWTKEVEEAS--KRLDKWSNK-----KQEVVPE----------------------  506
Sobic.001G185800.1    ITFRTDWDKEVEEAR--KRLNQWEDN-----KQPLLLV----------------------  498
Sobic.001G185500.1    VTYRTNWTKEVQEAQ--KRLNKWNDG-----KAPLLSA----------------------  514
Sobic.001G185500.2    VTYRTNWTKEVQEAQ--KRLNKWNDG-----KAPLLSA----------------------  397
Sobic.007G165500.2    VIWRRDWKSEAAQAS--SRVQKWGGK-----GTDEVKPLLQ*------------------  532
```

```
AT4G00350.1          MIYITNWNKEVEQAS--ERMKQWGAG-----YEKLEKIAT--------------------  542
Manes.01G255000.1    IICKTNWNKEVEEAS--ERMKTWGVQ-----DDSHN*-----------------------  600
Sobic.004G349550.1   LIWRTDWEAEAALAK--ERISAWSGE-----CRHVDVVKQQQGDT-G----SDS--DLKA  411
Sobic.004G349600.1   IVWTTDWKAEASLAL--ERVRIWGGA-----HHEKLGSS-QDDDA-------------VI  507
Sobic.007G176000.1   ITLRTNWDKEASEAH--SRIQKWGGS-----SPAAAKVSD------DC*-----------  508
Sobic.007G176100.1   ITLRTNWDKEAGEAH--SRIQKWGGS-----SAAARGEGT------L*------------  512
AT1G47530.1          IIYFTNWNKEAEQAE--SRVQRWGGT-----AQE--------------------------  484
Manes.05G164500.1    VTSITNWQKEAEEAE--SRVRKWGGS-----IAEDRSEW*--------------------  485
Manes.05G164500.2    VTSITNWQKEAEEAE--SRVRKWGGS-----IAEDRSEW*--------------------  485
Manes.18G030900.1    VTSVTNWKREAEEAE--SRVRKWGGS-----IAED*------------------------  482
AT1G23300.1          IIYRTNWKKEASLAE--ARIKKWGDQ-----SNKREEIDL------CEEDENNS--NGEN  511
AT3G26590.1          MICKTNWDTEASMAE--DRIREWGGE-----VSEIKQLIN--------------------  500
AT5G38030.1          MICRTNWDTEAAMAE--GRIREWGGE-----VSDQLL--N--------------------  498
Manes.12G023400.1    IIYRTNWNKEASVAE--DRIRKWGG------QIGSKDNNT------ETVTFRLE--M---  511
Manes.12G023500.1    IIYRTNWNKEASVAE--DRIKKWGGG-----QISSKENNI------GTVTFA*-------  509
Manes.12G023600.1    MIYRTNWNKEASIAG--DRIKRWGGH-----VDSE-EKNS------GKLSLSSN--RREM  531
Manes.13G025200.1    MIYRTNWNKEASIAE--DRIKRWGGH-----RLPR-E*----------------------  516
Manes.13G025200.2    MIYRTNWNKEASIAE--DRIKRWGGH-----RLPR-E*----------------------  515
Sobic.001G273100.1   IIFTTKWDKQAALAE--VRMAEWGGK-----NENLPLMETTHTADNHMAPAQEK--ILAH  516
Sobic.001G273100.2   IIFTTKWDKQAALAE--VRMAEWGGK-----NENLPLMETTHTADNHMAPAQEK--ILAH  516
Sobic.001G273000.2   VIFTTNWDKQAALTE--ERMAEWGGK-----E-KLPLMKSPHT-DDQMTPALEK--MLAQ  486
Sobic.001G273000.1   VIFTTNWDKQAALTE--ERMAEWGGK-----E-KLPLMKSPHT-DDQMTPALEK--MLAQ  497
Sobic.001G273000.3   VIFTTNWDKQAALTE--ERMAEWGGK-----E-KLPLMKSPHT-DDQMTPALEK--MLAQ  468
Manes.15G147800.1    ------------------------------------------------------------  428
Manes.15G147900.1    GFLRTNWQKEAAKAE--KHVRTWGGS-----QESQQ--SSSENIMNR*------------  491
Manes.17G098800.1    VFLRANWKKEALKAE--ERIRTWGGS-----ASVEPRQSSFEENMN*-------------  497
Manes.17G098900.1    VFLRANWKKEALKAE--ERIRTWGGS-------VEPRQSSFEENMN*-------------  495
Manes.17G098900.2    VFLRANWKKEALKAE--ERIRTWGGS-------VEPRQSSFEENMN*-------------  402
Sobic.001G162400.1   ISYRTKWEKQAMRAE--ERVREWGGR-----SDALPSATQVAPAVKDADPSSNA--AAI*  493
Sobic.003G307600.2   ILSRTKWQKEAMLAE--ERIRVWGGN-----VEL-PQTQETRPSENTAATVS*-------  409
Sobic.002G232200.1   ITLRCDWNKEALKAG--NRVRQWSST-----K*---------------------------  486
Sobic.002G232500.1   ITLRCDWNEEALKAS--TRMRRWSNS-----K*---------------------------  482
Sobic.002G232600.1   ITLRCDWNEEALKAS--SRMRTWSSS-----K*---------------------------  477
Sobic.007G160700.1   LTVKCDWDEEARLAS--MRMQKWADD-----LK*--------------------------  495
Sobic.003G126200.2   ITVRCDWEKQAIIAS--TRMDKLSQV-----R*---------------------------  505
Sobic.001G476700.1   MTVRCDWEKEAMVAS--TRMDNMSEV-----R*---------------------------  502
Sobic.001G476700.2   MTVRCDWEKEAMVAS--TRMDNMSEV-----R*---------------------------  525
Manes.18G062800.1    ITMKSDWDKEAEKAR--ARVAKWSSP-----HPDDQPAEPARR*----------------  496
AT5G44050.1          ITMRCDWEKEAQNAK--VRVNKWSVS-----DARK-------------------------  491
AT5G10420.1          ITTRCDWDNEAHKSS--VRIKKWLVS-----DAGN-------------------------  489
AT5G65380.1          ITMRCDWEKEAQKAS--ARINKWSNT-----IK---------------------------  486
Manes.12G129000.1    ITIRCDWEKEAEKAA--LHLKKWSEV-----K*---------------------------  487
Manes.13G097900.1    ITIRCDWEKEAEKAA--LHLKKWSEV-----K*---------------------------  461
Sobic.005G020700.1   ITLKTDWEKQVAVAQ--ERLKRWYMQ-----ENRRLQGLSGNS*----------------  529
Sobic.008G019400.1   ITLRTDWEKQVVTAQ--ERLKKWYME-----ENRRLQASRRNP*----------------  525
Manes.09G135300.1    VTCKTDWEKQVSLAR--NRINKWYVA-----DSEEQPNTQP*------------------  486
AT1G33080.1          MTMRTDWDQQVSSSL--KRLNRWVEP-----ESPSRNQTLQNE-----------------  494
AT1G33090.1          MTLRTDWDQQVSTSL--KNINRWVVP-----ESRDANQISSEE-----------------  494
AT1G33100.1          MTLRTDWDQQVSTSL--RNINRWVVP-----ESRDANQISSEE-----------------  491
AT1G33110.1          MTLRTDWDQQVSTSL--RRLNRWVVP-----ESRDVNQVSSEE-----------------  494
AT3G03620.1          IIYKTDWELEVKRTC--ERMKVWSLK-----PSNEESNPIIREESR--------------  498
AT5G17700.1          VIYKTDWELEVKKTN--ERMKTWTLN-----LPAVQSTTISTRDEE--------------  495
Manes.09G135200.1    LIWNTNWDEQVKIAS--ERLNRWFLK-----NP*--------------------------  476
Manes.09G134800.1    ITFRTDWDEQVKKAT--ERLNIFLKP-----S*---------------------------  472
Manes.08G150400.1    FTSKTNWDEQVIKVS--KHLDRWLLS-----KSEESNNENSI*-----------------  480
Manes.09G134900.1    ITSTTNWDEQVRKAS--ERLDRWLLR-----HSEESSNGNSIRERL--------------  485
Manes.09G135000.1    ITSTTNWDEQVKKAS--ERLDCWFLR-----PSEESSNGNSIQEIL--------------  485
Manes.09G134900.2    ITSTTNWDEQVRKAS--ERLDRWLLR-----HSEESSNGNSIRERL--------------  456
Manes.09G135000.2    ITSTTNWDEQVKKAS--ERLDCWFLR-----PSEESSNGNSIQEIL--------------  456


Sobic.002G099300.1   ------------------------------------------------------------  100
AT4G39030.1          ------------------------------------------------------------  543
Manes.04G084700.1    ------------------------------------------------------------  563
Manes.11G091900.1    ------------------------------------------------------------  557
Sobic.004G019800.2   ------------------------------------------------------------  424
Sobic.004G019800.3   ------------------------------------------------------------  424
Sobic.004G019800.4   ------------------------------------------------------------  409
AT2G21340.1          ------------------------------------------------------------  559
Manes.11G092000.1    ------------------------------------------------------------  558
Sobic.001G476700.3   ------------------------------------------------------------  164
```

```
Sobic.009G077000.1      ------------------------------------------------------------  80
Sobic.001G454900.1      ------------------------------------------------------------  565
Sobic.003G403000.1      ------------------------------------------------------------  600
Sobic.003G403000.2      ------------------------------------------------------------  600
Sobic.003G403000.3      ------------------------------------------------------------  600
Sobic.007G020600.1      ------------------------------------------------------------  525
Sobic.007G020600.2      ------------------------------------------------------------  486
AT3G08040.1             ------------------------------------------------------------  526
Manes.09G027700.1       ------------------------------------------------------------  211
Manes.09G027800.1       ------------------------------------------------------------  57
Manes.09G027900.1       ------------------------------------------------------------  57
Manes.09G026900.1       ------------------------------------------------------------  508
Manes.09G026900.2       ------------------------------------------------------------  508
Manes.09G027000.1       ------------------------------------------------------------  162
Manes.07G006000.1       ------------------------------------------------------------  547
Manes.07G006000.2       ------------------------------------------------------------  547
Manes.10G143000.1       ------------------------------------------------------------  540
AT1G51340.2             ------------------------------------------------------------  515
Manes.06G164500.1       ------------------------------------------------------------  509
Manes.06G164500.2       ------------------------------------------------------------  509
Manes.06G164500.3       ------------------------------------------------------------  509
Manes.06G164500.4       ------------------------------------------------------------  509
Manes.06G164500.5       ------------------------------------------------------------  509
Manes.14G002600.1       ------------------------------------------------------------  493
Sobic.008G006100.1      ------------------------------------------------------------  553
Sobic.005G005400.1      ------------------------------------------------------------  542
Sobic.005G005400.3      ------------------------------------------------------------  542
Sobic.005G005400.2      ------------------------------------------------------------  527
AT2G38330.1             ------------------------------------------------------------  521
Manes.08G096600.1       ----------SCTHA------SKFLCQAT*------------------------------  568
Manes.08G096600.2       ------------------------------------------------------------  551
AT4G38380.1             ------------------------------------------------------------  560
Sobic.002G286800.1      ------------------------------------------------------------  583
Sobic.003G149300.2      ------------------------------------------------------------  489
Sobic.003G149300.1      ------------------------------------------------------------  543
Sobic.003G149300.3      ------------------------------------------------------------  543
Sobic.003G149300.4      ------------------------------------------------------------  419
Manes.01G153400.1       ------------------------------------------------------------  274
Manes.04G064900.1       ------------------------------------------------------------  576
Manes.04G064900.2       ------------------------------------------------------------  576
Sobic.001G003700.1      ----------AC*-----------------------------------------------  484
AT4G22790.1             ------------------------------------------------------------  491
Manes.02G032300.1       ------------------------------------------------------------  508
Manes.10G000400.1       ---------------------------------DLEA--NLLYSTN*-------------  501
AT2G38510.1             ---------------------------------DLEI--GLLQNTN--------------  486
Manes.08G172300.1       ------------------------------------------------------------  496
Manes.09G117000.1       ------------------------------------------------------------  466
Sobic.010G167800.1      GVV-----ERSCYDHE-PLISNSGEDDPETV*----------------------------  559
AT5G19700.1             --------------SE-PLIYVVTVATD--------------------------------  508
AT4G29140.1             --------------AE-PLIRITVLY----------------------------------  532
Manes.03G026500.1       --------------EH-RLIPITVISP*--------------------------------  535
Manes.16G109300.1       ------------------------------------------------------------  512
Sobic.007G181100.1      TVLT*-------------------------------------------------------  532
Manes.13G127800.1       VTLVD------*------------------------------------------------  524
Sobic.001G446800.1      TVIATSNSAAGCKKNNGGYVPISESCSNDSELEKLEE--GLMTSDDIPSASVSGSACGGD  578
AT5G52050.1             ------------------------------------------------------------  505
AT1G58340.1             DMMRT-MLV---------------------------------------------------  532
Manes.05G186700.1       ANLEE-ILCINDEPVKSTSLETDPLI----------------------------------  533
Manes.18G054300.1       PQLEE-ILSIDHELVKSSSLETDPLL----------------------------------  544
Sobic.001G320900.1      TGPHA-KVAASHGDED-SSLLITVQG----------------------------------  569
Sobic.001G320900.2      TGPHA-KVAASHGDED-SSLLITVQG----------------------------------  418
Sobic.004G283500.1      -------VTAAGGDEE-LGLPIDVVI----------------------------------  560
Sobic.006G184400.1      -------LLL--GDTD-M-----EKA----------------------------------  560
AT4G23030.1             LDIEE-NLVF--------------------------------------------------  502
Manes.01G067000.1       --------LSTSSDHN-D------------------------------------------  525
Manes.02G027800.1       ------------------------------------------------------------  520
Manes.06G143600.1       PEIKE-DSLYNCGDSD-Q------------------------------------------  545
Manes.14G029100.1       PEIKV-DSLYYFGDQS--------------------------------------------  544
AT5G49130.1             ------------------------------------------------------------  502
Manes.03G198000.1       EEEEE-----GVG-----FLR----LKIEGAVD*--------------------------  514
Manes.15G011000.1       DEEFQ-----GVG-----FLG----LKIEYGVVDFEK--*--------------------  517
```

```
Sobic.001G019700.1      RLVVA-----GTG-------E----PAAEGGV*---------------------------  529
AT1G71870.1             DVL---------------------------------------------------------  510
Manes.02G189800.1       RLLVN-----GNG-------N----IP*--------------------------------  503
Manes.18G098800.1       RLLVN-----GKH-------I----RLIFSDY*---------------------------  502
Sobic.010G256932.1      ------------------------------------------------------------  411
Sobic.010G256700.1      ------------------------------------------------------------  461
Sobic.010G256700.5      ------------------------------------------------------------  461
Sobic.010G256700.6      ------------------------------------------------------------  461
Sobic.010G256700.4      ------------------------------------------------------------  483
Sobic.009G106900.1      ------------------------------------------------------------  85
Manes.S031500.1         ------------------------------------------------------------  487
Manes.14G109900.1       ------------------------------------------------------------  498
Manes.14G109900.2       ------------------------------------------------------------  397
Manes.14G109900.3       ------------------------------------------------------------  303
AT1G73700.1             ------------------------------------------------------------  476
AT2G34360.1             ------------------------------------------------------------  480
AT5G52450.1             ------------------------------------------------------------  486
Sobic.009G106960.1      ------------------------------------------------------------  67
Manes.14G060800.1       ------------------------------------------------------------  492
Sobic.009G106800.1      ------------------------------------------------------------  450
Sobic.007G074300.2      ------------------------------------------------------------  297
Sobic.009G106700.1      ------------------------------------------------------------  474
Sobic.010G138400.4      ------------------------------------------------------------  401
Sobic.010G138400.1      ------------------------------------------------------------  479
Sobic.010G138400.5      ------------------------------------------------------------  398
Sobic.004G129900.1      GAIE-----RTNGLNK-----------------------G*-------------------  511
Sobic.004G129900.2      GAIE-----RTNGLNK-----------------------G*-------------------  429
Sobic.006G042200.1      ------------------------------------------------------------  480
Sobic.004G129700.1      ------------------------------------------------------------  487
Sobic.004G129800.1      ------------------------------------------------------------  480
Sobic.004G129800.2      ------------------------------------------------------------  395
AT3G23550.1             ------------------------------------------------------------  469
AT3G23560.1             ------------------------------------------------------------  477
Manes.15G088100.1       ------------------------------------------------------------  499
Manes.03G109100.1       ------------------------------------------------------------  482
Manes.03G109100.2       ------------------------------------------------------------  443
Sobic.002G311200.1      ------------------------------------------------------------  592
Sobic.002G006500.3      ------------------------------------------------------------  506
Sobic.002G006500.4      ------------------------------------------------------------  487
AT2G04090.1             ------------------------------------------------------------  477
AT2G04100.1             ------------------------------------------------------------  483
AT2G04066.1             ------------------------------------------------------------  171
AT2G04040.1             ------------------------------------------------------------  476
AT2G04080.1             ------------------------------------------------------------  476
AT2G04050.1             ------------------------------------------------------------  476
AT2G04070.1             ------------------------------------------------------------  476
Sobic.001G481800.1      ------------------------------------------------------------  505
Sobic.003G260200.1      ------------------------------------------------------------  489
Sobic.009G224800.1      ------------------------------------------------------------  408
Sobic.009G224800.2      ------------------------------------------------------------  500
AT1G66760.2             ------------------------------------------------------------  482
AT1G64820.1             ------------------------------------------------------------  502
AT1G66780.1             ------------------------------------------------------------  485
Manes.02G072400.1       ------------------------------------------------------------  477
Manes.01G113500.1       ------------------------------------------------------------  470
Manes.01G113500.2       ------------------------------------------------------------  378
AT1G15150.1             ------------------------------------------------------------  487
AT1G15160.1             ------------------------------------------------------------  487
AT1G15170.1             ------------------------------------------------------------  481
AT1G15180.1             ------------------------------------------------------------  482
AT1G71140.1             ------------------------------------------------------------  485
Manes.06G063000.1       ------------------------------------------------------------  479
Manes.14G109700.1       ------------------------------------------------------------  488
Manes.14G109800.1       ------------------------------------------------------------  489
Sobic.008G171600.1      ------------------------------------------------------------  497
Sobic.008G171600.2      ------------------------------------------------------------  511
Sobic.008G171600.3      ------------------------------------------------------------  497
AT3G59030.1             ------------------------------------------------------------  507
Manes.01G182000.1       ------------------------------------------------------------  505
Manes.02G142000.1       ------------------------------------------------------------  505
AT4G21903.2             ------------------------------------------------------------  517
```

```
AT4G21910.4          ----------NKEKE------ISVVCAGDFNVRSFSWLLCLFYGNR----TIKIYFCLMD  566
AT1G11670.1          ------------------------------------------------------------  503
AT1G61890.1          ------------------------------------------------------------  501
Manes.16G008000.1    ------------------------------------------------------------  504
Manes.17G038200.1    ------------------------------------------------------------  507
Sobic.002G318300.1   ------------------------------------------------------------  496
Manes.17G038300.1    ------------------------------------------------------------  490
Manes.16G007900.1    ----------DRRDE------SE-------N*----------------------------  515
Manes.17G038400.1    -------------------------------*----------------------------  507
Sobic.001G012600.1   ---------------------DD-------*-----------------------------  505
Sobic.001G012600.2   ---------------------*--------------------------------------  498
Sobic.001G185600.1   ------------*-----------------------------------------------  524
Sobic.001G185400.1   ------------GVG------NG-------N*----------------------------  503
AT3G21690.1          ------------------------------------------------------------  506
Sobic.001G185800.1   -------------PS------D*-------------------------------------  501
Sobic.001G185500.1   -------------QE------*--------------------------------------  516
Sobic.001G185500.2   -------------QE------*--------------------------------------  399
Sobic.007G165500.2   ------------------------------------------------------------  532
AT4G00350.1          ------------------------------------------------------------  542
Manes.01G255000.1    ------------------------------------------------------------  600
Sobic.004G349550.1   ----------NLRV*---------------------------------------------  415
Sobic.004G349600.1   ----------*-------------------------------------------------  507
Sobic.007G176000.1   ------------------------------------------------------------  508
Sobic.007G176100.1   ------------------------------------------------------------  512
AT1G47530.1          ------------------------------------------------------------  484
Manes.05G164500.1    ------------------------------------------------------------  485
Manes.05G164500.2    ------------------------------------------------------------  485
Manes.18G030900.1    ------------------------------------------------------------  482
AT1G23300.1          ----------NHRK----------------------------------------------  515
AT3G26590.1          ------------------------------------------------------------  500
AT5G38030.1          ------------------------------------------------------------  498
Manes.12G023400.1    ----------*-------------------------------------------------  511
Manes.12G023500.1    ------------------------------------------------------------  509
Manes.12G023600.1    ----------DSDSL------VEL*-----------------------------------  539
Manes.13G025200.1    ------------------------------------------------------------  516
Manes.13G025200.2    ------------------------------------------------------------  515
Sobic.001G273100.1   ----------DSQKN------VELVRTD*-------------------------------  528
Sobic.001G273100.2   ----------DSQKN------VELVRTD*-------------------------------  528
Sobic.001G273000.2   ----------DSKKN------VDLLCTE*-------------------------------  498
Sobic.001G273000.1   ----------DSKKN------VDLLCTE*-------------------------------  509
Sobic.001G273000.3   ----------DSKKN------VDLLCTE*-------------------------------  480
Manes.15G147800.1    ------------------------------------------------------------  428
Manes.15G147900.1    ------------------------------------------------------------  491
Manes.17G098800.1    ------------------------------------------------------------  497
Manes.17G098900.1    ------------------------------------------------------------  495
Manes.17G098900.2    ------------------------------------------------------------  402
Sobic.001G162400.1   ------------------------------------------------------------  493
Sobic.003G307600.2   ------------------------------------------------------------  409
Sobic.002G232200.1   ------------------------------------------------------------  486
Sobic.002G232500.1   ------------------------------------------------------------  482
Sobic.002G232600.1   ------------------------------------------------------------  477
Sobic.007G160700.1   ------------------------------------------------------------  495
Sobic.003G126200.2   ------------------------------------------------------------  505
Sobic.001G476700.1   ------------------------------------------------------------  502
Sobic.001G476700.2   ------------------------------------------------------------  525
Manes.18G062800.1    ------------------------------------------------------------  496
AT5G44050.1          ------------------------------------------------------------  491
AT5G10420.1          ------------------------------------------------------------  489
AT5G65380.1          ------------------------------------------------------------  486
Manes.12G129000.1    ------------------------------------------------------------  487
Manes.13G097900.1    ------------------------------------------------------------  461
Sobic.005G020700.1   ------------------------------------------------------------  529
Sobic.008G019400.1   ------------------------------------------------------------  525
Manes.09G135300.1    ------------------------------------------------------------  486
AT1G33080.1          ------------------------------------------------------------  494
AT1G33090.1          ------------------------------------------------------------  494
AT1G33100.1          ------------------------------------------------------------  491
AT1G33110.1          ------------------------------------------------------------  494
AT3G03620.1          ------------SK----------------------------------------------  500
AT5G17700.1          ------------RK----------------------------------------------  497
Manes.09G135200.1    ------------------------------------------------------------  476
```

```
Manes.09G134800.1      ------------------------------------------------------------ 472
Manes.08G150400.1      ------------------------------------------------------------ 480
Manes.09G134900.1      ------------KS*--------------------------------------------- 487
Manes.09G135000.1      ------------NG*--------------------------------------------- 487
Manes.09G134900.2      ------------KS*--------------------------------------------- 458
Manes.09G135000.2      ------------NG*--------------------------------------------- 458


Sobic.002G099300.1     ------------------------------------------------------------ 100
AT4G39030.1            ------------------------------------------------------------ 543
Manes.04G084700.1      ------------------------------------------------------------ 563
Manes.11G091900.1      ------------------------------------------------------------ 557
Sobic.004G019800.2     ------------------------------------------------------------ 424
Sobic.004G019800.3     ------------------------------------------------------------ 424
Sobic.004G019800.4     ------------------------------------------------------------ 409
AT2G21340.1            ------------------------------------------------------------ 559
Manes.11G092000.1      ------------------------------------------------------------ 558
Sobic.001G476700.3     ------------------------------------------------------------ 164
Sobic.009G077000.1     ------------------------------------------------------------ 80
Sobic.001G454900.1     ------------------------------------------------------------ 565
Sobic.003G403000.1     ------------------------------------------------------------ 600
Sobic.003G403000.2     ------------------------------------------------------------ 600
Sobic.003G403000.3     ------------------------------------------------------------ 600
Sobic.007G020600.1     ------------------------------------------------------------ 525
Sobic.007G020600.2     ------------------------------------------------------------ 486
AT3G08040.1            ------------------------------------------------------------ 526
Manes.09G027700.1      ------------------------------------------------------------ 211
Manes.09G027800.1      ------------------------------------------------------------ 57
Manes.09G027900.1      ------------------------------------------------------------ 57
Manes.09G026900.1      ------------------------------------------------------------ 508
Manes.09G026900.2      ------------------------------------------------------------ 508
Manes.09G027000.1      ------------------------------------------------------------ 162
Manes.07G006000.1      ------------------------------------------------------------ 547
Manes.07G006000.2      ------------------------------------------------------------ 547
Manes.10G143000.1      ------------------------------------------------------------ 540
AT1G51340.2            ------------------------------------------------------------ 515
Manes.06G164500.1      ------------------------------------------------------------ 509
Manes.06G164500.2      ------------------------------------------------------------ 509
Manes.06G164500.3      ------------------------------------------------------------ 509
Manes.06G164500.4      ------------------------------------------------------------ 509
Manes.06G164500.5      ------------------------------------------------------------ 509
Manes.14G002600.1      ------------------------------------------------------------ 493
Sobic.008G006100.1     ------------------------------------------------------------ 553
Sobic.005G005400.1     ------------------------------------------------------------ 542
Sobic.005G005400.3     ------------------------------------------------------------ 542
Sobic.005G005400.2     ------------------------------------------------------------ 527
AT2G38330.1            ------------------------------------------------------------ 521
Manes.08G096600.1      ------------------------------------------------------------ 568
Manes.08G096600.2      ------------------------------------------------------------ 551
AT4G38380.1            ------------------------------------------------------------ 560
Sobic.002G286800.1     ------------------------------------------------------------ 583
Sobic.003G149300.2     ------------------------------------------------------------ 489
Sobic.003G149300.1     ------------------------------------------------------------ 543
Sobic.003G149300.3     ------------------------------------------------------------ 543
Sobic.003G149300.4     ------------------------------------------------------------ 419
Manes.01G153400.1      ------------------------------------------------------------ 274
Manes.04G064900.1      ------------------------------------------------------------ 576
Manes.04G064900.2      ------------------------------------------------------------ 576
Sobic.001G003700.1     ------------------------------------------------------------ 484
AT4G22790.1            ------------------------------------------------------------ 491
Manes.02G032300.1      ------------------------------------------------------------ 508
Manes.10G000400.1      ------------------------------------------------------------ 501
AT2G38510.1            ------------------------------------------------------------ 486
Manes.08G172300.1      ------------------------------------------------------------ 496
Manes.09G117000.1      ------------------------------------------------------------ 466
Sobic.010G167800.1     ------------------------------------------------------------ 559
AT5G19700.1            ------------------------------------------------------------ 508
AT4G29140.1            ------------------------------------------------------------ 532
Manes.03G026500.1      ------------------------------------------------------------ 535
Manes.16G109300.1      ------------------------------------------------------------ 512
Sobic.007G181100.1     ------------------------------------------------------------ 532
```

```
Manes.13G127800.1       ------------------------------------------------------------  524
Sobic.001G446800.1      TDAVVREQNRGSGNSCNDSGAAGTAATEGKEQRKGGSEERGPLISVSDGEHDGDSRGGGG  638
AT5G52050.1             ------------------------------------------------------------  505
AT1G58340.1             ------------------------------------------------------------  532
Manes.05G186700.1       ---------STASTVH*-------------------------------------------  540
Manes.18G054300.1       ---------STKRTVH*-------------------------------------------  551
Sobic.001G320900.1      ---------*--------------------------------------------------  569
Sobic.001G320900.2      ---------*--------------------------------------------------  418
Sobic.004G283500.1      ---------ERPKDQC*-------------------------------------------  567
Sobic.006G184400.1      ---------NAHSDQC*-------------------------------------------  567
AT4G23030.1             ------------------------------------------------------------  502
Manes.01G067000.1       ----------SS---A*-------------------------------------------  528
Manes.02G027800.1       ---------------A*-------------------------------------------  521
Manes.06G143600.1       ----------SP---V*-------------------------------------------  548
Manes.14G029100.1       -----------P---V*-------------------------------------------  546
AT5G49130.1             ------------------------------------------------------------  502
Manes.03G198000.1       ------------------------------------------------------------  514
Manes.15G011000.1       ------------------------------------------------------------  517
Sobic.001G019700.1      ------------------------------------------------------------  529
AT1G71870.1             ------------------------------------------------------------  510
Manes.02G189800.1       ------------------------------------------------------------  503
Manes.18G098800.1       ------------------------------------------------------------  502
Sobic.010G256932.1      ------------------------------------------------------------  411
Sobic.010G256700.1      ------------------------------------------------------------  461
Sobic.010G256700.5      ------------------------------------------------------------  461
Sobic.010G256700.6      ------------------------------------------------------------  461
Sobic.010G256700.4      ------------------------------------------------------------  483
Sobic.009G106900.1      ------------------------------------------------------------  85
Manes.S031500.1         ------------------------------------------------------------  487
Manes.14G109900.1       ------------------------------------------------------------  498
Manes.14G109900.2       ------------------------------------------------------------  397
Manes.14G109900.3       ------------------------------------------------------------  303
AT1G73700.1             ------------------------------------------------------------  476
AT2G34360.1             ------------------------------------------------------------  480
AT5G52450.1             ------------------------------------------------------------  486
Sobic.009G106960.1      ------------------------------------------------------------  67
Manes.14G060800.1       ------------------------------------------------------------  492
Sobic.009G106800.1      ------------------------------------------------------------  450
Sobic.007G074300.2      ------------------------------------------------------------  297
Sobic.009G106700.1      ------------------------------------------------------------  474
Sobic.010G138400.4      ------------------------------------------------------------  401
Sobic.010G138400.1      ------------------------------------------------------------  479
Sobic.010G138400.5      ------------------------------------------------------------  398
Sobic.004G129900.1      ------------------------------------------------------------  511
Sobic.004G129900.2      ------------------------------------------------------------  429
Sobic.006G042200.1      ------------------------------------------------------------  480
Sobic.004G129700.1      ------------------------------------------------------------  487
Sobic.004G129800.1      ------------------------------------------------------------  480
Sobic.004G129800.2      ------------------------------------------------------------  395
AT3G23550.1             ------------------------------------------------------------  469
AT3G23560.1             ------------------------------------------------------------  477
Manes.15G088100.1       ------------------------------------------------------------  499
Manes.03G109100.1       ------------------------------------------------------------  482
Manes.03G109100.2       ------------------------------------------------------------  443
Sobic.002G311200.1      ------------------------------------------------------------  592
Sobic.002G006500.3      ------------------------------------------------------------  506
Sobic.002G006500.4      ------------------------------------------------------------  487
AT2G04090.1             ------------------------------------------------------------  477
AT2G04100.1             ------------------------------------------------------------  483
AT2G04066.1             ------------------------------------------------------------  171
AT2G04040.1             ------------------------------------------------------------  476
AT2G04080.1             ------------------------------------------------------------  476
AT2G04050.1             ------------------------------------------------------------  476
AT2G04070.1             ------------------------------------------------------------  476
Sobic.001G481800.1      ------------------------------------------------------------  505
Sobic.003G260200.1      ------------------------------------------------------------  489
Sobic.009G224800.1      ------------------------------------------------------------  408
Sobic.009G224800.2      ------------------------------------------------------------  500
AT1G66760.2             ------------------------------------------------------------  482
AT1G64820.1             ------------------------------------------------------------  502
AT1G66780.1             ------------------------------------------------------------  485
```

```
Manes.02G072400.1      ------------------------------------------------------------  477
Manes.01G113500.1      ------------------------------------------------------------  470
Manes.01G113500.2      ------------------------------------------------------------  378
AT1G15150.1            ------------------------------------------------------------  487
AT1G15160.1            ------------------------------------------------------------  487
AT1G15170.1            ------------------------------------------------------------  481
AT1G15180.1            ------------------------------------------------------------  482
AT1G71140.1            ------------------------------------------------------------  485
Manes.06G063000.1      ------------------------------------------------------------  479
Manes.14G109700.1      ------------------------------------------------------------  488
Manes.14G109800.1      ------------------------------------------------------------  489
Sobic.008G171600.1     ------------------------------------------------------------  497
Sobic.008G171600.2     ------------------------------------------------------------  511
Sobic.008G171600.3     ------------------------------------------------------------  497
AT3G59030.1            ------------------------------------------------------------  507
Manes.01G182000.1      ------------------------------------------------------------  505
Manes.02G142000.1      ------------------------------------------------------------  505
AT4G21903.2            ------------------------------------------------------------  517
AT4G21910.4            YSL--------FTKSCS-------------------------------------------  575
AT1G11670.1            ------------------------------------------------------------  503
AT1G61890.1            ------------------------------------------------------------  501
Manes.16G008000.1      ------------------------------------------------------------  504
Manes.17G038200.1      ------------------------------------------------------------  507
Sobic.002G318300.1     ------------------------------------------------------------  496
Manes.17G038300.1      ------------------------------------------------------------  490
Manes.16G007900.1      ------------------------------------------------------------  515
Manes.17G038400.1      ------------------------------------------------------------  507
Sobic.001G012600.1     ------------------------------------------------------------  505
Sobic.001G012600.2     ------------------------------------------------------------  498
Sobic.001G185600.1     ------------------------------------------------------------  524
Sobic.001G185400.1     ------------------------------------------------------------  503
AT3G21690.1            ------------------------------------------------------------  506
Sobic.001G185800.1     ------------------------------------------------------------  501
Sobic.001G185500.1     ------------------------------------------------------------  516
Sobic.001G185500.2     ------------------------------------------------------------  399
Sobic.007G165500.2     ------------------------------------------------------------  532
AT4G00350.1            ------------------------------------------------------------  542
Manes.01G255000.1      ------------------------------------------------------------  600
Sobic.004G349550.1     ------------------------------------------------------------  415
Sobic.004G349600.1     ------------------------------------------------------------  507
Sobic.007G176000.1     ------------------------------------------------------------  508
Sobic.007G176100.1     ------------------------------------------------------------  512
AT1G47530.1            ------------------------------------------------------------  484
Manes.05G164500.1      ------------------------------------------------------------  485
Manes.05G164500.2      ------------------------------------------------------------  485
Manes.18G030900.1      ------------------------------------------------------------  482
AT1G23300.1            ------------------------------------------------------------  515
AT3G26590.1            ------------------------------------------------------------  500
AT5G38030.1            ------------------------------------------------------------  498
Manes.12G023400.1      ------------------------------------------------------------  511
Manes.12G023500.1      ------------------------------------------------------------  509
Manes.12G023600.1      ------------------------------------------------------------  539
Manes.13G025200.1      ------------------------------------------------------------  516
Manes.13G025200.2      ------------------------------------------------------------  515
Sobic.001G273100.1     ------------------------------------------------------------  528
Sobic.001G273100.2     ------------------------------------------------------------  528
Sobic.001G273000.2     ------------------------------------------------------------  498
Sobic.001G273000.1     ------------------------------------------------------------  509
Sobic.001G273000.3     ------------------------------------------------------------  480
Manes.15G147800.1      ------------------------------------------------------------  428
Manes.15G147900.1      ------------------------------------------------------------  491
Manes.17G098800.1      ------------------------------------------------------------  497
Manes.17G098900.1      ------------------------------------------------------------  495
Manes.17G098900.2      ------------------------------------------------------------  402
Sobic.001G162400.1     ------------------------------------------------------------  493
Sobic.003G307600.2     ------------------------------------------------------------  409
Sobic.002G232200.1     ------------------------------------------------------------  486
Sobic.002G232500.1     ------------------------------------------------------------  482
Sobic.002G232600.1     ------------------------------------------------------------  477
Sobic.007G160700.1     ------------------------------------------------------------  495
Sobic.003G126200.2     ------------------------------------------------------------  505
```

```
Sobic.001G476700.1      ------------------------------------------------------------  502
Sobic.001G476700.2      ------------------------------------------------------------  525
Manes.18G062800.1       ------------------------------------------------------------  496
AT5G44050.1             ------------------------------------------------------------  491
AT5G10420.1             ------------------------------------------------------------  489
AT5G65380.1             ------------------------------------------------------------  486
Manes.12G129000.1       ------------------------------------------------------------  487
Manes.13G097900.1       ------------------------------------------------------------  461
Sobic.005G020700.1      ------------------------------------------------------------  529
Sobic.008G019400.1      ------------------------------------------------------------  525
Manes.09G135300.1       ------------------------------------------------------------  486
AT1G33080.1             ------------------------------------------------------------  494
AT1G33090.1             ------------------------------------------------------------  494
AT1G33100.1             ------------------------------------------------------------  491
AT1G33110.1             ------------------------------------------------------------  494
AT3G03620.1             ------------------------------------------------------------  500
AT5G17700.1             ------------------------------------------------------------  497
Manes.09G135200.1       ------------------------------------------------------------  476
Manes.09G134800.1       ------------------------------------------------------------  472
Manes.08G150400.1       ------------------------------------------------------------  480
Manes.09G134900.1       ------------------------------------------------------------  487
Manes.09G135000.1       ------------------------------------------------------------  487
Manes.09G134900.2       ------------------------------------------------------------  458
Manes.09G135000.2       ------------------------------------------------------------  458


Sobic.002G099300.1      ---  100
AT4G39030.1             ---  543
Manes.04G084700.1       ---  563
Manes.11G091900.1       ---  557
Sobic.004G019800.2      ---  424
Sobic.004G019800.3      ---  424
Sobic.004G019800.4      ---  409
AT2G21340.1             ---  559
Manes.11G092000.1       ---  558
Sobic.001G476700.3      ---  164
Sobic.009G077000.1      ---  80
Sobic.001G454900.1      ---  565
Sobic.003G403000.1      ---  600
Sobic.003G403000.2      ---  600
Sobic.003G403000.3      ---  600
Sobic.007G020600.1      ---  525
Sobic.007G020600.2      ---  486
AT3G08040.1             ---  526
Manes.09G027700.1       ---  211
Manes.09G027800.1       ---  57
Manes.09G027900.1       ---  57
Manes.09G026900.1       ---  508
Manes.09G026900.2       ---  508
Manes.09G027000.1       ---  162
Manes.07G006000.1       ---  547
Manes.07G006000.2       ---  547
Manes.10G143000.1       ---  540
AT1G51340.2             ---  515
Manes.06G164500.1       ---  509
Manes.06G164500.2       ---  509
Manes.06G164500.3       ---  509
Manes.06G164500.4       ---  509
Manes.06G164500.5       ---  509
Manes.14G002600.1       ---  493
Sobic.008G006100.1      ---  553
Sobic.005G005400.1      ---  542
Sobic.005G005400.3      ---  542
Sobic.005G005400.2      ---  527
AT2G38330.1             ---  521
Manes.08G096600.1       ---  568
Manes.08G096600.2       ---  551
AT4G38380.1             ---  560
Sobic.002G286800.1      ---  583
Sobic.003G149300.2      ---  489
Sobic.003G149300.1      ---  543
```

| | | |
|---|---|---|
| Sobic.003G149300.3 | --- | 543 |
| Sobic.003G149300.4 | --- | 419 |
| Manes.01G153400.1 | --- | 274 |
| Manes.04G064900.1 | --- | 576 |
| Manes.04G064900.2 | --- | 576 |
| Sobic.001G003700.1 | --- | 484 |
| AT4G22790.1 | --- | 491 |
| Manes.02G032300.1 | --- | 508 |
| Manes.10G000400.1 | --- | 501 |
| AT2G38510.1 | --- | 486 |
| Manes.08G172300.1 | --- | 496 |
| Manes.09G117000.1 | --- | 466 |
| Sobic.010G167800.1 | --- | 559 |
| AT5G19700.1 | --- | 508 |
| AT4G29140.1 | --- | 532 |
| Manes.03G026500.1 | --- | 535 |
| Manes.16G109300.1 | --- | 512 |
| Sobic.007G181100.1 | --- | 532 |
| Manes.13G127800.1 | --- | 524 |
| Sobic.001G446800.1 | QV* | 640 |
| AT5G52050.1 | --- | 505 |
| AT1G58340.1 | --- | 532 |
| Manes.05G186700.1 | --- | 540 |
| Manes.18G054300.1 | --- | 551 |
| Sobic.001G320900.1 | --- | 569 |
| Sobic.001G320900.2 | --- | 418 |
| Sobic.004G283500.1 | --- | 567 |
| Sobic.006G184400.1 | --- | 567 |
| AT4G23030.1 | --- | 502 |
| Manes.01G067000.1 | --- | 528 |
| Manes.02G027800.1 | --- | 521 |
| Manes.06G143600.1 | --- | 548 |
| Manes.14G029100.1 | --- | 546 |
| AT5G49130.1 | --- | 502 |
| Manes.03G198000.1 | --- | 514 |
| Manes.15G011000.1 | --- | 517 |
| Sobic.001G019700.1 | --- | 529 |
| AT1G71870.1 | --- | 510 |
| Manes.02G189800.1 | --- | 503 |
| Manes.18G098800.1 | --- | 502 |
| Sobic.010G256932.1 | --- | 411 |
| Sobic.010G256700.1 | --- | 461 |
| Sobic.010G256700.5 | --- | 461 |
| Sobic.010G256700.6 | --- | 461 |
| Sobic.010G256700.4 | --- | 483 |
| Sobic.009G106900.1 | --- | 85 |
| Manes.S031500.1 | --- | 487 |
| Manes.14G109900.1 | --- | 498 |
| Manes.14G109900.2 | --- | 397 |
| Manes.14G109900.3 | --- | 303 |
| AT1G73700.1 | --- | 476 |
| AT2G34360.1 | --- | 480 |
| AT5G52450.1 | --- | 486 |
| Sobic.009G106960.1 | --- | 67 |
| Manes.14G060800.1 | --- | 492 |
| Sobic.009G106800.1 | --- | 450 |
| Sobic.007G074300.2 | --- | 297 |
| Sobic.009G106700.1 | --- | 474 |
| Sobic.010G138400.4 | --- | 401 |
| Sobic.010G138400.1 | --- | 479 |
| Sobic.010G138400.5 | --- | 398 |
| Sobic.004G129900.1 | --- | 511 |
| Sobic.004G129900.2 | --- | 429 |
| Sobic.006G042200.1 | --- | 480 |
| Sobic.004G129700.1 | --- | 487 |
| Sobic.004G129800.1 | --- | 480 |
| Sobic.004G129800.2 | --- | 395 |
| AT3G23550.1 | --- | 469 |
| AT3G23560.1 | --- | 477 |
| Manes.15G088100.1 | --- | 499 |
| Manes.03G109100.1 | --- | 482 |

| | | |
|---|---|---|
| Manes.03G109100.2 | --- | 443 |
| Sobic.002G311200.1 | --- | 592 |
| Sobic.002G006500.3 | --- | 506 |
| Sobic.002G006500.4 | --- | 487 |
| AT2G04090.1 | --- | 477 |
| AT2G04100.1 | --- | 483 |
| AT2G04066.1 | --- | 171 |
| AT2G04040.1 | --- | 476 |
| AT2G04080.1 | --- | 476 |
| AT2G04050.1 | --- | 476 |
| AT2G04070.1 | --- | 476 |
| Sobic.001G481800.1 | --- | 505 |
| Sobic.003G260200.1 | --- | 489 |
| Sobic.009G224800.1 | --- | 408 |
| Sobic.009G224800.2 | --- | 500 |
| AT1G66760.2 | --- | 482 |
| AT1G64820.1 | --- | 502 |
| AT1G66780.1 | --- | 485 |
| Manes.02G072400.1 | --- | 477 |
| Manes.01G113500.1 | --- | 470 |
| Manes.01G113500.2 | --- | 378 |
| AT1G15150.1 | --- | 487 |
| AT1G15160.1 | --- | 487 |
| AT1G15170.1 | --- | 481 |
| AT1G15180.1 | --- | 482 |
| AT1G71140.1 | --- | 485 |
| Manes.06G063000.1 | --- | 479 |
| Manes.14G109700.1 | --- | 488 |
| Manes.14G109800.1 | --- | 489 |
| Sobic.008G171600.1 | --- | 497 |
| Sobic.008G171600.2 | --- | 511 |
| Sobic.008G171600.3 | --- | 497 |
| AT3G59030.1 | --- | 507 |
| Manes.01G182000.1 | --- | 505 |
| Manes.02G142000.1 | --- | 505 |
| AT4G21903.2 | --- | 517 |
| AT4G21910.4 | --- | 575 |
| AT1G11670.1 | --- | 503 |
| AT1G61890.1 | --- | 501 |
| Manes.16G008000.1 | --- | 504 |
| Manes.17G038200.1 | --- | 507 |
| Sobic.002G318300.1 | --- | 496 |
| Manes.17G038300.1 | --- | 490 |
| Manes.16G007900.1 | --- | 515 |
| Manes.17G038400.1 | --- | 507 |
| Sobic.001G012600.1 | --- | 505 |
| Sobic.001G012600.2 | --- | 498 |
| Sobic.001G185600.1 | --- | 524 |
| Sobic.001G185400.1 | --- | 503 |
| AT3G21690.1 | --- | 506 |
| Sobic.001G185800.1 | --- | 501 |
| Sobic.001G185500.1 | --- | 516 |
| Sobic.001G185500.2 | --- | 399 |
| Sobic.007G165500.2 | --- | 532 |
| AT4G00350.1 | --- | 542 |
| Manes.01G255000.1 | --- | 600 |
| Sobic.004G349550.1 | --- | 415 |
| Sobic.004G349600.1 | --- | 507 |
| Sobic.007G176000.1 | --- | 508 |
| Sobic.007G176100.1 | --- | 512 |
| AT1G47530.1 | --- | 484 |
| Manes.05G164500.1 | --- | 485 |
| Manes.05G164500.2 | --- | 485 |
| Manes.18G030900.1 | --- | 482 |
| AT1G23300.1 | --- | 515 |
| AT3G26590.1 | --- | 500 |
| AT5G38030.1 | --- | 498 |
| Manes.12G023400.1 | --- | 511 |
| Manes.12G023500.1 | --- | 509 |
| Manes.12G023600.1 | --- | 539 |
| Manes.13G025200.1 | --- | 516 |

| | | |
|---|---|---|
| Manes.13G025200.2 | --- | 515 |
| Sobic.001G273100.1 | --- | 528 |
| Sobic.001G273100.2 | --- | 528 |
| Sobic.001G273000.2 | --- | 498 |
| Sobic.001G273000.1 | --- | 509 |
| Sobic.001G273000.3 | --- | 480 |
| Manes.15G147800.1 | --- | 428 |
| Manes.15G147900.1 | --- | 491 |
| Manes.17G098800.1 | --- | 497 |
| Manes.17G098900.1 | --- | 495 |
| Manes.17G098900.2 | --- | 402 |
| Sobic.001G162400.1 | --- | 493 |
| Sobic.003G307600.2 | --- | 409 |
| Sobic.002G232200.1 | --- | 486 |
| Sobic.002G232500.1 | --- | 482 |
| Sobic.002G232600.1 | --- | 477 |
| Sobic.007G160700.1 | --- | 495 |
| Sobic.003G126200.2 | --- | 505 |
| Sobic.001G476700.1 | --- | 502 |
| Sobic.001G476700.2 | --- | 525 |
| Manes.18G062800.1 | --- | 496 |
| AT5G44050.1 | --- | 491 |
| AT5G10420.1 | --- | 489 |
| AT5G65380.1 | --- | 486 |
| Manes.12G129000.1 | --- | 487 |
| Manes.13G097900.1 | --- | 461 |
| Sobic.005G020700.1 | --- | 529 |
| Sobic.008G019400.1 | --- | 525 |
| Manes.09G135300.1 | --- | 486 |
| AT1G33080.1 | --- | 494 |
| AT1G33090.1 | --- | 494 |
| AT1G33100.1 | --- | 491 |
| AT1G33110.1 | --- | 494 |
| AT3G03620.1 | --- | 500 |
| AT5G17700.1 | --- | 497 |
| Manes.09G135200.1 | --- | 476 |
| Manes.09G134800.1 | --- | 472 |
| Manes.08G150400.1 | --- | 480 |
| Manes.09G134900.1 | --- | 487 |
| Manes.09G135000.1 | --- | 487 |
| Manes.09G134900.2 | --- | 458 |
| Manes.09G135000.2 | --- | 458 |
