## Supplementary Figures for "Genetic architecture and gene mapping of cyanide in cassava (*Manihot esculenta Crantz*.)"

The following Supplementary Information is available for this article:

**Supplementary Note 1:**

### Population structure analysis

Population stratification analysis was conducted on a larger population of 3354 individuals using Discriminant Analysis of Principal Components (DAPC) (Jombart, Devillard, and Balloux 2010)and parametric Admixture (Alexander, Novembre, and Lange 2009)with 5 folds cross validations for 20 K (assumed number of ancestral populations) to define the optimal number of clusters and examine patterns of relatedness and sub-ancestry among individuals in the Brazilian dataset as earlier described by Ogbonna et al., (in press).

Population structure for these materials have been extensively discussed in a parallel study by Ogbonna et al., (in press). Principal component analysis for 1,246 HCN assayed individuals with 9,686 SNPs (with Hardy-Weinberg and LD filtering) in the Brazilian germplasm shows structure patterns in our population with the first three PCs accounting for over 15.3% genetic variation.

**Supplementary Method 1:**

### Proportion of variance explained by markers

To determine the proportion of variance explained by the discovered loci for HCN, we used a parametric mixed model (SPMM) multiple kernel approach as previously described by Akdemir and Jannink(Akdemir and Jannink 2015). Briefly, the approach incorporates the marginal variance contribution from each kernel matrix.

**Supplementary Method 2:**

### Genome-wide epistasis interaction analysis

Genome-wide epistatic interactions between pairs of SNPs across the genome were carried out using FaST-LMM (Lippert et al. 2011, 2013), an approach that scales linearly with cohort size (sample size) in both run time and memory use. The method is based on maximum-likelihood estimates (REML for epistasis interactions) and accounts for the problem of confounding by population structure, family structure and cryptic relatedness (Widmer et al. 2014). The Bonferroni threshold was used to test for interactions that were significant and the observed -log10(p-value) was compared against the expected using the quantile-quantile plot.

**Supplementary Method 3:**

**Cultivated and cassava progenitor differentiating loci analysis**

To assess fixed or nearly fixed loci differentiating between cultivated (*M. esculenta*) and wild cassava (*M. flabellifolia*) to investigate if cassava domestication targeted upstream or downstream genetic regulation steps of cyanide bio-synthesis. We compared differentiating loci using method earlier described by (Bredeson et al. 2016; Wolfe et al. 2019; Ogbonna et al. 2020, in press) using Whole-Genome sequencing HapMap II dataset (Ramu et al. 2017) and contrasted groups (cultivated and progenitors) of 5 representative accessions each based on the phylogenetic tree from Ramu et al (Ramu et al. 2017), see **Supplementary Table 6**.

**Supplementary Method 4:**

**Kaspar Marker Design and Assessment**

Based on association peaks, local linkage disequilibrium and allelic effect on HCN content, 6 KASP SNP markers (Supplemental table 7) were designed from available genome sequences (v6.1) including positions. Flanking regions (100 bp) were extracted for each SNP and submitted for designability following the manufacturer's recommendation (LGC genomics, Malden, MA, USA).

**Supplementary Method 5:**

**Candidate gene protein Topology and Structure Prediction**

Transmembrane protein topology of our candidate gene (Manes.16G007900) was predicted using transmembrane hidden Markov model (TMHMM, https://services.healthtech.dtu.dk/service.php?TMHMM-2.0) according to the method described by (Krogh et al. 2001). The protein structure was modelled using the Phyre2 server (http://www.sbg.bio.ic.ac.uk/phyre2) following the procedure outlined in Kelley et al. (Kelley et al. 2015).Subsequent protein molecules fold stability changes (ddG) upon single point-mutations were predicted using STRUM (https://zhanglab.ccmb.med.umich.edu/STRUM/) following the approach outlined in Quan et al. (Quan, Lv, and Zhang 2016).

**Supplementary Method 6:**

**Single Point Mutation Prediction**

Allele Mining and mutation prediction in whole genome resequencing data for Manes.16G007900 and Manes.16G008000 proteins. STRUM (https://zhanglab.ccmb.med.umich.edu/STRUM/) was used for predicting the fold stability change (ΔΔG) of protein molecules upon single-point SNP mutations. STRUM adopts a gradient boosting regression approach to train the Gibbs free-energy changes on a variety of features at different levels of sequence and structure properties. Change in free energy (ranges between -5 to 5) below zero means that the mutation causes destabilization. Mutations with sensitive stability changes can affect the motion and fluctuation of the target residues.

Point Mutation Prediction: Prediction on the SNP mutation-induced stability changes is important to protein function annotation. Wild-type amino acids are usually more stable and adoptable to the protein environments due to the long-term evolution than the new mutations. The relatively uniform stability from the wide-type amino acids when compared to the identity of the mutated amino acids should provide more information with regard to the stability changes upon new mutations (Quan, Lv, and Zhang 2016).

**Supplementary Method 7:**

**Geographical Distribution of HCN**

Using Tess3 (Caye et al. 2016)and available georeferenced data for our GWAS dataset (Ogbonna et al, in press), we plotted the geospatial allele frequency distribution for the candidates associated with SNP on chromosomes 14 and 16, respectively. Individual ancestry coefficients, the proportions of an individual genome that originate from multiple ancestral gene pools, were estimated from their allelic frequency (Frichot et al. 2014).

**Supplementary Method 8:**

**GWAS in African Population and Joint Africa, Latin America Analysis**

To validate our findings in African cassava we sourced HCN trial experiments (228) from the cassava breeding database cassavabase.org. These trials were conducted by the International Institute for Tropical Agriculture (IITA) in West Africa between 1996 through 2017 across multiple locations (18) with 18,794 plots assayed for HCN. We carried out phenotypic and GWAS analysis as earlier described for Brazilian germplasm. GWAS analysis was performed on 636 unique individuals with phenotypic and genotypic information along with 53,547 SNPs. In addition, we performed joint phenotypic and GWAS analysis on African and Brazilian populations. The individuals with both phenotypic and genotypic information were 1,875 (Brazil, 1239; Africa, 636; Supplementary Table 15) and were used for the GWAS analysis along with 17,773 common SNP loci.

We finally performed a whole genome imputation of the African-Brazilian dataset using using beagle4.0 (Browning and Browning 2009)with 10 iterations, a window of 5000 markers and an overlap window of 500 markers and the HapMap as a reference panel for chromosome 16 and this dataset was used for GWAS analysis for cyanide.

**Supplementary Figure 1**

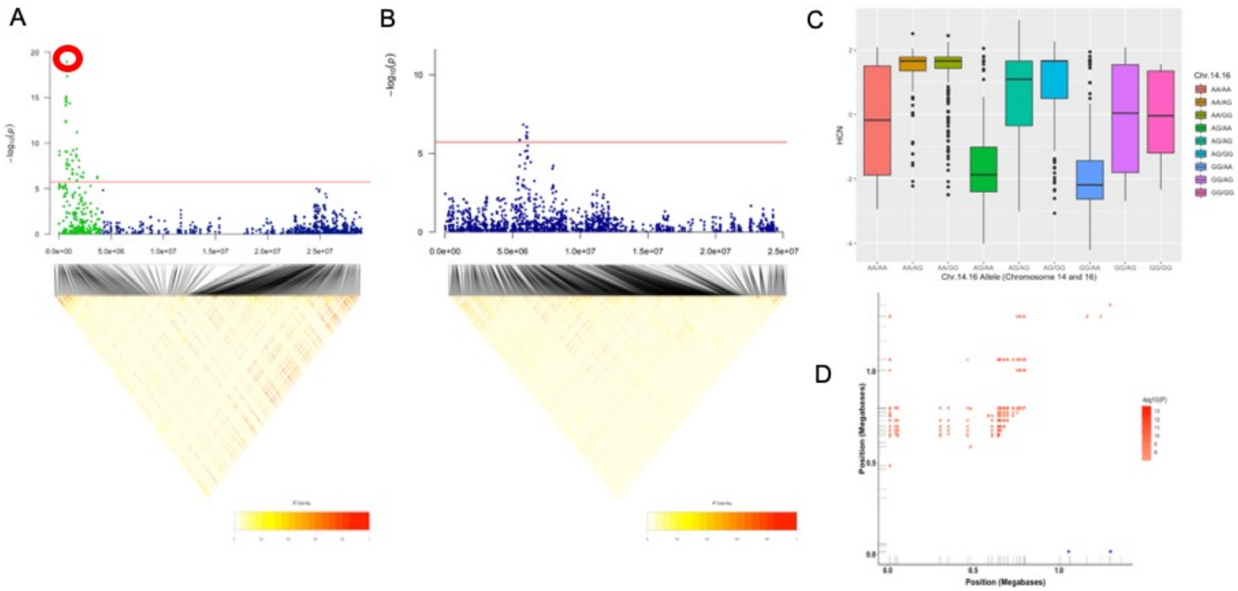


Manhattan plot from mixed linear model (MLM-LOCO) of chromosomes associated with cyanide variation in Latin American cassava. Below each Manhattan plot is a linkage disequilibrium (LD) heatmap for each chromosome showing pairwise squared correlation of alleles between markers. Bonferroni significance threshold is shown in red. **(A)** Manhattan plot of chromosome 16 showing candidate SNP for cyanide variation. The red circle indicates the candidate SNP. **(B)** Manhattan plot of chromosome 14 showing peak for cyanide variation. **(C)** Box plot showing the distribution of HCN for the combined effect (epistatic interaction) of the top significant markers for chromosome 16 and 14. HCN BLUP values were plotted on the Y-axis, while allelic effects of the candidate SNPs from chromosomes 14 and 16 combined on the X-axis. **(D)** Epistasis Interactions for HCN variation in LA germplasm. The upper triangle of the plot (red) represents 242 significant epistasis interactions (Bonferroni correction threshold, 0.05/1131*(1131-1)/2) for chromosome 16. The lower triangle of the plot (blue) shows the 3 separated interactions, with 2 of them overlapping at 1.286336 Megabases. The Y and X-axis of the plot are the positions in Megabases of the set of SNP1 and SNP2 interacting markers, respectively.

**Supplementary Figure 2.**

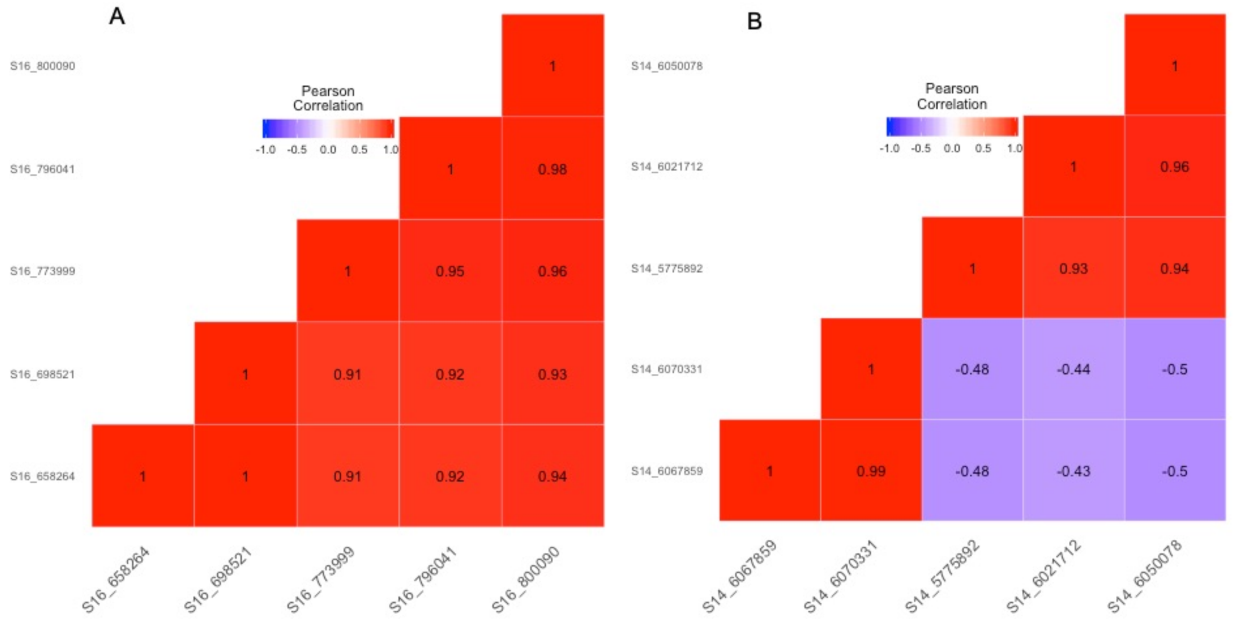


Pearson correlation using corrplot function in R package version 3.6.3 (2020-02-29) of top 5 significant SNPs in chromosome 16 (A) and 14 (B).

**Supplementary Figure 3.**

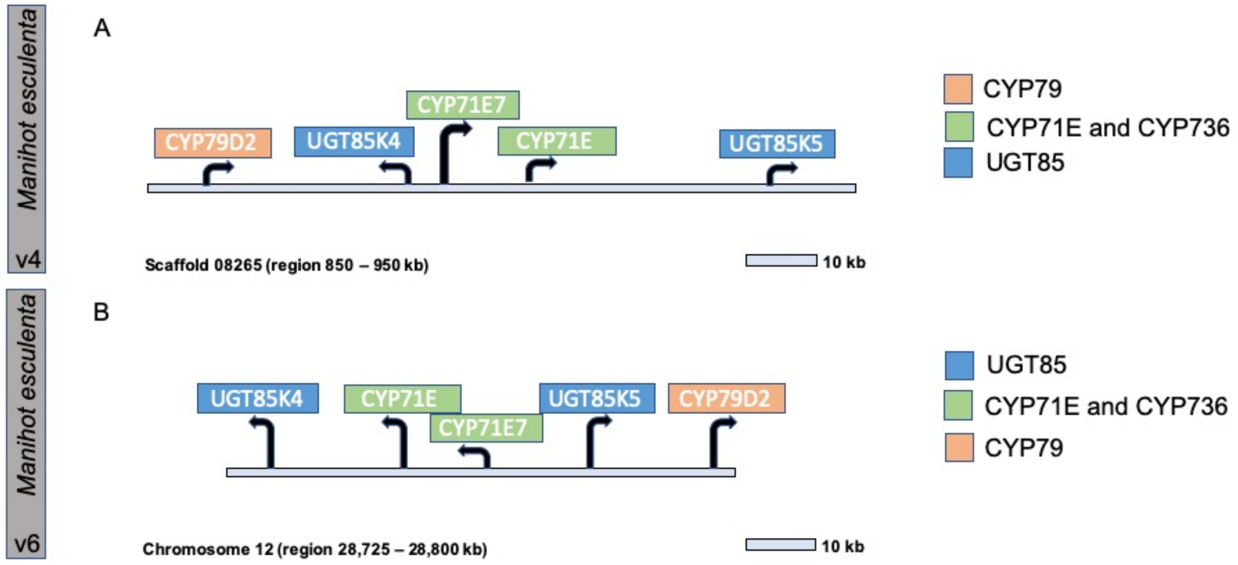


Schematic representation of the clustering of cyanogenic glucoside biosynthetic genes in the genome of *M. esculenta*. Functional genes are presented by arrows indicating their orientation. Confirmed genes in cyanogenic glucoside biosynthesis are labelled above each bar, with *CYP79* genes in pink, *CYP71E* and *CYP736* genes in green, and *UGT85* genes in blue. The sequences of the genes were retrieved and blasted against cassava genome version 6.1 on phytozome (http://phytozome.jgi.doe.gov). (A) Genome version Cassava4.1 (B) Genome draft version Cassava6.1. ***Adapted from Takoset al., the plant journal, 2011.***

**Supplementary Figure 4.**

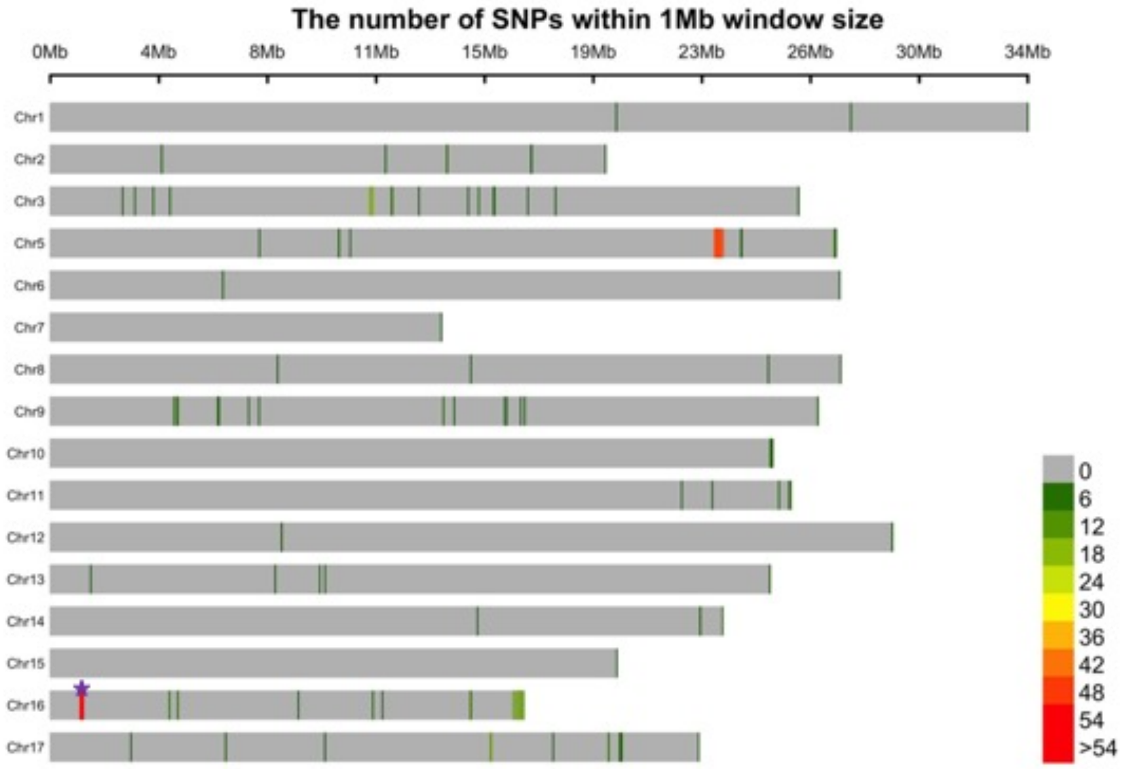


The plot shows the distribution of 294 biallelic ancestry-informative single-nucleotide markers that represent fixed, or nearly fixed, differences between *M. esculenta* and *M. flabellifolia in the hapmap II WGS dataset*. The legend scale indicates the number of SNPs within 1Mb window size. The purple star represents the identified region for cyanide regulation in cassava.

**Supplementary Figure 5.**

A

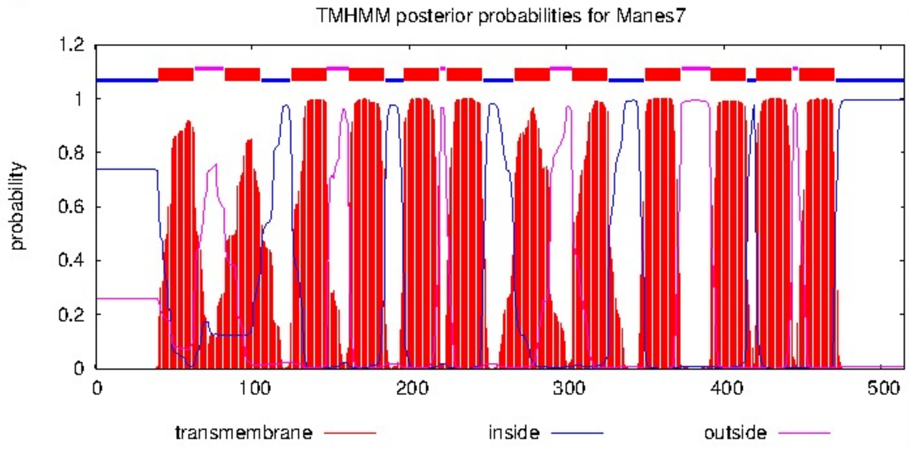


**TMHMM result**

```
# Manes7 Length: 515
# Manes7 Number of predicted TMHs:  12
# Manes7 Exp number of AAs in TMHs: 265.87223
# Manes7 Exp number, first 60 AAs:  13.97659
# Manes7 Total prob of N-in:        0.73944
# Manes7 POSSIBLE N-term signal sequence
Manes7  TMHMM2.0        inside       1    40
Manes7  TMHMM2.0        TMhelix     41    63
Manes7  TMHMM2.0        outside     64    82
Manes7  TMHMM2.0        TMhelix     83   105
Manes7  TMHMM2.0        inside     106   124
Manes7  TMHMM2.0        TMhelix    125   147
Manes7  TMHMM2.0        outside    148   161
Manes7  TMHMM2.0        TMhelix    162   184
Manes7  TMHMM2.0        inside     185   196
Manes7  TMHMM2.0        TMhelix    197   219
Manes7  TMHMM2.0        outside    220   223
Manes7  TMHMM2.0        TMhelix    224   246
Manes7  TMHMM2.0        inside     247   266
Manes7  TMHMM2.0        TMhelix    267   289
Manes7  TMHMM2.0        outside    290   303
Manes7  TMHMM2.0        TMhelix    304   326
Manes7  TMHMM2.0        inside     327   349
Manes7  TMHMM2.0        TMhelix    350   372
Manes7  TMHMM2.0        outside    373   391
Manes7  TMHMM2.0        TMhelix    392   414
Manes7  TMHMM2.0        inside     415   420
Manes7  TMHMM2.0        TMhelix    421   443
Manes7  TMHMM2.0        outside    444   447
Manes7  TMHMM2.0        TMhelix    448   470
Manes7  TMHMM2.0        inside     471   515
```

B

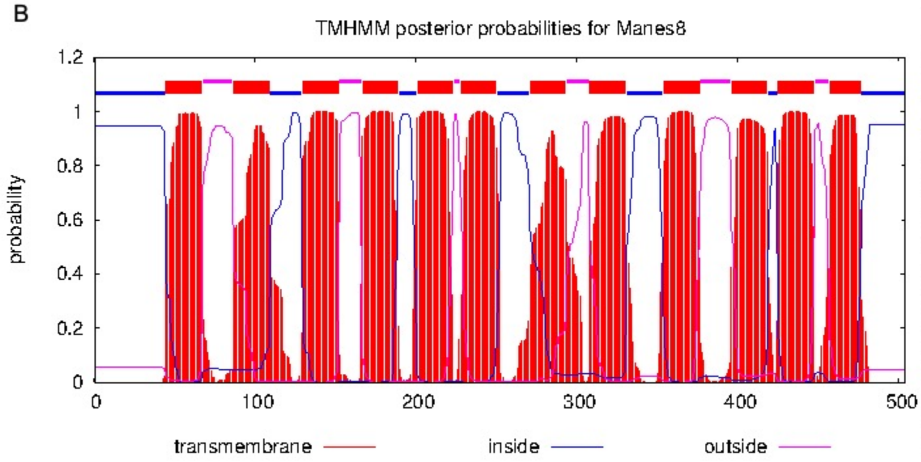


**TMHMM result**

```
# Manes8 Length: 504
# Manes8 Number of predicted TMHs:  12
# Manes8 Exp number of AAs in TMHs: 264.87498
# Manes8 Exp number, first 60 AAs:  14.37589
# Manes8 Total prob of N-in:        0.94595
# Manes8 POSSIBLE N-term signal sequence
Manes8  TMHMM2.0        inside       1    44
Manes8  TMHMM2.0        TMhelix     45    67
Manes8  TMHMM2.0        outside     68    86
Manes8  TMHMM2.0        TMhelix     87   109
Manes8  TMHMM2.0        inside     110   129
Manes8  TMHMM2.0        TMhelix    130   152
Manes8  TMHMM2.0        outside    153   166
Manes8  TMHMM2.0        TMhelix    167   189
Manes8  TMHMM2.0        inside     190   200
Manes8  TMHMM2.0        TMhelix    201   223
Manes8  TMHMM2.0        outside    224   227
Manes8  TMHMM2.0        TMhelix    228   250
Manes8  TMHMM2.0        inside     251   270
Manes8  TMHMM2.0        TMhelix    271   293
Manes8  TMHMM2.0        outside    294   307
Manes8  TMHMM2.0        TMhelix    308   330
Manes8  TMHMM2.0        inside     331   353
Manes8  TMHMM2.0        TMhelix    354   376
Manes8  TMHMM2.0        outside    377   395
Manes8  TMHMM2.0        TMhelix    396   418
Manes8  TMHMM2.0        inside     419   424
Manes8  TMHMM2.0        TMhelix    425   447
Manes8  TMHMM2.0        outside    448   456
Manes8  TMHMM2.0        TMhelix    457   476
Manes8  TMHMM2.0        inside     477   504
```

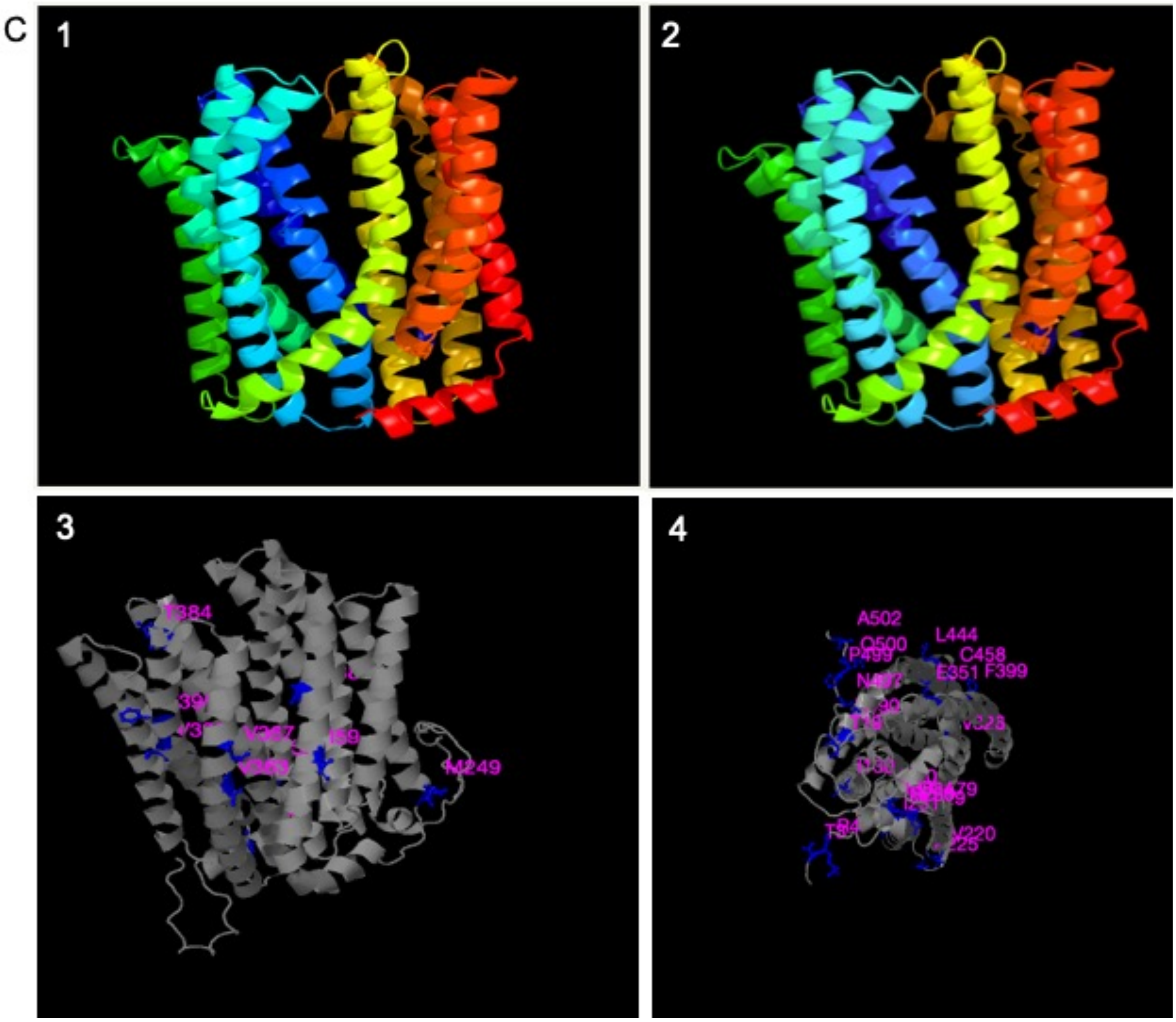


Transmembrane Helices prediction using Hidden Markov Model (TMHMM) posterior probability for transmembrane helix, inside, or outside displayed for the MATE transport family protein (Manes.16G007900.1 and Manes.16G008000.1) from cassava via (https://services.healthtech.dtu.dk/service.php?TMHMM-2.0). TMHMM is described in (Krogh et al. 2001), *Journal of Molecular Biology)*. **(A)** TMHMM results for Manes.16G007900.1 (here named Manes7), statistics and a list of the location of the predicted transmembrane helices and the predicted location of the intervening loop regions. Protein length: 515 amino acids, Number of predicted transmembrane (TM) helices: 12, Expected number of amino acids in TM helices: 265.87, Expected number of amino acids in TM helices in the first 60 amino acids of the protein: 13.98, Probability that the N-term is on the cytoplasmic side of the membrane: 0.74. The prediction gives the most probable location and orientation of transmembrane helices in the sequence. **(B)** TMHMM results for Manes.16G008000.1 protein (here named Manes8), statistics and a list of the location of the predicted transmembrane helices and the predicted location of the intervening loop regions. Protein length: 504, Number of predicted TM helices:

12, Expected number of amino acids in TM helices: 264.87, Expected number of amino acids in TM helices in the first 60 amino acids of the protein: 14.38, Probability that the N-term is on the cytoplasmic side of the membrane: 0.95. The prediction gives the most probable location and orientation of transmembrane helices in the sequence. **(C)** The protein structure was modelled using the Phyre2 server (http://www.sbg.bio.ic.ac.uk/phyre2) following the procedure outlined in Kelley *et al.* (2015). **(1)** Manes.16G007900: the model is based on the Crystal structure of eukaryotic MATE transporter AtDTX14(PDB ID: 5Y50A) with 100% confidence over 443 residues (36% of the sequence) (He et al. 2010). **(2)** Manes.16G008000**:** the model is based on the Crystal structure of eukaryotic MATE transporter AtDTX14(PDB ID: 5Y50A, (Miyauchi et al. 2017)) with 100% confidence over 446 residues (88% of the sequence) (He et al. 2010). **(3)** Single point mutation prediction positions for Manes.16G007900. **(4)** Single point mutation prediction positions for Manes.16G008000. STRUM (https://zhanglab.ccmb.med.umich.edu/STRUM/) was used to predict structural changes based on single point mutations.

**Supplementary Figure 6.**

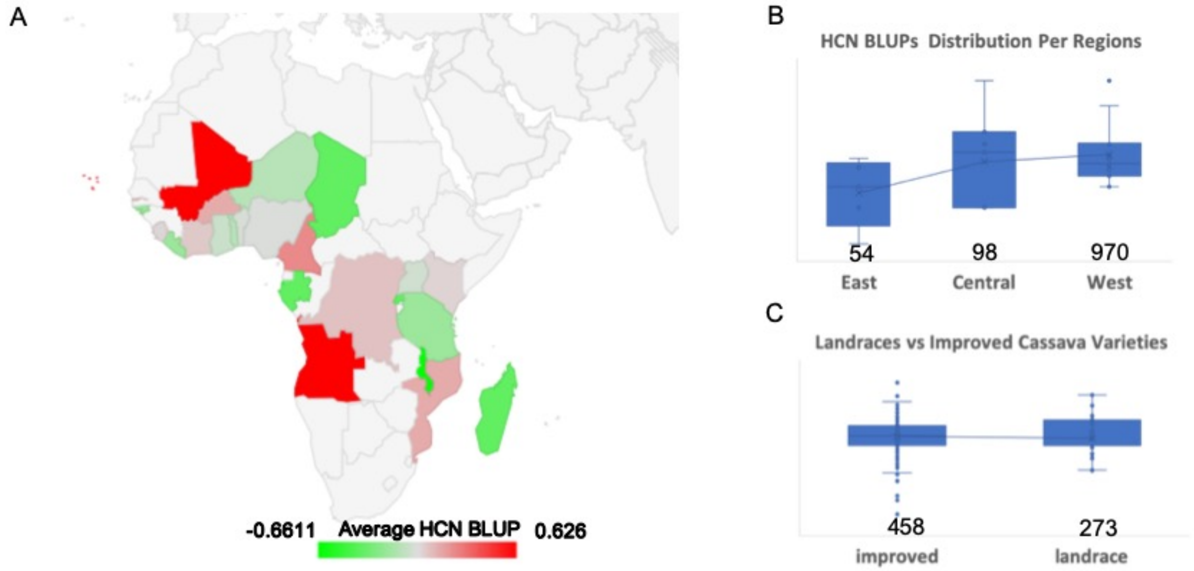


Cyanide (HCN) distribution by country and region for 1,156 accessions with country of origin in our African dataset (**Supplementary Table 11**). **(A)** Distribution of accessions based on average best linear unbiased prediction (BLUP) of HCN across countries in sub-Saharan Africa. Average HCN is higher in Central Africa where Konzo disease has prevailed. **(B)** Distribution of accessions based on average best linear unbiased prediction (BLUP) of HCN across regions of Africa, including data for 54, 98 and 970 accessions for East, Central and West Africa respectively. **(C) L**andraces vs improved accessions comparison including 458 improved and 273 landrace accessions.

**Supplementary Figure 7.**

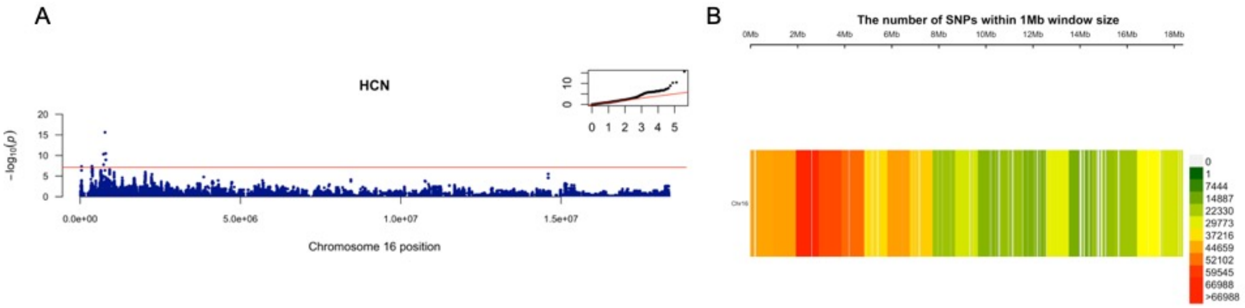


(A) Manhattan plot of Whole-Genome imputed chromosome 16 based on HapMapII (up to about 18 Megabases) from mixed linear model (MLM) using raw GBS from joint Latin American + African (LA+AF) germplasm. A total of 643,750 SNPs was imputed around 18 Megabases of chromosome 16 using 1877 individuals. The Bonferroni significance threshold is shown in red [7.109747, -log10(0.05/643750)]. A quantile-quantile plot is inserted to demonstrate the observed and expected -log10 of P-value for HCN. (B) Imputed single nucleotide polymorphism (SNP) density of joint Latin American + African (LA+AF) germplasm. The legend scale indicates the number of SNPs within 1Mb window size. The plot shows the distribution of SNPs across chromosome 16.

**Supplementary Figure 8.**

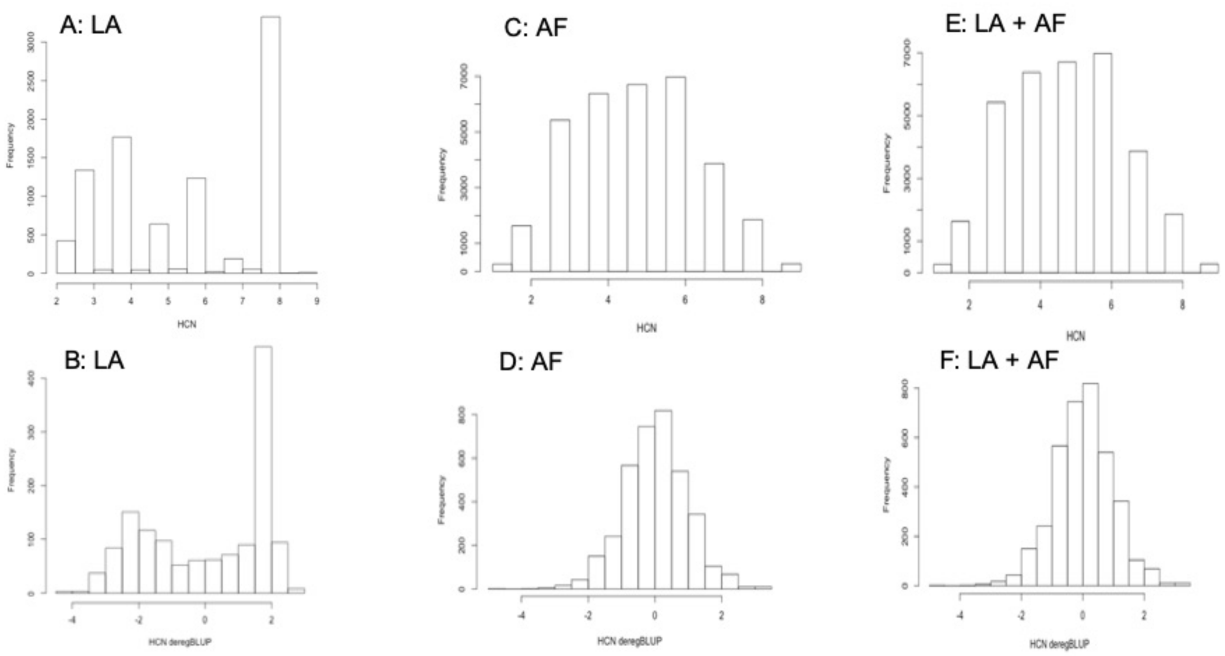


Distribution of HCN assayed for Latin American (LA, Brazilian), African (AF) and joint Latin American + African (LA+AF) germplasms. **(A)** Raw HCN phenotype scores for LA. **(B)** De-regressed HCN BLUPs for LA. **(C)** Raw HCN phenotype scores for AF. **(D)** De-regressed HCN BLUPs for AF. **(E)** Raw HCN phenotype scores for LA + AF. **(F)** De-regressed HCN BLUPs for LA + AF.

**Supplementary Figure 9.**

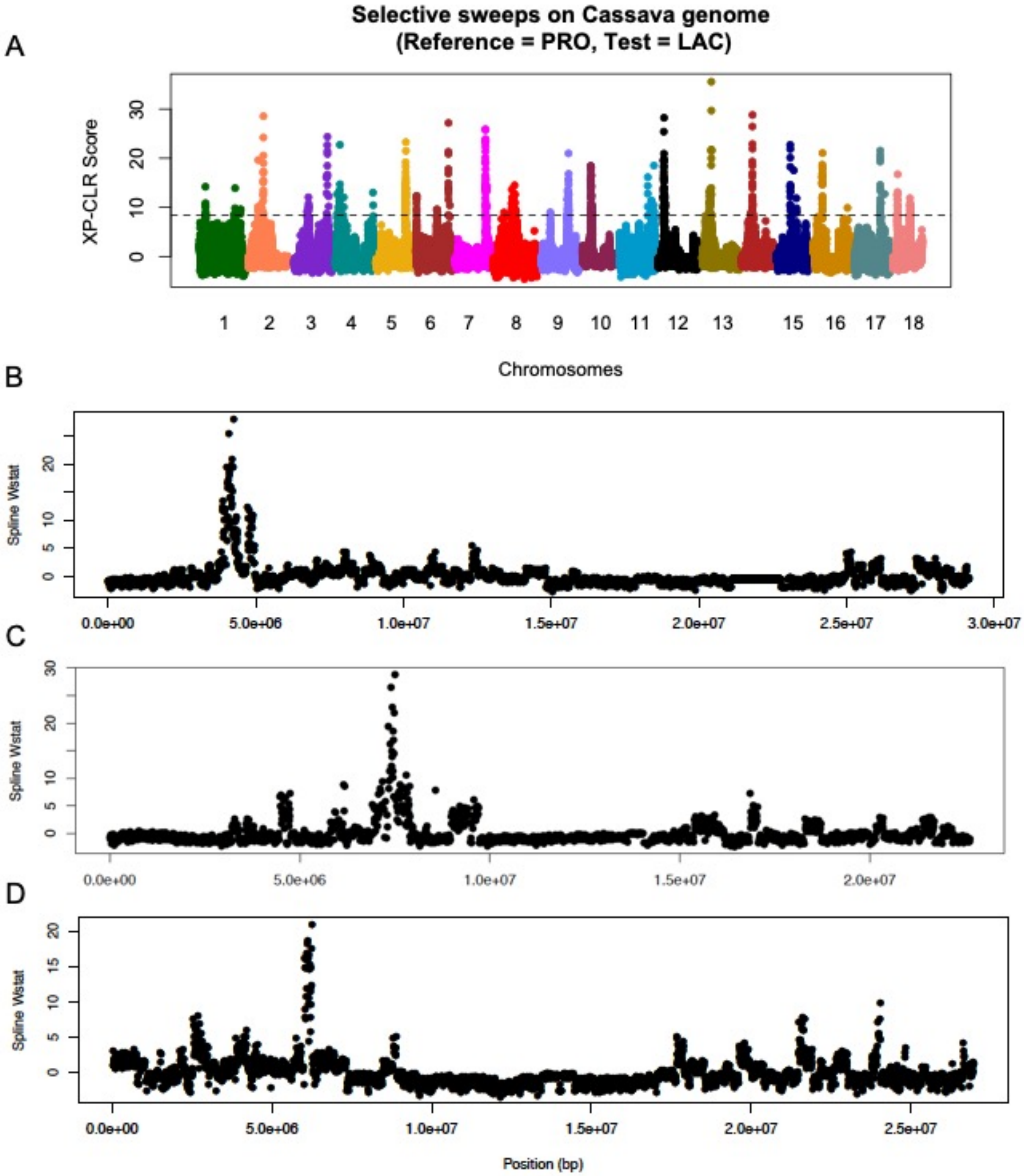


Selective sweep detection in Cassava HapMap II between Progenitors (PRO) and Latin American (LAC) accessions, following the approach described in Ramu et al (Ramu et al. 2017) (A) whole genome scan. (B) chromosome 12. (C) chromosome 14. (D) chromosome 16.

**Supplementary Figure 10.**

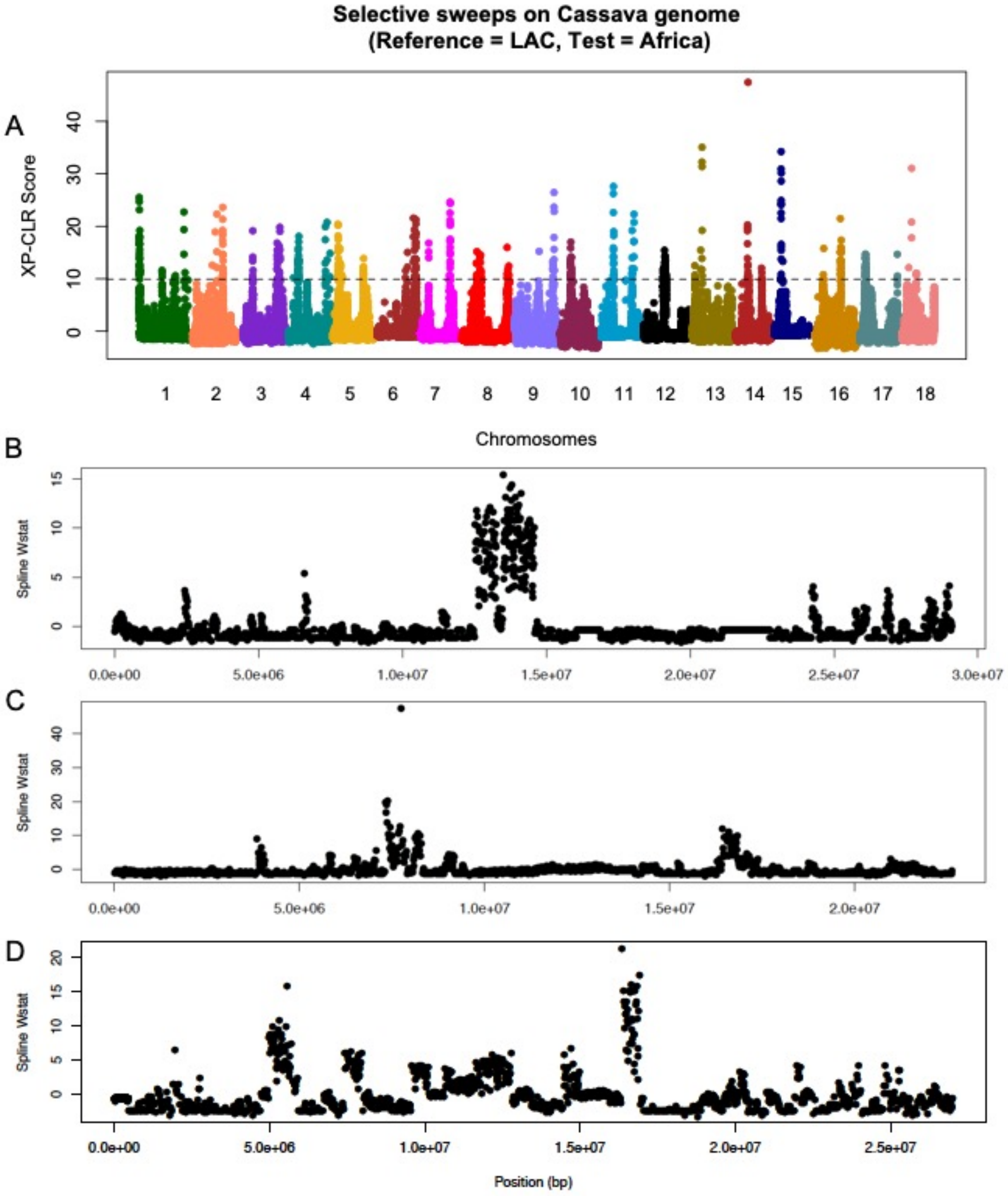


Selective sweep detection between Latin American (LAC) versus African accessions using the Cassava HapMap dataset following the approach described in Ramu et al (Ramu et al. 2017). (A) across the genome. (B) chromosome 12. (C) chromosome 14. (D) chromosome 16.

**Supplementary Figure 11.**

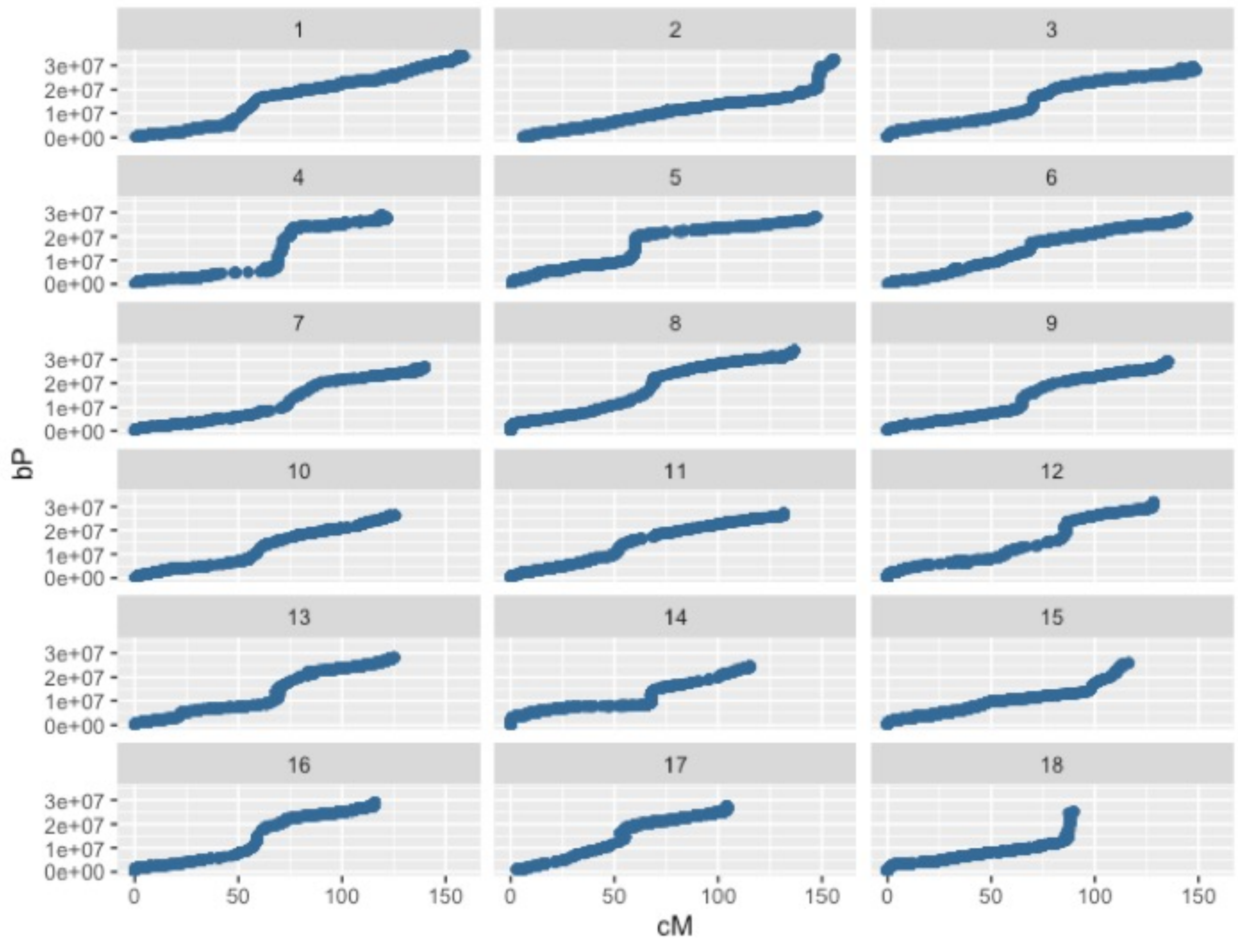


Projection of physical (bp) positions of Brazilian GBS marker set physical positions on updated International Cassava Genetic Map Consortium (ICGMC) linkage map based on method described in Wolfe et al (Wolfe et al. 2019) Scatterplots of genetic position vs. reference genome v6 physical positions for all 18 linkage groups for use in sweep detection analysis.

**Supplementary NOTE 2:**

**Phylogenetic tree**

Genome wide phylogenetic analysis of MATE genes in cassava, sorghum and arabidopsis suggested homology between our candidate, SbMATE2 and AT3G21690 genes. While SbMATE2 was discussed in the main text, additional discussion is provided for AT3G21690. SbMATE2 (Darbani et al. 2016) and AT3G21690 (Liu et al. 2009) are characterized as vacuolar membrane transporters in sorghum and Arabidopsis for cyanogenic glucoside, respectively. Koh and colleague (2010), suggested that, in suspension-cultured cells, overexpression of AT3G21690 affects the vacuolar accumulation of flavonoids or affects the biosynthesis of flavonoids by unknown mechanisms. In addition, a significant increase was observed in the accumulation of putative glucosinolates and sinapate derivatives in rosette leaves of the transgenic plants of Arabidopsis. Implying that a transporter encoded by AT3G21690 may transport multiple substrates according to the cell-type dependent metabolic activities (Koh et al. 2010). Manes.16G00800 showed closer sequence homology with AT1G61890 and AT1G11670. Within the same tree cluster, AT3G59030 is characterized as TT12 (Debeaujon et al. 2001; Marinova et al. 2007)and homologue in tobacco (Shoji et al. 2009).

**Supplementary NOTE 3:**

**Sweet and Bitter cassava geographical distribution (further discussion)**

Our finding was congruent with previous studies and presents new insights on cyanide spatial genetics. Clement and colleagues (2010), reported that bitter cassava cultivation was associated with the courses of the major Amazonian rivers, as well as the coastal areas of South America, where population densities were highest before conquest, while sweet cassava is the main crop throughout the headwaters of these same rivers in western Amazonia (Clement et al. 2010). In addition, Mckey and Beckerman (1993), reported that sweet cassava is commonly grown on a smaller scale where bitter cassava is the major crop. This may be due to the costs and benefits of toxicity, with greater benefits for large sedentary populations usually having semi-permanent fields, attracting greater pest and pathogen pressure. However, the cost is greater for smaller, more mobile populations (McKey, D. and Beckerman, S. 1993; Clement et al. 2010). Mckeyand Beckerman (1993), speculated that while these ideas may explain pre-conquest distributions, it is not clear if they explain current distributions of bitter and sweet cassava in Brazil. However, we found in our current dataset that the distribution of sweet and bitter cassava still reflects pre-conquest distribution and in addition, modern breeding activities in the last 40 years within Northern, Central and Southern regions of Brazil (Ogbonna et al, in press) had created a mixed population of low, intermediate and high cyanide varieties.

Jombart, Thibaut, Sébastien Devillard, and François Balloux. 2010. "Discriminant Analysis of Principal Components: A New Method for the Analysis of Genetically Structured Populations." *BMC Genetics* 11 (October): 94.

Kelley, Lawrence A., Stefans Mezulis, Christopher M. Yates, Mark N. Wass, and Michael J. E. Sternberg. 2015. "The Phyre2 Web Portal for Protein Modeling, Prediction and Analysis." *Nature Protocols* 10 (6): 845–58.

Krogh, A., B. Larsson, G. von Heijne, and E. L. Sonnhammer. 2001. "Predicting Transmembrane Protein Topology with a Hidden Markov Model: Application to Complete Genomes." *Journal of Molecular Biology* 305 (3): 567–80.

Lippert, Christoph, Jennifer Listgarten, Robert I. Davidson, Scott Baxter, Hoifung Poon, Carl M. Kadie, and David Heckerman. 2013. "An Exhaustive Epistatic SNP Association Analysis on Expanded Wellcome Trust Data." *Scientific Reports* 3 (January): 1099.

Lippert, Christoph, Jennifer Listgarten, Ying Liu, Carl M. Kadie, Robert I. Davidson, and David Heckerman. 2011. "FaST Linear Mixed Models for Genome-Wide Association Studies." *Nature Methods*. https://doi.org/10.1038/nmeth.1681.

Liu, Jiping, Jurandir V. Magalhaes, Jon Shaff, and Leon V. Kochian. 2009. "Aluminum-Activated Citrate and Malate Transporters from the MATE and ALMT Families Function Independently to Confer Arabidopsis Aluminum Tolerance." *The Plant Journal: For Cell and Molecular Biology* 57 (3): 389–99.

Marinova, Krasimira, Lucille Pourcel, Barbara Weder, Michael Schwarz, Denis Barron, Jean-Marc Routaboul, Isabelle Debeaujon, and Markus Klein. 2007. "The Arabidopsis MATE Transporter TT12 Acts as a Vacuolar flavonoid/H+ -Antiporter Active in Proanthocyanidin-Accumulating Cells of the Seed Coat." *The Plant Cell* 19 (6): 2023–38.

Miyauchi, Hirotake, Satomi Moriyama, Tsukasa Kusakizako, Kaoru Kumazaki, Takanori Nakane, Keitaro Yamashita, Kunio Hirata, et al. 2017. "Structural Basis for Xenobiotic Extrusion by Eukaryotic MATE Transporter." *Nature Communications* 8 (1): 1633.

Quan, Lijun, Qiang Lv, and Yang Zhang. 2016. “STRUM: Structure-Based Prediction of Protein Stability Changes upon Single-Point Mutation.” *Bioinformatics*. https://doi.org/10.1093/bioinformatics/btw361.

Ramu, Punna, Williams Esuma, Robert Kawuki, Ismail Y. Rabbi, Chiedozie Egesi, Jessen V. Bredeson, Rebecca S. Bart, Janu Verma, Edward S. Buckler, and Fei Lu. 2017. “Cassava Haplotype Map Highlights Fixation of Deleterious Mutations during Clonal Propagation.” *Nature Genetics* 49 (6): 959–63.

Shoji, Tsubasa, Koji Inai, Yoshiaki Yazaki, Yasutaka Sato, Hisabumi Takase, Nobukazu Shitan, Kazufumi Yazaki, et al. 2009. “Multidrug and Toxic Compound Extrusion-Type Transporters Implicated in Vacuolar Sequestration of Nicotine in Tobacco Roots.” *Plant Physiology* 149 (2): 708–18.

Widmer, Christian, Christoph Lippert, Omer Weissbrod, Nicolo Fusi, Carl Kadie, Robert Davidson, Jennifer Listgarten, and David Heckerman. 2014. “Further Improvements to Linear Mixed Models for Genome-Wide Association Studies.” *Scientific Reports* 4 (November): 6874.

Wolfe, Marnin D., Guillaume J. Bauchet, Ariel W. Chan, Roberto Lozano, Punna Ramu, Chiedozie Egesi, Robert Kawuki, Peter Kulakow, Ismail Rabbi, and Jean-Luc Jannink. 2019. “Historical Introgressions from a Wild Relative of Modern Cassava Improved Important Traits and May Be Under Balancing Selection.” *Genetics* 213 (4): 1237–53.
